# RE3DB: A multi-omics phylogenomics platform for rice E3 ubiquitin ligases identifies novel regulators of pollen germination

**DOI:** 10.1101/2025.09.18.676983

**Authors:** Wonjae Hwang, Ki-Hong Jung

**Affiliations:** Graduate School of Green-Bio Science and Crop Biotech Institute, Kyung Hee University, Yongin 17104, Republic of Korea; Research Centre for Plant Plasticity, Kyung Hee University, Seoul 08826, Republic of Korea

**Keywords:** E3 ubiquitin ligase, rice, database, multi-omics, phylogenomics, pollen germination

## Abstract

The ubiquitin proteasome system (UPS) shapes rice development and stress responses through E3 ubiquitin ligases, yet roles of E3 ligases in male gametophyte function remain undiscovered and plant-focused functional informatics resource is absent. We developed the Rice E3 Ubiquitin Ligase Database (RE3DB; https://re3db.khu.ac.kr/) and used it to nominate F-box ligases involved in rice (*Oryza sativa*) pollen germination. RE3DB combines 15 published datasets comprising 1,602 rice E3-ligase genes; RNA-seq across 25 tissues and stress conditions; protein-to-transcript ratios from 14 tissues; six high-confidence protein-protein interaction resources; and family level phylogenies. Integrated modules provide (i) classification/annotation, (ii) E3-substrate prediction from interaction evidence, co-expression and coordinated proteolytic turnover, and (iii) functional-redundancy assessment using phylogenetic heatmaps. Applying RE3DB, we found extensive reproductive enrichment: 284 F-box genes are preferentially expressed in mature anthers. Comparative transcriptomics across nine male-sterile rice mutants identified 111 F-box genes significantly downregulated in at least one mutant (log_2_ fold change < -1), resolving distinct co-regulation clusters. Integration with a literature-curated regulatory network positions previously uncharacterized F-box genes within modules governing pollen-tube initiation and elongation, consistent with regulation by the RUPO receptor-like kinase and the OsMADS62/63/68 transcription-factor complex. RE3DB is the first comprehensive multi-omics web platform dedicated to functional characterization of rice E3 ligases. By unifying classification, interaction and expression-based inference, and redundancy analysis, it accelerates hypothesis-driven discovery and guides targeted mutagenesis in molecular plant science. The prioritized F-box cohort provides testable candidates for UPS-mediated control of pollen germination and tube growth, and the framework is readily extensible to other crops, holding significant potential for agricultural biotechnology.

## Introduction

The ubiquitin-proteasome system (UPS) is a selective proteolytic system in which cellular proteins targeted for degradation by ATP-dependent 26S proteasomes are labeled with ubiquitin (Nandi *et al*., 2006; Kwon & Ciechanover, 2017). Ubiquitination involves three sequential enzymatic steps: ubiquitin activation by E1, transfer to E2 conjugating enzymes, and substrate-specific ligation via E3 ubiquitin ligases, which recognize target proteins and catalyze ubiquitin attachment to lysine residues (Sadanandom *et al*., 2012).

E3 ubiquitin ligases serve as essential specificity determinants within the UPS, with recent studies revealing their multifaceted regulatory functions in rice (*Oryza sativa*). The E3-mediated protein ubiquitination significantly influences plant development, environmental adaptations, hormone signaling pathways, and stress response mechanisms (Craig *et al*., 2009; Shu & Yang, 2017; Kelley, 2018). In developmental regulation, E3 ligases control critical transitions such as flowering time. The RING-type E3 ligase HAF1 (Heading date Associated Factor 1) ubiquitinates the flowering regulator Hd1 (Heading Date 1) and promotes its 26S proteasome-mediated degradation (Yang *et al*., 2015). By reducing daytime Hd1, when Hd1 represses a florigen homolog Hd3a even under short days, HAF1 alleviates this light-dependent repression and thereby promotes earlier flowering (Yang *et al*., 2015). Environmental adaptability in rice involves specialized E3 ligases; the cytosolic U-box E3 ubiquitin ligase OsPUB15 (*Oryza sativa* Plant U-box 15) limits reactive oxygen species (ROS) damage during seedling establishment, such that knockout seedlings are rootless and accumulate H_2_O_2_ and oxidized proteins, whereas overexpression lowers H_2_O_2_/oxidized proteins and enhances tolerance to salinity, oxidative stress, and drought (Park *et al*., 2011). In contrast, the drought-induced RING E3 ligase OsDIS1 (*Oryza sativa* drought-induced SINA protein 1) promotes 26S proteasome-dependent degradation of the tubulin-complex kinase OsNek6 (*Oryza sativa* NIMA-related kinase 6) and broadly reprograms drought-responsive transcription, so that overexpression reduces drought tolerance whereas RNAi silencing enhances it (Ning *et al*., 2011). E3-mediated ubiquitination also modulates hormone signaling and development as demonstrated by the RING E3 ligase OsDSG1 (*Oryza sativa* Delayed Seed Germination 1) which physically associates with the abscisic acid (ABA) regulator OsABI3 (*Oryza sativa* ABA Insensitive 3) and its loss-of-function elevates ABA-signaling outputs while suppressing GA-dependent hydrolase genes, resulting in delayed seed germination (Park *et al*., 2010). In stress response mechanisms, RING-H2 ubiquitin E3 ligase OsHTAS (Oryza sativa HEAT TOLERANCE AT SEEDLING STAGE) enhances rice seedling thermotolerance by promoting H_2_O_2_ induced stomatal closure through modulation of ROS accumulation and stimulation of ABA biosynthesis; OsHTAS participates in both the ABA-dependent pathway and the DROUGHT AND SALT TOLERANCE (DST)-mediated, ABA-independent route, thereby reducing water loss during heat stress (Liu *et al*., 2016).

Despite the critical importance of E3 ligases in plant biology, there remains no publicly available web resource that supports functional characterization of plant E3 ligases. Earlier efforts were confined to basic cataloging of the UPS genes rather than enabling functional analysis: plantsUPS provided annotations across seven plant genomes but is no longer publicly accessible, and iUUCD 2.0 broadened coverage to 39 plant species yet likewise focused on genome-wide identification. Most publicly available E3 ligase web resources are restricted to human E3 ligases with studies focused on discovering new drug targets and targeted therapies (Bielskienė *et al*., 2015; Sampson *et al*., 2023). For instance, the human E3 Ubiquitin Ligases database is an online resource of 377 human E3 ubiquitin ligases, predominantly HECT, RING, and U-box proteins, created to identify the E3 ligases most likely responsible for Aquaporin-2 (AQP2) ubiquitination using Bayes’ theorem (Medvar *et al*., 2016). Additionally, databases like UbiHub visualize biological, structural, and chemical data on phylogenetic trees of human protein families involved in ubiquitination signaling, aiding in target prioritization and drug design (Liu *et al*., 2019). UbiNet 2.0, which is currently inaccessible, offers detailed information on 3332 experimentally verified E3-substrate interactions across 54 organisms, with 73% of these interactions originating from human data, providing enriched annotations and interactive tools for visualizing ubiquitination networks (Li *et al*., 2021). Similarly, UbiBrowser 2.0 includes comprehensive data on predicted and known E3/deubiquitinase-substrate interactions across 39 non-plant organisms, with a confidence scoring system and enhanced web interface functionalities (Wang *et al*., 2022a).

The lack of web-based functional analysis resources on plant E3 ligase poses a significant gap in rice agricultural research, particularly because reproductive success and thus grain yield depends on efficient pollen hydration, germination, and sustained pollen tube growth that enables double fertilization; genetic disruption of key pollen-tube regulators causes near-complete male sterility with defects in hydration, germination, and elongation (Kim *et al*., 2023). A direct link to ubiquitin signaling is exemplified by *POLLEN TUBE BLOCKED 1* (*PTB1*), a RING-type E3 ligase in rice that promotes pollen-tube growth and elevates the panicle seed-setting rate; in 32 nucleotide mRNA deletion mutant *ptb1*, pollen tube growth is blocked in the style transmission tract and female-specific sterility results despite a morphologically normal female apparatus (Li *et al*., 2013). At the systems level, proteome-wide ubiquitinome profiling of young panicles mapped 1,638 lysine-ubiquitination sites on 916 proteins, including pollen and grain development, implying a critical role of UPS in the physiological functions of the reproductive tissues (Zhu *et al*., 2020). Yet discovering the function and role of the UPS in rice pollen maturation and germination remain incompletely resolved a network scale; out of 1,515 genes encoding E3 ligases, only 69 have been functionally characterized and the E3s for most ubiquitinated proteins are still unknown (Wang *et al*., 2022b).

Here, we present the Rice E3 Ubiquitin Ligase Database (RE3DB; https://re3db.khu.ac.kr/), the first comprehensive, multi-omics, web-based platform dedicated to the study of E3 ligases in rice. RE3DB provides an integrated analytics platform for (i) functional classification, (ii) E3-substrate prediction, and (iii) functional redundancy analysis of rice E3 ligases. It incorporates data from 15 published datasets identifying 1,602 E3 ligase genes, RNA-Seq data across 25 tissues and various stress conditions, protein-to-transcript ratios (PTRs) from 14 tissues, six high-confidence protein-protein interaction (PPI) datasets, and phylogenetic information from representative protein sequences. The database offers user-interactive visualization tools including word clouds, dynamic tables, network graphs, and phylogenetic heatmaps which significantly enhance data interpretation and experimental design capabilities for researchers investigating ubiquitin-mediated protein degradation in rice. As a proof-of-concept, we applied RE3DB to identify novel F-box genes involved in pollen germination in rice. Through tissue-specific expression profiling, over-representation analysis, differential gene expression analysis, and comparative transcriptome analysis across nine male-sterile mutant lines, we identified co-regulated F-box genes, predicted their upstream regulators, and positioned them within key rice pollen tube germination pathway modules. These results provide new insights into UPS-mediated regulation of male fertility and demonstrate the capability of RE3DB for extensive functional discoveries.

## Results

### Overview of RE3DB construction and functionality

RE3DB comprises three interactive modules: (i) the classification and annotation module; (ii) the functional redundancy assessment module; and (iii) the E3-substrate prediction module. To implement these modules, we unified multi-omics resources including genomic, transcriptomic and proteomic data obtained from public repositories and peer-reviewed studies (Fig. 1). For the classification and annotation module, gene lists for each E3-ligase family were gathered from 15 independent publications. Only loci with valid Rice Genome Annotation Project (RGAP, MSU7) identifiers were retained, and their representative protein sequences were retrieved. These sequences were re-annotated against the Pfam and InterPro databases to ensure consistent domain assignments. For the E3-substrate prediction module, PPI datasets from six published rice resources were merged. Interactions were accepted only when they exceeded the high confidence thresholds specified by the original studies. Transcript abundance was quantified from RNA-seq datasets covering anatomical series, abiotic and biotic stresses, hormone treatments and nutrient deprivation. Raw reads were downloaded from the NCBI SRA and EMBL-EBI ArrayExpress, processed into log_2_ normalized read counts (Table S1; Table S2; Table S3; Table S4; Table S5). Pearson correlation coefficients (PCC) and associated P-values were calculated across 25 rice tissues to derive co-expression networks. Additionally, we analyzed PTRs across 14 Nipponbare tissues to identify genes exhibiting low protein abundance relative to their transcript levels, indicative of post-transcriptional regulation. For the phylogenetic module, multiple-sequence alignments were generated for each E3 ligase family and maximum-likelihood trees were constructed. Anatomical expression profiles with PCC values and *p*-values were then mapped onto the trees to create combined phylogenetic heatmaps, facilitating visual inspection of redundant or diverged paralogues.

**Fig. 1.**
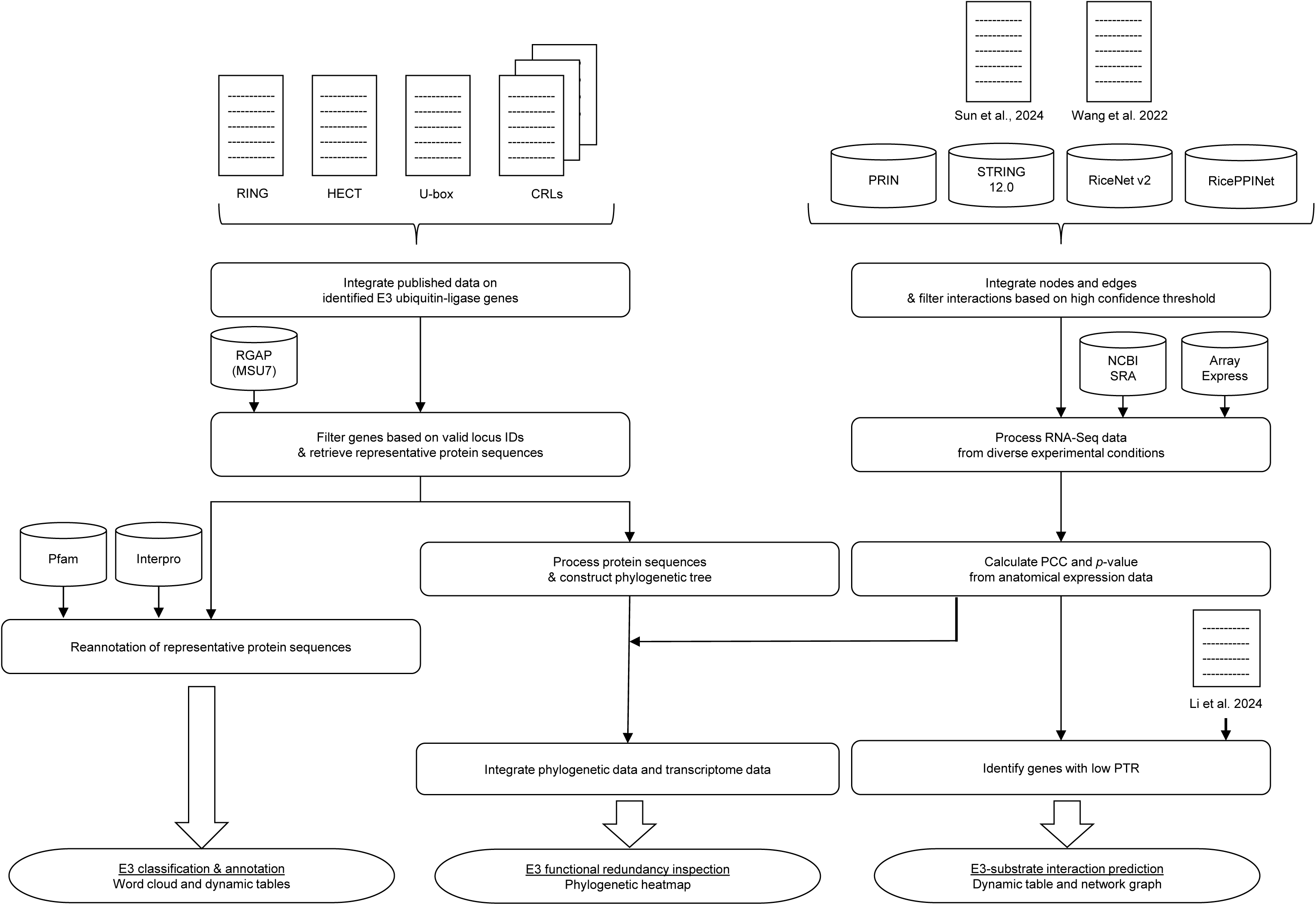
Overview of RE3DB Construction and Functionality. RE3DB integrates multi-omics resources to support three interactive modules: (i) classification and annotation; (ii) functional redundancy assessment; and (iii) E3–substrate prediction. Rice E3 ubiquitin ligase genes were collated from 15 publications; loci carrying valid Rice Genome Annotation Project identifiers (RGAP; MSU7) were retained and representative proteins re-annotated against Pfam and InterPro to harmonize domain assignments. Protein-protein interaction (PPI) datasets from six rice resources were merged; only interactions meeting the original studies’ high-confidence thresholds were included. RNA sequencing (RNA-seq) reads from the NCBI Sequence Read Archive (SRA) and EMBL-EBI ArrayExpress spanning anatomical series, abiotic/biotic stresses, hormone treatments, and nutrient deprivation were processed to log□ normalized read counts. Pearson correlation coefficients (PCC) across 25 rice tissues were used to construct co-expression networks; two-sided p-values associated with PCC are reported. Protein-to-transcript ratios (PTRs) across 14 tissues of Nipponbare (Oryza sativa ssp. japonica) highlight candidates with low protein abundance relative to transcript levels, consistent with post-transcriptional regulation. For each E3 family, multiple-sequence alignments and maximum-likelihood (ML) trees were generated; anatomical expression profiles are mapped as heatmaps onto phylogenies to visualize putative redundant versus diverged paralogues. PPI, protein–protein interaction; SRA, Sequence Read Archive; PCC, Pearson correlation coefficient; PTR, protein-to-transcript ratio; RGAP (MSU7), Rice Genome Annotation Project release 7; RNA-seq, RNA sequencing; ML, maximum-likelihood.

### Comprehensive classification and annotation of rice E3 ligases provided by the E3 ligase word cloud and search query

RE3DB enables in-depth exploration of all 1,602 rice E3 ligase genes by classifying genes into different families and presenting their distribution in an interactive word cloud. In this visualization, the font size of each family name is proportional to the number of genes it contains: APC (18 genes), BTB (160 genes), CDC20 (3 genes), Cullin (13 genes), DDB1 (1 genes), DWD (77 genes), F-box (729 genes), HECT (8 genes), RBX1 (2 genes), RING (483 genes), SKP1 (30 genes), TAD1 (1 genes), and U-box (77 genes) (Table 1). Selecting a family in the word cloud opens an interactive table that provides detailed annotations for every gene in that group. The rice gene OsBBI1 (BLAST AND BTH-INDUCED 1; LOC_Os06g03580), highlighted with a red square, encodes a 261-amino-acid RING-H2 E3 ligase (Fig. 2). Functional studies show that OsBBI1 positively regulates broad-spectrum resistance to rice blast: Tos17 insertion mutants display increased susceptibility to several *Magnaporthe oryzae* races, whereas 35S::OsBBI1 over-expression lines exhibit significantly smaller lesions and lower disease scores, presumably through enhanced reactive-oxygen-species-mediated cell-wall fortification and alterations in cell-wall defense responses (Li *et al*., 2011). Classification and annotation of individual genes can also be retrieved by their locus identifier using a search query in RE3DB.

**Fig. 2.**
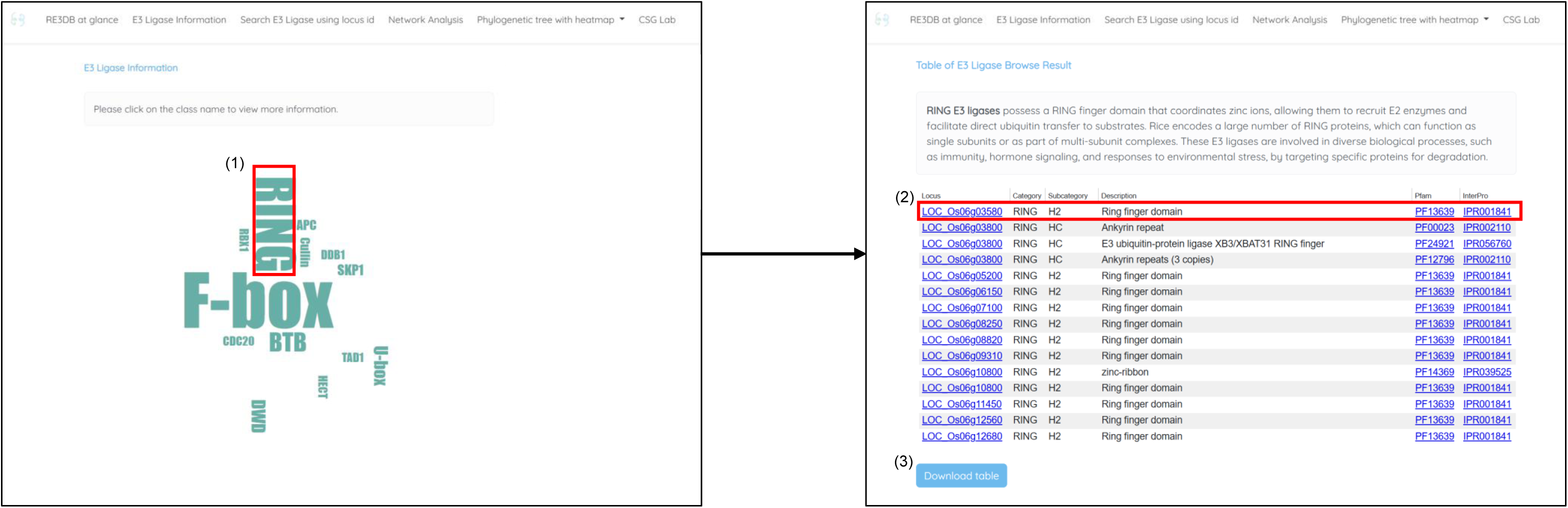
Comprehensive classification and annotation of rice E3 ubiquitin ligases in RE3DB. Word-cloud overview of 1,602 rice E3 ligase genes in which font size is proportional to family size. (1) When users click the E3 ligase family that they are interested in, (2) The family-specific interactive result table displays detailed information such as the RGAP (MSU7) locus id, E3 family category and subcategory, description based on Pfam annotation, and Pfam and InterPro entry. Clicking on to the hyperlinked locus id leads to RGAP locus information page, clicking on Pfam or InterPro entry leads to Pfam or Interpro information page. As an example, the rice gene OsBBI1 (BLAST AND BTH-INDUCED 1; LOC_Os06g03580) is marked by a red square. (3) At the bottom left corner, the sky-blue colored download button allows users to download the result table into a csv file.

**Table 1.**
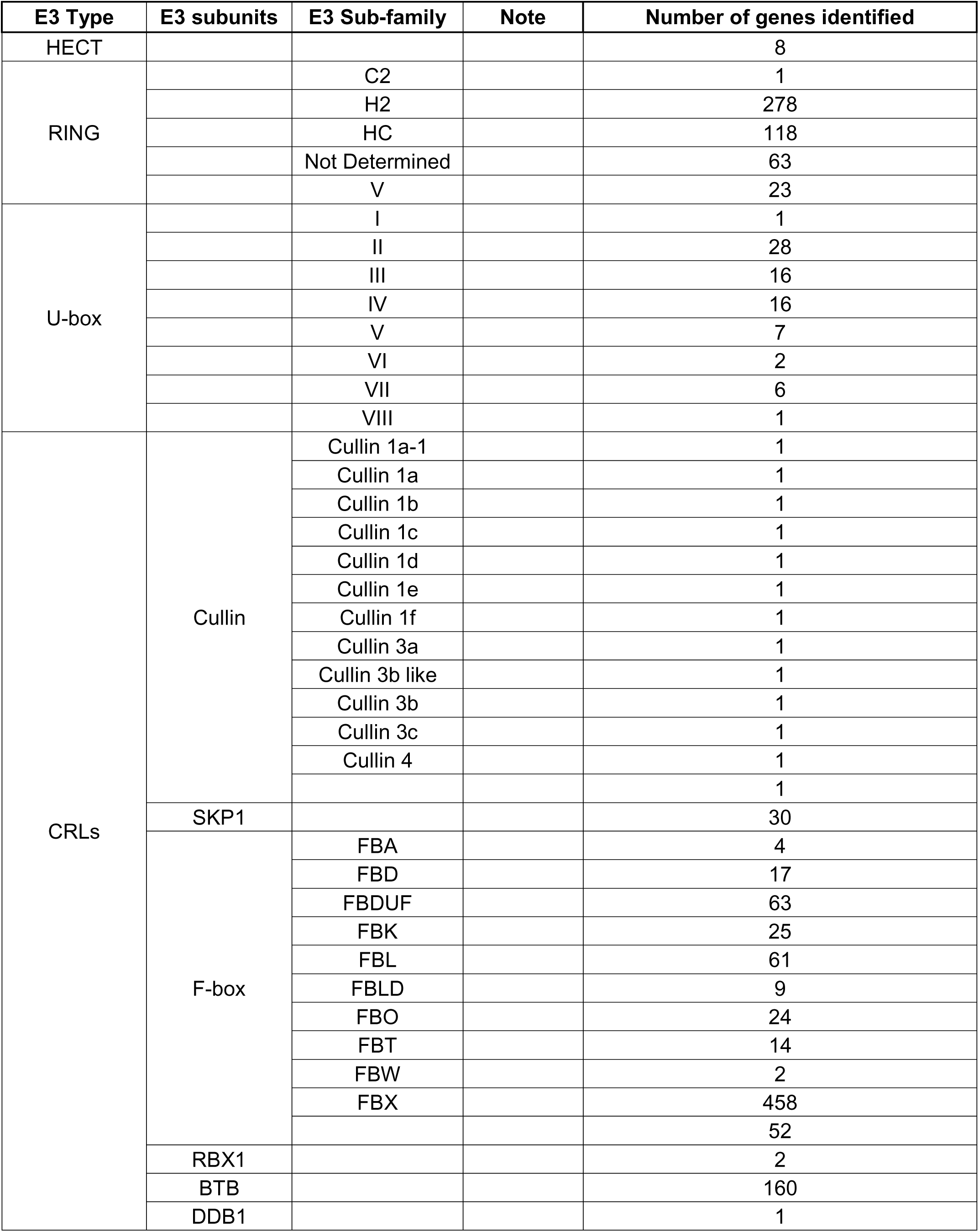
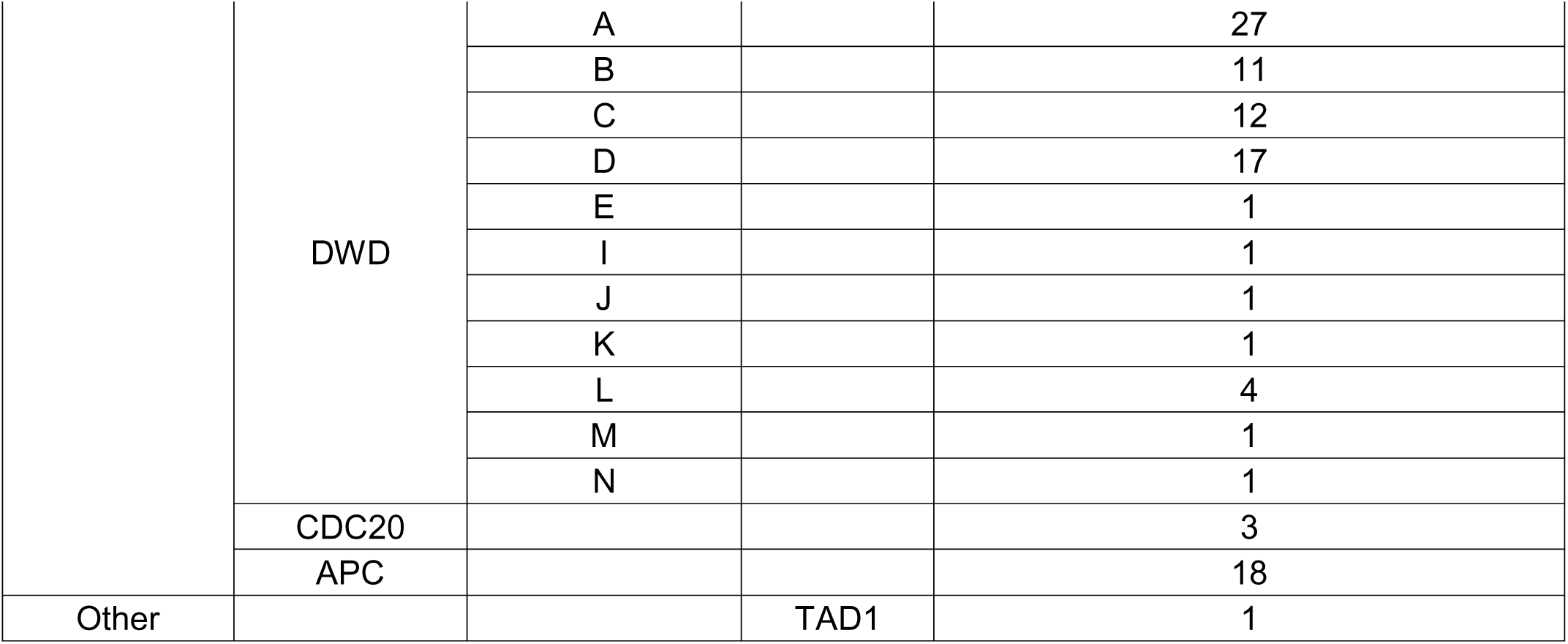
Categories and number of E3 ubiquitin ligase genes identified in RE3DB. HECT, Homologous to E6AP C-terminus; RING, Really interesting new gene; CRL, Cullin-RING E3 ubiquitin ligase; RBX1, RING box protein 1; BTB, Broad-complex, Tramtrack, and Bric-à-brac; DWD, DDB1 binding WD40; CDC20, Cell division cycle 20; APC, Anaphase-promoting complex.

| E3 Type | E3 subunits | E3 Sub-family | Note | Number of genes identified |
| --- | --- | --- | --- | --- |
| HECT |  |  |  | 8 |
| RING |  | C2 |  | 1 |
|  |  | H2 |  | 278 |
|  |  | HC |  | 118 |
|  |  | Not Determined |  | 63 |
|  |  | V |  | 23 |
| U-box |  | I |  | 1 |
|  |  | II |  | 28 |
|  |  | III |  | 16 |
|  |  | IV |  | 16 |
|  |  | V |  | 7 |
|  |  | VI |  | 2 |
|  |  | VII |  | 6 |
|  |  | VIII |  | 1 |
| CRLs | Cullin | Cullin 1a-1 |  | 1 |
|  |  | Cullin 1a |  | 1 |
|  |  | Cullin 1b |  | 1 |
|  |  | Cullin 1c |  | 1 |
|  |  | Cullin 1d |  | 1 |
|  |  | Cullin 1e |  | 1 |
|  |  | Cullin 1f |  | 1 |
|  |  | Cullin 3a |  | 1 |
|  |  | Cullin 3b like |  | 1 |
|  |  | Cullin 3b |  | 1 |
|  |  | Cullin 3c |  | 1 |
|  |  | Cullin 4 |  | 1 |
|  |  |  |  | 1 |
|  | SKP1 |  |  | 30 |
|  | F-box | FBA |  | 4 |
|  |  | FBD |  | 17 |
|  |  | FBDUF |  | 63 |
|  |  | FBK |  | 25 |
|  |  | FBL |  | 61 |
|  |  | FBLD |  | 9 |
|  |  | FBO |  | 24 |
|  |  | FBT |  | 14 |
|  |  | FBW |  | 2 |
|  |  | FBX |  | 458 |
|  |  |  |  | 52 |
|  | RBX1 |  |  | 2 |
|  | BTB |  |  | 160 |
|  | DDB1 |  |  | 1 |
|  | DWD | A |  | 27 |
|  |  | B |  | 11 |
|  |  | C |  | 12 |
|  |  | D |  | 17 |
|  |  | E |  | 1 |
|  |  | I |  | 1 |
|  |  | J |  | 1 |
|  |  | K |  | 1 |
|  |  | L |  | 4 |
|  |  | M |  | 1 |
|  |  | N |  | 1 |
|  | CDC20 |  |  | 3 |
|  | APC |  |  | 18 |
| Other |  |  | TAD1 | 1 |

### Phylogenetic heatmaps in RE3DB for assessing functional redundancy among rice E3 ubiquitin ligase families

The functional redundancy assessment module is designed to address gene redundancy within each E3 ligase family. It displays phylogenetic heatmaps that overlay protein sequence similarity with gene expression patterns across 25 tissues and under various conditions, including abiotic stress, biotic stress, hormone deprivation, and nutrient starvation. Users can generate a phylogenetic heatmap through an individual gene search query or for selecting an entire E3 ubiquitin ligase family. When using the gene search query, RE3DB also indicates the PCC and corresponding *p*-value for genes within the phylogenetic tree. This integrated approach allows researchers to identify co-expressed paralogs that may have compensatory functions, as well as unique family members with non-redundant expression profiles. Such information facilitates more precise and effective targeted mutagenesis strategies using gene-editing tools like CRISPR/Cas9. For example, the module clearly visualizes the functional redundancy between the F-box genes OsCOI1a (CORONATINE INSENSITIVE1a; LOC_Os01g63420) and OsCOI1b (LOC_Os05g37690) (Fig. 3). Consistent with the heatmap prediction, functional studies demonstrated that by generating CRISPR/Cas9 single knock-outs and two independent double mutants (oscoi1a-1/b-1 and oscoi1a-1/b-2), only the double mutants displayed distinctive traits, taller plants with significantly elongated tillers and internodes, whereas each single mutant was phenotypically similar to wild type, confirming their overlapping functions in shoot development (Nguyen *et al*., 2023).

**Fig. 3.**
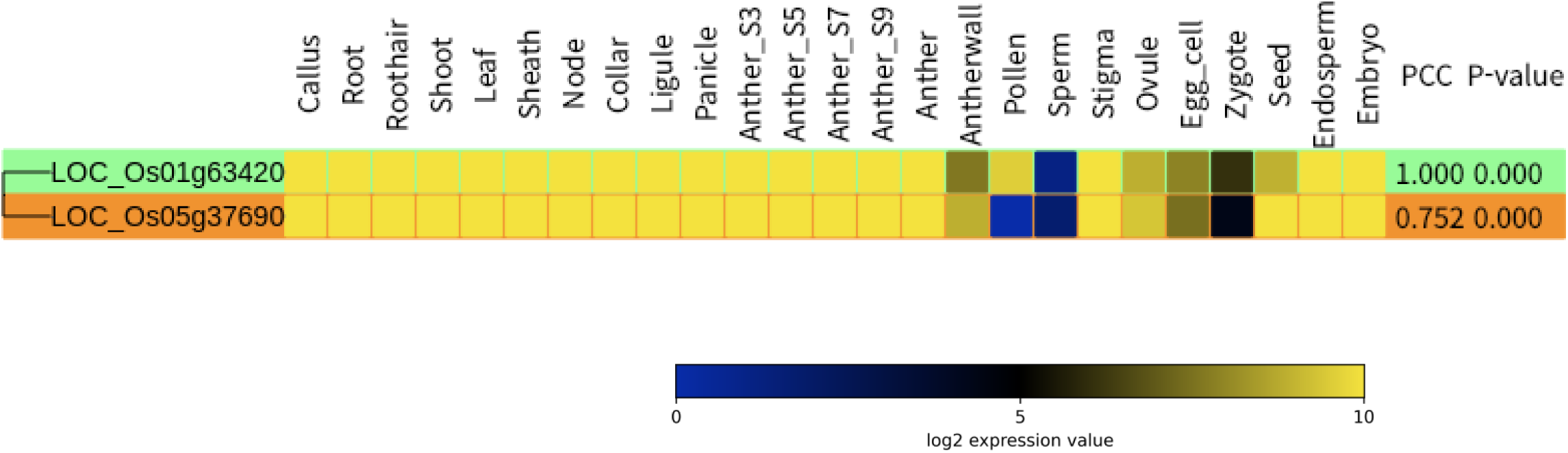
Phylogenetic heatmaps in RE3DB reveal redundant and divergent genes of rice E3 ubiquitin ligase families. For a selected E3 family, or a single gene query, RE3DB builds a maximum-likelihood protein phylogeny and overlays a heatmap of transcript abundance across 25 rice tissues and diverse conditions such as abiotic/biotic stresses, hormone and nutrient treatments. The color gradient indicates expression level in linear color scale. As for single gene query, each node also displays the Pearson correlation coefficient (PCC) for co-expression analysis along with the corresponding p-value. The prediction of the F-box paralogs OsCOI1a (CORONATINE INSENSITIVE1a; LOC_Os01g63420) and OsCOI1b (LOC_Os05g37690) have been validated through previous functional studies. PCC, Pearson correlation coefficient.

### Integrated multi-omics approach for prediction of E3-substrate interactions in RE3DB

The E3-substrate prediction module identifies high-confidence E3–substrate pairs by integrating four key criteria: (1) high-confidence protein-protein interaction (PPI) prediction networks based on diverse approaches including interolog mapping, protein structure docking, experimental high-throughput screening, machine learning with structural and functional features, network-based algorithms, and integration of multiple evidence types including co-expression and experimental data, (2) transcriptional co-expression across 25 tissues measured by Pearson correlation coefficients (PCC), (3) statistical significance filtering based on *p*-values, and (4) coordinated proteolytic turnover, inferred from low PTRs across 14 tissues. The result highlights potential E3-substrate pairs that are co-expressed within the same tissues and exhibit coordinated proteolytic dynamics. For example, querying the F-box protein OsFBK16 (F-box Kelch repeat protein 16; LOC_Os06g39370) returns a table and a network graph of predicted substrates, including experimentally validated interactors such as OsPAL1 (phenylalanine ammonia-lyase gene 1; LOC_Os02g41630) and OsPAL2 (LOC_Os02g41650) (Fig. 4). Along with yeast two-hybrid screens in which OsPAL1 as bait yielded 111 of 124 positive clones and OsPAL2 as bait yielded 93 of 105 positive clones encoding OsFBK16, the OsPAL1-OsFBK16 interaction was further confirmed by yeast co-transformation and by co-immunoprecipitation in *Nicotiana benthamiana*; over-expression of OsPAL1 displayed enhanced resistance to *Magnaporthe oryzae*, indicating that OsFBK16-mediated turnover of PALs negatively regulates rice blast immunity (Wang *et al*., 2022b).

**Fig. 4.**
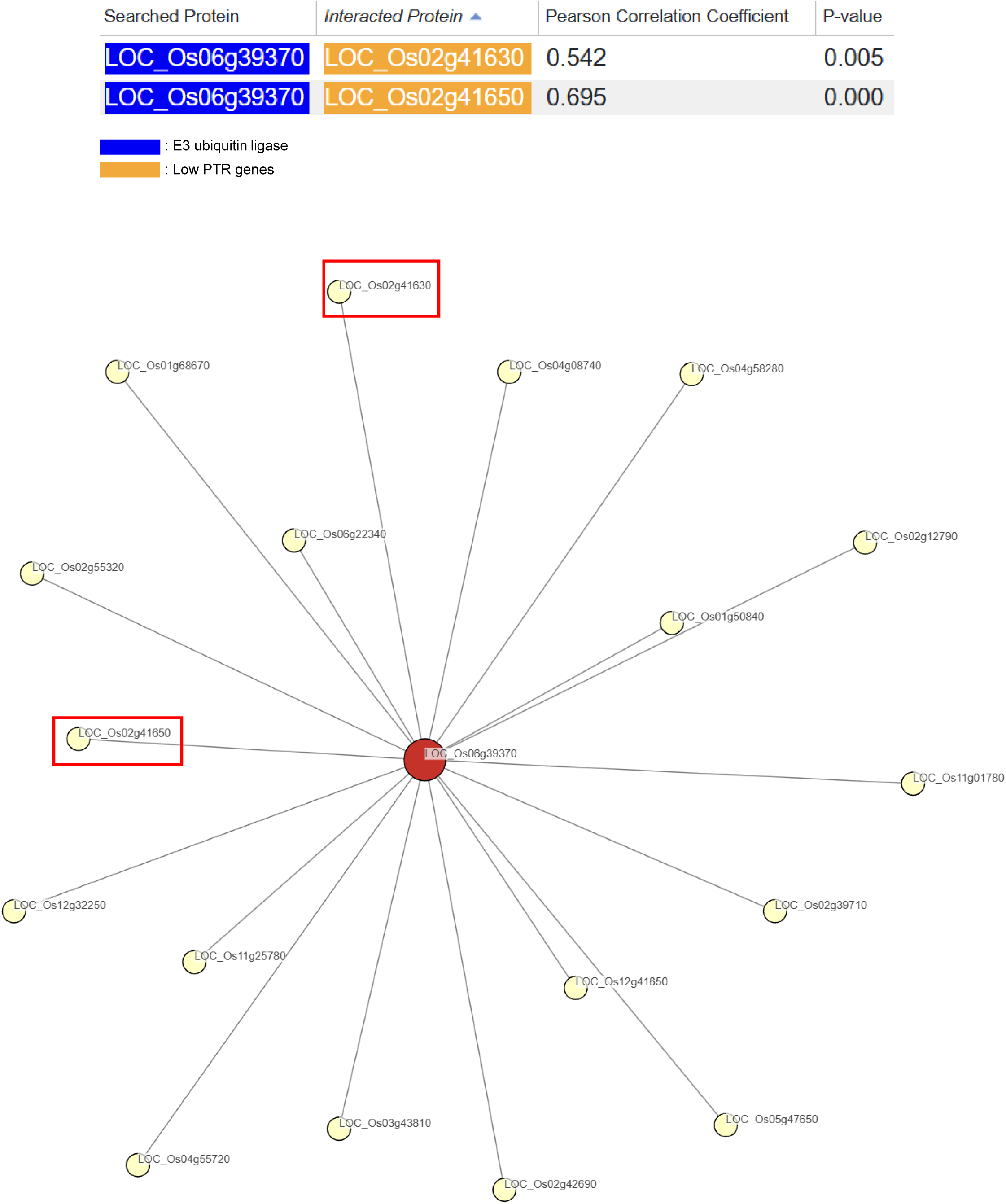
Integrated multi-omics workflow in RE3DB for predicting E3-substrate interactions. E3-substrate interactions are predicted by combining four evidence layers: high-confidence protein-protein interaction (PPI) resources, transcript co-expression across 25 tissues, statistical filtering by p-value, and proteolysis inferred from low protein-to-transcript ratios (PTRs) across 14 tissues. The result of an example query OsFBK16 (F-box Kelch repeat protein 16; LOC_Os06g39370) returns a table and network of predicted substrates including OsPAL1 (phenylalanine ammonia-lyase gene 1; LOC_Os02g41630) and OsPAL2 (LOC_Os02g41650) which has been validated through experimental studies. PPI, protein-protein interaction; PTR, protein-to-transcript ratio.

### Global identification and transcriptome analysis of E3 ligase encoding genes

Transcriptome profiling across 25 rice tissues and organs enabled the systematic identification of E3 ubiquitin-ligase-encoding genes with tissue- or organ-preferential expression. The result is organized into nested hierarchical groups based on developmental traits: (i) the vegetative body, subdivided into root system, shoot system (further partitioned into shoot and leaf tissues, node, and collar region), and callus; (ii) the inflorescence and floral sporophyte, encompassing panicle, anther developmental series (early and mature phases), and gynoecium organs; (iii) male (subdivided into pollen and sperm); and (iv) female gametophytic lineages (subdivided into egg cell and post-fertilization); and (v) seed development, which includes embryo, endosperm, and seed (Fig. 5; Fig. S1). A gene was considered tissue-preferential if at least one tissue in the corresponding group exhibited a Z-score ≥ 2, while no tissues outside the group reached this threshold (Table S6; Table S7; Fig. S2; Fig. S3; Fig. S4; Fig. S5; Fig. S6; Fig. S7; Fig. S8; Fig. S9; Fig. S10; Fig. S11)

**Fig. 5.**
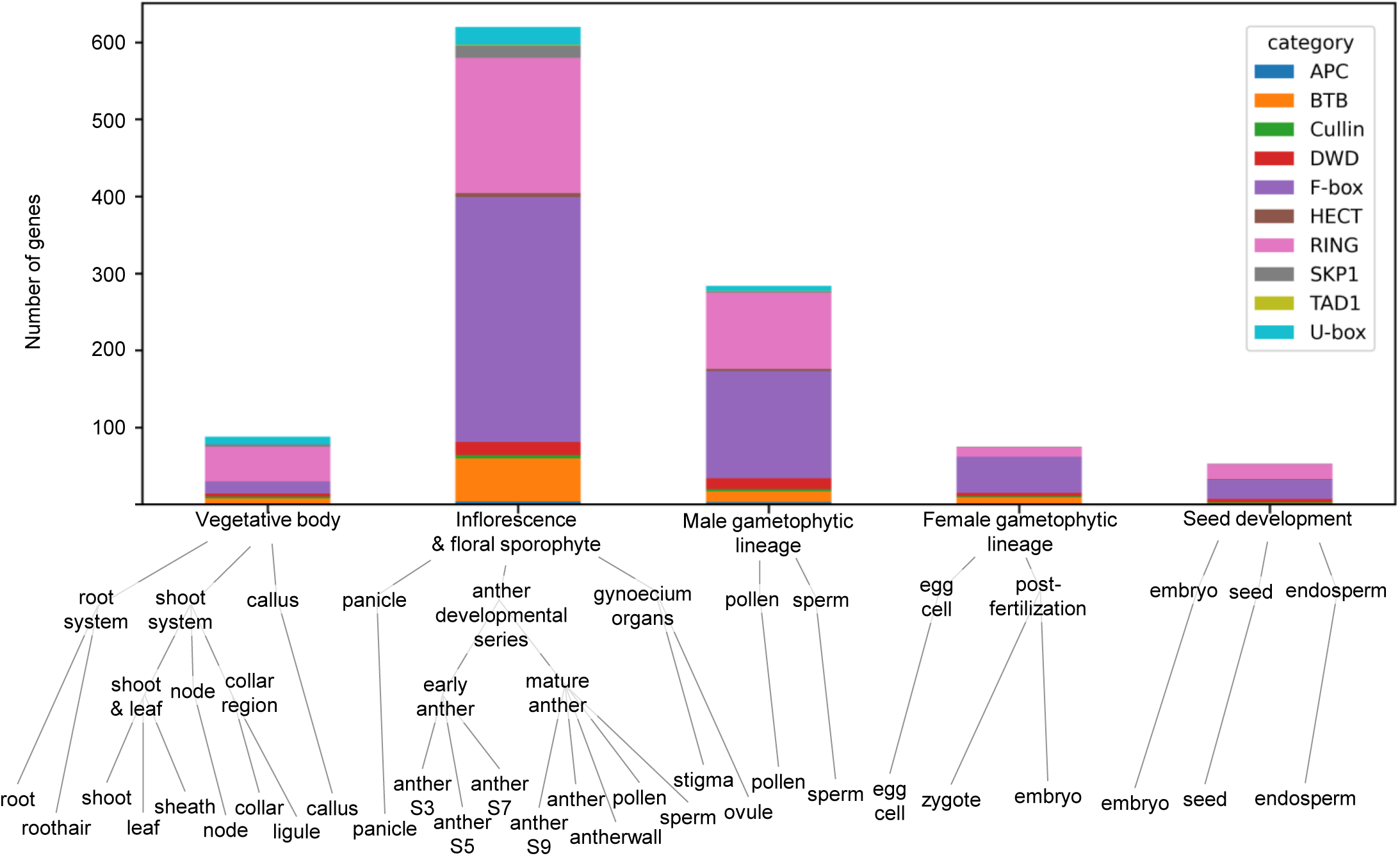
Global identification of rice E3 ubiquitin-ligase genes with tissue-preferential expression. Anatomical hierarchy used for transcriptome profiling of E3 genes across 25 rice tissues: vegetative body (root system; shoot system subdivided into shoot and leaf, node, and collar; callus), inflorescence and floral sporophyte (panicle; anthers at early and mature stages; gynoecium), male lineage (pollen, sperm), female lineage (egg cell, post-fertilization), and seed development (embryo, endosperm, seed). The tissue-preferential E3s are defined as genes with Z ≥ 2 in one or more tissues within a given group and no tissues outside that group reaching Z ≥ 2.

### Enrichment and functional profiling of E3 ligase families

An over-representation analysis (hypergeometric test), with false-discovery-rate (FDR) adjusted *p*-value ≤ 0.05 applied, was performed separately for each E3-ligase family (Table S8). For every family, the five annotation terms with the highest odds ratios and a FDR adjusted *p*-value ≤ 0.05 are displayed in dot-plots and further summarized in a bar plot (Fig. 6; Fig. S12). The result revealed notable enrichments of E3 ligase genes; (i) F-box genes in inflorescence and floral sporophyte (318 genes), specifically sperm (73 genes) and mature anther (284 genes) within the anther developmental series (304 genes), as well as female gametophytic lineage (47 genes), (ii) RING genes in sperm (45 genes) and pollen (47 genes) within the male gametophytic lineage (100 genes) and mature anther (165 genes), and (iii) U-box genes in root system (10 genes). Furthermore, based on the enriched gene list, we also investigated the list of functionally characterized tissue preferential E3 ligase genes (Table S9).

**Fig. 6.**
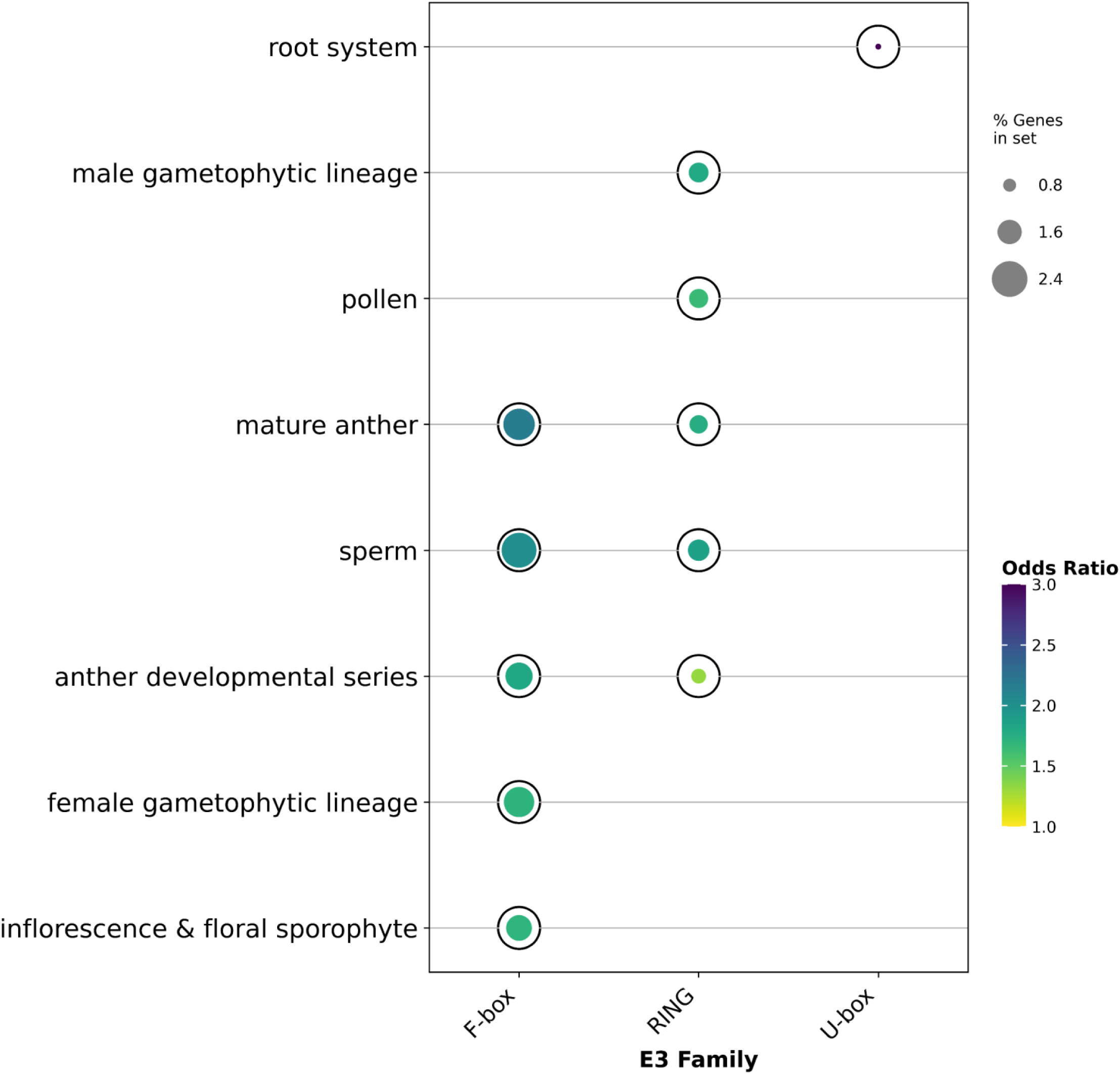
Enrichment result of rice E3-ligase families. For each E3 family, the top five annotation terms ranked by odds ratio that passes false-discovery-rate (FDR) adjusted p-value ≤ 0.05 are shown as dot plots. Dot size reflects the percentage of genes annotated to the term, and dot color shows the odds ratio. FDR, false-discovery-rate.

### Identification of mature anther preferential F-box E3 ligases in rice pollen germination pathways

In our previous study, we established a genetic resource comprising nine male-sterile rice mutant lines (*rupo*, *madstri*, *osmtd2*, *gori*, *ralf17/19*, *tape*, *osrac6*, *abcg16/28*, and *gtrd5*) by combining high-throughput screening of T-DNA insertion lines with CRISPR/Cas9-based gene editing (Kim *et al*., 2025a). For the 284 highly enriched mature anther preferential F-box genes, we conducted comparative RNA-seq analysis across the nine mutant lines and identified 111 downregulated differentially expressed genes (DEGs) in at least one mutant (log□ fold change < –1 compared to wild type). The regulation patterns were analyzed relative to wild-type controls (Table 2).

**Table 2.**
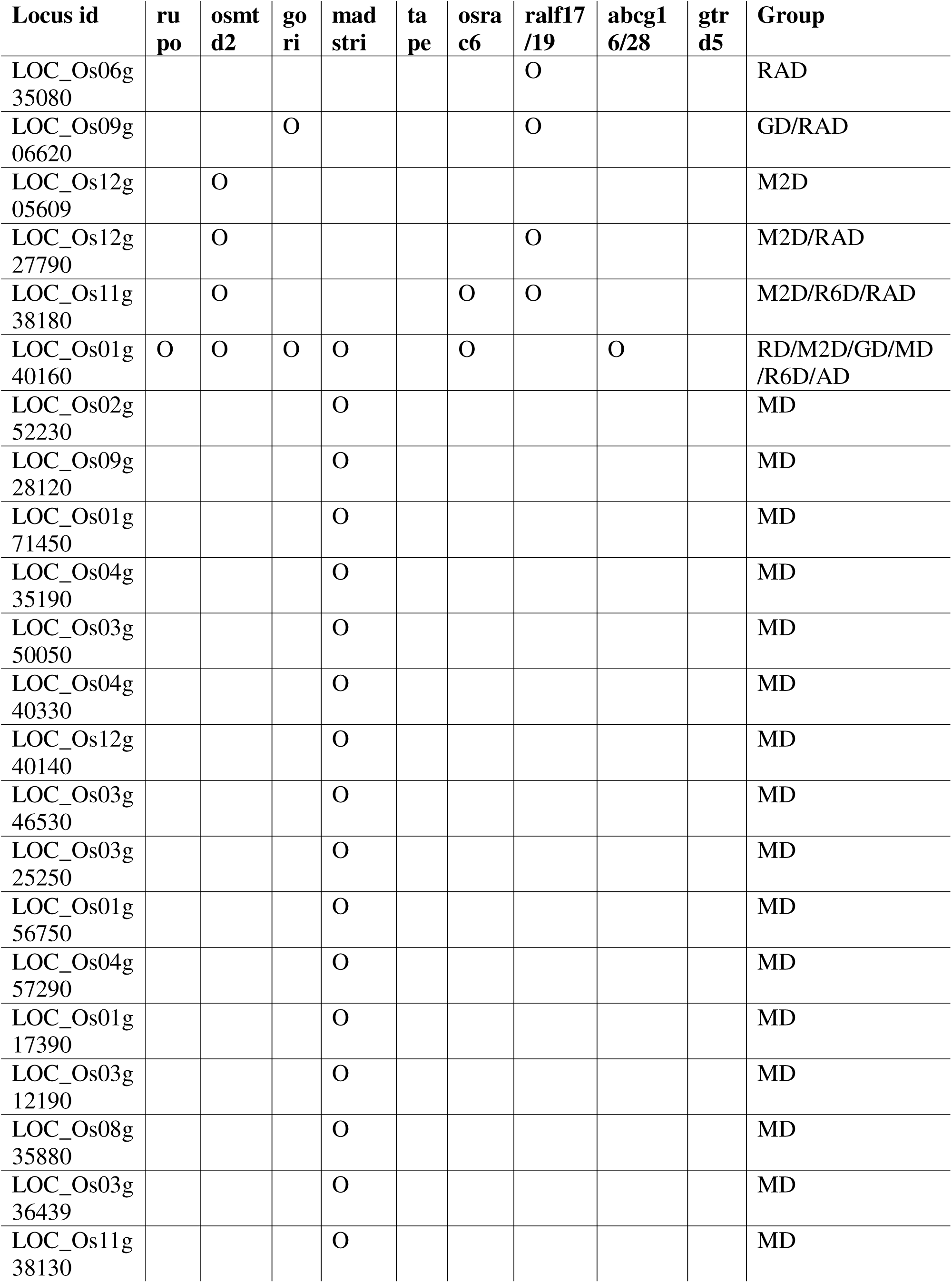
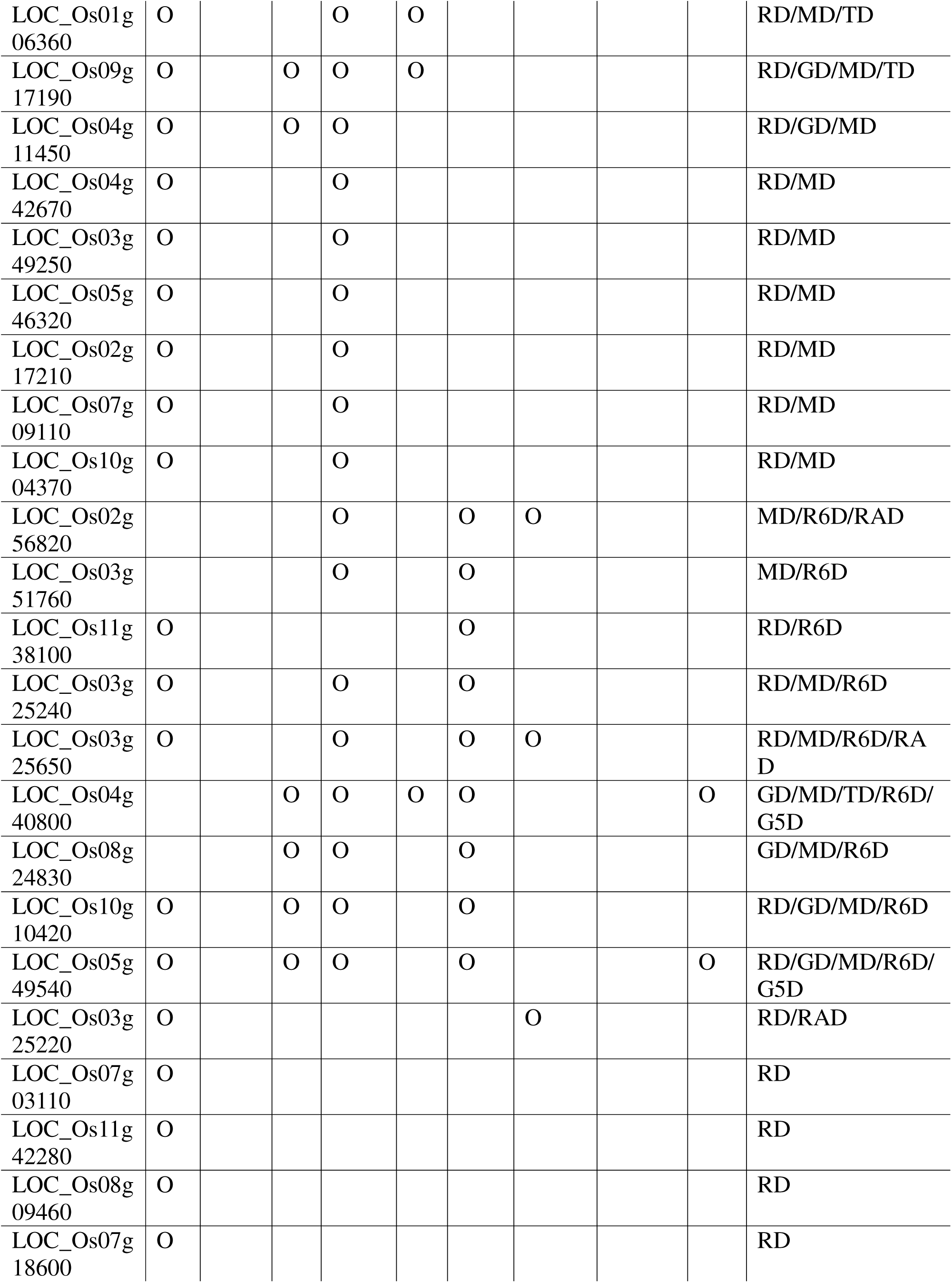
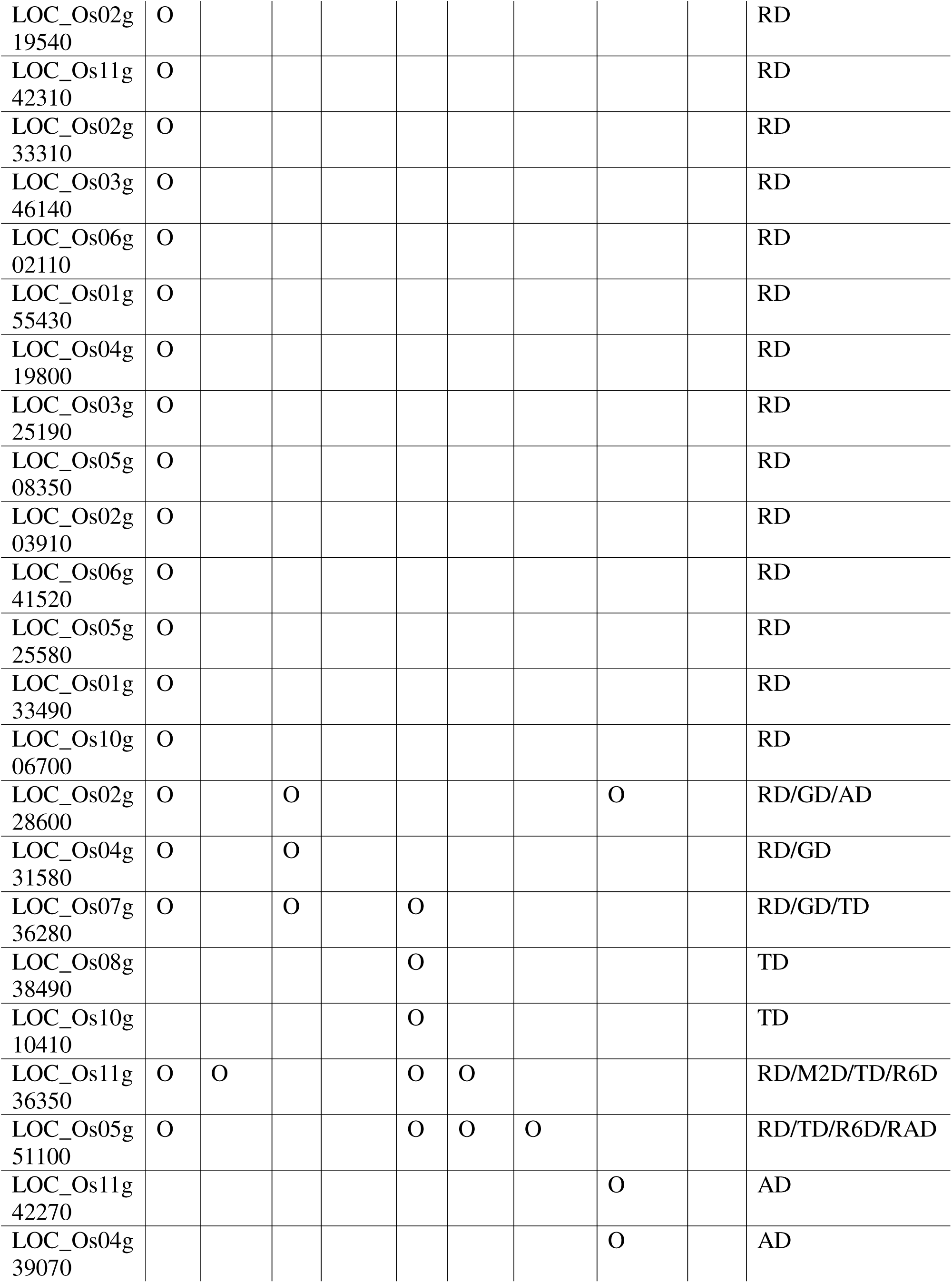
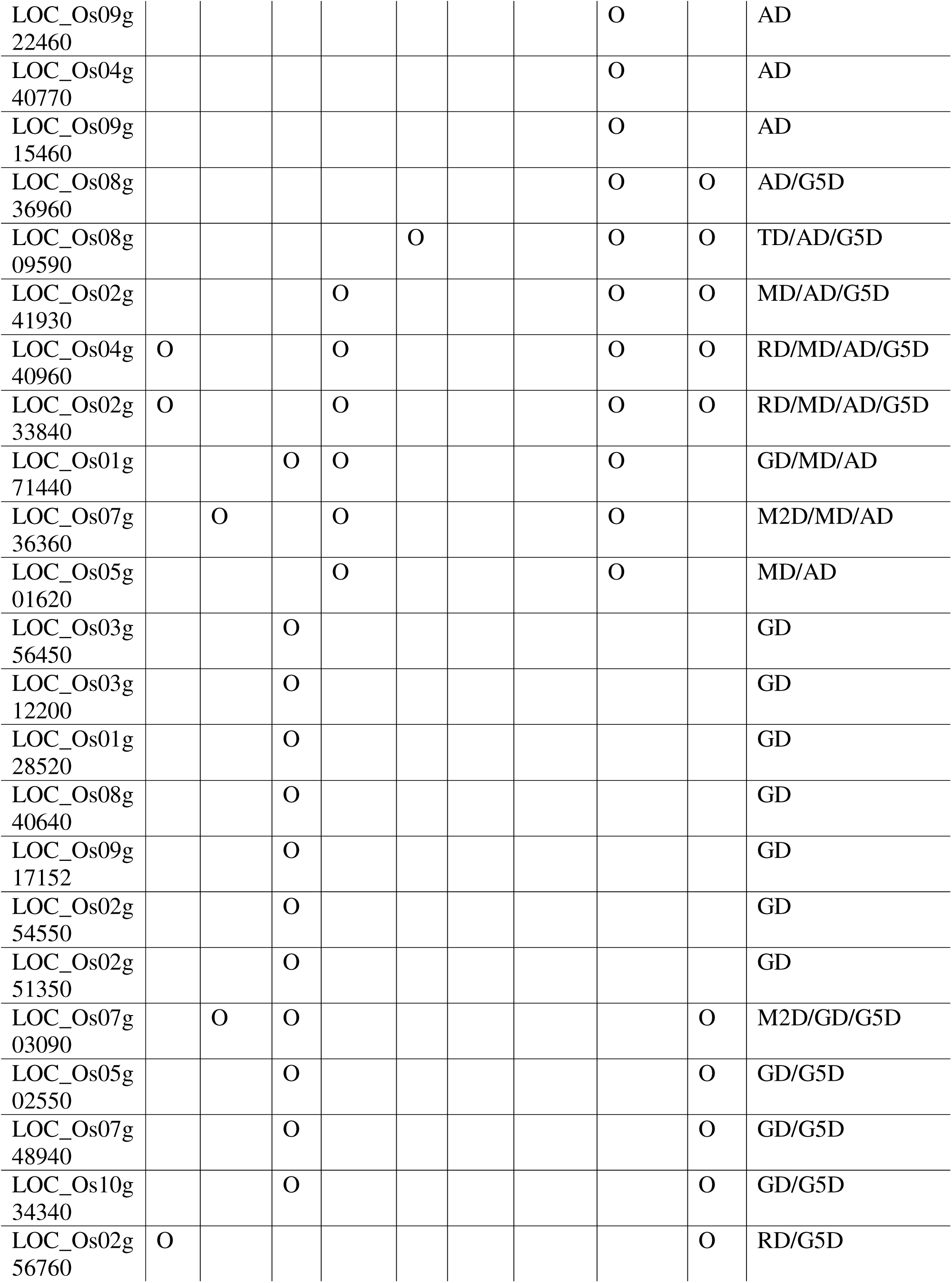
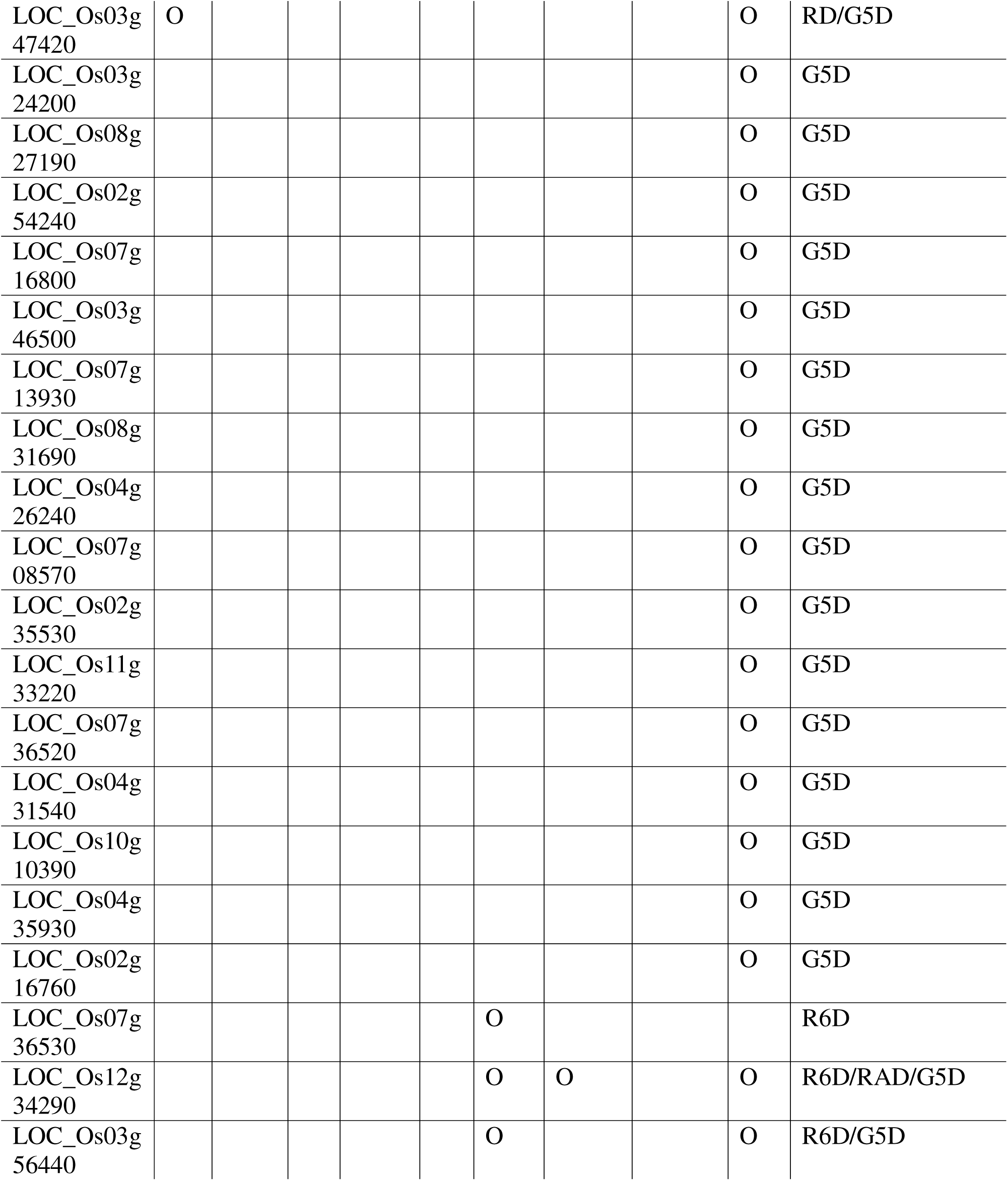
Regulation pattern of downregulated mature anther preferential F-box. RD, rupo; M2D, osmtd2; GD, gori; MD, madstri; TD, tape; R6D, osrac6; AD, abcg16/28; G5D, gtrd5; RAD, ralf17/19.

## Discussion

To demonstrate the utility of RE3DB in uncovering novel insights into ubiquitination in plant biology, we investigated key F-box genes involved in rice pollen germination. By applying a more stringent threshold (log□ fold change ≤ -2) and visualizing the overlaps using an upset plot, we uncovered distinct clusters of F-box genes in the rice pollen germination process regulated by upstream signaling components such as the RUPO receptor-like kinase and OsMADS62/63/68 transcription factors which have shown to have the broadest impact (Fig. 7-A; Table S10). In addition, F-box genes that are suppressed throughout mutant combinations are expected to help resolve specific pathway branches and stage-specific control points during rice pollen germination.

**Figure 7.**
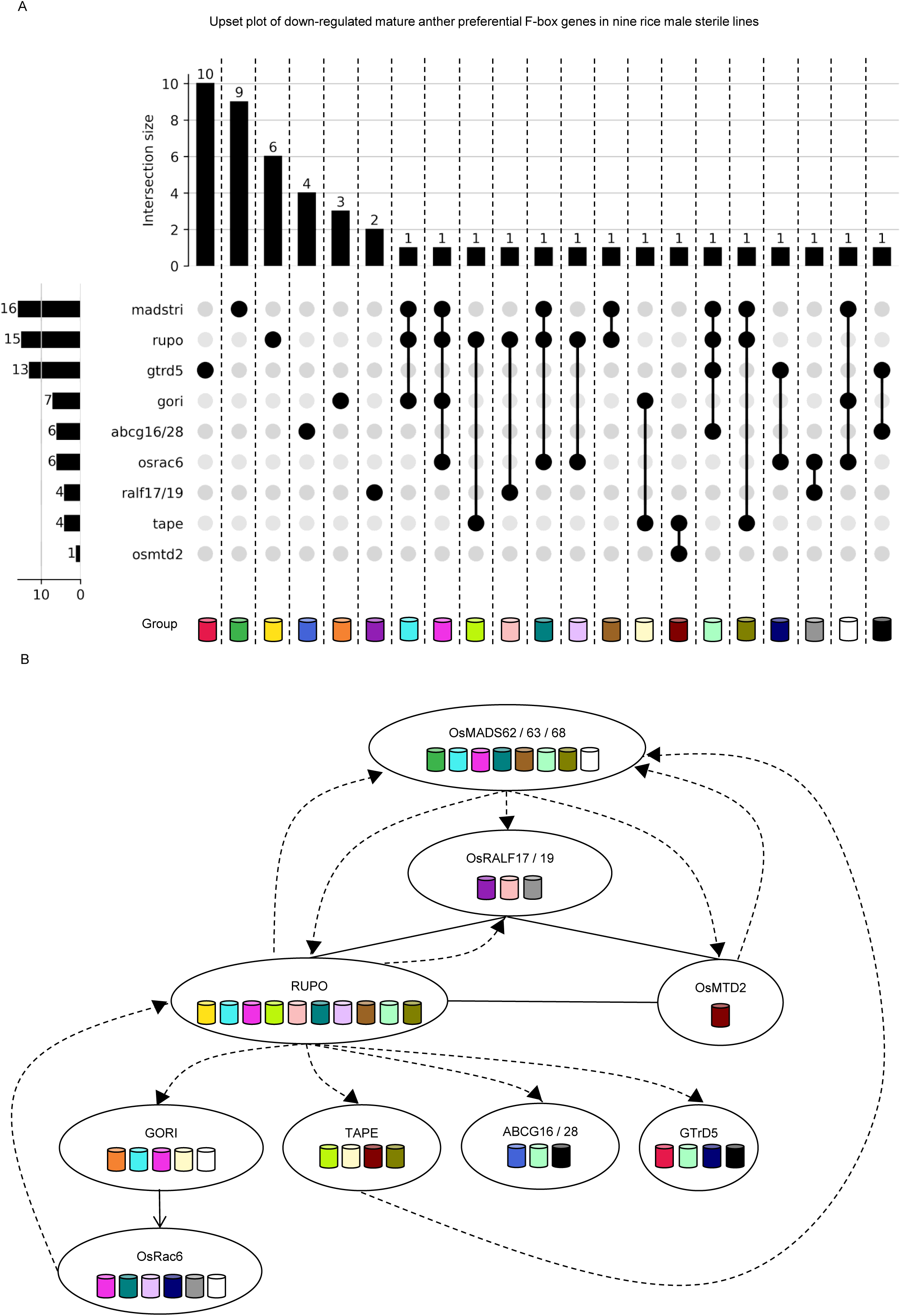
Regulatory network model of down-regulated mature anther preferential F-box genes in rice pollen germination. (A) UpSet plot of co-regulated F-box genes. Intersections show F-box loci that are down-regulated (log□ fold change ≤ −2) across nine male sterile mutants in rice pollen germination, highlighting clusters are most broadly affected by the RUPO receptor-like kinase and the OsMADS62/63/68 transcription-factor complex. Bars give gene counts per intersection and per set. Colored cylinders represent each intersection groups in the UpSet plot. (B) Regulatory network model integrating stringently co-regulated F-box genes with literature-curated targets. Solid lines represent experimentally validated regulatory relationships and interactions. Dotted lines are predicted based on regulation pattern shown through RNA sequencing (RNA-seq). The model illustrates the transcriptional hierarchy controlling pollen germination and tube growth. Colored cylinders in the model groups previously uncharacterized F-box genes with predicted upstream regulators and pathway modules for pollen tube initiation and elongation. RNA-seq, RNA sequencing.

Finally, we integrated these stringently co-regulated F-box genes with a literature-curated regulatory network comprising 13 known target genes, constructing a functional model that positions previously uncharacterized F-box genes within specific branches of the pollen tube initiation and elongation pathway (Fig. 7-B; Table S11). RNA-seq analysis of mature anthers from osmadstri (OsMADS62/63/68) triple-knockout mutants demonstrated that the OsMADS62/63/68 transcription-factor complex functions as an upstream positive regulator of RUPO during rice pollen germination, with RUPO expression being strongly down-regulated in these mutants (Kim *et al*., 2022). Furthermore, in a high-throughput screening of T-DNA insertion and CRISPR/Cas9 knockout lines, more than 50% expression reduction were observed in the following genes: (i) in *rupo* knockout lines, *GORI*, *OsMADS62*, *OsMADS68*, *TAPE*, *OsRac6*, *OsRALF19*, *OsABCG16*, *OsABCG28*, and *GTrD5* showed significant expression decreases, (ii) in *madstri* knockout mutants showed reduction in *TAPE*, *RUPO*, *OsMTD2*, *GORI*, *OsRALF17*, *OsRALF19*, *OsABCG16*, *OsABCG28*, and *GTrD5*, (iii) *OsMADS63* in *osmtd2* and *tape*, and (iv) *RUPO* in *osrac6* (Kim *et al*., 2025a). Using co-immunoprecipitation (Co-IP) assays, bimolecular fluorescence complementation (BiFC) and yeast-two-hybrid (Y2H) assays, the mature OsRALF17/19 peptides bind the OsMTD2 extracellular domain, and complementary Co-IP assays and Y2H assays show binding to RUPO plus an RUPO-OsMTD2 extracellular interaction, consistent with a heteromeric receptor complex in rice pollen tubes (Kim *et al*., 2023, 2025b). Moreover, physical interaction studies and genetic analyses confirmed that GORI acts upstream to positively modulate OsRac6 activity at the pollen-tube tip, with Co-IP revealing direct protein interaction, and over-expression crossing lines showing that extra GORI dampens Rac6-driven hyper-elongation, thereby promoting both pollen grain germination and pollen tube elongation (Kim & and Jung, 2023). Mapping these hierarchical and cross-regulatory interactions yielded a regulatory network that prioritizes a previously uncharacterized cohort of F-box genes by grouping co-regulated loci, predicting their upstream regulators, and situating them within discrete pathway modules. Collectively, the results implicate UPS mediated regulation in rice male-gametophyte function and underscore the utility of RE3DB for hypothesis-driven functional discovery.

Taken together, RE3DB constitutes the first interactive phylogenomics web platform dedicated to exploring rice E3 ligases, coupling integrated analytical tools with curated multi-omics datasets to accelerate discovery in molecular plant biology. While our case study implicating F-box regulators in pollen germination illustrates its utility, the resource more broadly enables systematic exploration and hypothesis generation for previously uncharacterized E3 ligases in rice. Moreover, because the underlying framework is readily extensible to additional crop species, RE3DB positions as a key infrastructure with broad utility for molecular plant science and future applications in agricultural biotechnology.

## Methods

### Identification, classification, and annotation of the E3 ligase genes

Candidate E3 ubiquitin-ligase loci were collated from primary literature for each major family: F-box (Jain *et al*., 2007; Wang *et al*., 2022c), U-box (Zeng *et al*., 2008), RING (Lim *et al*., 2010), CDC20 (Lin *et al*., 2022), TAD1 (Xu *et al*., 2012), Cullin (Moin *et al*., 2019), SKP1 (Kahloul *et al*., 2013), RBX1 (Liu *et al*., 2017b), BTB (Shalmani *et al*., 2021; Wang *et al*., 2022c), DDB1 (Ishibashi *et al*., 2003; Zang *et al*., 2016), DWD (Lee *et al*., 2008), APC (Eloy *et al*., 2015; Wang *et al*., 2022c), and HECT (Marín, 2013). After integration, the list was matched against the RGAP (MSU7) catalogue, and only genes with valid RGAP identifiers were retained. For each retained locus, the protein sequence of the longest annotated transcript was extracted from the RGAP release dated 1 September 2024 (https://rice.uga.edu/osa1r7_download/osa1_r7.all_models.pep.fa.gz) (Kawahara *et al*., 2013; Hamilton *et al*., 2025). Domain repertoires including Pfam v36.0 (Pei *et al*., 2024; Paysan-Lafosse *et al*., 2025) and InterPro (Blum *et al*., 2025) were reassessed with InterProScan v5.73-104.0 (Jones *et al*., 2014).

### Protein-sequence processing and phylogenetic reconstruction

Representative protein sequences were grouped by E3-ligase family and written to individual FASTA files with the SeqIO module of Biopython v1.85 (Cock *et al*., 2009). Families containing at least two members were aligned with MAFFT v7.526 using the *--auto* setting, which selects the most suitable algorithm and reorders sequences to maximize alignment accuracy (Katoh *et al*., 2002; Katoh & Standley, 2013). Phylogenies were inferred on an alignment-specific basis. For families with ≥ 4 members, IQ-TREE v3.0.0 (Wong *et al*., 2025) was executed with automatic model selection via ModelFinder (Kalyaanamoorthy *et al*., 2017), branch support assessed using 1,000 ultrafast bootstrap replicates (Hoang *et al*., 2018) and 1000 SH-aLRT tests (Guindon *et al*., 2010). For smaller families comprising two or three genes, FastTree v2.1.9 was employed with 100 bootstrap replicates (Price *et al*., 2010). All resulting trees were exported in Newick format for downstream visualization and comparative analyses.

### RNA-seq processing and co-expression analysis

RNA-seq libraries representing anatomical tissues, abiotic and biotic-stress treatments, hormonal and nutrient-starvation conditions were collected from the NCBI Sequence Read Archive (NCBI SRA; https://www.ncbi.nlm.nih.gov/sra) and EMBL-EBI ArrayExpress (https://www.ebi.ac.uk/biostudies/arrayexpress). FASTQ files were downloaded in parallel with parallel-fastq-dump v2.9.1 (https://github.com/rvalieris/parallel-fastq-dump). Adapter sequences and bases with Phred scores < 20 were removed using Cutadapt v1.18 (Martin, 2011). Quality-filtered reads were aligned to the Oryza sativa MSU7 reference genome with HISAT2 v2.1.0 (Kim *et al*., 2019) under default parameters, and gene-level counts were obtained with featureCounts v1.6.3 (Liao *et al*., 2014). Count matrices for each experimental series were normalized independently with DESeq2 v1.20.0 (Love *et al*., 2014) and merged to extend the expression heatmap in the CRISPR applicable functional redundancy inspector to accelerate functional genomics in rice (CAFRI-Rice; https://cafri-rice.khu.ac.kr/) (Hong *et al*., 2020) and the Rice Online Expression Profiles Array Database version 2 (ROADv2; https://roadv2.khu.ac.kr/) (Hwang *et al*., 2024) framework to the 25 tissues and multi-stress compendium used in RE3DB. PCC between a query gene and the remaining loci were computed with Pandas v2.2.3 (McKinney, 2010; The pandas development team, 2024). Two-tailed P-values were calculated with SciPy v1.15.2 (McKinney, 2010). These PCC and P-value estimates form the basis of the co-expression analysis provided in RE3DB.

### Processing of PPI data for predicting E3-substrate associations

Nodes and edges were collected from six public resources: STRING database (Szklarczyk *et al*., 2023), RicePPINet (Liu *et al*., 2017a), PRIN (Gu *et al*., 2011), RiceNet v2 (Lee *et al*., 2015), UbE3-ORFeome (Wang *et al*., 2022d), and protein structural interactome predicted based on protein docking (Sun *et al*., 2024). Dataset-specific thresholds were applied to retain only high-confidence interactions: combined score ≥ 0.7 for STRING, confidence score ≥ 0.5 for RicePPINet, the high-confidence subset supplied by PRIN, summed log-likelihood score ≥ 2 for RiceNet v2 and Z-score ≥ 8.8 for the docking-based interactome. UbE3-ORFeome interactions were incorporated without additional filtering. To identify substrates whose protein turnover is uncoupled from transcript abundance, the PPI network was intersected with paired transcriptomic and proteomic measurements from 14 rice tissues (Li *et al*., 2024): root tip, whole root, culm and leaves, stem, leaf sheath, auricle, ligule and lamina joint, leaf blade, spike neck, mature spikelet, pistil, pollen, early-stage immature seed, grain-filling seed and mature seed. Genes expressed at < 1 TPM in any tissue were excluded. For the remaining genes, PTRs were calculated in linear space and log□-transformed. Within each tissue, the median (M) and standard deviation (SD) of log□ PTR were derived; a gene was designated as low PTR in that tissue if log□ PTR < M - 2 SD. Genes meeting this criterion in at least one tissue were classified as low-PTR candidates. Each retained edge was subsequently annotated with the previously computed Pearson correlation coefficient and its associated P-value, yielding an integrated, confidence-ranked network for E3-substrate interaction prediction.

### Interactive E3 Ligase Family Visualization

We employed the d3-cloud (Davies, 2013) layout to generate an interactive word cloud for the E3 Ligase family. In this visualization, the font size of each E3 family term is dynamically adjusted based on the number of genes associated with that family. Additionally, we created an interactive table containing detailed information about the corresponding E3 ligase using a DataTable widget from the Bokeh package version 3.2.2 (Bokeh Development Team, 2025).

### PPI Network Visualization

PCC values of a user entered locus id and the rest of the rice gene loci are queried and filtered by a user entered minimum PCC value. We visualized an interactive PPI network graph by implementing networkx package, version 3.0 (Hagberg *et al*., 2008), and Bokeh package, version 3.2.2 (Bokeh Development Team, 2025). In the interactive table, genes are categorized by different colors: entries annotated as E3 were highlighted in blue, those classified as low protein-to-RNA ratio (low-PTR) in orange, and entries meeting both criteria in grey.

### Phylogenetic Heatmaps Visualization

Phylogenetic and transcriptomic datasets were integrated by constructing tree-anchored heat maps with the ETE3 framework (Huerta-Cepas *et al*., 2016). The phylogeny was pruned to its nearest clade using ETE3’s Prune function. PCC and p-value were computed between the user searched gene’s expression profile and those of all genes in the tree. Expression matrices were visualized as heatmaps with linear color mapping using Bokeh version 3.2.2 (Bokeh Development Team, 2025). For the “anatomy” dataset, log_2_ expression values from 0 to 10 were rendered on a blue to yellow gradient. For the “biotic,” “abiotic,” “nutrient,” and “hormone” conditions, log_2_ fold changes spanning −3 to 3 were rendered on a light green to light red gradient.

### Assignment of E3 Ubiquitin-Ligase Genes to Tissue and Developmental Groups

We classified tissue[preferential E3 ubiquitin-ligase genes from a log□ scale DESeq2-normalized expression matrix spanning 25 rice tissues and organs by applying per-gene Z-score standardization across tissues. For each gene, the mean and standard deviation across all tissue columns were computed using NumPy version 2.1.3 (Harris *et al*., 2020) and Pandas v2.2.3 (McKinney, 2010; The pandas development team, 2024); Z-scores were obtained by mean-centering and scaling the gene’s profile. Genes with zero variance were assigned missing denominators to avoid division by zero and were consequently not called as preferential. Anatomical groupings were defined to reflect developmental and lineage relationships. Group membership was evaluated independently: a gene was called tissue-preferential for a given group if one or more member tissues in that group exhibited Z ≥ 2 and no tissue outside the group reached this threshold. Because groups are nested, a gene could be reported for both a parent and its subgroups when the criterion was simultaneously satisfied.

### Over-representation analysis of E3 ubiquitin-ligase families

We performed over-representation analysis separately for each E3 ubiquitin-ligase family. First, we constructed a gene-set library from a tissue-preferential gene and group assignment file. We then mapped each tissue term to its unique set of associated gene loci. For each family, we deduplicated loci and applied hypergeometric test against the custom library, obtaining odds ratios and FDR adjusted P-values. Using Matplotlib version 3.10 (Hunter, 2007) and GSEApy version 1.1.9 (Fang *et al*., 2023), we visualized, for every family, the five most over-represented terms ranked by odds ratio.

### Detection of down-regulated genes and intersection analysis

For each mutant, genes were classified as down-regulated when log2 fold-change < -1 or a stricter threshold −2. The overlap structure among mutants was summarized by grouping genes that shared identical membership patterns across the nine contrasts; group labels were derived from the subset of mutants in which a gene was downregulated. Using UpSetPlot version 0.9.0 (Lex *et al*., 2014), an UpSet diagram was generated from these sets with counts displayed and intersections ordered by cardinality to visualize set sizes and their overlaps. We generated a gene table in which each mutant column was marked “O” if the gene met the down-regulation criterion in that mutant and left blank otherwise, and an additional “Group” field for implicated mutants; rupo as RD, osmtd2 as M2D, gori as GD, madstri as MD, tape as TD, osrac6 as R6D, abcg16/28 as AD, gtrd5 as G5D, and ralf17/19 as RAD.

### RE3DB Application Implementation and Deployment Architecture

RE3DB was implemented in Python v3.11.5 (https://www.python.org/downloads/release/python-3115/) with the Django framework v4.2.4 (https://www.djangoproject.com/) and deployed on Ubuntu 22.04 (Jammy; https://releases.ubuntu.com/jammy/). The production stack uses Nginx v1.22.1 (https://nginx.org) as a reverse proxy and static file server in front of a Gunicorn v20.1.0 WSGI application (https://gunicorn.org), with persistent storage in MySQL v8.0 (https://dev.mysql.com/doc/refman/8.0/en/). To support ingestion of diverse file formats, we integrated django-import-export v3.0.2 (https://github.com/django-import-export/django-import-export); long-running imports are dispatched to an asynchronous task queue via Celery v5.2.7 (https://github.com/celery/celery) using the django-import-export-celery bridge v1.3 (https://github.com/auto-mat/django-import-export-celery), thereby preserving responsiveness for interactive web requests. The client interface employs Bootstrap v5.3.1 (https://getbootstrap.com) to provide consistent layout across screen sizes and operating systems. The application has been verified on desktop platforms such as macOS, Windows, and Ubuntu, and functions with current versions of major browsers such as Chrome, Firefox, Microsoft Edge, and Opera. RE3DB is publicly available at https://re3db.khu.ac.kr/.

## Acknowledgements

This work was supported by the grant received from the National Research Foundation of Korea (RS-2021-NR060084 and RS-2021-NR059380 to KHJ). We are grateful to Professor Woo-Jong Hong for generously providing both the source code used to implement phylogenetic heatmap and server infrastructure that enabled us to host and maintain RE3DB.

## Author contributions

KHJ: Conceptualization, Resources, Supervision, Writing (review & editing), Funding acquisition. WH: Conceptualization, Data curation, Formal analysis, Visualization, Methodology, Software development, Writing (original draft, review & editing). All authors reviewed and approved the final manuscript.

## Supplemental Data

Fig. S1. E3 family composition in tissue preferential groups.

Fig. S2. Line plot of tissue preferential U-box groups.

Fig. S3. Line plot of tissue preferential SKP1 groups.

Fig. S4. Line plot of tissue preferential RING groups.

Fig. S5. Line plot of tissue preferential HECT groups.

Fig. S6. Line plot of tissue preferential F-box groups.

Fig. S7. Line plot of tissue preferential DWD groups.

Fig. S8. Line plot of tissue preferential Cullin groups.

Fig. S9. Line plot of tissue preferential BTB groups.

Fig. S10. Line plot of tissue preferential APC groups.

Fig. S11. Line plot of tissue preferential TAD1 groups.

Fig. S12. Filtered tissue preferential E3 ligase enrichment results in bar plot.

Table S1. Anatomy metadata of RNA-seq data used in RE3DB.

Table S2. Abiotic metadata of RNA-seq data used in RE3DB.

Table S3. Biotic metadata of RNA-seq data used in RE3DB.

Table S4. Nutrient metadata of RNA-seq data used in RE3DB.

Table S5. Hormone metadata of RNA-seq data used in RE3DB.

Table S6. List of tissue preferential E3 ligase genes.

Table S7. Number statistics of tissue preferential E3 ligase genes.

Table S8. E3 ligase enrichment result.

Table S9. Functionally characterized enriched E3 ligase genes.

Table S10. Downregulated F-box genes in upset plot.

Table S11. Literature evidence for the functional regulatory model of rice pollen germination.

