## Supplementary figures and images for "RE3DB: A multi-omics phylogenomics platform for rice E3 ubiquitin ligases identifies novel regulators of pollen germination"

### Fig.S1

#### Slide 1
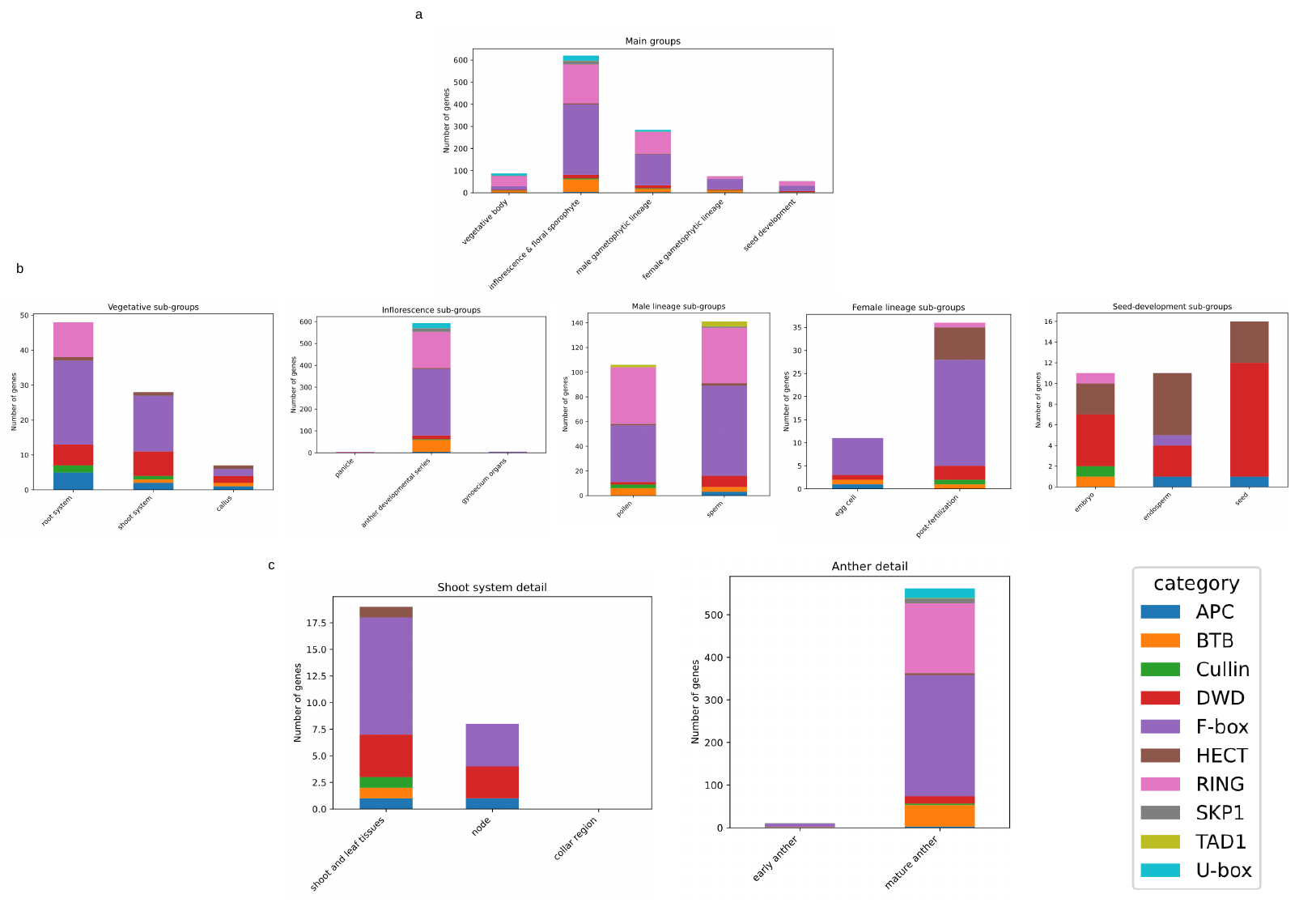

a
b
c

### Fig.S2

#### Slide 1
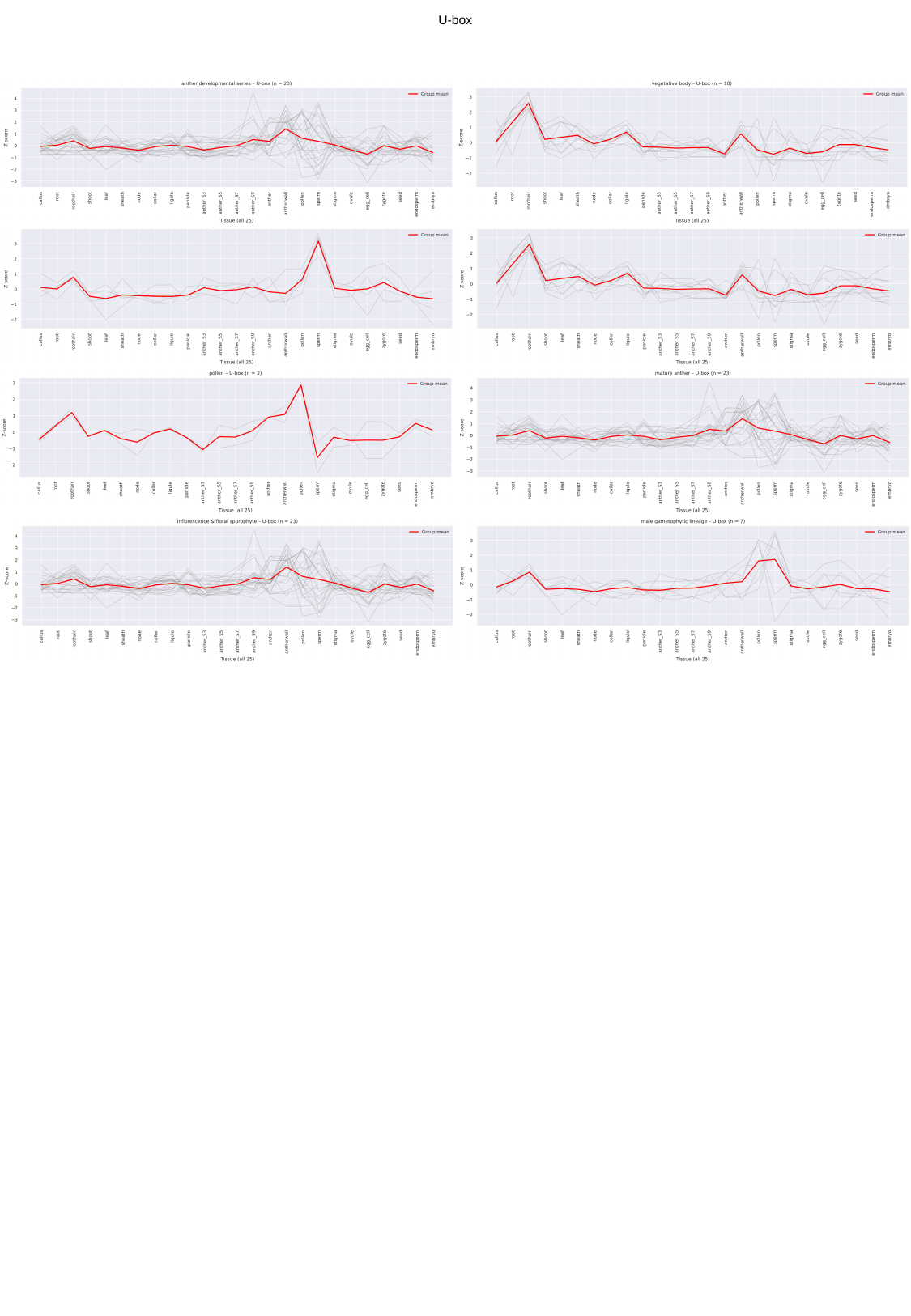

U-box

### Fig.S3

#### Slide 1
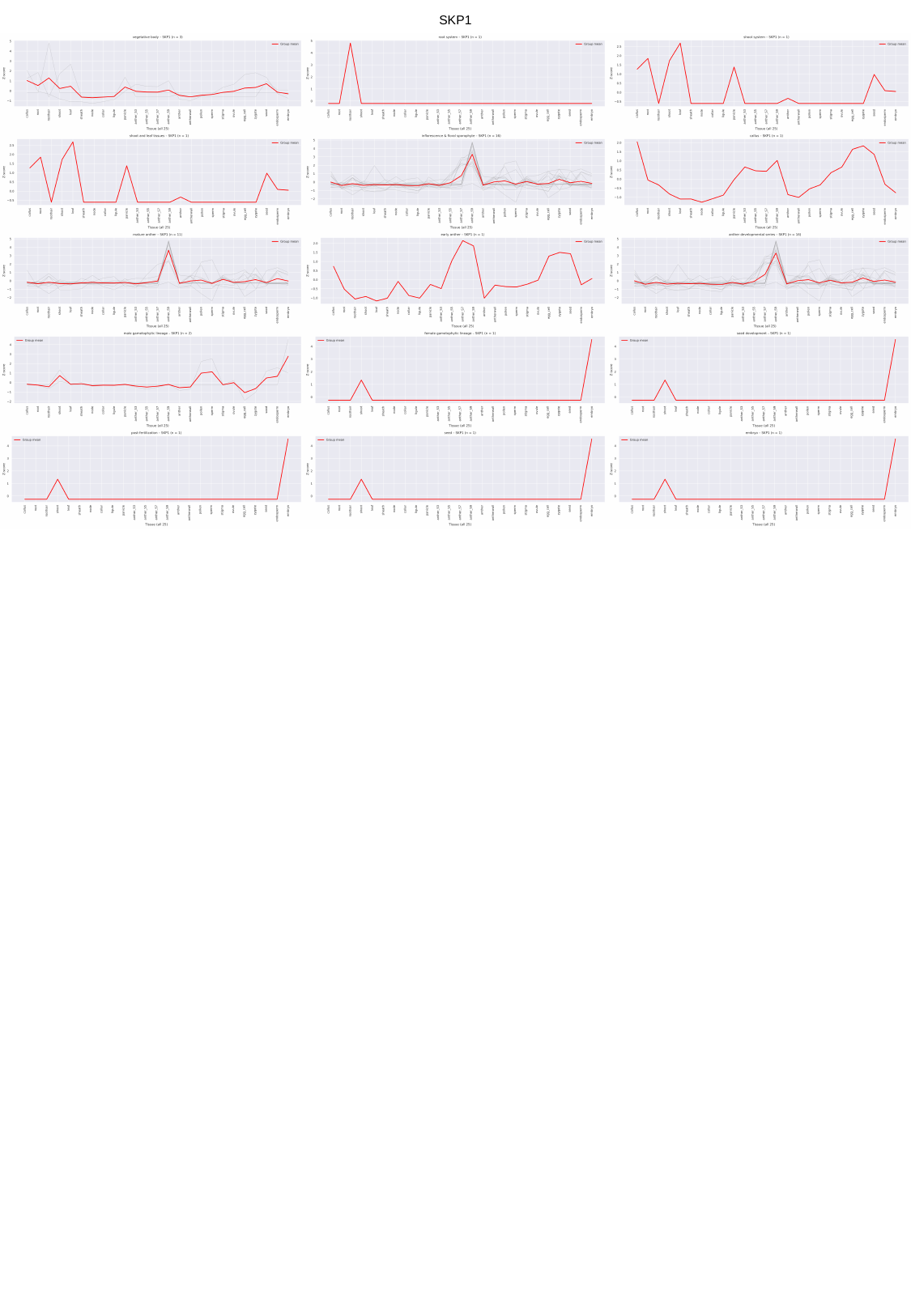

SKP1

### Fig.S4

#### Slide 1
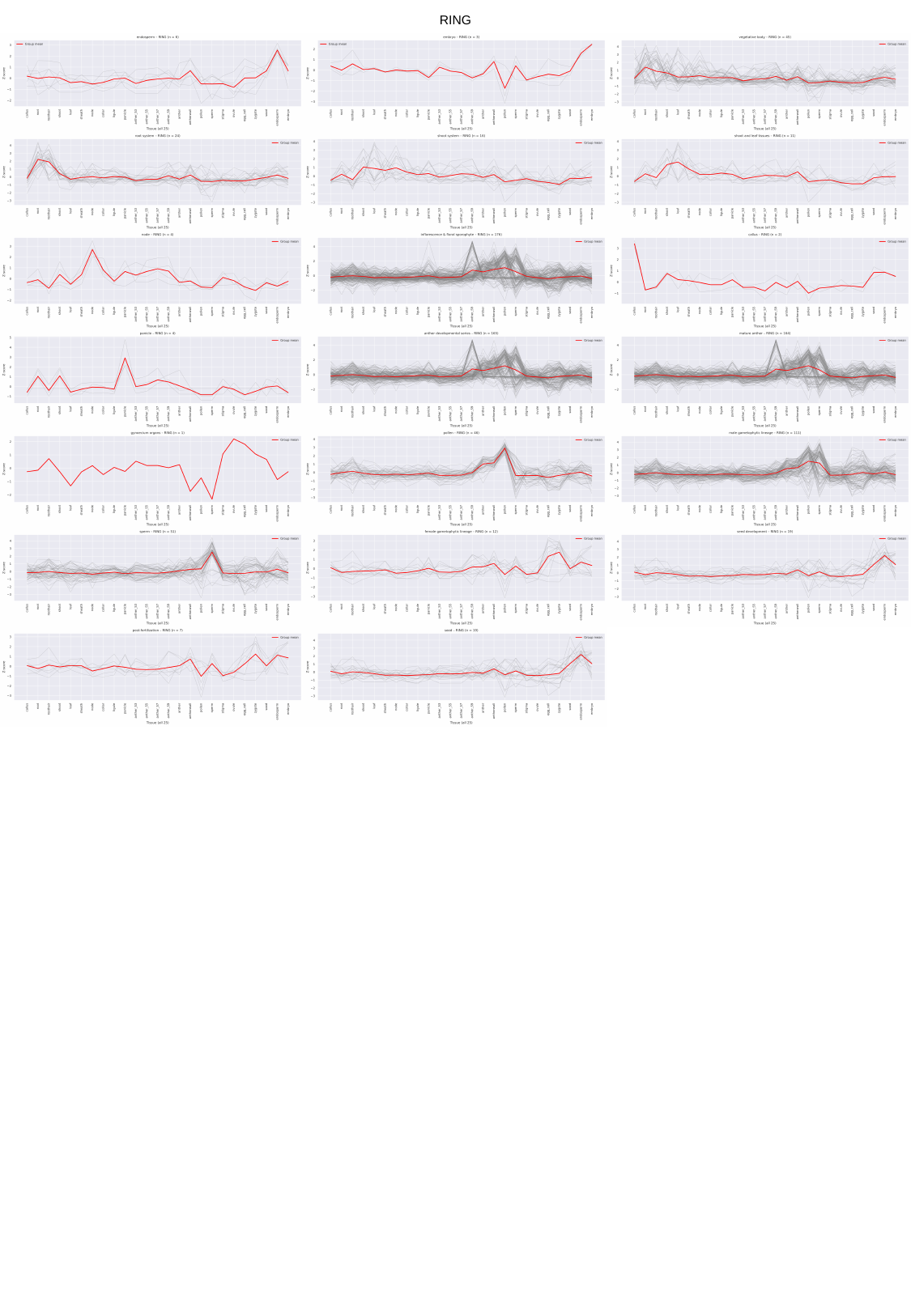

RING

### Fig.S5

#### Slide 1
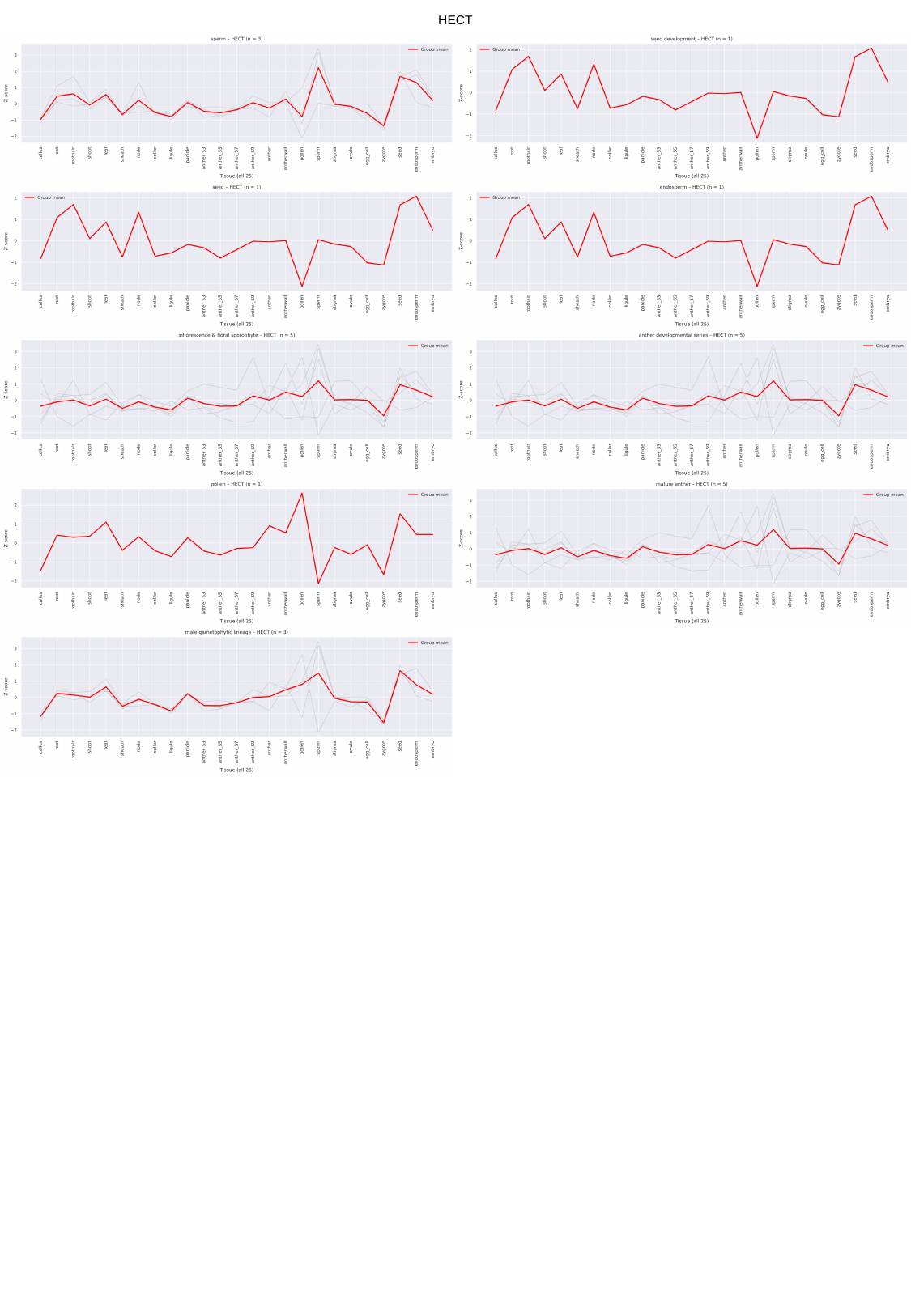

HECT

### Fig.S6

#### Slide 1
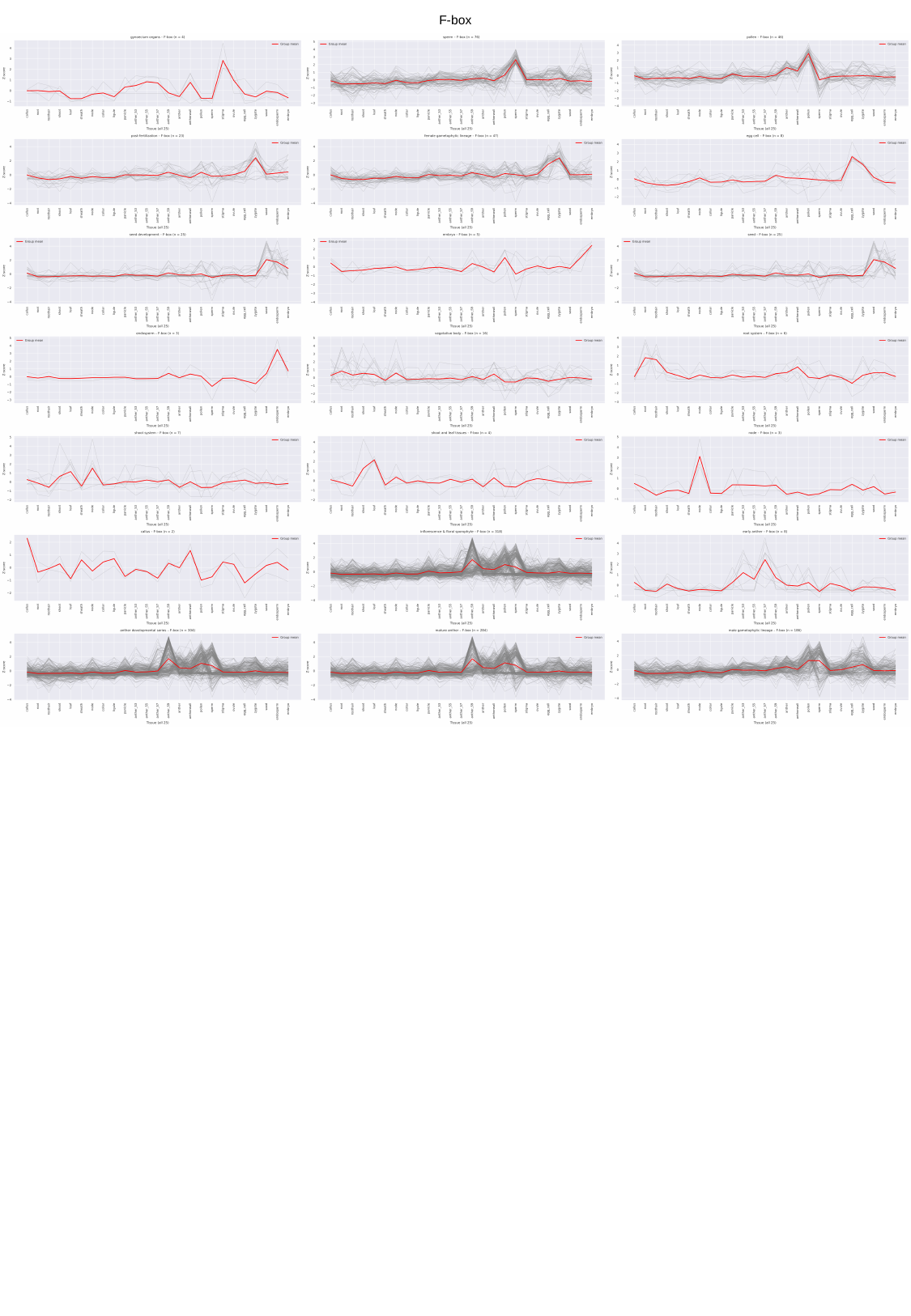

F-box

### Fig.S7

#### Slide 1
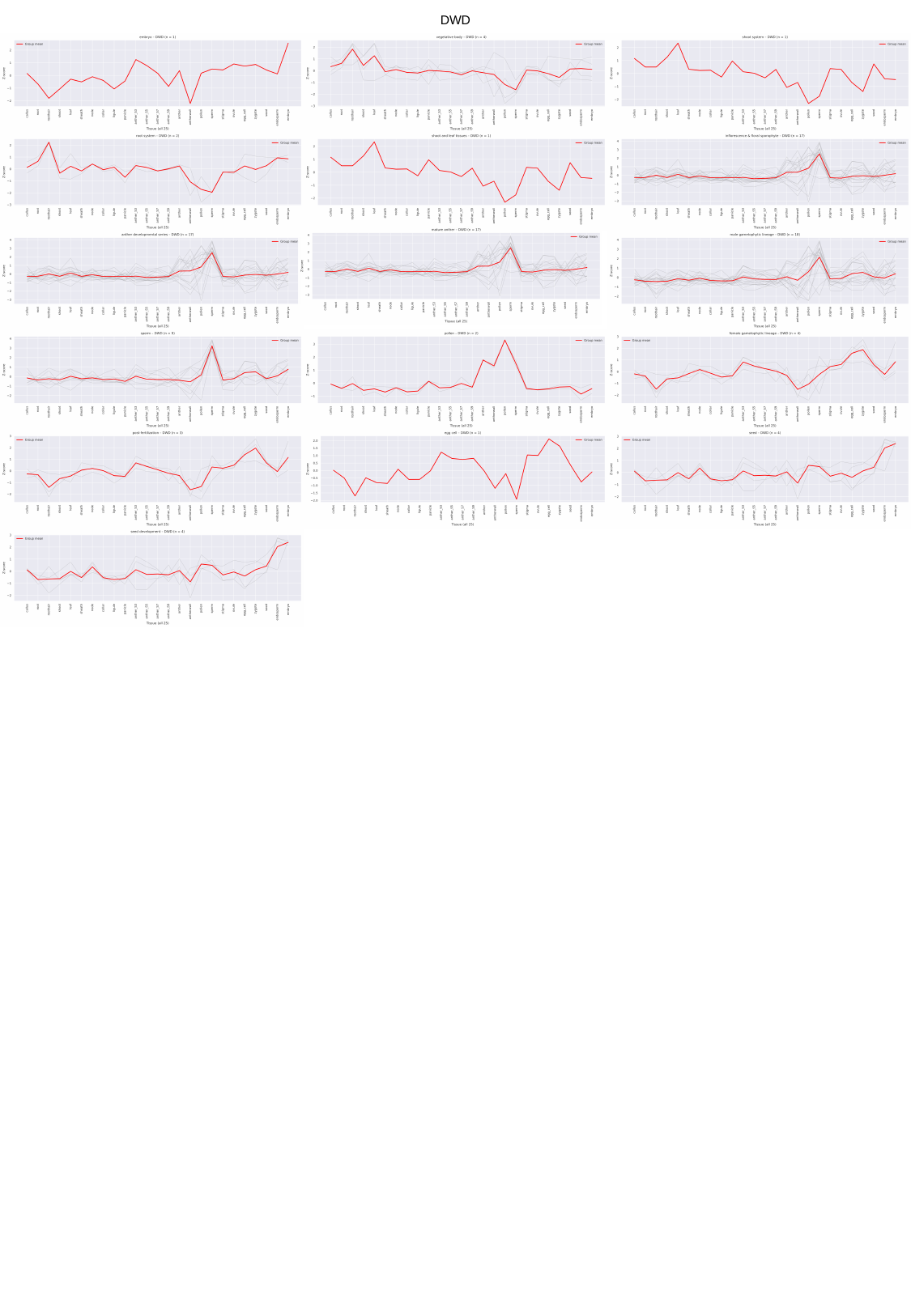

DWD

### Fig.S8

#### Slide 1
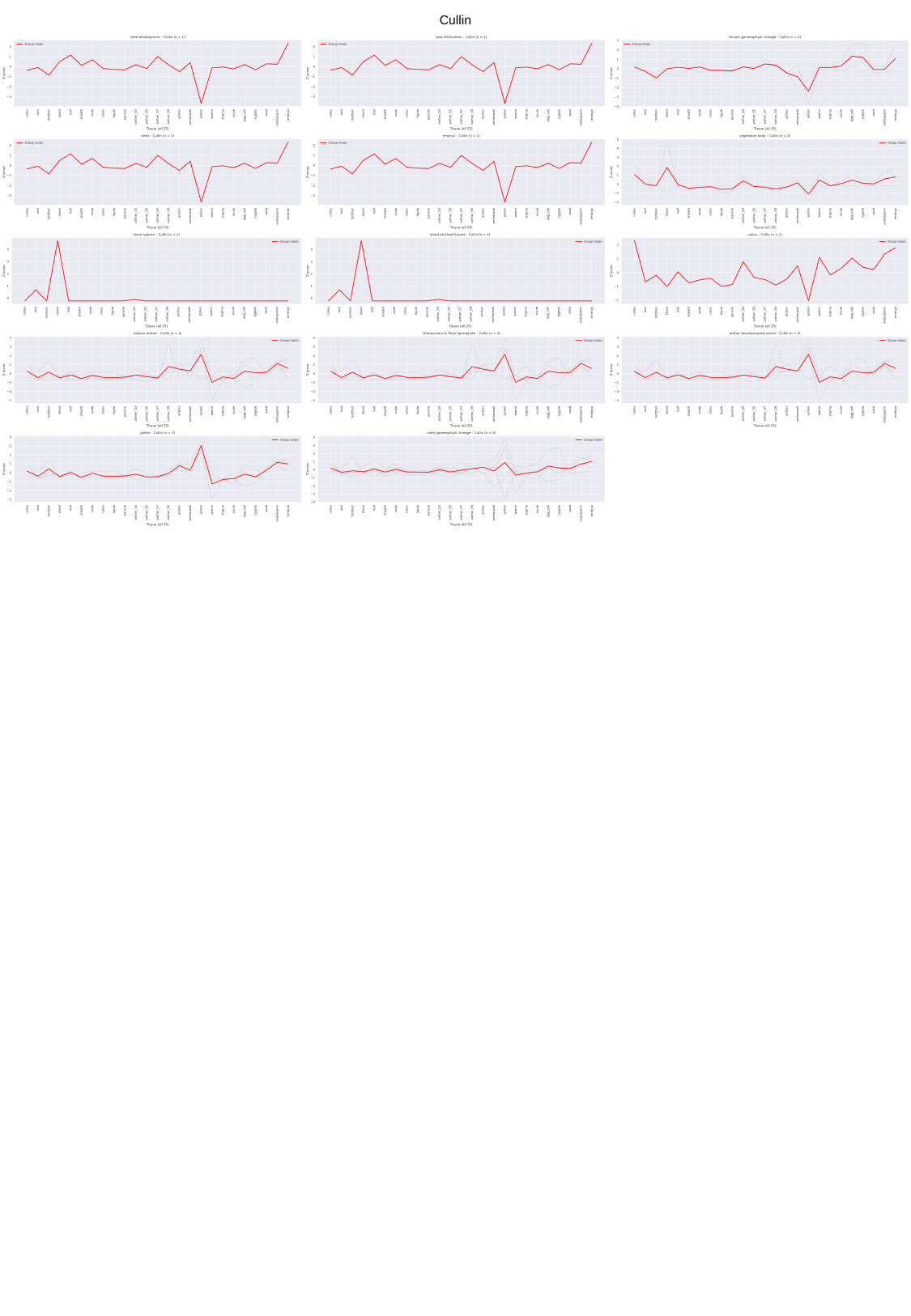

Cullin

### Fig.S9

#### Slide 1
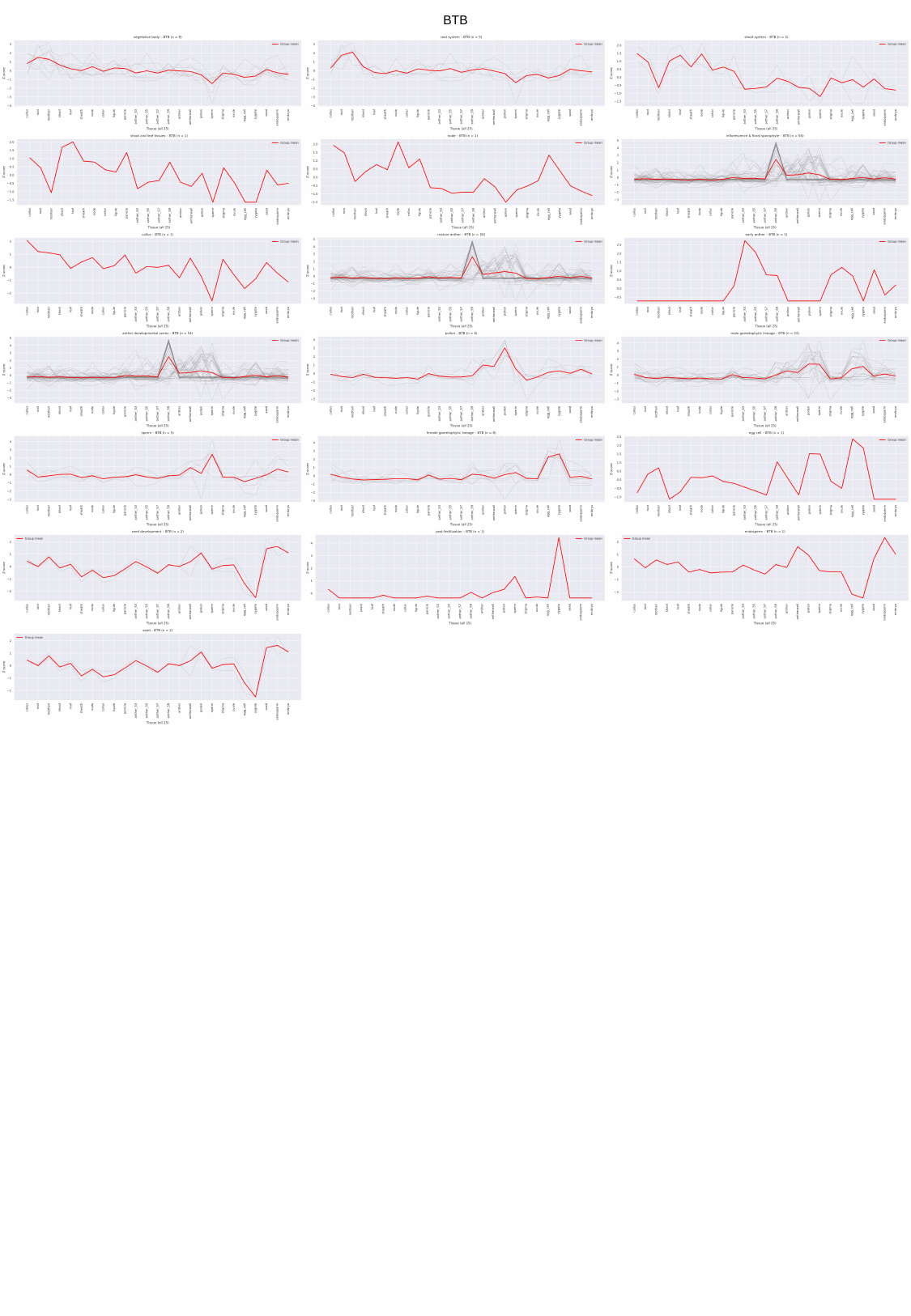

BTB

### Fig.S10

#### Slide 1
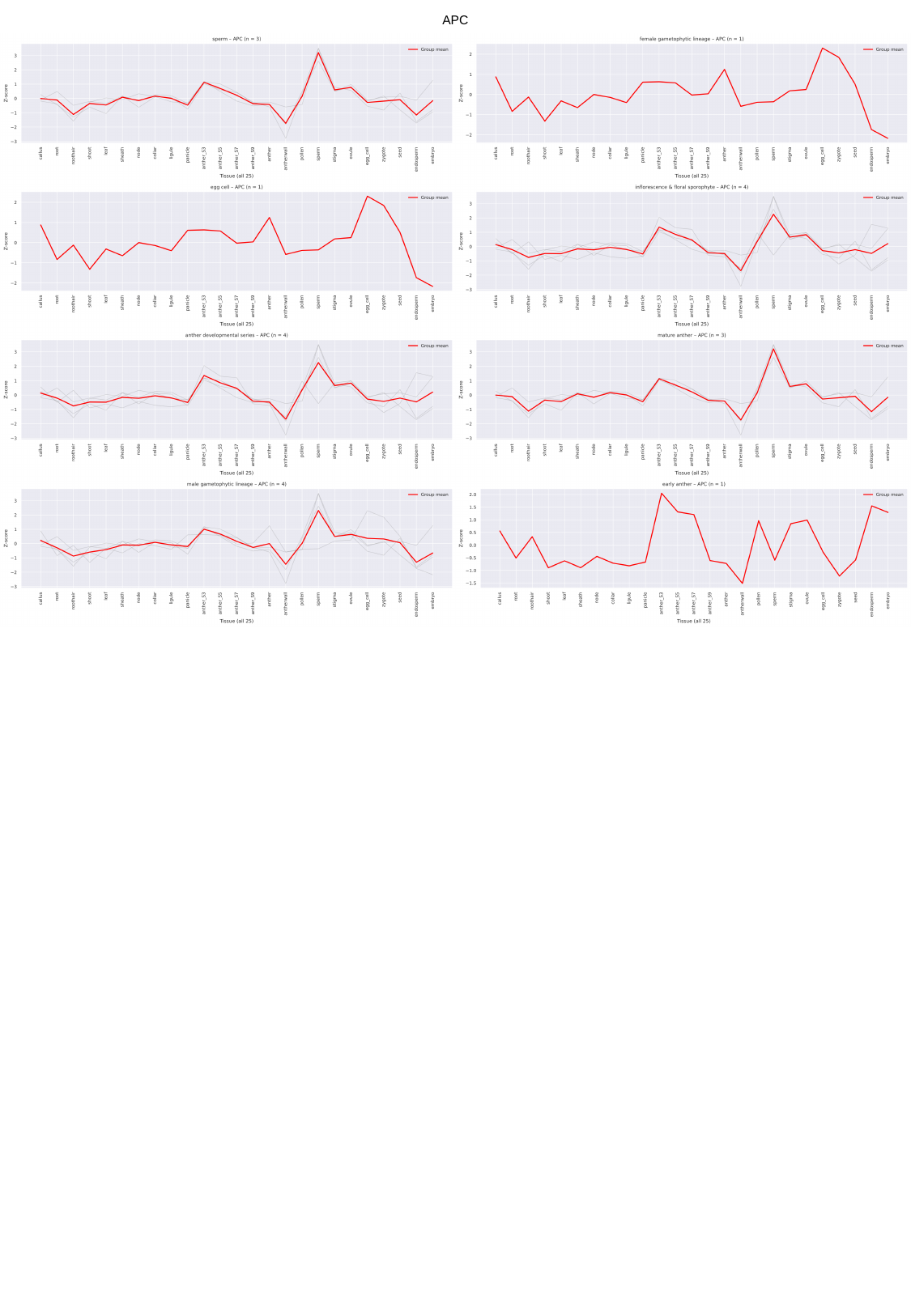

APC

### Fig.S11

#### Slide 1
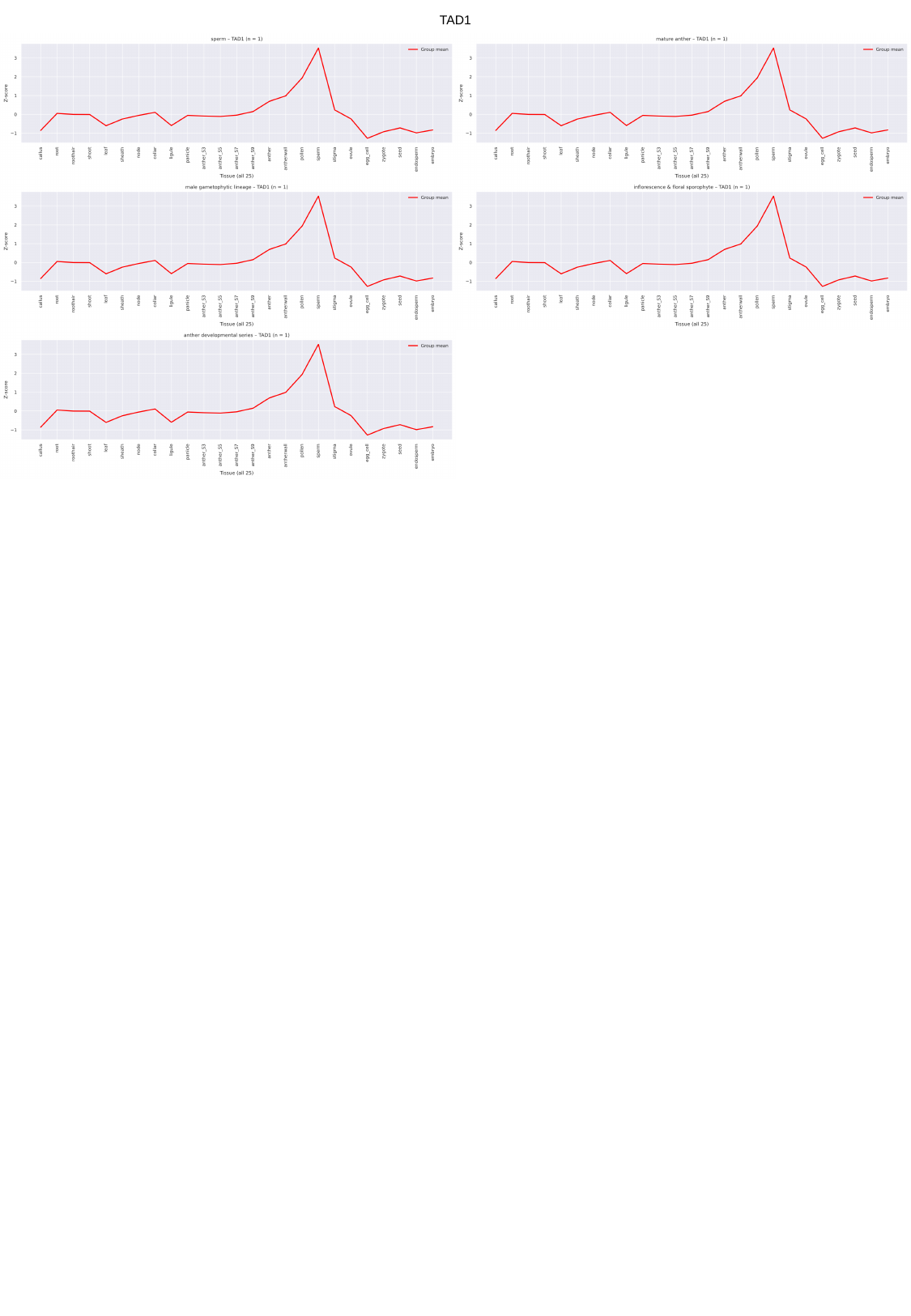

TAD1

### Fig.S12

#### Slide 1
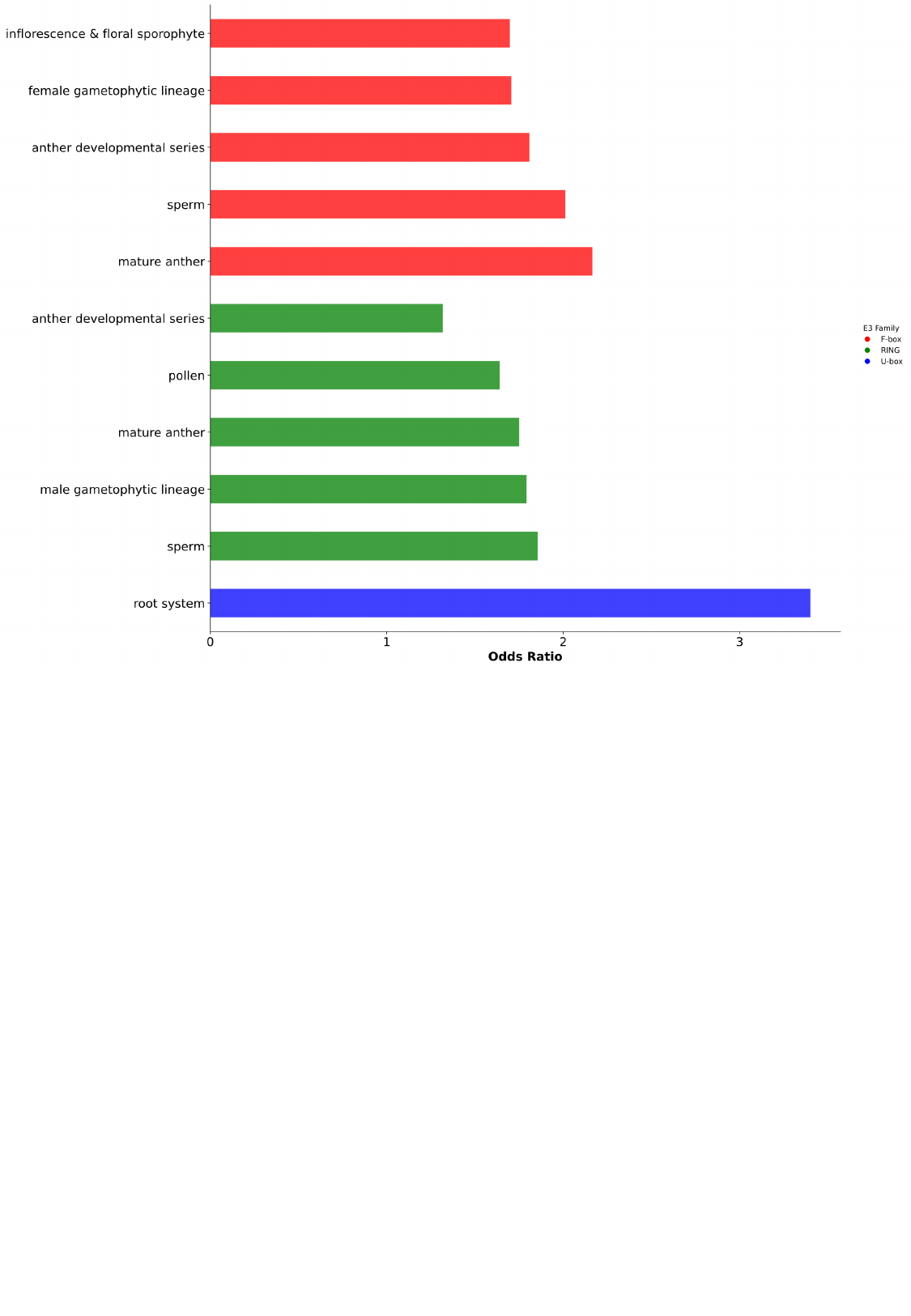
