## Supplementary material for "RE3DB: A multi-omics phylogenomics platform for rice E3 ubiquitin ligases identifies novel regulators of pollen germination": Table S1

| **Run** | **Tissue_simplify** | **Tissue_original** | **Stage** | **Cultivar** | **Subspecies** | **Layout** | **Strategy** | **Source** | **ETC** |
| --- | --- | --- | --- | --- | --- | --- | --- | --- | --- |
| DRR001029 | Callus_1 | Callus |  | Nipponbare | Japonica | Single | mRNA-Seq | 10.1093/gbe/evr111 |  |
| DRR001036 | Callus_2 | Callus |  | Nipponbare | Japonica | Single | mRNA-Seq | 10.1093/gbe/evr111 |  |
| DRR001043 | Callus_3 | Callus |  | Nipponbare | Japonica | Single | mRNA-Seq | 10.1093/gbe/evr111 |  |
| DRR001050 | Callus_4 | Callus |  | Nipponbare | Japonica | Single | mRNA-Seq | 10.1093/gbe/evr111 |  |
| DRR001026 | Shoot_1 | Shoot | 7 days after germination | Nipponbare | Japonica | Single | mRNA-Seq | 10.1093/gbe/evr111 |  |
| DRR001033 | Shoot_2 | Shoot | 7 days after germination | Nipponbare | Japonica | Single | mRNA-Seq | 10.1093/gbe/evr111 |  |
| DRR001040 | Shoot_3 | Shoot | 7 days after germination | Nipponbare | Japonica | Single | mRNA-Seq | 10.1093/gbe/evr111 |  |
| DRR001047 | Shoot_4 | Shoot | 7 days after germination | Nipponbare | Japonica | Single | mRNA-Seq | 10.1093/gbe/evr111 |  |
| DRR001024 | Leaf_1 | Leaf | 7 days before flowering to 7 days after flowering | Nipponbare | Japonica | Single | mRNA-Seq | 10.1093/gbe/evr111 |  |
| DRR001031 | Leaf_2 | Leaf | 7 days before flowering to 7 days after flowering | Nipponbare | Japonica | Single | mRNA-Seq | 10.1093/gbe/evr111 |  |
| DRR001038 | Leaf_3 | Leaf | 7 days before flowering to 7 days after flowering | Nipponbare | Japonica | Single | mRNA-Seq | 10.1093/gbe/evr111 |  |
| DRR001045 | Leaf_4 | Leaf | 7 days before flowering to 7 days after flowering | Nipponbare | Japonica | Single | mRNA-Seq | 10.1093/gbe/evr111 |  |
| E-MTAB-11005 | Sheath_1 | Leaf sheath | 6 weeks after germination | Dongjin | Japonica | Paired | mRNA-Seq |  |  |
| E-MTAB-11005 | Sheath_2 | Leaf sheath | 6 weeks after germination | Dongjin | Japonica | Paired | mRNA-Seq |  |  |
| CRR063403 | Node_1 | Node I | 0.3cm length Node I at grain-filling stage | Xiangwanxian No.12 | Indica | Paired | mRNA-Seq | 10.1186/s12864-020-6474-7 |  |
| CRR063404 | Node_2 | Node I | 0.3cm length Node I at grain-filling stage | Xiangwanxian No.12 | Indica | Paired | mRNA-Seq | 10.1186/s12864-020-6474-7 |  |
| CRR063405 | Node_3 | Node I | 0.3cm length Node I at grain-filling stage | Xiangwanxian No.12 | Indica | Paired | mRNA-Seq | 10.1186/s12864-020-6474-7 |  |
| CRR063415 | Node_4 | Node I | 0.3cm length Node I at grain-filling stage | Yuzhenxian | Indica | Paired | mRNA-Seq | 10.1186/s12864-020-6474-7 |  |
| CRR063416 | Node_5 | Node I | 0.3cm length Node I at grain-filling stage | Yuzhenxian | Indica | Paired | mRNA-Seq | 10.1186/s12864-020-6474-7 |  |
| CRR063417 | Node_6 | Node I | 0.3cm length Node I at grain-filling stage | Yuzhenxian | Indica | Paired | mRNA-Seq | 10.1186/s12864-020-6474-7 |  |
| E-MTAB-11005 | Collar_1 | Collar | 6 weeks after germination | Dongjin | Japonica | Paired | mRNA-Seq |  |  |
| E-MTAB-11005 | Collar_2 | Collar | 6 weeks after germination | Dongjin | Japonica | Paired | mRNA-Seq |  |  |
| E-MTAB-11005 | Collar_3 | L.joint | 6 weeks after germination | Dongjin | Japonica | Paired | mRNA-Seq |  |  |
| E-MTAB-11005 | Collar_4 | L.joint | 6 weeks after germination | Dongjin | Japonica | Paired | mRNA-Seq |  |  |
| E-MTAB-11005 | Ligule_1 | Ligule | 6 weeks after germination | Dongjin | Japonica | Paired | mRNA-Seq |  |  |
| E-MTAB-11005 | Ligule_2 | Ligule | 6 weeks after germination | Dongjin | Japonica | Paired | mRNA-Seq |  |  |
| DRR001025 | Root_1 | Root | 7 days after germination | Nipponbare | Japonica | Single | mRNA-Seq | 10.1093/gbe/evr111 |  |
| DRR001032 | Root_2 | Root | 7 days after germination | Nipponbare | Japonica | Single | mRNA-Seq | 10.1093/gbe/evr111 |  |
| DRR001039 | Root_3 | Root | 7 days after germination | Nipponbare | Japonica | Single | mRNA-Seq | 10.1093/gbe/evr111 |  |
| DRR001046 | Root_4 | Root | 7 days after germination | Nipponbare | Japonica | Single | mRNA-Seq | 10.1093/gbe/evr111 |  |
| E-MTAB-11005 | Roothair_1 | Roothair | 3 days after germination on solid MSO plate | Dongjin | Japonica | Paired | mRNA-Seq |  |  |
| E-MTAB-11005 | Roothair_2 | Roothair | 3 days after germination on solid MSO plate | Dongjin | Japonica | Paired | mRNA-Seq |  |  |
| E-MTAB-11005 | Roothair_3 | Roothair | 3 days after germination on solid MSO plate | Dongjin | Japonica | Paired | mRNA-Seq |  |  |
| DRR001027 | Panicle_1 | Pre-flowering panicle | 7 days before flowering | Nipponbare | Japonica | Single | mRNA-Seq | 10.1093/gbe/evr111 |  |
| DRR001034 | Panicle_2 | Pre-flowering panicle | 7 days before flowering | Nipponbare | Japonica | Single | mRNA-Seq | 10.1093/gbe/evr111 |  |
| DRR001041 | Panicle_3 | Pre-flowering panicle | 7 days before flowering | Nipponbare | Japonica | Single | mRNA-Seq | 10.1093/gbe/evr111 |  |
| DRR001048 | Panicle_4 | Pre-flowering panicle | 7 days before flowering | Nipponbare | Japonica | Single | mRNA-Seq | 10.1093/gbe/evr111 |  |
| SRR3129121 | Anther_S3_1 | Developing anther | Anther stage 3 (spikelet length 0.5~0.6mm) | 9522 | Japonica | Paired | mRNA-Seq | 10.1093/jxb/erw361 |  |
| SRR3129122 | Anther_S3_2 | Developing anther | Anther stage 3 (spikelet length 0.5~0.6mm) | 9522 | Japonica | Paired | mRNA-Seq | 10.1093/jxb/erw361 |  |
| SRR3129123 | Anther_S3_3 | Developing anther | Anther stage 3 (spikelet length 0.5~0.6mm) | 9522 | Japonica | Paired | mRNA-Seq | 10.1093/jxb/erw361 |  |
| SRR3129124 | Anther_S5_1 | Developing anther | Anther stage 5 (spikelet length 1~1.5mm) | 9522 | Japonica | Paired | mRNA-Seq | 10.1093/jxb/erw361 |  |
| SRR3129125 | Anther_S5_2 | Developing anther | Anther stage 5 (spikelet length 1~1.5mm) | 9522 | Japonica | Paired | mRNA-Seq | 10.1093/jxb/erw361 |  |
| SRR3129126 | Anther_S5_3 | Developing anther | Anther stage 5 (spikelet length 1~1.5mm) | 9522 | Japonica | Paired | mRNA-Seq | 10.1093/jxb/erw361 |  |
| SRR3129127 | Anther_S7_1 | Developing anther | Anther stage 7 (spikelet length 2.5~3mm) | 9522 | Japonica | Paired | mRNA-Seq | 10.1093/jxb/erw361 |  |
| SRR3129128 | Anther_S7_2 | Developing anther | Anther stage 7 (spikelet length 2.5~3mm) | 9522 | Japonica | Paired | mRNA-Seq | 10.1093/jxb/erw361 |  |
| SRR3129129 | Anther_S7_3 | Developing anther | Anther stage 7 (spikelet length 2.5~3mm) | 9522 | Japonica | Paired | mRNA-Seq | 10.1093/jxb/erw361 |  |
| SRR9026265 | Anther_S9_1 | Developing anther | Anther stage 9 | Huanghuazhan | Indica | Paired | mRNA-Seq | 10.1096/pcp/pcaa025 |  |
| SRR9026264 | Anther_S9_2 | Developing anther | Anther stage 9 | Huanghuazhan | Indica | Paired | mRNA-Seq | 10.1096/pcp/pcaa025 |  |
| SRR9026267 | Anther_S9_3 | Developing anther | Anther stage 9 | Huanghuazhan | Indica | Paired | mRNA-Seq | 10.1096/pcp/pcaa025 |  |
| E-MTAB-7974 | Anther_1 | Anther | Mature Anther at anthesis time | Dongjin | Japonica | Paired | mRNA-Seq | 10.1007/s00299-022-02852-3 |  |
| E-MTAB-7974 | Anther_2 | Anther | Mature Anther at anthesis time | Dongjin | Japonica | Paired | mRNA-Seq | 10.1007/s00299-022-02852-3 |  |
| E-MTAB-7974 | Anther_3 | Anther | Mature Anther at anthesis time | Dongjin | Japonica | Paired | mRNA-Seq | 10.1007/s00299-022-02852-3 |  |
| E-MTAB-11005 | Antherwall_1 | Antherwall | Empty anther after releasing pollen | Dongjin | Japonica | Paired | mRNA-Seq | 10.1007/s00299-022-02852-3 |  |
| E-MTAB-11005 | Antherwall_2 | Antherwall | Empty anther after releasing pollen | Dongjin | Japonica | Paired | mRNA-Seq | 10.1007/s00299-022-02852-3 |  |
| E-MTAB-11005 | Antherwall_3 | Antherwall | Empty anther after releasing pollen | Dongjin | Japonica | Paired | mRNA-Seq | 10.1007/s00299-022-02852-3 |  |
| E-MTAB-7974 | Pollen_1 | Pollen | Mature pollen | Dongjin | Japonica | Paired | mRNA-Seq | 10.1007/s00299-022-02852-3 |  |
| E-MTAB-7974 | Pollen_2 | Pollen | Mature pollen | Dongjin | Japonica | Paired | mRNA-Seq | 10.1007/s00299-022-02852-3 |  |
| E-MTAB-7974 | Pollen_3 | Pollen | Mature pollen | Dongjin | Japonica | Paired | mRNA-Seq | 10.1007/s00299-022-02852-3 |  |
| DRR166409 | Sperm_1 | Sperm cell |  | Nipponbare | Japonica | Paired | mRNA-Seq | 10.1093/pcp/pcz030 |  |
| DRR166410 | Sperm_2 | Sperm cell |  | Nipponbare | Japonica | Paired | mRNA-Seq | 10.1093/pcp/pcz030 |  |
| SRR10546653 | Stigma_1 | Stigma,style | Functional megaspore (Spikelet length 3.8~4.2mm) | ZH11 | Japonica | Paired | mRNA-Seq | 10.1104/pp.19.01254 |  |
| SRR10546652 | Stigma_2 | Stigma,style | Functional megaspore (Spikelet length 3.8~4.2mm) | ZH11 | Japonica | Paired | mRNA-Seq | 10.1104/pp.19.01254 |  |
| SRR10546651 | Stigma_3 | Stigma,style | Functional megaspore (Spikelet length 3.8~4.2mm) | ZH11 | Japonica | Paired | mRNA-Seq | 10.1104/pp.19.01254 |  |
| SRR10546643 | Ovule_1 | Ovule | Mature Ovule (Spikelet length 7~8mm) | ZH11 | Japonica | Paired | mRNA-Seq | 10.1104/pp.19.01254 |  |
| SRR10546658 | Ovule_2 | Ovule | Mature Ovule (Spikelet length 7~8mm) | ZH11 | Japonica | Paired | mRNA-Seq | 10.1104/pp.19.01254 |  |
| SRR10546657 | Ovule_3 | Ovule | Mature Ovule (Spikelet length 7~8mm) | ZH11 | Japonica | Paired | mRNA-Seq | 10.1104/pp.19.01254 |  |
| DRR166400 | Egg_cell_1 | Egg cell |  | Nipponbare | Japonica | Paired | mRNA-Seq | 10.1093/pcp/pcz030 |  |
| DRR166401 | Egg_cell_2 | Egg cell |  | Nipponbare | Japonica | Paired | mRNA-Seq | 10.1093/pcp/pcz030 |  |
| DRR166402 | Egg_cell_3 | Egg cell |  | Nipponbare | Japonica | Paired | mRNA-Seq | 10.1093/pcp/pcz030 |  |
| DRR166403 | Zygote_1 | Zygote |  | Nipponbare | Japonica | Paired | mRNA-Seq | 10.1093/pcp/pcz030 |  |
| DRR166404 | Zygote_2 | Zygote |  | Nipponbare | Japonica | Paired | mRNA-Seq | 10.1093/pcp/pcz030 |  |
| DRR001030 | Seed_1 | seed |  | Nipponbare | Japonica | Single | mRNA-Seq | 10.1093/gbe/evr111 |  |
| DRR001037 | Seed_2 | seed |  | Nipponbare | Japonica | Single | mRNA-Seq | 10.1093/gbe/evr111 |  |
| DRR001044 | Seed_3 | seed |  | Nipponbare | Japonica | Single | mRNA-Seq | 10.1093/gbe/evr111 |  |
| DRR001051 | Seed_4 | seed |  | Nipponbare | Japonica | Single | mRNA-Seq | 10.1093/gbe/evr111 |  |
| SRR14301430 | Embryo_1 | Embryo | Embryo 45 DAH | Wuyujing3 | Japonica | Paired | mRNA-Seq | 10.1101/2021.04.29.441907 | Single-end data at SRA |
| SRR14301431 | Embryo_2 | Embryo | Embryo 45 DAH | Wuyujing3 | Japonica | Paired | mRNA-Seq | 10.1101/2021.04.29.441907 | Single-end data at SRA |
| SRR14301432 | Embryo_3 | Embryo | Embryo 45 DAH | Wuyujing3 | Japonica | Paired | mRNA-Seq | 10.1101/2021.04.29.441907 | Single-end data at SRA |
| SRR14301427 | Endosperm_1 | Endosperm | Endosperm upperpart 45 DAH | Wuyujing3 | Japonica | Paired | mRNA-Seq | 10.1101/2021.04.29.441907 | Single-end data at SRA |
| SRR14301428 | Endosperm_2 | Endosperm | Endosperm upperpart 45 DAH | Wuyujing3 | Japonica | Paired | mRNA-Seq | 10.1101/2021.04.29.441907 | Single-end data at SRA |
| SRR14301429 | Endosperm_3 | Endosperm | Endosperm upperpart 45 DAH | Wuyujing3 | Japonica | Paired | mRNA-Seq | 10.1101/2021.04.29.441907 | Single-end data at SRA |
| SRR14301423 | Endosperm_4 | Endosperm | Endosperm bottompart 45 DAH | Wuyujing3 | Japonica | Paired | mRNA-Seq | 10.1101/2021.04.29.441907 | Single-end data at SRA |
| SRR14301424 | Endosperm_5 | Endosperm | Endosperm bottompart 46 DAH | Wuyujing3 | Japonica | Paired | mRNA-Seq | 10.1101/2021.04.29.441907 | Single-end data at SRA |
| SRR14301425 | Endosperm_6 | Endosperm | Endosperm bottompart 47 DAH | Wuyujing3 | Japonica | Paired | mRNA-Seq | 10.1101/2021.04.29.441907 | Single-end data at SRA |
