## Supplementary material for "RE3DB: A multi-omics phylogenomics platform for rice E3 ubiquitin ligases identifies novel regulators of pollen germination": Table S2

| **Run** | **Treatment** | **Tissue_original** | **Description** | **Cultivar** | **Layout** | **Strategy** | **Source** |
| --- | --- | --- | --- | --- | --- | --- | --- |
| ERR2061590 | Cold_control_1 | 15 days seedling development stage | none | Oryza sativa Japonica Group | PAIRED | mRNA-Seq | 10.1007/s10142-018-0615-y |
| ERR2061591 | Cold_control_2 | 15 days seedling development stage | none | Oryza sativa Japonica Group | PAIRED | mRNA-Seq | 10.1007/s10142-018-0615-y |
| ERR2061592 | Cold_control_3 | 15 days seedling development stage | none | Oryza sativa Japonica Group | PAIRED | mRNA-Seq | 10.1007/s10142-018-0615-y |
| ERR2061593 | Cold_control_4 | 15 days seedling development stage | none | Oryza sativa Japonica Group | PAIRED | mRNA-Seq | 10.1007/s10142-018-0615-y |
| ERR2061594 | Cold_control_5 | 15 days seedling development stage | none | Oryza sativa Japonica Group | PAIRED | mRNA-Seq | 10.1007/s10142-018-0615-y |
| ERR2061595 | Cold_control_6 | 15 days seedling development stage | none | Oryza sativa Japonica Group | PAIRED | mRNA-Seq | 10.1007/s10142-018-0615-y |
| ERR2061596 | Cold_2h_1 | 15 days seedling development stage | cold temperature regimen2 (hour) | Oryza sativa Japonica Group | PAIRED | mRNA-Seq | 10.1007/s10142-018-0615-y |
| ERR2061597 | Cold_2h_2 | 15 days seedling development stage | cold temperature regimen2 (hour) | Oryza sativa Japonica Group | PAIRED | mRNA-Seq | 10.1007/s10142-018-0615-y |
| ERR2061598 | Cold_2h_3 | 15 days seedling development stage | cold temperature regimen2 (hour) | Oryza sativa Japonica Group | PAIRED | mRNA-Seq | 10.1007/s10142-018-0615-y |
| ERR2061599 | Cold_2h_4 | 15 days seedling development stage | cold temperature regimen2 (hour) | Oryza sativa Japonica Group | PAIRED | mRNA-Seq | 10.1007/s10142-018-0615-y |
| ERR2061600 | Cold_2h_5 | 15 days seedling development stage | cold temperature regimen2 (hour) | Oryza sativa Japonica Group | PAIRED | mRNA-Seq | 10.1007/s10142-018-0615-y |
| ERR2061601 | Cold_2h_6 | 15 days seedling development stage | cold temperature regimen2 (hour) | Oryza sativa Japonica Group | PAIRED | mRNA-Seq | 10.1007/s10142-018-0615-y |
| ERR2061602 | Cold_10h_1 | 15 days seedling development stage | cold temperature regimen10 (hour) | Oryza sativa Japonica Group | PAIRED | mRNA-Seq | 10.1007/s10142-018-0615-y |
| ERR2061603 | Cold_10h_2 | 15 days seedling development stage | cold temperature regimen10 (hour) | Oryza sativa Japonica Group | PAIRED | mRNA-Seq | 10.1007/s10142-018-0615-y |
| ERR2061604 | Cold_10h_3 | 15 days seedling development stage | cold temperature regimen10 (hour) | Oryza sativa Japonica Group | PAIRED | mRNA-Seq | 10.1007/s10142-018-0615-y |
| ERR2061605 | Cold_10h_4 | 15 days seedling development stage | cold temperature regimen10 (hour) | Oryza sativa Japonica Group | PAIRED | mRNA-Seq | 10.1007/s10142-018-0615-y |
| ERR2061606 | Cold_10h_5 | 15 days seedling development stage | cold temperature regimen10 (hour) | Oryza sativa Japonica Group | PAIRED | mRNA-Seq | 10.1007/s10142-018-0615-y |
| ERR2061607 | Cold_10h_6 | 15 days seedling development stage | cold temperature regimen10 (hour) | Oryza sativa Japonica Group | PAIRED | mRNA-Seq | 10.1007/s10142-018-0615-y |
| SRR1761788 | Drought_1 | Leaf tissue | WT-DR-1 (OsHSFA2e control) | Oryza Sativa Japonica | SINGLE | mRNA-Seq | GSE65024 |
| SRR1761789 | Drought_2 | Leaf tissue | WT-DR-2 (OsHSFA2e control) | Oryza Sativa Japonica | SINGLE | mRNA-Seq | GSE65024 |
| SRR1761780 | Drought_3 | Leaf tissue | WT-DR-1 (OsbHLH148 control) | Oryza Sativa Japonica | SINGLE | mRNA-Seq | GSE65024 |
| SRR1761781 | Drought_4 | Leaf tissue | WT-DR-2 (OsbHLH148 control) | Oryza Sativa Japonica | SINGLE | mRNA-Seq | GSE65024 |
| SRR1761528 | Drought_5 | Leaf tissue | WT-DR-1 | Oryza Sativa Japonica | SINGLE | mRNA-Seq | GSE65024 |
| SRR1761529 | Drought_6 | Leaf tissue | WT-DR-2 | Oryza Sativa Japonica | SINGLE | mRNA-Seq | GSE65024 |
| SRR1761790 | Drought_control_1 | Leaf tissue | WT-WW-1 (OsHSFA2e control) | Oryza Sativa Japonica | SINGLE | mRNA-Seq | GSE65024 |
| SRR1761791 | Drought_control_2 | Leaf tissue | WT-WW-2 (OsHSFA2e control) | Oryza Sativa Japonica | SINGLE | mRNA-Seq | GSE65024 |
| SRR1761782 | Drought_control_3 | Leaf tissue | WT-WW-1 (OsbHLH148 control) | Oryza Sativa Japonica | SINGLE | mRNA-Seq | GSE65024 |
| SRR1761783 | Drought_control_4 | Leaf tissue | WT-WW-2 (OsbHLH148 control) | Oryza Sativa Japonica | SINGLE | mRNA-Seq | GSE65024 |
| SRR1761530 | Drought_control_5 | Leaf tissue | WT-WW-1 | Oryza Sativa Japonica | SINGLE | mRNA-Seq | GSE65024 |
| SRR1761531 | Drought_control_6 | Leaf tissue | WT-WW-2 | Oryza Sativa Japonica | SINGLE | mRNA-Seq | GSE65024 |
| SRR5134063 | Drought_R_control_1 | Root, 4weeks seedling | Control_Replicate 1 | Oryza Sativa Japonica | PAIRED | mRNA-Seq | 10.3389/fpls.2017.00580 |
| SRR5134064 | Drought_R_control_2 | Root, 5weeks seedling | Control_Replicate 2 | Oryza Sativa Japonica | PAIRED | mRNA-Seq | 10.3389/fpls.2017.00580 |
| SRR5134065 | Drought_R_2day_1 | Root, 6weeks seedling | Drought_2 days_Replicate 1 | Oryza Sativa Japonica | PAIRED | mRNA-Seq | 10.3389/fpls.2017.00580 |
| SRR5134066 | Drought_R_2day_2 | Root, 7weeks seedling | Drought_2 days_Replicate 2 | Oryza Sativa Japonica | PAIRED | mRNA-Seq | 10.3389/fpls.2017.00580 |
| SRR5134067 | Drought_R_3day_1 | Root, 8weeks seedling | Drought_3 days_Replicate 1 | Oryza Sativa Japonica | PAIRED | mRNA-Seq | 10.3389/fpls.2017.00580 |
| SRR5134068 | Drought_R_3day_2 | Root, 7weeks seedling | Drought_3 days_Replicate 2 | Oryza Sativa Japonica | PAIRED | mRNA-Seq | 10.3389/fpls.2017.00580 |
| SRR2931040 | CONTROL_15min | Rice leaf | ID001 | Oryza sativa Japonica Group | PAIRED | mRNA-Seq | 10.1105/tpc.16.00158 |
| SRR2931041 | CONTROL_15min | Rice leaf | ID145 | Oryza sativa Japonica Group | PAIRED | mRNA-Seq | 10.1105/tpc.16.00158 |
| SRR2931075 | HEAT_15min_1 | Rice leaf | ID360 | Oryza sativa Japonica Group | PAIRED | mRNA-Seq | 10.1105/tpc.16.00158 |
| SRR2931076 | HEAT_15min_2 | Rice leaf | ID491 | Oryza sativa Japonica Group | PAIRED | mRNA-Seq | 10.1105/tpc.16.00158 |
| SRR2931077 | HEAT_30min_1 | Rice leaf | ID361-2 | Oryza sativa Japonica Group | PAIRED | mRNA-Seq | 10.1105/tpc.16.00158 |
| SRR2931078 | HEAT_30min_2 | Rice leaf | ID492-2 | Oryza sativa Japonica Group | PAIRED | mRNA-Seq | 10.1105/tpc.16.00158 |
| SRR2931079 | HEAT_45min_1 | Rice leaf | ID362 | Oryza sativa Japonica Group | PAIRED | mRNA-Seq | 10.1105/tpc.16.00158 |
| SRR2931080 | HEAT_45min_2 | Rice leaf | ID493-2 | Oryza sativa Japonica Group | PAIRED | mRNA-Seq | 10.1105/tpc.16.00158 |
| SRR2931081 | HEAT_60min_1 | Rice leaf | ID363-2 | Oryza sativa Japonica Group | PAIRED | mRNA-Seq | 10.1105/tpc.16.00158 |
| SRR2931082 | HEAT_60min_2 | Rice leaf | ID494-2 | Oryza sativa Japonica Group | PAIRED | mRNA-Seq | 10.1105/tpc.16.00158 |
| SRR2931083 | HEAT_75min_1 | Rice leaf | ID364 | Oryza sativa Japonica Group | PAIRED | mRNA-Seq | 10.1105/tpc.16.00158 |
| SRR2931084 | HEAT_75min_2 | Rice leaf | ID495 | Oryza sativa Japonica Group | PAIRED | mRNA-Seq | 10.1105/tpc.16.00158 |
| SRR2931085 | HEAT_90min_1 | Rice leaf | ID365-2 | Oryza sativa Japonica Group | PAIRED | mRNA-Seq | 10.1105/tpc.16.00158 |
| SRR2931086 | HEAT_90min_2 | Rice leaf | ID496-2 | Oryza sativa Japonica Group | PAIRED | mRNA-Seq | 10.1105/tpc.16.00158 |
| SRR2931087 | HEAT_105min_1 | Rice leaf | ID366 | Oryza sativa Japonica Group | PAIRED | mRNA-Seq | 10.1105/tpc.16.00158 |
| SRR2931088 | HEAT_105min_2 | Rice leaf | ID497 | Oryza sativa Japonica Group | PAIRED | mRNA-Seq | 10.1105/tpc.16.00158 |
| SRR2931089 | HEAT_120min_1 | Rice leaf | ID367-2 | Oryza sativa Japonica Group | PAIRED | mRNA-Seq | 10.1105/tpc.16.00158 |
| SRR2931090 | HEAT_120min_2 | Rice leaf | ID498-2 | Oryza sativa Japonica Group | PAIRED | mRNA-Seq | 10.1105/tpc.16.00158 |
| SRR2931091 | HEAT_135min_1 | Rice leaf | ID368 | Oryza sativa Japonica Group | PAIRED | mRNA-Seq | 10.1105/tpc.16.00158 |
| SRR2931092 | HEAT_135min_2 | Rice leaf | ID499 | Oryza sativa Japonica Group | PAIRED | mRNA-Seq | 10.1105/tpc.16.00158 |
| SRR2931093 | HEAT_150min_1 | Rice leaf | ID369 | Oryza sativa Japonica Group | PAIRED | mRNA-Seq | 10.1105/tpc.16.00158 |
| SRR2931094 | HEAT_150min_2 | Rice leaf | ID500 | Oryza sativa Japonica Group | PAIRED | mRNA-Seq | 10.1105/tpc.16.00158 |
| SRR2931095 | HEAT_165min_1 | Rice leaf | ID370 | Oryza sativa Japonica Group | PAIRED | mRNA-Seq | 10.1105/tpc.16.00158 |
| SRR2931096 | HEAT_165min_2 | Rice leaf | ID501 | Oryza sativa Japonica Group | PAIRED | mRNA-Seq | 10.1105/tpc.16.00158 |
| SRR2931097 | HEAT_180min_1 | Rice leaf | ID068-2 | Oryza sativa Japonica Group | PAIRED | mRNA-Seq | 10.1105/tpc.16.00158 |
| SRR2931098 | HEAT_180min_2 | Rice leaf | ID371 | Oryza sativa Japonica Group | PAIRED | mRNA-Seq | 10.1105/tpc.16.00158 |
| SRR2931099 | HEAT_195min_1 | Rice leaf | ID469 | Oryza sativa Japonica Group | PAIRED | mRNA-Seq | 10.1105/tpc.16.00158 |
| SRR2931100 | HEAT_195min_2 | Rice leaf | ID803 | Oryza sativa Japonica Group | PAIRED | mRNA-Seq | 10.1105/tpc.16.00158 |
| SRR2931101 | HEAT_210min_1 | Rice leaf | ID470 | Oryza sativa Japonica Group | PAIRED | mRNA-Seq | 10.1105/tpc.16.00158 |
| SRR2931102 | HEAT_210min_2 | Rice leaf | ID504 | Oryza sativa Japonica Group | PAIRED | mRNA-Seq | 10.1105/tpc.16.00158 |
| SRR2931103 | HEAT_225min_1 | Rice leaf | ID374 | Oryza sativa Japonica Group | PAIRED | mRNA-Seq | 10.1105/tpc.16.00158 |
| SRR2931104 | HEAT_225min_2 | Rice leaf | ID505 | Oryza sativa Japonica Group | PAIRED | mRNA-Seq | 10.1105/tpc.16.00158 |
| SRR2931105 | HEAT_240min_1 | Rice leaf | ID072-2 | Oryza sativa Japonica Group | PAIRED | mRNA-Seq | 10.1105/tpc.16.00158 |
| SRR2931106 | HEAT_240min_2 | Rice leaf | ID375 | Oryza sativa Japonica Group | PAIRED | mRNA-Seq | 10.1105/tpc.16.00158 |
| ERR266228 | salt_control_1h_1 | seedling, two leaves visible, three leaves visible | nippon_control_1hr_rep1 | Oryza Sativa Japonica | SINGLE | mRNA-Seq | E-MTAB-1625 |
| ERR266233 | salt_control_1h_2 | seedling, two leaves visible, three leaves visible | nippon_control_1hr_rep2 | Oryza Sativa Japonica | SINGLE | mRNA-Seq | E-MTAB-1625 |
| ERR266230 | salt_control_1h_3 | seedling, two leaves visible, three leaves visible | nippon_control_1hr_rep3 | Oryza Sativa Japonica | SINGLE | mRNA-Seq | E-MTAB-1625 |
| ERR266225 | salt_control_24h_1 | seedling, two leaves visible, three leaves visible | nippon_control_24hr_rep1 | Oryza Sativa Japonica | SINGLE | mRNA-Seq | E-MTAB-1625 |
| ERR266234 | salt_control_24h_2 | seedling, two leaves visible, three leaves visible | nippon_control_24hr_rep2 | Oryza Sativa Japonica | SINGLE | mRNA-Seq | E-MTAB-1625 |
| ERR266232 | salt_control_24h_3 | seedling, two leaves visible, three leaves visible | nippon_control_24hr_rep3 | Oryza Sativa Japonica | SINGLE | mRNA-Seq | E-MTAB-1625 |
| ERR266229 | salt_control_5h_1 | seedling, two leaves visible, three leaves visible | nippon_control_5hr_rep1 | Oryza Sativa Japonica | SINGLE | mRNA-Seq | E-MTAB-1625 |
| ERR266223 | salt_control_5h_2 | seedling, two leaves visible, three leaves visible | nippon_control_5hr_rep2 | Oryza Sativa Japonica | SINGLE | mRNA-Seq | E-MTAB-1625 |
| ERR266222 | salt_control_5h_3 | seedling, two leaves visible, three leaves visible | nippon_control_5hr_rep3 | Oryza Sativa Japonica | SINGLE | mRNA-Seq | E-MTAB-1625 |
| ERR266237 | salt_1h_1 | seedling, two leaves visible, three leaves visible | nippon_salt_1hr_rep1 | Oryza Sativa Japonica | SINGLE | mRNA-Seq | E-MTAB-1625 |
| ERR266236 | salt_1h_2 | seedling, two leaves visible, three leaves visible | nippon_salt_1hr_rep2 | Oryza Sativa Japonica | SINGLE | mRNA-Seq | E-MTAB-1625 |
| ERR266235 | salt_1h_3 | seedling, two leaves visible, three leaves visible | nippon_salt_1hr_rep3 | Oryza Sativa Japonica | SINGLE | mRNA-Seq | E-MTAB-1625 |
| ERR266238 | salt_24h_1 | seedling, two leaves visible, three leaves visible | nippon_salt_24hr_rep1 | Oryza Sativa Japonica | SINGLE | mRNA-Seq | E-MTAB-1625 |
| ERR266226 | salt_24h_2 | seedling, two leaves visible, three leaves visible | nippon_salt_24hr_rep2 | Oryza Sativa Japonica | SINGLE | mRNA-Seq | E-MTAB-1625 |
| ERR266231 | salt_24h_3 | seedling, two leaves visible, three leaves visible | nippon_salt_24hr_rep3 | Oryza Sativa Japonica | SINGLE | mRNA-Seq | E-MTAB-1625 |
| ERR266227 | salt_5h_1 | seedling, two leaves visible, three leaves visible | nippon_salt_5hr_rep1 | Oryza Sativa Japonica | SINGLE | mRNA-Seq | E-MTAB-1625 |
| ERR266224 | salt_5h_2 | seedling, two leaves visible, three leaves visible | nippon_salt_5hr_rep2 | Oryza Sativa Japonica | SINGLE | mRNA-Seq | E-MTAB-1625 |
| ERR266221 | salt_5h_3 | seedling, two leaves visible, three leaves visible | nippon_salt_5hr_rep3 | Oryza Sativa Japonica | SINGLE | mRNA-Seq | E-MTAB-1625 |
| ERR986071 | normal_air_1 | 0 seed germination stage | normal air_1 | Oryza Sativa (mixed) | SINGLE | mRNA-Seq | 10.3389/fpls.2017.00762 |
| ERR986073 | normal_air_2 | 0 seed germination stage | normal air_2 | Oryza Sativa (mixed) | SINGLE | mRNA-Seq | 10.3389/fpls.2017.00762 |
| ERR986076 | normal_air_3 | 0 seed germination stage | normal air_3 | Oryza Sativa (mixed) | SINGLE | mRNA-Seq | 10.3389/fpls.2017.00762 |
| ERR986078 | normal_air_4 | 0 seed germination stage | normal air_4 | Oryza Sativa (mixed) | SINGLE | mRNA-Seq | 10.3389/fpls.2017.00762 |
| ERR986080 | normal_air_5 | 0 seed germination stage | normal air_5 | Oryza Sativa (mixed) | SINGLE | mRNA-Seq | 10.3389/fpls.2017.00762 |
| ERR986082 | normal_air_6 | 0 seed germination stage | normal air_6 | Oryza Sativa (mixed) | SINGLE | mRNA-Seq | 10.3389/fpls.2017.00762 |
| ERR986072 | submergence_1 | 0 seed germination stage | submergence_1 | Oryza Sativa (mixed) | SINGLE | mRNA-Seq | 10.3389/fpls.2017.00762 |
| ERR986074 | submergence_2 | 0 seed germination stage | submergence_2 | Oryza Sativa (mixed) | SINGLE | mRNA-Seq | 10.3389/fpls.2017.00762 |
| ERR986075 | submergence_3 | 0 seed germination stage | submergence_3 | Oryza Sativa (mixed) | SINGLE | mRNA-Seq | 10.3389/fpls.2017.00762 |
| ERR986077 | submergence_4 | 0 seed germination stage | submergence_4 | Oryza Sativa (mixed) | SINGLE | mRNA-Seq | 10.3389/fpls.2017.00762 |
| ERR986079 | submergence_5 | 0 seed germination stage | submergence_5 | Oryza Sativa (mixed) | SINGLE | mRNA-Seq | 10.3389/fpls.2017.00762 |
| ERR986081 | submergence_6 | 0 seed germination stage | submergence_6 | Oryza Sativa (mixed) | SINGLE | mRNA-Seq | 10.3389/fpls.2017.00762 |
