## Supplementary material for "RE3DB: A multi-omics phylogenomics platform for rice E3 ubiquitin ligases identifies novel regulators of pollen germination": Table S3

| **Run** | **Treatment** | **Tissue_original** | **Description** | **Cultivar** | **Layout** | **Strategy** | **Source** |
| --- | --- | --- | --- | --- | --- | --- | --- |
| SRR561003 | B.glumae_Mock_1 | Seedling | Rice inoculation at the seedling stage | Oryza sativa f. spontanea | SINGLE | mRNA-Seq | https://doi.org/10.1186/1471-2164-15-755 |
| SRR638899 | B.glumae_Mock_2 | Seedling | Rice inoculation at the seedling stage | Oryza sativa f. spontanea | SINGLE | mRNA-Seq | https://doi.org/10.1186/1471-2164-15-755 |
| SRR638900 | B.glumae_Mock_3 | Seedling | Rice inoculation at the seedling stage | Oryza sativa f. spontanea | SINGLE | mRNA-Seq | https://doi.org/10.1186/1471-2164-15-755 |
| SRR638911 | B.glumae_1 | Seedling | Rice inoculation at the seedling stage | Oryza sativa f. spontanea | SINGLE | mRNA-Seq | https://doi.org/10.1186/1471-2164-15-755 |
| SRR638913 | B.glumae_2 | Seedling | Rice inoculation at the seedling stage | Oryza sativa f. spontanea | SINGLE | mRNA-Seq | https://doi.org/10.1186/1471-2164-15-755 |
| SRR638916 | B.glumae_3 | Seedling | Rice inoculation at the seedling stage | Oryza sativa f. spontanea | SINGLE | mRNA-Seq | https://doi.org/10.1186/1471-2164-15-755 |
| SRR1288367 | H.oryzae_3days_1 | 18days shoot | Shoots of Ho-infected plants, 3 days post infection, replicate 1 | Oryza sativa Japonica Group | PAIRED | mRNA-Seq | https://doi.org/10.1371/journal.pone.0106858 |
| SRR1288368 | H.oryzae_3days_2 | 18days shoot | Shoots of Ho-infected plants, 3 days post infection, replicate 2 | Oryza sativa Japonica Group | PAIRED | mRNA-Seq | https://doi.org/10.1371/journal.pone.0106858 |
| SRR1288369 | H.oryzae_7days_1 | 18days shoot | Shoots of Ho-infected plants, 7 days post infection, replicate 1 | Oryza sativa Japonica Group | PAIRED | mRNA-Seq | https://doi.org/10.1371/journal.pone.0106858 |
| SRR1288370 | H.oryzae_7days_2 | 18days shoot | Shoots of Ho-infected plants, 7 days post infection, replicate 2 | Oryza sativa Japonica Group | PAIRED | mRNA-Seq | https://doi.org/10.1371/journal.pone.0106858 |
| SRR1288363 | H.oryzae_Mock_3days_1 | 18days shoot | Shoots of uninfected plants, 3 days post infection, replicate 1 | Oryza sativa Japonica Group | PAIRED | mRNA-Seq | https://doi.org/10.1371/journal.pone.0106858 |
| SRR1288364 | H.oryzae_Mock_3days_2 | 18days shoot | Shoots of uninfected plants, 3 days post infection, replicate 2 | Oryza sativa Japonica Group | PAIRED | mRNA-Seq | https://doi.org/10.1371/journal.pone.0106858 |
| SRR1288365 | H.oryzae_Mock_7days_1 | 18days shoot | Shoots of uninfected plants, 7 days post infection, replicate 1 | Oryza sativa Japonica Group | PAIRED | mRNA-Seq | https://doi.org/10.1371/journal.pone.0106858 |
| SRR1288366 | H.oryzae_Mock_7days_2 | 18days shoot | Shoots of uninfected plants, 7 days post infection, replicate 2 | Oryza sativa Japonica Group | PAIRED | mRNA-Seq | https://doi.org/10.1371/journal.pone.0106858 |
| SRR5873827 | R.solani_12h_1 | leaf | the rice leaf induced by the AG1 IA strain of R. solani at 12h |  | PAIRED | mRNA-Seq | 10.3389/fpls.2017.01422 |
| SRR5873828 | R.solani_12h_2 | leaf | the rice leaf induced by the AG1 IA strain of R. solani at 12h |  | PAIRED | mRNA-Seq | 10.3389/fpls.2017.01422 |
| SRR5873829 | R.solani_12h_3 | leaf | the rice leaf induced by the AG1 IA strain of R. solani at 12h |  | PAIRED | mRNA-Seq | 10.3389/fpls.2017.01422 |
| SRR5873822 | R.solani_24h_1 | leaf | the rice leaf induced by the AG1 IA strain of R. solani at 24h |  | PAIRED | mRNA-Seq | 10.3389/fpls.2017.01422 |
| SRR5873823 | R.solani_24h_2 | leaf | the rice leaf induced by the AG1 IA strain of R. solani at 24h |  | PAIRED | mRNA-Seq | 10.3389/fpls.2017.01422 |
| SRR5873824 | R.solani_24h_3 | leaf | the rice leaf induced by the AG1 IA strain of R. solani at 24h |  | PAIRED | mRNA-Seq | 10.3389/fpls.2017.01422 |
| SRR5873825 | R.solani_36h_1 | leaf | the rice leaf induced by the AG1 IA strain of R. solani at 36h |  | PAIRED | mRNA-Seq | 10.3389/fpls.2017.01422 |
| SRR5873830 | R.solani_36h_2 | leaf | the rice leaf induced by the AG1 IA strain of R. solani at 36h |  | PAIRED | mRNA-Seq | 10.3389/fpls.2017.01422 |
| SRR5873831 | R.solani_36h_3 | leaf | the rice leaf induced by the AG1 IA strain of R. solani at 36h |  | PAIRED | mRNA-Seq | 10.3389/fpls.2017.01422 |
| SRR5873807 | R.solani_48h_1 | leaf | the rice leaf induced by the AG1 IA strain of R. solani at 48h |  | PAIRED | mRNA-Seq | 10.3389/fpls.2017.01422 |
| SRR5873806 | R.solani_48h_2 | leaf | the rice leaf induced by the AG1 IA strain of R. solani at 48h |  | PAIRED | mRNA-Seq | 10.3389/fpls.2017.01422 |
| SRR5873811 | R.solani_48h_3 | leaf | the rice leaf induced by the AG1 IA strain of R. solani at 48h |  | PAIRED | mRNA-Seq | 10.3389/fpls.2017.01422 |
| SRR5873810 | R.solani_72h_1 | leaf | the rice leaf induced by the AG1 IA strain of R. solani at 72h |  | PAIRED | mRNA-Seq | 10.3389/fpls.2017.01422 |
| SRR5873809 | R.solani_72h_2 | leaf | the rice leaf induced by the AG1 IA strain of R. solani at 72h |  | PAIRED | mRNA-Seq | 10.3389/fpls.2017.01422 |
| SRR5873808 | R.solani_72h_3 | leaf | the rice leaf induced by the AG1 IA strain of R. solani at 72h |  | PAIRED | mRNA-Seq | 10.3389/fpls.2017.01422 |
| SRR5873805 | R.solani_MOCK_1 | leaf | the rice leaf if control |  | PAIRED | mRNA-Seq | 10.3389/fpls.2017.01422 |
| SRR5873804 | R.solani_MOCK_2 | leaf | the rice leaf if control |  | PAIRED | mRNA-Seq | 10.3389/fpls.2017.01422 |
| SRR5873826 | R.solani_MOCK_3 | leaf | the rice leaf if control |  | PAIRED | mRNA-Seq | 10.3389/fpls.2017.01422 |
| SRR950091 | RSV_MOCK | 3leaf stage seedling | Gene expression of a rice stripe virus susceptible cultivar in three-leaves stage | Oryza sativa Japonica Group | PAIRED | mRNA-Seq | 10.1371/journal.pone.0082126 |
| SRR949614 | RSV | 3leaf stage seedling | Gene exression of a rice stripe virus susceptible cultivar after being co-cultured with small black planthopper carried RSV for 48h | Oryza sativa Japonica Group | PAIRED | mRNA-Seq | 10.1371/journal.pone.0082126 |
| SRR5909330 | RVD_MOCK_1 | 42 days old seedling | 42 days old seedling (4 weeks post inoculation) & wild type (inoculated with virus free insect) |  | PAIRED | mRNA-Seq | https://elifesciences.org/articles/27529 |
| SRR5909331 | RVD_MOCK_2 | 42 days old seedling | 42 days old seedling (4 weeks post inoculation) & wild type (inoculated with virus free insect) |  | PAIRED | mRNA-Seq | https://elifesciences.org/articles/27529 |
| SRR5909332 | RVD_MOCK_3 | 42 days old seedling | 42 days old seedling (4 weeks post inoculation) & wild type (inoculated with virus free insect) |  | PAIRED | mRNA-Seq | https://elifesciences.org/articles/27529 |
| SRR5909333 | RVD_1 | 42 days old seedling | 42 days old seedling (4 weeks post inoculation) & RDV-infected (inoculated with insect carrying RDV) |  | PAIRED | mRNA-Seq | https://elifesciences.org/articles/27529 |
| SRR5909334 | RVD_2 | 42 days old seedling | 42 days old seedling (4 weeks post inoculation) & RDV-infected (inoculated with insect carrying RDV) |  | PAIRED | mRNA-Seq | https://elifesciences.org/articles/27529 |
| SRR5909335 | RVD_3 | 42 days old seedling | 42 days old seedling (4 weeks post inoculation) & RDV-infected (inoculated with insect carrying RDV) |  | PAIRED | mRNA-Seq | https://elifesciences.org/articles/27529 |
| SRR1952778 | X.Oryzae_BLS256_1 | Leaf | 48 hrs after inoculation with BLS256 | Oryza sativa Japonica Group | SINGLE | mRNA-Seq | https://doi.org/10.3389/fpls.2015.00536 |
| SRR1952779 | X.Oryzae_BLS256_2 | Leaf | 48 hrs after inoculation with BLS256 | Oryza sativa Japonica Group | SINGLE | mRNA-Seq | https://doi.org/10.3389/fpls.2015.00536 |
| SRR1952780 | X.Oryzae_BLS256_3 | Leaf | 48 hrs after inoculation with BLS256 | Oryza sativa Japonica Group | SINGLE | mRNA-Seq | https://doi.org/10.3389/fpls.2015.00536 |
| SRR1952781 | X.Oryzae_BLS279_1 | Leaf | 48 hrs after inoculation with BLS279 | Oryza sativa Japonica Group | SINGLE | mRNA-Seq | https://doi.org/10.3389/fpls.2015.00536 |
| SRR1952782 | X.Oryzae_BLS279_2 | Leaf | 48 hrs after inoculation with BLS279 | Oryza sativa Japonica Group | SINGLE | mRNA-Seq | https://doi.org/10.3389/fpls.2015.00536 |
| SRR1952783 | X.Oryzae_BLS279_3 | Leaf | 48 hrs after inoculation with BLS279 | Oryza sativa Japonica Group | SINGLE | mRNA-Seq | https://doi.org/10.3389/fpls.2015.00536 |
| SRR1952784 | X.Oryzae_CFBP2286_1 | Leaf | 48 hrs after inoculation with CFBP2286 | Oryza sativa Japonica Group | SINGLE | mRNA-Seq | https://doi.org/10.3389/fpls.2015.00536 |
| SRR1952785 | X.Oryzae_CFBP2286_2 | Leaf | 48 hrs after inoculation with CFBP2286 | Oryza sativa Japonica Group | SINGLE | mRNA-Seq | https://doi.org/10.3389/fpls.2015.00536 |
| SRR1952786 | X.Oryzae_CFBP2286_3 | Leaf | 48 hrs after inoculation with CFBP2286 | Oryza sativa Japonica Group | SINGLE | mRNA-Seq | https://doi.org/10.3389/fpls.2015.00536 |
| SRR1952787 | X.Oryzae_B8-12_1 | Leaf | 48 hrs after inoculation with B8-12 | Oryza sativa Japonica Group | SINGLE | mRNA-Seq | https://doi.org/10.3389/fpls.2015.00536 |
| SRR1952788 | X.Oryzae_B8-12_2 | Leaf | 48 hrs after inoculation with B8-12 | Oryza sativa Japonica Group | SINGLE | mRNA-Seq | https://doi.org/10.3389/fpls.2015.00536 |
| SRR1952789 | X.Oryzae_B8-12_3 | Leaf | 48 hrs after inoculation with B8-12 | Oryza sativa Japonica Group | SINGLE | mRNA-Seq | https://doi.org/10.3389/fpls.2015.00536 |
| SRR1952790 | X.Oryzae_L8_1 | Leaf | 48 hrs after inoculation with L8 | Oryza sativa Japonica Group | SINGLE | mRNA-Seq | https://doi.org/10.3389/fpls.2015.00536 |
| SRR1952791 | X.Oryzae_L8_2 | Leaf | 48 hrs after inoculation with L8 | Oryza sativa Japonica Group | SINGLE | mRNA-Seq | https://doi.org/10.3389/fpls.2015.00536 |
| SRR1952792 | X.Oryzae_L8_3 | Leaf | 48 hrs after inoculation with L8 | Oryza sativa Japonica Group | SINGLE | mRNA-Seq | https://doi.org/10.3389/fpls.2015.00536 |
| SRR1952793 | X.Oryzae_RS105_1 | Leaf | 48 hrs after inoculation with RS105 | Oryza sativa Japonica Group | SINGLE | mRNA-Seq | https://doi.org/10.3389/fpls.2015.00536 |
| SRR1952794 | X.Oryzae_RS105_2 | Leaf | 48 hrs after inoculation with RS105 | Oryza sativa Japonica Group | SINGLE | mRNA-Seq | https://doi.org/10.3389/fpls.2015.00536 |
| SRR1952795 | X.Oryzae_RS105_3 | Leaf | 48 hrs after inoculation with RS105 | Oryza sativa Japonica Group | SINGLE | mRNA-Seq | https://doi.org/10.3389/fpls.2015.00536 |
| SRR1952796 | X.Oryzae_BXOR1_1 | Leaf | 48 hrs after inoculation with BXOR1 | Oryza sativa Japonica Group | SINGLE | mRNA-Seq | https://doi.org/10.3389/fpls.2015.00536 |
| SRR1952797 | X.Oryzae_BXOR1_2 | Leaf | 48 hrs after inoculation with BXOR1 | Oryza sativa Japonica Group | SINGLE | mRNA-Seq | https://doi.org/10.3389/fpls.2015.00536 |
| SRR1952798 | X.Oryzae_BXOR1_3 | Leaf | 48 hrs after inoculation with BXOR1 | Oryza sativa Japonica Group | SINGLE | mRNA-Seq | https://doi.org/10.3389/fpls.2015.00536 |
| SRR1952799 | X.Oryzae_CFBP7331_1 | Leaf | 48 hrs after inoculation with CFBP7331 | Oryza sativa Japonica Group | SINGLE | mRNA-Seq | https://doi.org/10.3389/fpls.2015.00536 |
| SRR1952800 | X.Oryzae_CFBP7331_2 | Leaf | 48 hrs after inoculation with CFBP7331 | Oryza sativa Japonica Group | SINGLE | mRNA-Seq | https://doi.org/10.3389/fpls.2015.00536 |
| SRR1952801 | X.Oryzae_CFBP7331_3 | Leaf | 48 hrs after inoculation with CFBP7331 | Oryza sativa Japonica Group | SINGLE | mRNA-Seq | https://doi.org/10.3389/fpls.2015.00536 |
| SRR1952802 | X.Oryzae_CFBP7341_1 | Leaf | 48 hrs after inoculation with CFBP7341 | Oryza sativa Japonica Group | SINGLE | mRNA-Seq | https://doi.org/10.3389/fpls.2015.00536 |
| SRR1952803 | X.Oryzae_CFBP7341_2 | Leaf | 48 hrs after inoculation with CFBP7341 | Oryza sativa Japonica Group | SINGLE | mRNA-Seq | https://doi.org/10.3389/fpls.2015.00536 |
| SRR1952804 | X.Oryzae_CFBP7341_3 | Leaf | 48 hrs after inoculation with CFBP7341 | Oryza sativa Japonica Group | SINGLE | mRNA-Seq | https://doi.org/10.3389/fpls.2015.00536 |
| SRR1952805 | X.Oryzae_CFBP7342_1 | Leaf | 48 hrs after inoculation with CFBP7342 | Oryza sativa Japonica Group | SINGLE | mRNA-Seq | https://doi.org/10.3389/fpls.2015.00536 |
| SRR1952806 | X.Oryzae_CFBP7342_2 | Leaf | 48 hrs after inoculation with CFBP7342 | Oryza sativa Japonica Group | SINGLE | mRNA-Seq | https://doi.org/10.3389/fpls.2015.00536 |
| SRR1952807 | X.Oryzae_CFBP7342_3 | Leaf | 48 hrs after inoculation with CFBP7342 | Oryza sativa Japonica Group | SINGLE | mRNA-Seq | https://doi.org/10.3389/fpls.2015.00536 |
| SRR1952808 | Mock_1 | Leaf | 48 hrs after mock-inoculation | Oryza sativa Japonica Group | SINGLE | mRNA-Seq | https://doi.org/10.3389/fpls.2015.00536 |
| SRR1952809 | Mock_2 | Leaf | 48 hrs after mock-inoculation | Oryza sativa Japonica Group | SINGLE | mRNA-Seq | https://doi.org/10.3389/fpls.2015.00536 |
| SRR1952810 | Mock_3 | Leaf | 48 hrs after mock-inoculation | Oryza sativa Japonica Group | SINGLE | mRNA-Seq | https://doi.org/10.3389/fpls.2015.00536 |
