## Supplementary material for "RE3DB: A multi-omics phylogenomics platform for rice E3 ubiquitin ligases identifies novel regulators of pollen germination": Table S4

| **Run** | **Treatment** | **Tissue_original** | **Description** | **Cultivar** | **Layout** | **Strategy** | **Source** |
| --- | --- | --- | --- | --- | --- | --- | --- |
| SRR1562061 | Nitro_S_0h_1 | treated by N sufficient solution | root | Oryza Sativa Japonica | PAIRED | mRNA-Seq | 10.1186/s12870-015-0425-5 |
| SRR1562069 | Nitro_S_0h_2 | treated by N sufficient solution | root | Oryza Sativa Japonica | PAIRED | mRNA-Seq | 10.1186/s12870-015-0425-5 |
| SRR1562070 | Nitro_D_12h_1 | treated by N deficient solution | root | Oryza Sativa Japonica | PAIRED | mRNA-Seq | 10.1186/s12870-015-0425-5 |
| SRR1562072 | Nitro_D_12h_2 | treated by N deficient solution | root | Oryza Sativa Japonica | PAIRED | mRNA-Seq | 10.1186/s12870-015-0425-5 |
| SRR1005290 | Pi_21d_R_control_1 | grown in 0.323 millimolar sodium dihydrogenphosphate for 21 days | root | Oryza Sativa Japonica | PAIRED | mRNA-Seq | 10.1105/tpc.113.117325 |
| SRR1005291 | Pi_21d_R_control_2 | grown in 0.323 millimolar sodium dihydrogenphosphate for 21 days | root | Oryza Sativa Japonica | PAIRED | mRNA-Seq | 10.1105/tpc.113.117325 |
| SRR1005292 | Pi_21d_R_control_3 | grown in 0.323 millimolar sodium dihydrogenphosphate for 21 days | root | Oryza Sativa Japonica | PAIRED | mRNA-Seq | 10.1105/tpc.113.117325 |
| SRR1005293 | Pi_21d_R_Recov_0h_1 | grown without sodium dihydrogenphosphate for 21 days | root | Oryza Sativa Japonica | PAIRED | mRNA-Seq | 10.1105/tpc.113.117325 |
| SRR1005294 | Pi_21d_R_Recov_0h_2 | grown without sodium dihydrogenphosphate for 21 days | root | Oryza Sativa Japonica | PAIRED | mRNA-Seq | 10.1105/tpc.113.117325 |
| SRR1005295 | Pi_21d_R_Recov_0h_3 | grown without sodium dihydrogenphosphate for 21 days | root | Oryza Sativa Japonica | PAIRED | mRNA-Seq | 10.1105/tpc.113.117325 |
| SRR1005269 | Pi_21d_R_Recov_1h_1 | grown without sodium dihydrogenphosphate for 21 days and then grown in grown in 0.323 millimolar sodium dihydrogenphosphate for 1 hour | root | Oryza Sativa Japonica | PAIRED | mRNA-Seq | 10.1105/tpc.113.117325 |
| SRR1005270 | Pi_21d_R_Recov_1h_2 | grown without sodium dihydrogenphosphate for 21 days and then grown in grown in 0.323 millimolar sodium dihydrogenphosphate for 1 hour | root | Oryza Sativa Japonica | PAIRED | mRNA-Seq | 10.1105/tpc.113.117325 |
| SRR1005271 | Pi_21d_R_Recov_1h_3 | grown without sodium dihydrogenphosphate for 21 days and then grown in grown in 0.323 millimolar sodium dihydrogenphosphate for 1 hour | root | Oryza Sativa Japonica | PAIRED | mRNA-Seq | 10.1105/tpc.113.117325 |
| SRR1005278 | Pi_21d_R_Recov_24h_1 | grown without sodium dihydrogenphosphate for 21 days and then grown in grown in 0.323 millimolar sodium dihydrogenphosphate for 24 hours | root | Oryza Sativa Japonica | PAIRED | mRNA-Seq | 10.1105/tpc.113.117325 |
| SRR1005279 | Pi_21d_R_Recov_24h_2 | grown without sodium dihydrogenphosphate for 21 days and then grown in grown in 0.323 millimolar sodium dihydrogenphosphate for 24 hours | root | Oryza Sativa Japonica | PAIRED | mRNA-Seq | 10.1105/tpc.113.117325 |
| SRR1005280 | Pi_21d_R_Recov_24h_3 | grown without sodium dihydrogenphosphate for 21 days and then grown in grown in 0.323 millimolar sodium dihydrogenphosphate for 24 hours | root | Oryza Sativa Japonica | PAIRED | mRNA-Seq | 10.1105/tpc.113.117325 |
| SRR1005287 | Pi_21d_R_Recov_6h_1 | grown without sodium dihydrogenphosphate for 21 days and then grown in grown in 0.323 millimolar sodium dihydrogenphosphate for 6 hours | root | Oryza Sativa Japonica | PAIRED | mRNA-Seq | 10.1105/tpc.113.117325 |
| SRR1005288 | Pi_21d_R_Recov_6h_2 | grown without sodium dihydrogenphosphate for 21 days and then grown in grown in 0.323 millimolar sodium dihydrogenphosphate for 6 hours | root | Oryza Sativa Japonica | PAIRED | mRNA-Seq | 10.1105/tpc.113.117325 |
| SRR1005289 | Pi_21d_R_Recov_6h_3 | grown without sodium dihydrogenphosphate for 21 days and then grown in grown in 0.323 millimolar sodium dihydrogenphosphate for 6 hours | root | Oryza Sativa Japonica | PAIRED | mRNA-Seq | 10.1105/tpc.113.117325 |
| SRR1005353 | Pi_21d_S_control_1 | grown in 0.323 millimolar sodium dihydrogenphosphate for 21 days | Shoot | Oryza Sativa Japonica | PAIRED | mRNA-Seq | 10.1105/tpc.113.117325 |
| SRR1005354 | Pi_21d_S_control_2 | grown in 0.323 millimolar sodium dihydrogenphosphate for 21 days | Shoot | Oryza Sativa Japonica | PAIRED | mRNA-Seq | 10.1105/tpc.113.117325 |
| SRR1005355 | Pi_21d_S_control_3 | grown in 0.323 millimolar sodium dihydrogenphosphate for 21 days | Shoot | Oryza Sativa Japonica | PAIRED | mRNA-Seq | 10.1105/tpc.113.117325 |
| SRR1005356 | Pi_21d_S_Recov_0h_1 | grown without sodium dihydrogenphosphate for 21 days | Shoot | Oryza Sativa Japonica | PAIRED | mRNA-Seq | 10.1105/tpc.113.117325 |
| SRR1005357 | Pi_21d_S_Recov_0h_2 | grown without sodium dihydrogenphosphate for 21 days | Shoot | Oryza Sativa Japonica | PAIRED | mRNA-Seq | 10.1105/tpc.113.117325 |
| SRR1005358 | Pi_21d_S_Recov_0h_3 | grown without sodium dihydrogenphosphate for 21 days | Shoot | Oryza Sativa Japonica | PAIRED | mRNA-Seq | 10.1105/tpc.113.117325 |
| SRR1005332 | Pi_21d_S_Recov_1h_1 | grown without sodium dihydrogenphosphate for 21 days and then grown in grown in 0.323 millimolar sodium dihydrogenphosphate for 1 hour | Shoot | Oryza Sativa Japonica | PAIRED | mRNA-Seq | 10.1105/tpc.113.117325 |
| SRR1005333 | Pi_21d_S_Recov_1h_2 | grown without sodium dihydrogenphosphate for 21 days and then grown in grown in 0.323 millimolar sodium dihydrogenphosphate for 1 hour | Shoot | Oryza Sativa Japonica | PAIRED | mRNA-Seq | 10.1105/tpc.113.117325 |
| SRR1005334 | Pi_21d_S_Recov_1h_3 | grown without sodium dihydrogenphosphate for 21 days and then grown in grown in 0.323 millimolar sodium dihydrogenphosphate for 1 hour | Shoot | Oryza Sativa Japonica | PAIRED | mRNA-Seq | 10.1105/tpc.113.117325 |
| SRR1005341 | Pi_21d_S_Recov_24h_1 | grown without sodium dihydrogenphosphate for 21 days and then grown in grown in 0.323 millimolar sodium dihydrogenphosphate for 24 hours | Shoot | Oryza Sativa Japonica | PAIRED | mRNA-Seq | 10.1105/tpc.113.117325 |
| SRR1005342 | Pi_21d_S_Recov_24h_2 | grown without sodium dihydrogenphosphate for 21 days and then grown in grown in 0.323 millimolar sodium dihydrogenphosphate for 24 hours | Shoot | Oryza Sativa Japonica | PAIRED | mRNA-Seq | 10.1105/tpc.113.117325 |
| SRR1005343 | Pi_21d_S_Recov_24h_3 | grown without sodium dihydrogenphosphate for 21 days and then grown in grown in 0.323 millimolar sodium dihydrogenphosphate for 24 hours | Shoot | Oryza Sativa Japonica | PAIRED | mRNA-Seq | 10.1105/tpc.113.117325 |
| SRR1005350 | Pi_21d_S_Recov_6h_1 | grown without sodium dihydrogenphosphate for 21 days and then grown in grown in 0.323 millimolar sodium dihydrogenphosphate for 6 hours | Shoot | Oryza Sativa Japonica | PAIRED | mRNA-Seq | 10.1105/tpc.113.117325 |
| SRR1005351 | Pi_21d_S_Recov_6h_2 | grown without sodium dihydrogenphosphate for 21 days and then grown in grown in 0.323 millimolar sodium dihydrogenphosphate for 6 hours | Shoot | Oryza Sativa Japonica | PAIRED | mRNA-Seq | 10.1105/tpc.113.117325 |
| SRR1005352 | Pi_21d_S_Recov_6h_3 | grown without sodium dihydrogenphosphate for 21 days and then grown in grown in 0.323 millimolar sodium dihydrogenphosphate for 6 hours | Shoot | Oryza Sativa Japonica | PAIRED | mRNA-Seq | 10.1105/tpc.113.117325 |
