## Supplementary material for "RE3DB: A multi-omics phylogenomics platform for rice E3 ubiquitin ligases identifies novel regulators of pollen germination": Table S5

| **Run** | **Treatment** | **Tissue_original** | **Description** | **Cultivar** | **Layout** | **Strategy** | **Source** |
| --- | --- | --- | --- | --- | --- | --- | --- |
| DRR006631 | ABA_0_1 | root |  | Oryza Sativa Japonica | Single | mRNA-Seq | https://trace.ncbi.nlm.nih.gov/Traces/sra/?study=DRP000997 |
| DRR006713 | ABA_0_2 | root |  | Oryza Sativa Japonica | Single | mRNA-Seq | https://trace.ncbi.nlm.nih.gov/Traces/sra/?study=DRP000997 |
| DRR006632 | ABA_1h_1 | root |  | Oryza Sativa Japonica | Single | mRNA-Seq | https://trace.ncbi.nlm.nih.gov/Traces/sra/?study=DRP000997 |
| DRR006714 | ABA_1h_2 | root |  | Oryza Sativa Japonica | Single | mRNA-Seq | https://trace.ncbi.nlm.nih.gov/Traces/sra/?study=DRP000997 |
| DRR006633 | ABA_3h_1 | root |  | Oryza Sativa Japonica | Single | mRNA-Seq | https://trace.ncbi.nlm.nih.gov/Traces/sra/?study=DRP000997 |
| DRR006715 | ABA_3h_2 | root |  | Oryza Sativa Japonica | Single | mRNA-Seq | https://trace.ncbi.nlm.nih.gov/Traces/sra/?study=DRP000997 |
| DRR006634 | ABA_6h_1 | root |  | Oryza Sativa Japonica | Single | mRNA-Seq | https://trace.ncbi.nlm.nih.gov/Traces/sra/?study=DRP000997 |
| DRR006716 | ABA_6h_2 | root |  | Oryza Sativa Japonica | Single | mRNA-Seq | https://trace.ncbi.nlm.nih.gov/Traces/sra/?study=DRP000997 |
| DRR006635 | ABA_12h_1 | root |  | Oryza Sativa Japonica | Single | mRNA-Seq | https://trace.ncbi.nlm.nih.gov/Traces/sra/?study=DRP000997 |
| DRR006717 | ABA_12h_2 | root |  | Oryza Sativa Japonica | Single | mRNA-Seq | https://trace.ncbi.nlm.nih.gov/Traces/sra/?study=DRP000997 |
| DRR006636 | ABA_1day_1 | root |  | Oryza Sativa Japonica | Single | mRNA-Seq | https://trace.ncbi.nlm.nih.gov/Traces/sra/?study=DRP000997 |
| DRR006718 | ABA_1day_2 | root |  | Oryza Sativa Japonica | Single | mRNA-Seq | https://trace.ncbi.nlm.nih.gov/Traces/sra/?study=DRP000997 |
| SRR1264023 | Ethylene_mock_1 | seedling |  | Oryza Sativa Japonica | PAIRED | mRNA-Seq | 10.1371/journal.pgen.1004701 |
| SRR1264024 | Ethylene_mock_2 | seedling |  | Oryza Sativa Japonica | PAIRED | mRNA-Seq | 10.1371/journal.pgen.1004701 |
| SRR1985011 | Ethylene_mock_3 | seedling |  | Oryza Sativa Japonica | PAIRED | mRNA-Seq | 10.1371/journal.pgen.1004701 |
| SRR1985012 | Ethylene_mock_4 | seedling |  | Oryza Sativa Japonica | PAIRED | mRNA-Seq | 10.1371/journal.pgen.1004701 |
| SRR1987330 | Ethylene_mock_5 | seedling |  | Oryza Sativa Japonica | PAIRED | mRNA-Seq | 10.1371/journal.pgen.1004701 |
| SRR1987331 | Ethylene_mock_6 | seedling |  | Oryza Sativa Japonica | PAIRED | mRNA-Seq | 10.1371/journal.pgen.1004701 |
| SRR1264026 | Ethylene_1 | seedling |  | Oryza Sativa Japonica | PAIRED | mRNA-Seq | 10.1371/journal.pgen.1004701 |
| SRR1264028 | Ethylene_2 | seedling |  | Oryza Sativa Japonica | PAIRED | mRNA-Seq | 10.1371/journal.pgen.1004701 |
| SRR1985015 | Ethylene_3 | seedling |  | Oryza Sativa Japonica | PAIRED | mRNA-Seq | 10.1371/journal.pgen.1004701 |
| SRR1985017 | Ethylene_4 | seedling |  | Oryza Sativa Japonica | PAIRED | mRNA-Seq | 10.1371/journal.pgen.1004701 |
| SRR1987332 | Ethylene_5 | seedling |  | Oryza Sativa Japonica | PAIRED | mRNA-Seq | 10.1371/journal.pgen.1004701 |
| SRR1987333 | Ethylene_6 | seedling |  | Oryza Sativa Japonica | PAIRED | mRNA-Seq | 10.1371/journal.pgen.1004701 |
| DRR006579 | JA_0_1 | root |  | Oryza Sativa Japonica | Single | mRNA-Seq | https://trace.ncbi.nlm.nih.gov/Traces/sra/?study=DRP000997 |
| DRR006643 | JA_0_2 | root |  | Oryza Sativa Japonica | Single | mRNA-Seq | https://trace.ncbi.nlm.nih.gov/Traces/sra/?study=DRP000997 |
| DRR006580 | JA_1h_1 | root |  | Oryza Sativa Japonica | Single | mRNA-Seq | https://trace.ncbi.nlm.nih.gov/Traces/sra/?study=DRP000997 |
| DRR006644 | JA_1h_2 | root |  | Oryza Sativa Japonica | Single | mRNA-Seq | https://trace.ncbi.nlm.nih.gov/Traces/sra/?study=DRP000997 |
| DRR006581 | JA_3h_1 | root |  | Oryza Sativa Japonica | Single | mRNA-Seq | https://trace.ncbi.nlm.nih.gov/Traces/sra/?study=DRP000997 |
| DRR006645 | JA_3h_2 | root |  | Oryza Sativa Japonica | Single | mRNA-Seq | https://trace.ncbi.nlm.nih.gov/Traces/sra/?study=DRP000997 |
| DRR006582 | JA_6h_1 | root |  | Oryza Sativa Japonica | Single | mRNA-Seq | https://trace.ncbi.nlm.nih.gov/Traces/sra/?study=DRP000997 |
| DRR006646 | JA_6h_2 | root |  | Oryza Sativa Japonica | Single | mRNA-Seq | https://trace.ncbi.nlm.nih.gov/Traces/sra/?study=DRP000997 |
| DRR006583 | JA_12h_1 | root |  | Oryza Sativa Japonica | Single | mRNA-Seq | https://trace.ncbi.nlm.nih.gov/Traces/sra/?study=DRP000997 |
| DRR006647 | JA_12h_2 | root |  | Oryza Sativa Japonica | Single | mRNA-Seq | https://trace.ncbi.nlm.nih.gov/Traces/sra/?study=DRP000997 |
| DRR006584 | JA_1day_1 | root |  | Oryza Sativa Japonica | Single | mRNA-Seq | https://trace.ncbi.nlm.nih.gov/Traces/sra/?study=DRP000997 |
| DRR006648 | JA_1day_2 | root |  | Oryza Sativa Japonica | Single | mRNA-Seq | https://trace.ncbi.nlm.nih.gov/Traces/sra/?study=DRP000997 |
| DRR006743 | JA_1day_3 | root |  | Oryza Sativa Japonica | Single | mRNA-Seq | https://trace.ncbi.nlm.nih.gov/Traces/sra/?study=DRP000997 |
