## Supplementary material for "RE3DB: A multi-omics phylogenomics platform for rice E3 ubiquitin ligases identifies novel regulators of pollen germination": Table S6

| **Gene** | **Group** | **category** | **subcategory** |
| --- | --- | --- | --- |
| LOC_Os01g01420 | inflorescence & floral sporophyte, anther developmental series, mature anther, male gametophytic lineage | RING | HC |
| LOC_Os01g03100 | inflorescence & floral sporophyte, anther developmental series, mature anther, male gametophytic lineage, sperm | RING | V |
| LOC_Os01g03510 | inflorescence & floral sporophyte, anther developmental series, mature anther, male gametophytic lineage, pollen | DWD | D |
| LOC_Os01g04870 | inflorescence & floral sporophyte, anther developmental series, mature anther, male gametophytic lineage, sperm | DWD | B |
| LOC_Os01g05880 | female gametophytic lineage, post-fertilization | F-box | FBX |
| LOC_Os01g05890 | inflorescence & floral sporophyte, anther developmental series, mature anther, male gametophytic lineage | F-box | FBX |
| LOC_Os01g05970 | inflorescence & floral sporophyte, anther developmental series, mature anther, male gametophytic lineage, sperm | F-box | FBO |
| LOC_Os01g06360 | inflorescence & floral sporophyte, anther developmental series, mature anther | F-box |  |
| LOC_Os01g06590 | inflorescence & floral sporophyte, anther developmental series, mature anther | RING | H2 |
| LOC_Os01g07160 | inflorescence & floral sporophyte, anther developmental series, mature anther | F-box | FBX |
| LOC_Os01g08340 | inflorescence & floral sporophyte, anther developmental series, mature anther, male gametophytic lineage, pollen | RING | H2 |
| LOC_Os01g08830 | female gametophytic lineage, post-fertilization, seed development, embryo | F-box | FBDUF |
| LOC_Os01g09252 | inflorescence & floral sporophyte, anther developmental series, mature anther, male gametophytic lineage, sperm | DWD | D |
| LOC_Os01g11480 | seed development | RING | H2 |
| LOC_Os01g11490 | inflorescence & floral sporophyte, anther developmental series, mature anther | RING | H2 |
| LOC_Os01g11500 | vegetative body, root system | RING | H2 |
| LOC_Os01g12930 | inflorescence & floral sporophyte, anther developmental series, mature anther | U-box | VI |
| LOC_Os01g13370 | inflorescence & floral sporophyte, anther developmental series, mature anther, male gametophytic lineage, pollen | RING | HC |
| LOC_Os01g16950 | inflorescence & floral sporophyte, anther developmental series, mature anther, male gametophytic lineage, pollen | RING | H2 |
| LOC_Os01g17390 | inflorescence & floral sporophyte, anther developmental series, mature anther, male gametophytic lineage, pollen | F-box | FBX |
| LOC_Os01g24880 | seed development, seed | RING | H2 |
| LOC_Os01g27150 | inflorescence & floral sporophyte, anther developmental series, mature anther, male gametophytic lineage, pollen | Cullin | Cullin 1a |
| LOC_Os01g27680 | inflorescence & floral sporophyte, anther developmental series, mature anther | F-box |  |
| LOC_Os01g28150 | inflorescence & floral sporophyte, anther developmental series, mature anther, male gametophytic lineage, sperm | F-box | FBX |
| LOC_Os01g28520 | inflorescence & floral sporophyte, anther developmental series, mature anther | F-box | FBX |
| LOC_Os01g33490 | inflorescence & floral sporophyte, anther developmental series, mature anther, male gametophytic lineage, pollen | F-box | FBX |
| LOC_Os01g34220 | seed development, seed | F-box | FBX |
| LOC_Os01g35100 | inflorescence & floral sporophyte, anther developmental series, mature anther | RING | H2 |
| LOC_Os01g35120 | inflorescence & floral sporophyte, anther developmental series, mature anther | RING | H2 |
| LOC_Os01g36940 | inflorescence & floral sporophyte, anther developmental series, mature anther, male gametophytic lineage, sperm | F-box | FBX |
| LOC_Os01g38690 | inflorescence & floral sporophyte, anther developmental series, mature anther | RING | HC |
| LOC_Os01g38700 | inflorescence & floral sporophyte, anther developmental series, mature anther | RING | HC |
| LOC_Os01g38710 | inflorescence & floral sporophyte, anther developmental series, mature anther | RING | HC |
| LOC_Os01g38720 | inflorescence & floral sporophyte, anther developmental series, mature anther | RING | HC |
| LOC_Os01g38760 | inflorescence & floral sporophyte, anther developmental series, mature anther | RING | HC |
| LOC_Os01g39380 | inflorescence & floral sporophyte, anther developmental series, mature anther | DWD | I |
| LOC_Os01g40160 | inflorescence & floral sporophyte, anther developmental series, mature anther | F-box | FBX |
| LOC_Os01g42500 | inflorescence & floral sporophyte, anther developmental series, mature anther, male gametophytic lineage, sperm | RING | V |
| LOC_Os01g44240 | inflorescence & floral sporophyte | RING | H2 |
| LOC_Os01g44394 | female gametophytic lineage, egg cell | DWD | D |
| LOC_Os01g45020 | inflorescence & floral sporophyte, anther developmental series, mature anther, male gametophytic lineage, sperm | F-box |  |
| LOC_Os01g46500 | inflorescence & floral sporophyte, anther developmental series, mature anther, male gametophytic lineage, sperm | F-box | FBX |
| LOC_Os01g47740 | inflorescence & floral sporophyte, anther developmental series, mature anther | RING | H2 |
| LOC_Os01g48310 | inflorescence & floral sporophyte, anther developmental series, mature anther | RING | Not Determined |
| LOC_Os01g49280 | vegetative body, shoot system, shoot and leaf tissues | RING | Not Determined |
| LOC_Os01g49470 | vegetative body, shoot system, shoot and leaf tissues | RING | H2 |
| LOC_Os01g49770 | inflorescence & floral sporophyte, anther developmental series, mature anther, male gametophytic lineage, sperm | RING | H2 |
| LOC_Os01g50750 | inflorescence & floral sporophyte, anther developmental series, mature anther | RING | H2 |
| LOC_Os01g52110 | vegetative body, callus | RING | H2 |
| LOC_Os01g52640 | vegetative body, shoot system, shoot and leaf tissues | DWD | L |
| LOC_Os01g52980 | inflorescence & floral sporophyte, anther developmental series, mature anther, male gametophytic lineage, pollen | F-box | FBX |
| LOC_Os01g53130 | inflorescence & floral sporophyte, anther developmental series, mature anther, male gametophytic lineage, pollen | RING | V |
| LOC_Os01g55210 | inflorescence & floral sporophyte, anther developmental series, early anther | F-box | FBX |
| LOC_Os01g55430 | inflorescence & floral sporophyte, anther developmental series, mature anther, male gametophytic lineage, sperm | F-box | FBT |
| LOC_Os01g56740 | female gametophytic lineage, post-fertilization | F-box | FBDUF |
| LOC_Os01g56750 | inflorescence & floral sporophyte, anther developmental series, mature anther | F-box | FBDUF |
| LOC_Os01g57110 | inflorescence & floral sporophyte, anther developmental series, mature anther, male gametophytic lineage, sperm | RING | Not Determined |
| LOC_Os01g57230 | inflorescence & floral sporophyte | BTB |  |
| LOC_Os01g58780 | inflorescence & floral sporophyte, anther developmental series, mature anther, male gametophytic lineage | RING | H2 |
| LOC_Os01g59910 | female gametophytic lineage, post-fertilization, seed development, embryo | F-box | FBX |
| LOC_Os01g60920 | inflorescence & floral sporophyte, anther developmental series, mature anther, male gametophytic lineage, sperm | F-box | FBX |
| LOC_Os01g61420 | inflorescence & floral sporophyte, anther developmental series, mature anther, male gametophytic lineage, pollen | RING | HC |
| LOC_Os01g61470 | vegetative body, shoot system, shoot and leaf tissues | RING | H2 |
| LOC_Os01g62640 | vegetative body, shoot system, node | RING | H2 |
| LOC_Os01g64030 | inflorescence & floral sporophyte, anther developmental series, mature anther, male gametophytic lineage, sperm | F-box | FBL |
| LOC_Os01g64570 | inflorescence & floral sporophyte, anther developmental series, mature anther | U-box | III |
| LOC_Os01g64620 | vegetative body, root system | RING | H2 |
| LOC_Os01g64700 | vegetative body, root system | F-box | FBT |
| LOC_Os01g66130 | inflorescence & floral sporophyte, anther developmental series, mature anther, male gametophytic lineage, sperm | U-box | II |
| LOC_Os01g68060 | inflorescence & floral sporophyte, anther developmental series, mature anther, male gametophytic lineage, sperm | RING | HC |
| LOC_Os01g68900 | inflorescence & floral sporophyte, anther developmental series, mature anther, male gametophytic lineage, pollen | RING | HC |
| LOC_Os01g69040 | inflorescence & floral sporophyte, anther developmental series, mature anther | RING | HC |
| LOC_Os01g70670 | seed development, seed | BTB |  |
| LOC_Os01g70920 | female gametophytic lineage | Cullin | Cullin 1d |
| LOC_Os01g71430 | inflorescence & floral sporophyte, anther developmental series, early anther | F-box | FBDUF |
| LOC_Os01g71440 | inflorescence & floral sporophyte, anther developmental series, mature anther | F-box | FBDUF |
| LOC_Os01g71450 | inflorescence & floral sporophyte, anther developmental series, mature anther, male gametophytic lineage, pollen | F-box | FBX |
| LOC_Os01g71460 | inflorescence & floral sporophyte, anther developmental series, early anther | F-box | FBDUF |
| LOC_Os01g72310 | inflorescence & floral sporophyte, anther developmental series, mature anther, male gametophytic lineage, pollen | F-box | FBO |
| LOC_Os01g72480 | inflorescence & floral sporophyte, anther developmental series, mature anther, male gametophytic lineage, sperm | RING | HC |
| LOC_Os10g03600 | inflorescence & floral sporophyte, anther developmental series, mature anther, male gametophytic lineage, sperm | F-box | FBX |
| LOC_Os10g03620 | inflorescence & floral sporophyte, anther developmental series, mature anther, male gametophytic lineage, sperm | F-box | FBX |
| LOC_Os10g03730 | female gametophytic lineage | F-box | FBX |
| LOC_Os10g03740 | female gametophytic lineage | F-box | FBX |
| LOC_Os10g03750 | female gametophytic lineage | F-box | FBX |
| LOC_Os10g03870 | seed development, seed | F-box | FBX |
| LOC_Os10g04370 | inflorescence & floral sporophyte, anther developmental series, mature anther | F-box | FBX |
| LOC_Os10g04590 | inflorescence & floral sporophyte, anther developmental series, mature anther, male gametophytic lineage, sperm | F-box | FBX |
| LOC_Os10g04600 | inflorescence & floral sporophyte, anther developmental series, mature anther, male gametophytic lineage, sperm | F-box | FBX |
| LOC_Os10g04700 | inflorescence & floral sporophyte, anther developmental series, mature anther, male gametophytic lineage, pollen | F-box | FBX |
| LOC_Os10g04750 | inflorescence & floral sporophyte, anther developmental series, mature anther, male gametophytic lineage, sperm | F-box | FBX |
| LOC_Os10g04850 | female gametophytic lineage, post-fertilization | F-box | FBO |
| LOC_Os10g04900 | seed development | F-box | FBX |
| LOC_Os10g04980 | seed development | F-box | FBX |
| LOC_Os10g05000 | inflorescence & floral sporophyte, anther developmental series, mature anther, male gametophytic lineage | F-box | FBX |
| LOC_Os10g05200 | inflorescence & floral sporophyte, anther developmental series, mature anther, male gametophytic lineage, sperm | F-box | FBX |
| LOC_Os10g05240 | inflorescence & floral sporophyte, anther developmental series, mature anther | F-box | FBX |
| LOC_Os10g05500 | seed development, seed | F-box | FBX |
| LOC_Os10g05520 | inflorescence & floral sporophyte, anther developmental series, mature anther, male gametophytic lineage, sperm | F-box | FBX |
| LOC_Os10g05800 | inflorescence & floral sporophyte, anther developmental series, mature anther, male gametophytic lineage, sperm | F-box |  |
| LOC_Os10g06700 | inflorescence & floral sporophyte, anther developmental series, mature anther | F-box | FBX |
| LOC_Os10g10390 | inflorescence & floral sporophyte, anther developmental series, mature anther, male gametophytic lineage, sperm | F-box | FBX |
| LOC_Os10g10410 | inflorescence & floral sporophyte, anther developmental series, mature anther, male gametophytic lineage, sperm | F-box | FBX |
| LOC_Os10g10420 | inflorescence & floral sporophyte, anther developmental series, mature anther | F-box | FBX |
| LOC_Os10g17930 | inflorescence & floral sporophyte, anther developmental series, mature anther, male gametophytic lineage, pollen | F-box | FBX |
| LOC_Os10g17940 | inflorescence & floral sporophyte, anther developmental series, mature anther | F-box | FBX |
| LOC_Os10g20600 | inflorescence & floral sporophyte, anther developmental series, mature anther, male gametophytic lineage, sperm | RING | H2 |
| LOC_Os10g24010 | inflorescence & floral sporophyte, anther developmental series, mature anther | F-box | FBX |
| LOC_Os10g24900 | inflorescence & floral sporophyte, anther developmental series, mature anther, male gametophytic lineage, sperm | F-box | FBK |
| LOC_Os10g25210 | seed development, endosperm | F-box | FBX |
| LOC_Os10g25660 | inflorescence & floral sporophyte, anther developmental series, mature anther, male gametophytic lineage, pollen | F-box | FBX |
| LOC_Os10g25680 | inflorescence & floral sporophyte, anther developmental series, mature anther, male gametophytic lineage, sperm | F-box | FBL |
| LOC_Os10g26610 | inflorescence & floral sporophyte, anther developmental series, mature anther, male gametophytic lineage, pollen | RING | H2 |
| LOC_Os10g28760 | inflorescence & floral sporophyte, anther developmental series, mature anther | BTB |  |
| LOC_Os10g28770 | inflorescence & floral sporophyte, anther developmental series, mature anther | BTB |  |
| LOC_Os10g28790 | inflorescence & floral sporophyte, anther developmental series, mature anther | BTB |  |
| LOC_Os10g28810 | inflorescence & floral sporophyte, anther developmental series, mature anther | BTB |  |
| LOC_Os10g28860 | inflorescence & floral sporophyte, anther developmental series, mature anther | BTB |  |
| LOC_Os10g28870 | inflorescence & floral sporophyte, anther developmental series, mature anther | BTB |  |
| LOC_Os10g29060 | female gametophytic lineage | BTB |  |
| LOC_Os10g29100 | inflorescence & floral sporophyte, anther developmental series, mature anther | BTB |  |
| LOC_Os10g29110 | inflorescence & floral sporophyte, anther developmental series, mature anther | BTB |  |
| LOC_Os10g29150 | inflorescence & floral sporophyte, anther developmental series, mature anther | BTB |  |
| LOC_Os10g29180 | female gametophytic lineage | BTB |  |
| LOC_Os10g29230 | inflorescence & floral sporophyte, anther developmental series, mature anther, male gametophytic lineage, pollen | BTB |  |
| LOC_Os10g29380 | female gametophytic lineage | BTB |  |
| LOC_Os10g29410 | inflorescence & floral sporophyte, anther developmental series | BTB |  |
| LOC_Os10g29750 | inflorescence & floral sporophyte, anther developmental series, mature anther | BTB |  |
| LOC_Os10g29790 | inflorescence & floral sporophyte, anther developmental series, mature anther | BTB |  |
| LOC_Os10g29810 | inflorescence & floral sporophyte, anther developmental series, mature anther | BTB |  |
| LOC_Os10g29840 | inflorescence & floral sporophyte, anther developmental series, mature anther | BTB |  |
| LOC_Os10g30190 | inflorescence & floral sporophyte, anther developmental series, mature anther, male gametophytic lineage, pollen | BTB |  |
| LOC_Os10g30200 | inflorescence & floral sporophyte, anther developmental series, mature anther | SKP1 |  |
| LOC_Os10g30310 | inflorescence & floral sporophyte, panicle | RING | H2 |
| LOC_Os10g31850 | inflorescence & floral sporophyte, anther developmental series, mature anther, male gametophytic lineage, pollen | RING | H2 |
| LOC_Os10g32710 | seed development | DWD | B |
| LOC_Os10g32740 | inflorescence & floral sporophyte, anther developmental series, mature anther | RING | H2 |
| LOC_Os10g32750 | inflorescence & floral sporophyte, anther developmental series, mature anther | RING | H2 |
| LOC_Os10g32980 | inflorescence & floral sporophyte, panicle | RING | Not Determined |
| LOC_Os10g34030 | inflorescence & floral sporophyte, anther developmental series, mature anther, male gametophytic lineage, pollen | RING | Not Determined |
| LOC_Os10g34340 | inflorescence & floral sporophyte, anther developmental series, mature anther | F-box | FBX |
| LOC_Os10g35190 | inflorescence & floral sporophyte, anther developmental series, mature anther, male gametophytic lineage, pollen | RING | HC |
| LOC_Os10g35920 | inflorescence & floral sporophyte, anther developmental series, mature anther, male gametophytic lineage, sperm | F-box | FBX |
| LOC_Os10g36310 | vegetative body, root system | F-box | FBX |
| LOC_Os10g37540 | female gametophytic lineage, post-fertilization | F-box | FBDUF |
| LOC_Os10g37570 | inflorescence & floral sporophyte, anther developmental series, mature anther | F-box | FBDUF |
| LOC_Os10g40120 | vegetative body, root system | U-box | IV |
| LOC_Os10g40490 | vegetative body, root system | U-box | VII |
| LOC_Os10g41220 | inflorescence & floral sporophyte, anther developmental series, mature anther, male gametophytic lineage, sperm | U-box | IV |
| LOC_Os10g41580 | inflorescence & floral sporophyte | RING | HC |
| LOC_Os10g41650 | inflorescence & floral sporophyte, anther developmental series, mature anther, male gametophytic lineage, sperm | F-box | FBX |
| LOC_Os10g41829 | inflorescence & floral sporophyte, anther developmental series, mature anther | F-box | FBX |
| LOC_Os10g42390 | inflorescence & floral sporophyte, anther developmental series, mature anther | RING | H2 |
| LOC_Os11g01240 | vegetative body, shoot system, shoot and leaf tissues | RING | Not Determined |
| LOC_Os11g02250 | female gametophytic lineage | RING | H2 |
| LOC_Os11g02260 | inflorescence & floral sporophyte, anther developmental series, mature anther | RING | H2 |
| LOC_Os11g03680 | inflorescence & floral sporophyte, anther developmental series, mature anther, male gametophytic lineage, pollen | F-box | FBX |
| LOC_Os11g04280 | inflorescence & floral sporophyte, anther developmental series, mature anther | RING | H2 |
| LOC_Os11g04281 | inflorescence & floral sporophyte, anther developmental series, mature anther | RING | H2 |
| LOC_Os11g04680 | inflorescence & floral sporophyte, anther developmental series, mature anther | RING | H2 |
| LOC_Os11g04690 | inflorescence & floral sporophyte, anther developmental series, mature anther | RING | Not Determined |
| LOC_Os11g05200 | inflorescence & floral sporophyte, anther developmental series, mature anther | RING | H2 |
| LOC_Os11g05300 | inflorescence & floral sporophyte, anther developmental series, mature anther, male gametophytic lineage, sperm | RING | H2 |
| LOC_Os11g05590 | inflorescence & floral sporophyte, anther developmental series, mature anther | F-box |  |
| LOC_Os11g06420 | inflorescence & floral sporophyte, anther developmental series, mature anther, male gametophytic lineage, sperm | F-box | FBT |
| LOC_Os11g07970 | inflorescence & floral sporophyte, anther developmental series, mature anther, male gametophytic lineage, sperm | F-box | FBW |
| LOC_Os11g09360 | vegetative body, shoot system, node | F-box | FBX |
| LOC_Os11g09580 | female gametophytic lineage | F-box | FBL |
| LOC_Os11g09600 | female gametophytic lineage, egg cell | F-box | FBX |
| LOC_Os11g09630 | female gametophytic lineage | F-box | FBX |
| LOC_Os11g09640 | inflorescence & floral sporophyte, anther developmental series, mature anther, male gametophytic lineage, sperm | F-box | FBX |
| LOC_Os11g09670 | female gametophytic lineage | F-box | FBX |
| LOC_Os11g10200 | female gametophytic lineage | F-box | FBX |
| LOC_Os11g10300 | vegetative body, shoot system, shoot and leaf tissues | F-box | FBX |
| LOC_Os11g10340 | inflorescence & floral sporophyte, anther developmental series, mature anther, male gametophytic lineage, sperm | F-box | FBX |
| LOC_Os11g12480 | female gametophytic lineage | F-box | FBL |
| LOC_Os11g16280 | inflorescence & floral sporophyte, anther developmental series, mature anther, male gametophytic lineage, sperm | F-box |  |
| LOC_Os11g16330 | female gametophytic lineage, egg cell | F-box |  |
| LOC_Os11g19500 | inflorescence & floral sporophyte, anther developmental series, mature anther | RING | Not Determined |
| LOC_Os11g32940 | inflorescence & floral sporophyte, anther developmental series, mature anther, male gametophytic lineage, sperm | F-box | FBX |
| LOC_Os11g33180 | inflorescence & floral sporophyte, anther developmental series, mature anther | F-box | FBL |
| LOC_Os11g33210 | inflorescence & floral sporophyte | F-box | FBL |
| LOC_Os11g33220 | inflorescence & floral sporophyte, anther developmental series, mature anther | F-box | FBX |
| LOC_Os11g35870 | seed development | RING | Not Determined |
| LOC_Os11g36350 | inflorescence & floral sporophyte, anther developmental series, mature anther | F-box | FBDUF |
| LOC_Os11g36430 | seed development, endosperm | RING | H2 |
| LOC_Os11g36560 | inflorescence & floral sporophyte, anther developmental series, mature anther | RING | HC |
| LOC_Os11g36610 | inflorescence & floral sporophyte, gynoecium organs | F-box | FBDUF |
| LOC_Os11g36970 | inflorescence & floral sporophyte, anther developmental series, mature anther, male gametophytic lineage, sperm | RING | Not Determined |
| LOC_Os11g37300 | inflorescence & floral sporophyte, anther developmental series, mature anther, male gametophytic lineage, pollen | F-box | FBDUF |
| LOC_Os11g37340 | female gametophytic lineage, post-fertilization | F-box | FBX |
| LOC_Os11g38100 | inflorescence & floral sporophyte, anther developmental series, mature anther | F-box | FBDUF |
| LOC_Os11g38130 | inflorescence & floral sporophyte, anther developmental series, mature anther, male gametophytic lineage, pollen | F-box | FBDUF |
| LOC_Os11g38180 | inflorescence & floral sporophyte, anther developmental series, mature anther | F-box | FBDUF |
| LOC_Os11g38800 | vegetative body, root system | RING | V |
| LOC_Os11g39609 | inflorescence & floral sporophyte, anther developmental series, mature anther | F-box | FBDUF |
| LOC_Os11g40220 | inflorescence & floral sporophyte, anther developmental series, mature anther | BTB |  |
| LOC_Os11g40680 | inflorescence & floral sporophyte, anther developmental series | BTB |  |
| LOC_Os11g41290 | inflorescence & floral sporophyte, anther developmental series, mature anther | BTB |  |
| LOC_Os11g41300 | inflorescence & floral sporophyte, anther developmental series, mature anther | BTB |  |
| LOC_Os11g41310 | inflorescence & floral sporophyte, anther developmental series, mature anther | BTB |  |
| LOC_Os11g41350 | inflorescence & floral sporophyte, anther developmental series, mature anther | BTB |  |
| LOC_Os11g41560 | inflorescence & floral sporophyte, anther developmental series, mature anther, male gametophytic lineage, pollen | F-box | FBX |
| LOC_Os11g41570 | inflorescence & floral sporophyte, anther developmental series, mature anther, male gametophytic lineage | F-box | FBX |
| LOC_Os11g42160 | inflorescence & floral sporophyte, anther developmental series, mature anther, male gametophytic lineage, pollen | F-box |  |
| LOC_Os11g42240 | inflorescence & floral sporophyte, anther developmental series, mature anther | F-box | FBX |
| LOC_Os11g42270 | inflorescence & floral sporophyte, anther developmental series, mature anther | F-box | FBX |
| LOC_Os11g42280 | inflorescence & floral sporophyte, anther developmental series, mature anther | F-box | FBX |
| LOC_Os11g42310 | inflorescence & floral sporophyte, anther developmental series, mature anther | F-box | FBL |
| LOC_Os11g45560 | inflorescence & floral sporophyte, anther developmental series, mature anther | BTB |  |
| LOC_Os12g01190 | inflorescence & floral sporophyte, anther developmental series, mature anther, male gametophytic lineage, sperm | RING | C2 |
| LOC_Os12g01230 | seed development, endosperm | RING | Not Determined |
| LOC_Os12g01340 | inflorescence & floral sporophyte, panicle | RING | Not Determined |
| LOC_Os12g02030 | vegetative body, root system | BTB |  |
| LOC_Os12g02210 | vegetative body, callus | RING | H2 |
| LOC_Os12g02220 | inflorescence & floral sporophyte, anther developmental series, mature anther | RING | H2 |
| LOC_Os12g03440 | female gametophytic lineage, post-fertilization | F-box | FBX |
| LOC_Os12g04130 | inflorescence & floral sporophyte, gynoecium organs | F-box | FBO |
| LOC_Os12g04660 | inflorescence & floral sporophyte, anther developmental series, mature anther, male gametophytic lineage | RING | H2 |
| LOC_Os12g05280 | inflorescence & floral sporophyte, anther developmental series, mature anther | RING | Not Determined |
| LOC_Os12g05370 | inflorescence & floral sporophyte, anther developmental series, mature anther, male gametophytic lineage, sperm | RING | H2 |
| LOC_Os12g05609 | inflorescence & floral sporophyte, anther developmental series, mature anther, male gametophytic lineage, pollen | F-box | FBDUF |
| LOC_Os12g05709 | inflorescence & floral sporophyte, anther developmental series, mature anther | F-box | FBO |
| LOC_Os12g06030 | vegetative body, root system | F-box | FBX |
| LOC_Os12g06410 | vegetative body, root system | U-box | III |
| LOC_Os12g06810 | inflorescence & floral sporophyte, anther developmental series, mature anther | DWD | C |
| LOC_Os12g10250 | inflorescence & floral sporophyte | RING | H2 |
| LOC_Os12g13360 | vegetative body, shoot system, shoot and leaf tissues | Cullin | Cullin 1e |
| LOC_Os12g16690 | inflorescence & floral sporophyte, anther developmental series, mature anther, male gametophytic lineage, pollen | RING | HC |
| LOC_Os12g24080 | inflorescence & floral sporophyte, anther developmental series, mature anther, male gametophytic lineage, sperm | HECT |  |
| LOC_Os12g24530 | inflorescence & floral sporophyte, anther developmental series, mature anther, male gametophytic lineage, pollen | RING | H2 |
| LOC_Os12g27760 | inflorescence & floral sporophyte, anther developmental series, mature anther | F-box | FBX |
| LOC_Os12g27790 | inflorescence & floral sporophyte, anther developmental series, mature anther, male gametophytic lineage | F-box | FBX |
| LOC_Os12g27810 | inflorescence & floral sporophyte, anther developmental series, mature anther | F-box | FBX |
| LOC_Os12g30940 | female gametophytic lineage, post-fertilization | F-box | FBX |
| LOC_Os12g30950 | inflorescence & floral sporophyte, anther developmental series, mature anther | F-box |  |
| LOC_Os12g30990 | inflorescence & floral sporophyte, anther developmental series, mature anther | F-box | FBX |
| LOC_Os12g31340 | inflorescence & floral sporophyte, anther developmental series, mature anther, male gametophytic lineage, pollen | F-box | FBD |
| LOC_Os12g33830 | vegetative body | F-box | FBX |
| LOC_Os12g34200 | inflorescence & floral sporophyte, anther developmental series, mature anther | F-box | FBX |
| LOC_Os12g34210 | inflorescence & floral sporophyte, anther developmental series, mature anther, male gametophytic lineage | F-box |  |
| LOC_Os12g34240 | inflorescence & floral sporophyte, anther developmental series, mature anther | F-box | FBX |
| LOC_Os12g34290 | inflorescence & floral sporophyte, anther developmental series, mature anther | F-box | FBX |
| LOC_Os12g35320 | inflorescence & floral sporophyte, anther developmental series, mature anther, male gametophytic lineage, pollen | RING | H2 |
| LOC_Os12g37130 | inflorescence & floral sporophyte, anther developmental series, early anther | F-box | FBD |
| LOC_Os12g38090 | inflorescence & floral sporophyte, anther developmental series, mature anther | F-box | FBX |
| LOC_Os12g39520 | inflorescence & floral sporophyte, anther developmental series, mature anther, male gametophytic lineage, pollen | F-box | FBDUF |
| LOC_Os12g40140 | inflorescence & floral sporophyte, anther developmental series, mature anther, male gametophytic lineage, pollen | F-box | FBX |
| LOC_Os12g40310 | inflorescence & floral sporophyte, anther developmental series | F-box | FBX |
| LOC_Os12g40320 | inflorescence & floral sporophyte, anther developmental series | F-box | FBX |
| LOC_Os12g40360 | inflorescence & floral sporophyte, anther developmental series | F-box | FBX |
| LOC_Os12g40370 | inflorescence & floral sporophyte, anther developmental series, early anther | F-box | FBX |
| LOC_Os12g41300 | inflorescence & floral sporophyte, anther developmental series, mature anther | F-box | FBX |
| LOC_Os12g41630 | inflorescence & floral sporophyte, anther developmental series, mature anther | F-box | FBX |
| LOC_Os12g42340 | inflorescence & floral sporophyte, anther developmental series, mature anther | F-box | FBX |
| LOC_Os12g42540 | vegetative body, shoot system, shoot and leaf tissues | RING | H2 |
| LOC_Os12g43770 | female gametophytic lineage, post-fertilization, seed development, embryo | F-box | FBX |
| LOC_Os12g43930 | inflorescence & floral sporophyte, anther developmental series, mature anther, male gametophytic lineage, pollen | RING | HC |
| LOC_Os02g01160 | vegetative body, shoot system, shoot and leaf tissues | SKP1 |  |
| LOC_Os02g01170 | inflorescence & floral sporophyte, anther developmental series, mature anther, male gametophytic lineage, pollen | HECT |  |
| LOC_Os02g02350 | inflorescence & floral sporophyte, anther developmental series, mature anther, male gametophytic lineage, sperm | F-box | FBK |
| LOC_Os02g02380 | vegetative body | DWD | D |
| LOC_Os02g03620 | inflorescence & floral sporophyte, anther developmental series, mature anther, male gametophytic lineage, sperm | RING | HC |
| LOC_Os02g03660 | inflorescence & floral sporophyte | F-box | FBDUF |
| LOC_Os02g03760 | inflorescence & floral sporophyte, anther developmental series, mature anther, male gametophytic lineage, sperm | RING | HC |
| LOC_Os02g03810 | inflorescence & floral sporophyte | F-box | FBX |
| LOC_Os02g03910 | inflorescence & floral sporophyte, anther developmental series, mature anther | F-box | FBX |
| LOC_Os02g03980 | inflorescence & floral sporophyte, anther developmental series, mature anther | F-box | FBX |
| LOC_Os02g04000 | inflorescence & floral sporophyte, anther developmental series, mature anther | F-box | FBX |
| LOC_Os02g05700 | inflorescence & floral sporophyte, anther developmental series, mature anther, male gametophytic lineage, pollen | F-box | FBO |
| LOC_Os02g06470 | seed development, endosperm | F-box | FBX |
| LOC_Os02g06520 | inflorescence & floral sporophyte, anther developmental series, mature anther, male gametophytic lineage, sperm | F-box | FBLD |
| LOC_Os02g08200 | inflorescence & floral sporophyte, anther developmental series, mature anther, male gametophytic lineage | RING | H2 |
| LOC_Os02g09580 | female gametophytic lineage, post-fertilization | F-box | FBX |
| LOC_Os02g09820 | inflorescence & floral sporophyte, anther developmental series, mature anther, male gametophytic lineage, pollen | RING | H2 |
| LOC_Os02g11790 | seed development, seed | F-box | FBK |
| LOC_Os02g13180 | inflorescence & floral sporophyte, anther developmental series, mature anther | SKP1 |  |
| LOC_Os02g13810 | inflorescence & floral sporophyte, anther developmental series, mature anther, male gametophytic lineage, sperm | RING | Not Determined |
| LOC_Os02g14720 | seed development, seed | RING | Not Determined |
| LOC_Os02g15010 | vegetative body | RING | H2 |
| LOC_Os02g15080 | seed development | RING | H2 |
| LOC_Os02g15100 | vegetative body, root system | RING | H2 |
| LOC_Os02g15160 | inflorescence & floral sporophyte, anther developmental series, mature anther, male gametophytic lineage, sperm | F-box | FBX |
| LOC_Os02g16000 | inflorescence & floral sporophyte, anther developmental series, mature anther, male gametophytic lineage, pollen | BTB |  |
| LOC_Os02g16760 | inflorescence & floral sporophyte, anther developmental series, mature anther | F-box | FBX |
| LOC_Os02g17180 | inflorescence & floral sporophyte, anther developmental series, mature anther | F-box | FBX |
| LOC_Os02g17210 | inflorescence & floral sporophyte, anther developmental series, mature anther, male gametophytic lineage, pollen | F-box | FBX |
| LOC_Os02g18820 | inflorescence & floral sporophyte, anther developmental series, mature anther, male gametophytic lineage | DWD | C |
| LOC_Os02g19140 | vegetative body, root system | RING | HC |
| LOC_Os02g19540 | inflorescence & floral sporophyte, anther developmental series, mature anther, male gametophytic lineage, pollen | F-box | FBX |
| LOC_Os02g19804 | seed development | RING | HC |
| LOC_Os02g20690 | female gametophytic lineage | BTB |  |
| LOC_Os02g20720 | vegetative body, root system | BTB |  |
| LOC_Os02g21110 | inflorescence & floral sporophyte, anther developmental series, mature anther, male gametophytic lineage, sperm | F-box | FBK |
| LOC_Os02g21230 | inflorescence & floral sporophyte, anther developmental series, mature anther | F-box |  |
| LOC_Os02g21240 | inflorescence & floral sporophyte, anther developmental series, mature anther, male gametophytic lineage, sperm | F-box | FBX |
| LOC_Os02g28600 | inflorescence & floral sporophyte, anther developmental series, mature anther, male gametophytic lineage, pollen | F-box | FBX |
| LOC_Os02g28720 | inflorescence & floral sporophyte, anther developmental series, mature anther | U-box | II |
| LOC_Os02g28870 | inflorescence & floral sporophyte, anther developmental series, mature anther | U-box | V |
| LOC_Os02g31150 | inflorescence & floral sporophyte, anther developmental series, mature anther | RING | HC |
| LOC_Os02g32570 | inflorescence & floral sporophyte, gynoecium organs | RING | HC |
| LOC_Os02g33310 | inflorescence & floral sporophyte, anther developmental series, mature anther, male gametophytic lineage, sperm | F-box | FBD |
| LOC_Os02g33400 | inflorescence & floral sporophyte, anther developmental series, mature anther, male gametophytic lineage, sperm | F-box | FBL |
| LOC_Os02g33590 | inflorescence & floral sporophyte, anther developmental series, mature anther | U-box | III |
| LOC_Os02g33680 | inflorescence & floral sporophyte, anther developmental series, mature anther | U-box | III |
| LOC_Os02g33720 | vegetative body, shoot system, node | RING | H2 |
| LOC_Os02g33840 | inflorescence & floral sporophyte, anther developmental series, mature anther | F-box | FBX |
| LOC_Os02g35329 | inflorescence & floral sporophyte | RING | H2 |
| LOC_Os02g35347 | inflorescence & floral sporophyte | RING | H2 |
| LOC_Os02g35440 | vegetative body, shoot system, shoot and leaf tissues | RING | H2 |
| LOC_Os02g35530 | inflorescence & floral sporophyte, anther developmental series, mature anther | F-box | FBK |
| LOC_Os02g36300 | inflorescence & floral sporophyte, anther developmental series, mature anther, male gametophytic lineage, pollen | RING | H2 |
| LOC_Os02g36320 | female gametophytic lineage, post-fertilization | RING | H2 |
| LOC_Os02g36520 | inflorescence & floral sporophyte, anther developmental series, mature anther, male gametophytic lineage, sperm | F-box | FBK |
| LOC_Os02g36740 | inflorescence & floral sporophyte, anther developmental series, mature anther, male gametophytic lineage, pollen | RING | V |
| LOC_Os02g37856 | female gametophytic lineage, post-fertilization, seed development, embryo | DWD | B |
| LOC_Os02g38029 | female gametophytic lineage, egg cell | APC |  |
| LOC_Os02g38320 | inflorescence & floral sporophyte, anther developmental series, mature anther | BTB |  |
| LOC_Os02g38499 | inflorescence & floral sporophyte, anther developmental series, mature anther | F-box | FBX |
| LOC_Os02g38589 | inflorescence & floral sporophyte, anther developmental series, mature anther | F-box | FBX |
| LOC_Os02g38720 | inflorescence & floral sporophyte, anther developmental series, mature anther | F-box | FBX |
| LOC_Os02g39910 | inflorescence & floral sporophyte, anther developmental series, mature anther, male gametophytic lineage, sperm | BTB |  |
| LOC_Os02g40810 | inflorescence & floral sporophyte, anther developmental series, mature anther, male gametophytic lineage, pollen | RING | HC |
| LOC_Os02g41930 | inflorescence & floral sporophyte, anther developmental series, mature anther | F-box | FBX |
| LOC_Os02g43920 | inflorescence & floral sporophyte, anther developmental series, early anther | APC |  |
| LOC_Os02g44599 | inflorescence & floral sporophyte, anther developmental series, mature anther, male gametophytic lineage | U-box | IV |
| LOC_Os02g44700 | inflorescence & floral sporophyte, anther developmental series, mature anther | RING | Not Determined |
| LOC_Os02g44990 | vegetative body, callus | F-box | FBDUF |
| LOC_Os02g45240 | seed development | RING | H2 |
| LOC_Os02g45320 | vegetative body, shoot system, shoot and leaf tissues | F-box | FBX |
| LOC_Os02g45780 | inflorescence & floral sporophyte, anther developmental series, mature anther | RING | H2 |
| LOC_Os02g46100 | inflorescence & floral sporophyte, anther developmental series, mature anther | RING | H2 |
| LOC_Os02g46690 | seed development, seed | F-box | FBD |
| LOC_Os02g46740 | inflorescence & floral sporophyte, anther developmental series, mature anther | RING | Not Determined |
| LOC_Os02g47870 | inflorescence & floral sporophyte, anther developmental series, mature anther, male gametophytic lineage, sperm | RING | Not Determined |
| LOC_Os02g48300 | inflorescence & floral sporophyte, anther developmental series, mature anther | F-box | FBX |
| LOC_Os02g49550 | inflorescence & floral sporophyte, anther developmental series, mature anther, male gametophytic lineage, pollen | RING | H2 |
| LOC_Os02g50460 | inflorescence & floral sporophyte, anther developmental series, mature anther | U-box | III |
| LOC_Os02g51350 | inflorescence & floral sporophyte, anther developmental series, mature anther, male gametophytic lineage, sperm | F-box | FBK |
| LOC_Os02g52210 | inflorescence & floral sporophyte, anther developmental series, mature anther | RING | H2 |
| LOC_Os02g52230 | inflorescence & floral sporophyte, anther developmental series, mature anther | F-box |  |
| LOC_Os02g52870 | seed development, endosperm | RING | H2 |
| LOC_Os02g54240 | inflorescence & floral sporophyte, anther developmental series, mature anther | F-box | FBX |
| LOC_Os02g54550 | inflorescence & floral sporophyte, anther developmental series, mature anther | F-box | FBX |
| LOC_Os02g54830 | inflorescence & floral sporophyte, anther developmental series, mature anther | RING | H2 |
| LOC_Os02g55050 | inflorescence & floral sporophyte, anther developmental series, mature anther, male gametophytic lineage, sperm | F-box | FBX |
| LOC_Os02g55340 | vegetative body, root system | DWD | L |
| LOC_Os02g56760 | inflorescence & floral sporophyte, anther developmental series, mature anther, male gametophytic lineage, sperm | F-box | FBX |
| LOC_Os02g56820 | inflorescence & floral sporophyte, anther developmental series, mature anther, male gametophytic lineage, sperm | F-box | FBX |
| LOC_Os02g57700 | vegetative body, root system | U-box | IV |
| LOC_Os02g57890 | inflorescence & floral sporophyte | F-box | FBX |
| LOC_Os02g58540 | vegetative body, shoot system, node | RING | H2 |
| LOC_Os03g01660 | inflorescence & floral sporophyte, anther developmental series, mature anther, male gametophytic lineage | SKP1 |  |
| LOC_Os03g01720 | vegetative body, shoot system | RING | HC |
| LOC_Os03g01790 | inflorescence & floral sporophyte, anther developmental series, mature anther, male gametophytic lineage | RING | Not Determined |
| LOC_Os03g02210 | female gametophytic lineage, egg cell | F-box |  |
| LOC_Os03g03150 | inflorescence & floral sporophyte, anther developmental series, mature anther, male gametophytic lineage, sperm | TAD1 |  |
| LOC_Os03g04270 | inflorescence & floral sporophyte, anther developmental series, mature anther, male gametophytic lineage, sperm | F-box | FBL |
| LOC_Os03g04890 | inflorescence & floral sporophyte, anther developmental series, mature anther | RING | H2 |
| LOC_Os03g05560 | vegetative body, shoot system, shoot and leaf tissues | RING | H2 |
| LOC_Os03g06100 | inflorescence & floral sporophyte, anther developmental series, mature anther, male gametophytic lineage, pollen | F-box |  |
| LOC_Os03g07130 | vegetative body, shoot system, shoot and leaf tissues | RING | H2 |
| LOC_Os03g07530 | inflorescence & floral sporophyte, anther developmental series, mature anther | F-box | FBK |
| LOC_Os03g08000 | inflorescence & floral sporophyte, anther developmental series, mature anther, male gametophytic lineage, pollen | RING | HC |
| LOC_Os03g08830 | seed development | DWD | N |
| LOC_Os03g10880 | vegetative body, shoot system, shoot and leaf tissues | BTB |  |
| LOC_Os03g10890 | inflorescence & floral sporophyte, anther developmental series, mature anther, male gametophytic lineage, pollen | RING | H2 |
| LOC_Os03g11260 | female gametophytic lineage, post-fertilization | RING | HC |
| LOC_Os03g12190 | inflorescence & floral sporophyte, anther developmental series, mature anther | F-box | FBX |
| LOC_Os03g12200 | inflorescence & floral sporophyte, anther developmental series, mature anther | F-box | FBX |
| LOC_Os03g12940 | inflorescence & floral sporophyte, anther developmental series, mature anther, male gametophytic lineage | F-box | FBO |
| LOC_Os03g13100 | inflorescence & floral sporophyte, anther developmental series, mature anther, male gametophytic lineage, pollen | RING | Not Determined |
| LOC_Os03g13370 | inflorescence & floral sporophyte, anther developmental series, mature anther, male gametophytic lineage, sperm | APC |  |
| LOC_Os03g13740 | inflorescence & floral sporophyte, anther developmental series, mature anther | U-box | III |
| LOC_Os03g15000 | inflorescence & floral sporophyte, anther developmental series, mature anther | RING | HC |
| LOC_Os03g15730 | inflorescence & floral sporophyte, anther developmental series, mature anther, male gametophytic lineage, sperm | RING | HC |
| LOC_Os03g16480 | inflorescence & floral sporophyte, anther developmental series, mature anther, male gametophytic lineage, sperm | RING | H2 |
| LOC_Os03g16570 | inflorescence & floral sporophyte, anther developmental series, mature anther, male gametophytic lineage, pollen | RING | H2 |
| LOC_Os03g17170 | female gametophytic lineage, post-fertilization, seed development, embryo | RING | HC |
| LOC_Os03g18360 | inflorescence & floral sporophyte, anther developmental series, mature anther | BTB |  |
| LOC_Os03g19020 | inflorescence & floral sporophyte, anther developmental series, mature anther | RING | HC |
| LOC_Os03g20980 | female gametophytic lineage, post-fertilization, seed development, embryo | RING | H2 |
| LOC_Os03g22080 | inflorescence & floral sporophyte, anther developmental series, mature anther | RING | H2 |
| LOC_Os03g22110 | inflorescence & floral sporophyte, anther developmental series, mature anther | RING | H2 |
| LOC_Os03g22600 | vegetative body, root system | BTB |  |
| LOC_Os03g22830 | inflorescence & floral sporophyte, anther developmental series, mature anther, male gametophytic lineage, sperm | RING | H2 |
| LOC_Os03g22990 | inflorescence & floral sporophyte, anther developmental series | F-box | FBX |
| LOC_Os03g24200 | inflorescence & floral sporophyte, anther developmental series, mature anther | F-box | FBX |
| LOC_Os03g25190 | inflorescence & floral sporophyte, anther developmental series, mature anther | F-box | FBX |
| LOC_Os03g25220 | inflorescence & floral sporophyte, anther developmental series, mature anther, male gametophytic lineage | F-box | FBX |
| LOC_Os03g25240 | inflorescence & floral sporophyte, anther developmental series, mature anther | F-box | FBX |
| LOC_Os03g25250 | inflorescence & floral sporophyte, anther developmental series, mature anther | F-box | FBX |
| LOC_Os03g25640 | female gametophytic lineage | F-box | FBX |
| LOC_Os03g25650 | inflorescence & floral sporophyte, anther developmental series, mature anther | F-box | FBX |
| LOC_Os03g26370 | inflorescence & floral sporophyte, anther developmental series, mature anther | RING | H2 |
| LOC_Os03g27250 | inflorescence & floral sporophyte, anther developmental series, mature anther, male gametophytic lineage, sperm | F-box | FBO |
| LOC_Os03g27360 | vegetative body, root system | RING | H2 |
| LOC_Os03g27970 | seed development | DWD | A |
| LOC_Os03g30160 | inflorescence & floral sporophyte, gynoecium organs | F-box | FBK |
| LOC_Os03g30920 | inflorescence & floral sporophyte, anther developmental series, mature anther, male gametophytic lineage, sperm | F-box | FBX |
| LOC_Os03g31000 | inflorescence & floral sporophyte, anther developmental series, mature anther | U-box | IV |
| LOC_Os03g31400 | vegetative body, root system | U-box | I |
| LOC_Os03g36439 | inflorescence & floral sporophyte, anther developmental series, mature anther, male gametophytic lineage, pollen | F-box |  |
| LOC_Os03g40370 | inflorescence & floral sporophyte, anther developmental series, mature anther | F-box | FBDUF |
| LOC_Os03g42780 | female gametophytic lineage, post-fertilization | RING | Not Determined |
| LOC_Os03g42790 | female gametophytic lineage | RING | Not Determined |
| LOC_Os03g43060 | inflorescence & floral sporophyte, anther developmental series, mature anther | F-box | FBX |
| LOC_Os03g43390 | inflorescence & floral sporophyte, anther developmental series, mature anther, male gametophytic lineage | F-box |  |
| LOC_Os03g43890 | inflorescence & floral sporophyte, anther developmental series, mature anther, male gametophytic lineage, sperm | DWD | A |
| LOC_Os03g44636 | inflorescence & floral sporophyte, anther developmental series, mature anther | RING | H2 |
| LOC_Os03g44642 | inflorescence & floral sporophyte | RING | H2 |
| LOC_Os03g44810 | inflorescence & floral sporophyte, anther developmental series, mature anther, male gametophytic lineage, sperm | RING | HC |
| LOC_Os03g44920 | inflorescence & floral sporophyte | F-box | FBX |
| LOC_Os03g44980 | inflorescence & floral sporophyte | F-box | FBX |
| LOC_Os03g46120 | inflorescence & floral sporophyte, anther developmental series, mature anther | F-box | FBX |
| LOC_Os03g46140 | inflorescence & floral sporophyte, anther developmental series, mature anther | F-box | FBX |
| LOC_Os03g46500 | inflorescence & floral sporophyte, anther developmental series, mature anther | F-box | FBX |
| LOC_Os03g46510 | inflorescence & floral sporophyte, anther developmental series, mature anther, male gametophytic lineage, sperm | F-box | FBX |
| LOC_Os03g46530 | inflorescence & floral sporophyte, anther developmental series, mature anther | F-box | FBX |
| LOC_Os03g47420 | inflorescence & floral sporophyte, anther developmental series, mature anther | F-box | FBX |
| LOC_Os03g47500 | inflorescence & floral sporophyte, anther developmental series, mature anther, male gametophytic lineage, sperm | RING | HC |
| LOC_Os03g48120 | inflorescence & floral sporophyte, anther developmental series, mature anther, male gametophytic lineage | BTB |  |
| LOC_Os03g49250 | inflorescence & floral sporophyte, anther developmental series, mature anther, male gametophytic lineage, pollen | F-box | FBO |
| LOC_Os03g50050 | inflorescence & floral sporophyte, anther developmental series, mature anther, male gametophytic lineage | F-box | FBDUF |
| LOC_Os03g51270 | inflorescence & floral sporophyte, anther developmental series, mature anther, male gametophytic lineage, sperm | F-box | FBX |
| LOC_Os03g51760 | inflorescence & floral sporophyte, anther developmental series, mature anther, male gametophytic lineage, sperm | F-box | FBX |
| LOC_Os03g53080 | inflorescence & floral sporophyte, anther developmental series, mature anther, male gametophytic lineage, pollen | RING | HC |
| LOC_Os03g53510 | vegetative body, root system | DWD | A |
| LOC_Os03g56440 | inflorescence & floral sporophyte, anther developmental series, mature anther | F-box | FBX |
| LOC_Os03g56450 | inflorescence & floral sporophyte, anther developmental series, mature anther | F-box | FBX |
| LOC_Os03g56510 | inflorescence & floral sporophyte, anther developmental series, mature anther | F-box | FBX |
| LOC_Os03g57290 | female gametophytic lineage, post-fertilization, seed development, embryo | Cullin | Cullin 4 |
| LOC_Os03g57410 | vegetative body, root system | RING | H2 |
| LOC_Os03g57500 | inflorescence & floral sporophyte, anther developmental series, mature anther, male gametophytic lineage, pollen | RING | H2 |
| LOC_Os03g58390 | inflorescence & floral sporophyte, anther developmental series, mature anther, male gametophytic lineage, sperm | RING | HC |
| LOC_Os03g59540 | inflorescence & floral sporophyte, anther developmental series, mature anther, male gametophytic lineage, sperm | RING | HC |
| LOC_Os04g08470 | seed development, seed | F-box | FBX |
| LOC_Os04g10680 | inflorescence & floral sporophyte, anther developmental series, mature anther, male gametophytic lineage, pollen | RING | H2 |
| LOC_Os04g11450 | inflorescence & floral sporophyte, anther developmental series, mature anther, male gametophytic lineage, pollen | F-box | FBX |
| LOC_Os04g11660 | female gametophytic lineage, post-fertilization | F-box | FBX |
| LOC_Os04g11890 | inflorescence & floral sporophyte, anther developmental series, mature anther, male gametophytic lineage, sperm | F-box | FBX |
| LOC_Os04g12990 | seed development | F-box | FBX |
| LOC_Os04g13010 | inflorescence & floral sporophyte, anther developmental series, mature anther | F-box | FBX |
| LOC_Os04g13040 | inflorescence & floral sporophyte, anther developmental series, mature anther | F-box | FBX |
| LOC_Os04g13170 | inflorescence & floral sporophyte, anther developmental series, early anther | F-box | FBD |
| LOC_Os04g19750 | inflorescence & floral sporophyte, anther developmental series, mature anther | F-box | FBL |
| LOC_Os04g19800 | inflorescence & floral sporophyte, anther developmental series, mature anther | F-box |  |
| LOC_Os04g19810 | inflorescence & floral sporophyte, anther developmental series, mature anther | F-box | FBL |
| LOC_Os04g22240 | seed development, seed | RING | HC |
| LOC_Os04g26240 | inflorescence & floral sporophyte, anther developmental series, mature anther | F-box | FBDUF |
| LOC_Os04g28100 | inflorescence & floral sporophyte, anther developmental series, mature anther, male gametophytic lineage, sperm | U-box | II |
| LOC_Os04g30470 | inflorescence & floral sporophyte, anther developmental series, mature anther | U-box | V |
| LOC_Os04g30810 | inflorescence & floral sporophyte | F-box | FBX |
| LOC_Os04g30830 | seed development, seed | F-box |  |
| LOC_Os04g31390 | inflorescence & floral sporophyte, anther developmental series, mature anther | RING | H2 |
| LOC_Os04g31540 | inflorescence & floral sporophyte, anther developmental series, mature anther | F-box | FBX |
| LOC_Os04g31580 | inflorescence & floral sporophyte, anther developmental series, mature anther | F-box | FBX |
| LOC_Os04g34030 | vegetative body, root system | U-box | III |
| LOC_Os04g34140 | inflorescence & floral sporophyte, anther developmental series, mature anther | U-box | III |
| LOC_Os04g35190 | inflorescence & floral sporophyte, anther developmental series, mature anther, male gametophytic lineage, pollen | F-box | FBX |
| LOC_Os04g35340 | inflorescence & floral sporophyte, anther developmental series, mature anther, male gametophytic lineage, pollen | BTB |  |
| LOC_Os04g35930 | inflorescence & floral sporophyte, anther developmental series, mature anther | F-box | FBX |
| LOC_Os04g35940 | inflorescence & floral sporophyte, anther developmental series, mature anther, male gametophytic lineage, sperm | F-box | FBX |
| LOC_Os04g35990 | female gametophytic lineage, post-fertilization | F-box | FBX |
| LOC_Os04g36000 | female gametophytic lineage, post-fertilization | F-box | FBX |
| LOC_Os04g36020 | female gametophytic lineage, egg cell | F-box | FBX |
| LOC_Os04g37730 | vegetative body, root system | RING | H2 |
| LOC_Os04g38970 | seed development, endosperm | RING | Not Determined |
| LOC_Os04g39070 | inflorescence & floral sporophyte, anther developmental series, mature anther | F-box | FBX |
| LOC_Os04g39080 | inflorescence & floral sporophyte, anther developmental series, mature anther, male gametophytic lineage, sperm | F-box | FBX |
| LOC_Os04g40030 | inflorescence & floral sporophyte, anther developmental series, mature anther, male gametophytic lineage, sperm | F-box | FBO |
| LOC_Os04g40330 | inflorescence & floral sporophyte, anther developmental series, mature anther, male gametophytic lineage, pollen | F-box | FBX |
| LOC_Os04g40630 | vegetative body, callus | BTB |  |
| LOC_Os04g40760 | inflorescence & floral sporophyte, anther developmental series, mature anther | F-box | FBX |
| LOC_Os04g40770 | inflorescence & floral sporophyte, anther developmental series, mature anther | F-box | FBX |
| LOC_Os04g40780 | inflorescence & floral sporophyte, anther developmental series, mature anther | F-box | FBX |
| LOC_Os04g40800 | inflorescence & floral sporophyte, anther developmental series, mature anther | F-box | FBL |
| LOC_Os04g40920 | inflorescence & floral sporophyte, anther developmental series, mature anther, male gametophytic lineage, pollen | F-box |  |
| LOC_Os04g40960 | inflorescence & floral sporophyte, anther developmental series, mature anther | F-box | FBX |
| LOC_Os04g41050 | vegetative body, shoot system, shoot and leaf tissues | RING | H2 |
| LOC_Os04g41070 | inflorescence & floral sporophyte, anther developmental series, mature anther | RING | H2 |
| LOC_Os04g41080 | inflorescence & floral sporophyte, anther developmental series | RING | H2 |
| LOC_Os04g41250 | inflorescence & floral sporophyte, anther developmental series, mature anther | U-box | II |
| LOC_Os04g41470 | inflorescence & floral sporophyte, anther developmental series, mature anther, male gametophytic lineage, pollen | RING | Not Determined |
| LOC_Os04g42670 | inflorescence & floral sporophyte, anther developmental series, mature anther, male gametophytic lineage, pollen | F-box | FBL |
| LOC_Os04g42980 | vegetative body, root system | RING | H2 |
| LOC_Os04g43220 | inflorescence & floral sporophyte, anther developmental series, mature anther, male gametophytic lineage, sperm | RING | H2 |
| LOC_Os04g43300 | inflorescence & floral sporophyte, anther developmental series, mature anther, male gametophytic lineage, pollen | RING | HC |
| LOC_Os04g44820 | inflorescence & floral sporophyte, anther developmental series, mature anther, male gametophytic lineage, pollen | RING | HC |
| LOC_Os04g46450 | inflorescence & floral sporophyte, anther developmental series, mature anther, male gametophytic lineage, sperm | RING | HC |
| LOC_Os04g48050 | female gametophytic lineage, post-fertilization, seed development, embryo | RING | H2 |
| LOC_Os04g48270 | vegetative body, callus | F-box | FBX |
| LOC_Os04g49550 | inflorescence & floral sporophyte, anther developmental series, mature anther | RING | H2 |
| LOC_Os04g49950 | vegetative body, shoot system, shoot and leaf tissues | F-box | FBX |
| LOC_Os04g49970 | vegetative body, root system | U-box | III |
| LOC_Os04g50200 | inflorescence & floral sporophyte, anther developmental series, mature anther | F-box | FBX |
| LOC_Os04g52870 | inflorescence & floral sporophyte, anther developmental series, mature anther | F-box | FBW |
| LOC_Os04g53400 | inflorescence & floral sporophyte, anther developmental series, mature anther | BTB |  |
| LOC_Os04g53430 | inflorescence & floral sporophyte, anther developmental series, mature anther | BTB |  |
| LOC_Os04g53720 | inflorescence & floral sporophyte, anther developmental series, mature anther, male gametophytic lineage, sperm | RING | Not Determined |
| LOC_Os04g55000 | inflorescence & floral sporophyte, anther developmental series, mature anther | Cullin | Cullin 3b like |
| LOC_Os04g55030 | inflorescence & floral sporophyte, anther developmental series, mature anther, male gametophytic lineage, pollen | Cullin | Cullin 3a |
| LOC_Os04g56460 | female gametophytic lineage | BTB |  |
| LOC_Os04g57290 | inflorescence & floral sporophyte, anther developmental series, mature anther, male gametophytic lineage, pollen | F-box | FBX |
| LOC_Os04g57920 | inflorescence & floral sporophyte, anther developmental series, mature anther, male gametophytic lineage, pollen | F-box | FBX |
| LOC_Os04g58920 | vegetative body, root system | U-box | III |
| LOC_Os05g01620 | inflorescence & floral sporophyte, anther developmental series, mature anther | F-box | FBX |
| LOC_Os05g02550 | inflorescence & floral sporophyte, anther developmental series, mature anther | F-box |  |
| LOC_Os05g02570 | inflorescence & floral sporophyte, anther developmental series, mature anther | F-box |  |
| LOC_Os05g03100 | inflorescence & floral sporophyte, anther developmental series, mature anther | HECT |  |
| LOC_Os05g05700 | vegetative body, callus | Cullin | Cullin 1a-1 |
| LOC_Os05g05720 | inflorescence & floral sporophyte, anther developmental series, mature anther, male gametophytic lineage, sperm | APC |  |
| LOC_Os05g06270 | inflorescence & floral sporophyte, anther developmental series, mature anther, male gametophytic lineage, pollen | RING | H2 |
| LOC_Os05g06690 | seed development, endosperm | HECT |  |
| LOC_Os05g07070 | inflorescence & floral sporophyte, anther developmental series, mature anther, male gametophytic lineage, sperm | RING | H2 |
| LOC_Os05g07140 | inflorescence & floral sporophyte, anther developmental series, mature anther, male gametophytic lineage, pollen | RING | H2 |
| LOC_Os05g08350 | inflorescence & floral sporophyte, anther developmental series, mature anther, male gametophytic lineage | F-box | FBX |
| LOC_Os05g11190 | inflorescence & floral sporophyte, anther developmental series, mature anther, male gametophytic lineage, pollen | BTB |  |
| LOC_Os05g11720 | inflorescence & floral sporophyte, anther developmental series, mature anther, male gametophytic lineage, pollen | RING | V |
| LOC_Os05g11860 | inflorescence & floral sporophyte, anther developmental series, mature anther | RING | H2 |
| LOC_Os05g14860 | inflorescence & floral sporophyte, anther developmental series, mature anther, male gametophytic lineage, pollen | RING | HC |
| LOC_Os05g15170 | vegetative body, root system | RING | H2 |
| LOC_Os05g16660 | female gametophytic lineage, post-fertilization | DWD | D |
| LOC_Os05g19480 | vegetative body, root system | RING | HC |
| LOC_Os05g19970 | inflorescence & floral sporophyte, anther developmental series, mature anther, male gametophytic lineage, sperm | RING | HC |
| LOC_Os05g23810 | inflorescence & floral sporophyte, anther developmental series, mature anther | F-box |  |
| LOC_Os05g25180 | inflorescence & floral sporophyte, anther developmental series, mature anther, male gametophytic lineage | RING | HC |
| LOC_Os05g25580 | inflorescence & floral sporophyte, anther developmental series, mature anther, male gametophytic lineage, pollen | F-box | FBX |
| LOC_Os05g27550 | inflorescence & floral sporophyte, anther developmental series, mature anther | F-box | FBX |
| LOC_Os05g28730 | inflorescence & floral sporophyte, anther developmental series, mature anther | RING | V |
| LOC_Os05g29710 | vegetative body, root system | RING | H2 |
| LOC_Os05g30920 | inflorescence & floral sporophyte, anther developmental series, mature anther | F-box | FBX |
| LOC_Os05g33710 | inflorescence & floral sporophyte, anther developmental series, mature anther, male gametophytic lineage, sperm | DWD | A |
| LOC_Os05g33830 | vegetative body, shoot system, node | RING | H2 |
| LOC_Os05g38830 | inflorescence & floral sporophyte, anther developmental series, mature anther | HECT |  |
| LOC_Os05g39260 | inflorescence & floral sporophyte, anther developmental series, mature anther | RING | H2 |
| LOC_Os05g39300 | vegetative body, root system | F-box | FBL |
| LOC_Os05g39380 | inflorescence & floral sporophyte, anther developmental series, mature anther, male gametophytic lineage, pollen | RING | HC |
| LOC_Os05g39930 | inflorescence & floral sporophyte, anther developmental series, mature anther | U-box | II |
| LOC_Os05g39940 | inflorescence & floral sporophyte, anther developmental series, mature anther | RING | H2 |
| LOC_Os05g40500 | inflorescence & floral sporophyte, anther developmental series | F-box |  |
| LOC_Os05g40520 | inflorescence & floral sporophyte, anther developmental series | F-box | FBT |
| LOC_Os05g43610 | inflorescence & floral sporophyte, anther developmental series, mature anther, male gametophytic lineage, sperm | RING | HC |
| LOC_Os05g43850 | inflorescence & floral sporophyte, anther developmental series, mature anther, male gametophytic lineage, pollen | F-box | FBT |
| LOC_Os05g44540 | inflorescence & floral sporophyte, anther developmental series, mature anther | BTB |  |
| LOC_Os05g45040 | inflorescence & floral sporophyte, anther developmental series, mature anther, male gametophytic lineage | F-box | FBX |
| LOC_Os05g46050 | inflorescence & floral sporophyte, anther developmental series | F-box | FBDUF |
| LOC_Os05g46060 | inflorescence & floral sporophyte, anther developmental series | F-box | FBDUF |
| LOC_Os05g46160 | inflorescence & floral sporophyte, anther developmental series | F-box | FBDUF |
| LOC_Os05g46300 | inflorescence & floral sporophyte, anther developmental series, mature anther, male gametophytic lineage, sperm | F-box | FBX |
| LOC_Os05g46320 | inflorescence & floral sporophyte, anther developmental series, mature anther, male gametophytic lineage, pollen | F-box | FBX |
| LOC_Os05g47900 | vegetative body, root system | RING | V |
| LOC_Os05g48970 | inflorescence & floral sporophyte, anther developmental series, mature anther, male gametophytic lineage, sperm | RING | H2 |
| LOC_Os05g49400 | inflorescence & floral sporophyte, anther developmental series, mature anther | F-box | FBX |
| LOC_Os05g49450 | seed development, endosperm | F-box | FBX |
| LOC_Os05g49530 | inflorescence & floral sporophyte, anther developmental series, mature anther | F-box |  |
| LOC_Os05g49540 | inflorescence & floral sporophyte, anther developmental series, mature anther | F-box |  |
| LOC_Os05g49590 | inflorescence & floral sporophyte, anther developmental series, mature anther | DWD | L |
| LOC_Os05g49680 | inflorescence & floral sporophyte, anther developmental series, mature anther | F-box | FBX |
| LOC_Os05g49980 | inflorescence & floral sporophyte, anther developmental series, mature anther, male gametophytic lineage, pollen | F-box | FBL |
| LOC_Os05g51100 | inflorescence & floral sporophyte, anther developmental series, mature anther | F-box | FBX |
| LOC_Os05g51780 | vegetative body, root system | RING | H2 |
| LOC_Os06g02054 | inflorescence & floral sporophyte | F-box | FBX |
| LOC_Os06g02100 | female gametophytic lineage, post-fertilization | F-box | FBX |
| LOC_Os06g02110 | inflorescence & floral sporophyte, anther developmental series, mature anther, male gametophytic lineage, sperm | F-box | FBX |
| LOC_Os06g02400 | inflorescence & floral sporophyte, anther developmental series, mature anther, male gametophytic lineage, sperm | F-box | FBO |
| LOC_Os06g03840 | inflorescence & floral sporophyte, anther developmental series, mature anther, male gametophytic lineage, sperm | BTB |  |
| LOC_Os06g04980 | inflorescence & floral sporophyte, anther developmental series, mature anther, male gametophytic lineage | F-box | FBX |
| LOC_Os06g05200 | inflorescence & floral sporophyte, anther developmental series, mature anther, male gametophytic lineage, pollen | RING | H2 |
| LOC_Os06g05580 | inflorescence & floral sporophyte, anther developmental series, mature anther, male gametophytic lineage, sperm | F-box | FBDUF |
| LOC_Os06g05620 | inflorescence & floral sporophyte, anther developmental series, mature anther, male gametophytic lineage, sperm | F-box | FBDUF |
| LOC_Os06g06150 | seed development | RING | H2 |
| LOC_Os06g06600 | female gametophytic lineage, post-fertilization, seed development, embryo | F-box | FBX |
| LOC_Os06g07380 | inflorescence & floral sporophyte, anther developmental series, mature anther | F-box | FBX |
| LOC_Os06g07390 | inflorescence & floral sporophyte, anther developmental series, mature anther, male gametophytic lineage, pollen | F-box |  |
| LOC_Os06g07430 | inflorescence & floral sporophyte, anther developmental series, mature anther | F-box | FBX |
| LOC_Os06g07460 | inflorescence & floral sporophyte, anther developmental series, mature anther | F-box | FBX |
| LOC_Os06g08820 | vegetative body, root system | RING | H2 |
| LOC_Os06g09310 | inflorescence & floral sporophyte, anther developmental series, mature anther | RING | H2 |
| LOC_Os06g10290 | inflorescence & floral sporophyte, anther developmental series, mature anther, male gametophytic lineage, sperm | F-box | FBX |
| LOC_Os06g12560 | inflorescence & floral sporophyte, anther developmental series, mature anther, male gametophytic lineage, pollen | RING | H2 |
| LOC_Os06g13080 | vegetative body, root system | U-box | II |
| LOC_Os06g13090 | vegetative body, root system | U-box | II |
| LOC_Os06g13850 | inflorescence & floral sporophyte, anther developmental series, mature anther | F-box | FBX |
| LOC_Os06g13870 | inflorescence & floral sporophyte, anther developmental series, mature anther | U-box | III |
| LOC_Os06g13990 | inflorescence & floral sporophyte, anther developmental series, early anther | F-box | FBX |
| LOC_Os06g14200 | vegetative body, root system | RING | HC |
| LOC_Os06g14640 | inflorescence & floral sporophyte, anther developmental series, mature anther | RING | H2 |
| LOC_Os06g14650 | inflorescence & floral sporophyte, anther developmental series, mature anther | RING | H2 |
| LOC_Os06g16060 | vegetative body | RING | H2 |
| LOC_Os06g16940 | inflorescence & floral sporophyte, anther developmental series, mature anther | RING | H2 |
| LOC_Os06g17280 | inflorescence & floral sporophyte, anther developmental series, mature anther, male gametophytic lineage, sperm | RING | HC |
| LOC_Os06g19670 | seed development | F-box | FBX |
| LOC_Os06g21390 | inflorescence & floral sporophyte, anther developmental series, mature anther, male gametophytic lineage, pollen | RING | HC |
| LOC_Os06g29740 | seed development, seed | F-box | FBX |
| LOC_Os06g31100 | inflorescence & floral sporophyte, anther developmental series, mature anther | BTB |  |
| LOC_Os06g34360 | inflorescence & floral sporophyte, anther developmental series, mature anther, male gametophytic lineage, pollen | RING | H2 |
| LOC_Os06g34400 | inflorescence & floral sporophyte, anther developmental series, mature anther | RING | H2 |
| LOC_Os06g34430 | vegetative body, root system | RING | H2 |
| LOC_Os06g34470 | inflorescence & floral sporophyte, anther developmental series, mature anther | RING | H2 |
| LOC_Os06g34620 | vegetative body, root system | RING | H2 |
| LOC_Os06g34640 | inflorescence & floral sporophyte, anther developmental series, mature anther | RING | H2 |
| LOC_Os06g34880 | vegetative body, root system | RING | H2 |
| LOC_Os06g35080 | inflorescence & floral sporophyte, anther developmental series, mature anther | F-box |  |
| LOC_Os06g41520 | inflorescence & floral sporophyte, anther developmental series, mature anther | F-box | FBX |
| LOC_Os06g41750 | inflorescence & floral sporophyte, anther developmental series, mature anther, male gametophytic lineage, sperm | APC |  |
| LOC_Os06g42700 | inflorescence & floral sporophyte, anther developmental series, mature anther, male gametophytic lineage, pollen | RING | H2 |
| LOC_Os06g43740 | inflorescence & floral sporophyte, anther developmental series, mature anther | F-box | FBX |
| LOC_Os06g44275 | inflorescence & floral sporophyte, panicle | RING | Not Determined |
| LOC_Os06g44370 | inflorescence & floral sporophyte, anther developmental series, mature anther, male gametophytic lineage, sperm | DWD | A |
| LOC_Os06g44920 | inflorescence & floral sporophyte, anther developmental series | F-box | FBDUF |
| LOC_Os06g45460 | inflorescence & floral sporophyte, anther developmental series, early anther | F-box | FBX |
| LOC_Os06g45720 | inflorescence & floral sporophyte, anther developmental series, mature anther | BTB |  |
| LOC_Os06g47270 | inflorescence & floral sporophyte, anther developmental series, mature anther | RING | Not Determined |
| LOC_Os06g47400 | inflorescence & floral sporophyte, anther developmental series, mature anther | F-box | FBX |
| LOC_Os06g48040 | inflorescence & floral sporophyte, anther developmental series, mature anther | RING | H2 |
| LOC_Os06g49530 | inflorescence & floral sporophyte, anther developmental series, mature anther, male gametophytic lineage, sperm | F-box | FBX |
| LOC_Os06g50280 | vegetative body, shoot system, node | F-box | FBL |
| LOC_Os06g50880 | inflorescence & floral sporophyte, anther developmental series, mature anther, male gametophytic lineage, sperm | DWD | B |
| LOC_Os06g51130 | inflorescence & floral sporophyte, anther developmental series, mature anther | U-box | II |
| LOC_Os07g01140 | vegetative body, root system | BTB |  |
| LOC_Os07g02290 | female gametophytic lineage | F-box | FBX |
| LOC_Os07g02770 | female gametophytic lineage, post-fertilization, seed development, embryo | F-box | FBL |
| LOC_Os07g02910 | inflorescence & floral sporophyte, anther developmental series, mature anther, male gametophytic lineage, sperm | F-box | FBL |
| LOC_Os07g03090 | inflorescence & floral sporophyte, anther developmental series, mature anther | F-box | FBX |
| LOC_Os07g03110 | inflorescence & floral sporophyte, anther developmental series, mature anther, male gametophytic lineage | F-box | FBX |
| LOC_Os07g04750 | female gametophytic lineage, post-fertilization | F-box | FBX |
| LOC_Os07g04790 | inflorescence & floral sporophyte, anther developmental series, mature anther | F-box | FBX |
| LOC_Os07g05150 | inflorescence & floral sporophyte, anther developmental series, mature anther | SKP1 |  |
| LOC_Os07g05160 | inflorescence & floral sporophyte, anther developmental series, mature anther | SKP1 |  |
| LOC_Os07g05180 | inflorescence & floral sporophyte, anther developmental series, mature anther | SKP1 |  |
| LOC_Os07g06540 | vegetative body, root system | RING | H2 |
| LOC_Os07g06560 | vegetative body, root system | RING | H2 |
| LOC_Os07g06730 | inflorescence & floral sporophyte, anther developmental series, mature anther, male gametophytic lineage, pollen | F-box | FBX |
| LOC_Os07g07270 | inflorescence & floral sporophyte, anther developmental series, mature anther, male gametophytic lineage, sperm | BTB |  |
| LOC_Os07g08570 | inflorescence & floral sporophyte, anther developmental series, mature anther | F-box | FBX |
| LOC_Os07g09110 | inflorescence & floral sporophyte, anther developmental series, mature anther | F-box | FBX |
| LOC_Os07g10710 | inflorescence & floral sporophyte, anther developmental series, mature anther | F-box | FBX |
| LOC_Os07g13870 | inflorescence & floral sporophyte, anther developmental series, mature anther | F-box | FBX |
| LOC_Os07g13900 | female gametophytic lineage, post-fertilization | F-box | FBLD |
| LOC_Os07g13930 | inflorescence & floral sporophyte, anther developmental series, mature anther, male gametophytic lineage, sperm | F-box | FBX |
| LOC_Os07g16800 | inflorescence & floral sporophyte, anther developmental series, mature anther | F-box | FBX |
| LOC_Os07g17570 | inflorescence & floral sporophyte | F-box |  |
| LOC_Os07g18600 | inflorescence & floral sporophyte, anther developmental series, mature anther, male gametophytic lineage | F-box | FBL |
| LOC_Os07g22220 | inflorescence & floral sporophyte, anther developmental series, mature anther, male gametophytic lineage, sperm | DWD | K |
| LOC_Os07g22680 | inflorescence & floral sporophyte, anther developmental series | SKP1 |  |
| LOC_Os07g24190 | vegetative body, root system | RING | Not Determined |
| LOC_Os07g26490 | inflorescence & floral sporophyte, anther developmental series, mature anther, male gametophytic lineage, pollen | RING | HC |
| LOC_Os07g27030 | inflorescence & floral sporophyte, anther developmental series, mature anther, male gametophytic lineage, sperm | F-box | FBX |
| LOC_Os07g27950 | inflorescence & floral sporophyte, anther developmental series, mature anther, male gametophytic lineage, pollen | RING | H2 |
| LOC_Os07g31650 | inflorescence & floral sporophyte, anther developmental series, mature anther, male gametophytic lineage, pollen | RING | HC |
| LOC_Os07g31680 | inflorescence & floral sporophyte, anther developmental series, mature anther | F-box | FBL |
| LOC_Os07g31850 | inflorescence & floral sporophyte, anther developmental series, mature anther, male gametophytic lineage, sperm | RING | H2 |
| LOC_Os07g32730 | inflorescence & floral sporophyte, anther developmental series, mature anther, male gametophytic lineage, sperm | RING | HC |
| LOC_Os07g35070 | vegetative body, shoot system, node | F-box | FBX |
| LOC_Os07g36280 | inflorescence & floral sporophyte, anther developmental series, mature anther | F-box | FBX |
| LOC_Os07g36300 | female gametophytic lineage | F-box | FBX |
| LOC_Os07g36320 | seed development, seed | F-box | FBX |
| LOC_Os07g36330 | inflorescence & floral sporophyte, anther developmental series | F-box | FBX |
| LOC_Os07g36360 | inflorescence & floral sporophyte, anther developmental series, mature anther | F-box | FBO |
| LOC_Os07g36370 | inflorescence & floral sporophyte, anther developmental series | F-box | FBX |
| LOC_Os07g36520 | inflorescence & floral sporophyte, anther developmental series, mature anther | F-box | FBX |
| LOC_Os07g36530 | inflorescence & floral sporophyte, anther developmental series, mature anther | F-box | FBX |
| LOC_Os07g36830 | female gametophytic lineage | F-box | FBX |
| LOC_Os07g36840 | female gametophytic lineage, egg cell | F-box | FBX |
| LOC_Os07g36870 | inflorescence & floral sporophyte, anther developmental series, mature anther, male gametophytic lineage, sperm | F-box | FBX |
| LOC_Os07g36900 | inflorescence & floral sporophyte, anther developmental series, mature anther | F-box | FBL |
| LOC_Os07g36910 | female gametophytic lineage, post-fertilization | F-box | FBX |
| LOC_Os07g36920 | seed development | F-box | FBX |
| LOC_Os07g37140 | inflorescence & floral sporophyte, anther developmental series, mature anther, male gametophytic lineage, sperm | RING | Not Determined |
| LOC_Os07g38430 | inflorescence & floral sporophyte, anther developmental series, mature anther, male gametophytic lineage, sperm | DWD | D |
| LOC_Os07g39530 | vegetative body, root system | BTB |  |
| LOC_Os07g40030 | inflorescence & floral sporophyte, anther developmental series, mature anther, male gametophytic lineage, sperm | DWD | D |
| LOC_Os07g43180 | inflorescence & floral sporophyte, anther developmental series, early anther | SKP1 |  |
| LOC_Os07g43220 | inflorescence & floral sporophyte, anther developmental series | SKP1 |  |
| LOC_Os07g43230 | inflorescence & floral sporophyte, anther developmental series | SKP1 |  |
| LOC_Os07g43260 | inflorescence & floral sporophyte, anther developmental series, mature anther | SKP1 |  |
| LOC_Os07g43270 | inflorescence & floral sporophyte, anther developmental series | SKP1 |  |
| LOC_Os07g43380 | inflorescence & floral sporophyte, anther developmental series, mature anther | RING | HC |
| LOC_Os07g43740 | inflorescence & floral sporophyte, anther developmental series, mature anther, male gametophytic lineage, pollen | RING | H2 |
| LOC_Os07g43830 | inflorescence & floral sporophyte, anther developmental series, mature anther | RING | HC |
| LOC_Os07g45350 | inflorescence & floral sporophyte, anther developmental series, mature anther, male gametophytic lineage, sperm | RING | HC |
| LOC_Os07g46160 | seed development, endosperm | BTB |  |
| LOC_Os07g46555 | inflorescence & floral sporophyte, anther developmental series, mature anther, male gametophytic lineage, sperm | F-box |  |
| LOC_Os07g47170 | vegetative body, shoot system, shoot and leaf tissues | F-box |  |
| LOC_Os07g47650 | vegetative body, root system | F-box | FBK |
| LOC_Os07g48680 | vegetative body, shoot system, shoot and leaf tissues | RING | H2 |
| LOC_Os07g48940 | inflorescence & floral sporophyte, anther developmental series, mature anther | F-box | FBDUF |
| LOC_Os07g49030 | inflorescence & floral sporophyte, anther developmental series, mature anther, male gametophytic lineage, sperm | RING | HC |
| LOC_Os08g01320 | inflorescence & floral sporophyte, anther developmental series, mature anther, male gametophytic lineage, pollen | BTB |  |
| LOC_Os08g01900 | inflorescence & floral sporophyte, anther developmental series, mature anther, male gametophytic lineage, sperm | U-box | II |
| LOC_Os08g02140 | inflorescence & floral sporophyte, anther developmental series, mature anther, male gametophytic lineage, pollen | U-box | VII |
| LOC_Os08g03300 | female gametophytic lineage, post-fertilization | F-box | FBDUF |
| LOC_Os08g03470 | female gametophytic lineage | BTB |  |
| LOC_Os08g03480 | inflorescence & floral sporophyte, anther developmental series, mature anther, male gametophytic lineage | BTB |  |
| LOC_Os08g03490 | inflorescence & floral sporophyte, anther developmental series, mature anther, male gametophytic lineage | BTB |  |
| LOC_Os08g03500 | inflorescence & floral sporophyte, anther developmental series, mature anther | BTB |  |
| LOC_Os08g03530 | inflorescence & floral sporophyte, anther developmental series, mature anther | BTB |  |
| LOC_Os08g04270 | inflorescence & floral sporophyte, anther developmental series, mature anther, male gametophytic lineage | DWD | A |
| LOC_Os08g05560 | inflorescence & floral sporophyte, anther developmental series, mature anther, male gametophytic lineage, sperm | RING | H2 |
| LOC_Os08g06090 | inflorescence & floral sporophyte, anther developmental series, mature anther, male gametophytic lineage, pollen | RING | H2 |
| LOC_Os08g06510 | inflorescence & floral sporophyte, anther developmental series, mature anther, male gametophytic lineage, sperm | RING | HC |
| LOC_Os08g06710 | inflorescence & floral sporophyte, anther developmental series, mature anther | F-box | FBDUF |
| LOC_Os08g06780 | inflorescence & floral sporophyte, anther developmental series, mature anther, male gametophytic lineage | F-box | FBDUF |
| LOC_Os08g07400 | inflorescence & floral sporophyte, anther developmental series, mature anther, male gametophytic lineage, pollen | Cullin | Cullin 3b |
| LOC_Os08g08220 | inflorescence & floral sporophyte, anther developmental series, mature anther, male gametophytic lineage, sperm | RING | Not Determined |
| LOC_Os08g09220 | female gametophytic lineage | F-box | FBX |
| LOC_Os08g09380 | female gametophytic lineage, post-fertilization | F-box | FBX |
| LOC_Os08g09420 | inflorescence & floral sporophyte, anther developmental series, mature anther, male gametophytic lineage, pollen | F-box | FBX |
| LOC_Os08g09460 | inflorescence & floral sporophyte, anther developmental series, mature anther, male gametophytic lineage | F-box | FBX |
| LOC_Os08g09480 | female gametophytic lineage | F-box | FBX |
| LOC_Os08g09590 | inflorescence & floral sporophyte, anther developmental series, mature anther, male gametophytic lineage, pollen | F-box | FBL |
| LOC_Os08g09650 | female gametophytic lineage | F-box | FBX |
| LOC_Os08g09720 | female gametophytic lineage | F-box | FBX |
| LOC_Os08g09730 | inflorescence & floral sporophyte, anther developmental series, mature anther | F-box | FBX |
| LOC_Os08g09830 | vegetative body, shoot system, node | BTB |  |
| LOC_Os08g12960 | inflorescence & floral sporophyte, anther developmental series, mature anther | BTB |  |
| LOC_Os08g13030 | inflorescence & floral sporophyte | BTB |  |
| LOC_Os08g13060 | inflorescence & floral sporophyte, anther developmental series, mature anther, male gametophytic lineage | BTB |  |
| LOC_Os08g13090 | inflorescence & floral sporophyte, anther developmental series, early anther | BTB |  |
| LOC_Os08g15840 | vegetative body, root system | RING | HC |
| LOC_Os08g16630 | seed development, seed | F-box | FBX |
| LOC_Os08g16710 | inflorescence & floral sporophyte, anther developmental series, mature anther, male gametophytic lineage, pollen | F-box | FBX |
| LOC_Os08g16860 | seed development, seed | F-box | FBX |
| LOC_Os08g20492 | inflorescence & floral sporophyte, anther developmental series, mature anther, male gametophytic lineage, sperm | F-box | FBX |
| LOC_Os08g21660 | female gametophytic lineage, post-fertilization | DWD | D |
| LOC_Os08g24140 | inflorescence & floral sporophyte, anther developmental series, mature anther, male gametophytic lineage, pollen | F-box | FBL |
| LOC_Os08g24190 | inflorescence & floral sporophyte, anther developmental series, mature anther, male gametophytic lineage, sperm | F-box | FBX |
| LOC_Os08g24370 | inflorescence & floral sporophyte, anther developmental series, mature anther | F-box | FBX |
| LOC_Os08g24830 | inflorescence & floral sporophyte, anther developmental series, mature anther, male gametophytic lineage, pollen | F-box | FBDUF |
| LOC_Os08g25240 | inflorescence & floral sporophyte, anther developmental series, mature anther | BTB |  |
| LOC_Os08g27190 | inflorescence & floral sporophyte, anther developmental series, mature anther | F-box | FBX |
| LOC_Os08g28780 | vegetative body, root system | SKP1 |  |
| LOC_Os08g28800 | inflorescence & floral sporophyte, anther developmental series, mature anther | SKP1 |  |
| LOC_Os08g28940 | seed development | F-box | FBX |
| LOC_Os08g29590 | inflorescence & floral sporophyte, anther developmental series, mature anther, male gametophytic lineage, sperm | RING | V |
| LOC_Os08g31450 | female gametophytic lineage, post-fertilization | BTB |  |
| LOC_Os08g31560 | inflorescence & floral sporophyte, anther developmental series, mature anther, male gametophytic lineage | DWD | A |
| LOC_Os08g31690 | inflorescence & floral sporophyte, anther developmental series, mature anther, male gametophytic lineage, sperm | F-box | FBX |
| LOC_Os08g31720 | inflorescence & floral sporophyte, anther developmental series, mature anther | RING | H2 |
| LOC_Os08g31930 | inflorescence & floral sporophyte, anther developmental series, mature anther, male gametophytic lineage, sperm | RING | HC |
| LOC_Os08g32060 | inflorescence & floral sporophyte, anther developmental series, mature anther, male gametophytic lineage, pollen | U-box | II |
| LOC_Os08g33010 | inflorescence & floral sporophyte, anther developmental series, mature anther | F-box | FBDUF |
| LOC_Os08g33860 | inflorescence & floral sporophyte, anther developmental series, mature anther | RING | V |
| LOC_Os08g34820 | inflorescence & floral sporophyte, gynoecium organs | F-box | FBX |
| LOC_Os08g34860 | inflorescence & floral sporophyte, anther developmental series, mature anther | F-box | FBX |
| LOC_Os08g35060 | female gametophytic lineage | RING | Not Determined |
| LOC_Os08g35070 | female gametophytic lineage | RING | Not Determined |
| LOC_Os08g35870 | inflorescence & floral sporophyte, anther developmental series, mature anther, male gametophytic lineage | F-box | FBL |
| LOC_Os08g35880 | inflorescence & floral sporophyte, anther developmental series, mature anther, male gametophytic lineage | F-box | FBX |
| LOC_Os08g35900 | inflorescence & floral sporophyte, anther developmental series, mature anther | F-box | FBL |
| LOC_Os08g35930 | inflorescence & floral sporophyte, anther developmental series, mature anther, male gametophytic lineage, sperm | F-box | FBL |
| LOC_Os08g35960 | inflorescence & floral sporophyte, anther developmental series, mature anther, male gametophytic lineage, pollen | F-box | FBD |
| LOC_Os08g36960 | inflorescence & floral sporophyte, anther developmental series, mature anther, male gametophytic lineage, sperm | F-box | FBX |
| LOC_Os08g37570 | inflorescence & floral sporophyte, anther developmental series, mature anther | U-box | II |
| LOC_Os08g37760 | inflorescence & floral sporophyte, anther developmental series, mature anther | RING | H2 |
| LOC_Os08g38060 | inflorescence & floral sporophyte, anther developmental series, mature anther, male gametophytic lineage, pollen | RING | H2 |
| LOC_Os08g38330 | inflorescence & floral sporophyte, anther developmental series, mature anther | F-box | FBX |
| LOC_Os08g38470 | inflorescence & floral sporophyte, anther developmental series, mature anther | F-box | FBX |
| LOC_Os08g38480 | inflorescence & floral sporophyte | F-box | FBX |
| LOC_Os08g38490 | inflorescence & floral sporophyte, anther developmental series, mature anther | F-box | FBX |
| LOC_Os08g38520 | inflorescence & floral sporophyte, anther developmental series, mature anther | F-box | FBX |
| LOC_Os08g40460 | inflorescence & floral sporophyte, anther developmental series | BTB |  |
| LOC_Os08g40490 | inflorescence & floral sporophyte, anther developmental series, mature anther | BTB |  |
| LOC_Os08g40640 | inflorescence & floral sporophyte, anther developmental series, mature anther | F-box |  |
| LOC_Os08g41120 | inflorescence & floral sporophyte, anther developmental series, mature anther | BTB |  |
| LOC_Os08g41150 | inflorescence & floral sporophyte, anther developmental series, mature anther, male gametophytic lineage, sperm | BTB |  |
| LOC_Os08g41170 | inflorescence & floral sporophyte, anther developmental series, mature anther | BTB |  |
| LOC_Os08g41190 | inflorescence & floral sporophyte, anther developmental series, mature anther | BTB |  |
| LOC_Os08g41220 | female gametophytic lineage, egg cell | BTB |  |
| LOC_Os08g42640 | inflorescence & floral sporophyte, anther developmental series, mature anther | RING | H2 |
| LOC_Os08g43480 | seed development, endosperm | RING | H2 |
| LOC_Os08g44950 | inflorescence & floral sporophyte, anther developmental series, mature anther | RING | H2 |
| LOC_Os09g06620 | inflorescence & floral sporophyte, anther developmental series, mature anther, male gametophytic lineage, sperm | F-box | FBX |
| LOC_Os09g07900 | inflorescence & floral sporophyte, anther developmental series, mature anther, male gametophytic lineage, sperm | HECT |  |
| LOC_Os09g10020 | female gametophytic lineage, post-fertilization, seed development, embryo | SKP1 |  |
| LOC_Os09g10200 | inflorescence & floral sporophyte, anther developmental series, mature anther | SKP1 |  |
| LOC_Os09g10230 | inflorescence & floral sporophyte, anther developmental series, mature anther | SKP1 |  |
| LOC_Os09g10270 | inflorescence & floral sporophyte, anther developmental series, mature anther | SKP1 |  |
| LOC_Os09g12550 | inflorescence & floral sporophyte, anther developmental series, mature anther, male gametophytic lineage, pollen | DWD | C |
| LOC_Os09g15430 | inflorescence & floral sporophyte, anther developmental series, mature anther, male gametophytic lineage, sperm | RING | H2 |
| LOC_Os09g15440 | inflorescence & floral sporophyte, anther developmental series, mature anther | F-box | FBX |
| LOC_Os09g15460 | inflorescence & floral sporophyte, anther developmental series, mature anther | F-box |  |
| LOC_Os09g15560 | vegetative body, root system | F-box | FBX |
| LOC_Os09g15570 | inflorescence & floral sporophyte, anther developmental series, mature anther, male gametophytic lineage | F-box | FBX |
| LOC_Os09g16850 | inflorescence & floral sporophyte, anther developmental series, mature anther | BTB |  |
| LOC_Os09g16870 | inflorescence & floral sporophyte, anther developmental series, mature anther | BTB |  |
| LOC_Os09g17152 | inflorescence & floral sporophyte, anther developmental series, mature anther | F-box | FBX |
| LOC_Os09g17190 | inflorescence & floral sporophyte, anther developmental series, mature anther | F-box | FBX |
| LOC_Os09g20550 | seed development, seed | RING | HC |
| LOC_Os09g20650 | inflorescence & floral sporophyte, anther developmental series, mature anther, male gametophytic lineage | F-box | FBX |
| LOC_Os09g21580 | inflorescence & floral sporophyte, anther developmental series, mature anther | F-box | FBX |
| LOC_Os09g21620 | inflorescence & floral sporophyte, anther developmental series, mature anther | F-box | FBX |
| LOC_Os09g22460 | inflorescence & floral sporophyte, anther developmental series, mature anther | F-box | FBX |
| LOC_Os09g24650 | inflorescence & floral sporophyte, anther developmental series, mature anther, male gametophytic lineage | RING | V |
| LOC_Os09g25190 | female gametophytic lineage | RING | HC |
| LOC_Os09g25200 | female gametophytic lineage, post-fertilization | RING | Not Determined |
| LOC_Os09g25220 | inflorescence & floral sporophyte, anther developmental series, mature anther, male gametophytic lineage, sperm | RING | Not Determined |
| LOC_Os09g25260 | inflorescence & floral sporophyte, anther developmental series, mature anther, male gametophytic lineage, sperm | RING | Not Determined |
| LOC_Os09g26400 | inflorescence & floral sporophyte, anther developmental series, mature anther, male gametophytic lineage, pollen | RING | H2 |
| LOC_Os09g27090 | female gametophytic lineage, post-fertilization | F-box | FBLD |
| LOC_Os09g27100 | inflorescence & floral sporophyte, anther developmental series, mature anther | F-box | FBX |
| LOC_Os09g27380 | vegetative body | RING | H2 |
| LOC_Os09g27570 | inflorescence & floral sporophyte, anther developmental series, mature anther, male gametophytic lineage, sperm | F-box | FBA |
| LOC_Os09g27660 | inflorescence & floral sporophyte, anther developmental series, mature anther, male gametophytic lineage, pollen | F-box | FBO |
| LOC_Os09g28120 | inflorescence & floral sporophyte, anther developmental series, mature anther, male gametophytic lineage, pollen | F-box | FBX |
| LOC_Os09g29310 | vegetative body, root system | RING | H2 |
| LOC_Os09g30180 | inflorescence & floral sporophyte, anther developmental series, mature anther, male gametophytic lineage, sperm | F-box | FBX |
| LOC_Os09g32410 | inflorescence & floral sporophyte, anther developmental series, mature anther, male gametophytic lineage, sperm | F-box | FBX |
| LOC_Os09g32600 | female gametophytic lineage, egg cell | F-box | FBX |
| LOC_Os09g32690 | inflorescence & floral sporophyte, anther developmental series, mature anther, male gametophytic lineage, sperm | RING | HC |
| LOC_Os09g32730 | seed development, endosperm | RING | HC |
| LOC_Os09g32870 | inflorescence & floral sporophyte, anther developmental series, mature anther, male gametophytic lineage, pollen | F-box | FBX |
| LOC_Os09g33670 | inflorescence & floral sporophyte, anther developmental series, mature anther, male gametophytic lineage | RING | H2 |
| LOC_Os09g33740 | inflorescence & floral sporophyte, anther developmental series, mature anther, male gametophytic lineage, sperm | RING | HC |
| LOC_Os09g34200 | inflorescence & floral sporophyte, anther developmental series, mature anther, male gametophytic lineage, sperm | F-box | FBX |
| LOC_Os09g36830 | vegetative body, callus | SKP1 |  |
| LOC_Os09g37050 | vegetative body, shoot system, shoot and leaf tissues | RING | H2 |
| LOC_Os09g37570 | female gametophytic lineage, egg cell | F-box | FBX |
| LOC_Os09g38630 | inflorescence & floral sporophyte, anther developmental series, mature anther | RING | Not Determined |
| LOC_Os09g38640 | inflorescence & floral sporophyte, anther developmental series, mature anther | RING | Not Determined |
| LOC_Os09g38900 | inflorescence & floral sporophyte, anther developmental series, mature anther, male gametophytic lineage, sperm | F-box |  |
| LOC_Os09g39000 | inflorescence & floral sporophyte, anther developmental series, mature anther | F-box | FBX |
| LOC_Os09g39050 | inflorescence & floral sporophyte, anther developmental series, mature anther | F-box | FBX |
| LOC_Os09g39190 | female gametophytic lineage, egg cell | F-box | FBO |
