## Supplementary material for "RE3DB: A multi-omics phylogenomics platform for rice E3 ubiquitin ligases identifies novel regulators of pollen germination": Table S7

| **Group** | **category** | **subcategory** | **gene_count** |
| --- | --- | --- | --- |
| female gametophytic lineage | BTB |  | 6 |
| female gametophytic lineage | Cullin | Cullin 1d | 1 |
| female gametophytic lineage | F-box | FBL | 2 |
| female gametophytic lineage | F-box | FBX | 14 |
| female gametophytic lineage | RING | H2 | 1 |
| female gametophytic lineage | RING | HC | 1 |
| female gametophytic lineage | RING | Not Determined | 3 |
| female gametophytic lineage, egg cell | APC |  | 1 |
| female gametophytic lineage, egg cell | BTB |  | 1 |
| female gametophytic lineage, egg cell | DWD | D | 1 |
| female gametophytic lineage, egg cell | F-box | FBO | 1 |
| female gametophytic lineage, egg cell | F-box | FBX | 5 |
| female gametophytic lineage, egg cell | F-box |  | 2 |
| female gametophytic lineage, post-fertilization | BTB |  | 1 |
| female gametophytic lineage, post-fertilization | DWD | D | 2 |
| female gametophytic lineage, post-fertilization | F-box | FBDUF | 3 |
| female gametophytic lineage, post-fertilization | F-box | FBLD | 2 |
| female gametophytic lineage, post-fertilization | F-box | FBO | 1 |
| female gametophytic lineage, post-fertilization | F-box | FBX | 12 |
| female gametophytic lineage, post-fertilization | RING | H2 | 1 |
| female gametophytic lineage, post-fertilization | RING | HC | 1 |
| female gametophytic lineage, post-fertilization | RING | Not Determined | 2 |
| female gametophytic lineage, post-fertilization, seed development, embryo | Cullin | Cullin 4 | 1 |
| female gametophytic lineage, post-fertilization, seed development, embryo | DWD | B | 1 |
| female gametophytic lineage, post-fertilization, seed development, embryo | F-box | FBDUF | 1 |
| female gametophytic lineage, post-fertilization, seed development, embryo | F-box | FBL | 1 |
| female gametophytic lineage, post-fertilization, seed development, embryo | F-box | FBX | 3 |
| female gametophytic lineage, post-fertilization, seed development, embryo | RING | H2 | 2 |
| female gametophytic lineage, post-fertilization, seed development, embryo | RING | HC | 1 |
| female gametophytic lineage, post-fertilization, seed development, embryo | SKP1 |  | 1 |
| inflorescence & floral sporophyte | BTB |  | 2 |
| inflorescence & floral sporophyte | F-box | FBDUF | 1 |
| inflorescence & floral sporophyte | F-box | FBL | 1 |
| inflorescence & floral sporophyte | F-box | FBX | 7 |
| inflorescence & floral sporophyte | F-box |  | 1 |
| inflorescence & floral sporophyte | RING | H2 | 5 |
| inflorescence & floral sporophyte | RING | HC | 1 |
| inflorescence & floral sporophyte, anther developmental series | BTB |  | 3 |
| inflorescence & floral sporophyte, anther developmental series | F-box | FBDUF | 4 |
| inflorescence & floral sporophyte, anther developmental series | F-box | FBT | 1 |
| inflorescence & floral sporophyte, anther developmental series | F-box | FBX | 6 |
| inflorescence & floral sporophyte, anther developmental series | F-box |  | 1 |
| inflorescence & floral sporophyte, anther developmental series | RING | H2 | 1 |
| inflorescence & floral sporophyte, anther developmental series | SKP1 |  | 4 |
| inflorescence & floral sporophyte, anther developmental series, early anther | APC |  | 1 |
| inflorescence & floral sporophyte, anther developmental series, early anther | BTB |  | 1 |
| inflorescence & floral sporophyte, anther developmental series, early anther | F-box | FBD | 2 |
| inflorescence & floral sporophyte, anther developmental series, early anther | F-box | FBDUF | 2 |
| inflorescence & floral sporophyte, anther developmental series, early anther | F-box | FBX | 4 |
| inflorescence & floral sporophyte, anther developmental series, early anther | SKP1 |  | 1 |
| inflorescence & floral sporophyte, anther developmental series, mature anther | BTB |  | 36 |
| inflorescence & floral sporophyte, anther developmental series, mature anther | Cullin | Cullin 3b like | 1 |
| inflorescence & floral sporophyte, anther developmental series, mature anther | DWD | C | 1 |
| inflorescence & floral sporophyte, anther developmental series, mature anther | DWD | I | 1 |
| inflorescence & floral sporophyte, anther developmental series, mature anther | DWD | L | 1 |
| inflorescence & floral sporophyte, anther developmental series, mature anther | F-box | FBDUF | 12 |
| inflorescence & floral sporophyte, anther developmental series, mature anther | F-box | FBK | 2 |
| inflorescence & floral sporophyte, anther developmental series, mature anther | F-box | FBL | 8 |
| inflorescence & floral sporophyte, anther developmental series, mature anther | F-box | FBO | 2 |
| inflorescence & floral sporophyte, anther developmental series, mature anther | F-box | FBW | 1 |
| inflorescence & floral sporophyte, anther developmental series, mature anther | F-box | FBX | 105 |
| inflorescence & floral sporophyte, anther developmental series, mature anther | F-box |  | 15 |
| inflorescence & floral sporophyte, anther developmental series, mature anther | HECT |  | 2 |
| inflorescence & floral sporophyte, anther developmental series, mature anther | RING | H2 | 42 |
| inflorescence & floral sporophyte, anther developmental series, mature anther | RING | HC | 12 |
| inflorescence & floral sporophyte, anther developmental series, mature anther | RING | Not Determined | 9 |
| inflorescence & floral sporophyte, anther developmental series, mature anther | RING | V | 2 |
| inflorescence & floral sporophyte, anther developmental series, mature anther | SKP1 |  | 10 |
| inflorescence & floral sporophyte, anther developmental series, mature anther | U-box | II | 5 |
| inflorescence & floral sporophyte, anther developmental series, mature anther | U-box | III | 7 |
| inflorescence & floral sporophyte, anther developmental series, mature anther | U-box | IV | 1 |
| inflorescence & floral sporophyte, anther developmental series, mature anther | U-box | V | 2 |
| inflorescence & floral sporophyte, anther developmental series, mature anther | U-box | VI | 1 |
| inflorescence & floral sporophyte, anther developmental series, mature anther, male gametophytic lineage | BTB |  | 4 |
| inflorescence & floral sporophyte, anther developmental series, mature anther, male gametophytic lineage | DWD | A | 2 |
| inflorescence & floral sporophyte, anther developmental series, mature anther, male gametophytic lineage | DWD | C | 1 |
| inflorescence & floral sporophyte, anther developmental series, mature anther, male gametophytic lineage | F-box | FBDUF | 2 |
| inflorescence & floral sporophyte, anther developmental series, mature anther, male gametophytic lineage | F-box | FBL | 2 |
| inflorescence & floral sporophyte, anther developmental series, mature anther, male gametophytic lineage | F-box | FBO | 1 |
| inflorescence & floral sporophyte, anther developmental series, mature anther, male gametophytic lineage | F-box | FBX | 13 |
| inflorescence & floral sporophyte, anther developmental series, mature anther, male gametophytic lineage | F-box |  | 2 |
| inflorescence & floral sporophyte, anther developmental series, mature anther, male gametophytic lineage | RING | H2 | 4 |
| inflorescence & floral sporophyte, anther developmental series, mature anther, male gametophytic lineage | RING | HC | 2 |
| inflorescence & floral sporophyte, anther developmental series, mature anther, male gametophytic lineage | RING | Not Determined | 1 |
| inflorescence & floral sporophyte, anther developmental series, mature anther, male gametophytic lineage | RING | V | 1 |
| inflorescence & floral sporophyte, anther developmental series, mature anther, male gametophytic lineage | SKP1 |  | 1 |
| inflorescence & floral sporophyte, anther developmental series, mature anther, male gametophytic lineage | U-box | IV | 1 |
| inflorescence & floral sporophyte, anther developmental series, mature anther, male gametophytic lineage, pollen | BTB |  | 6 |
| inflorescence & floral sporophyte, anther developmental series, mature anther, male gametophytic lineage, pollen | Cullin | Cullin 1a | 1 |
| inflorescence & floral sporophyte, anther developmental series, mature anther, male gametophytic lineage, pollen | Cullin | Cullin 3a | 1 |
| inflorescence & floral sporophyte, anther developmental series, mature anther, male gametophytic lineage, pollen | Cullin | Cullin 3b | 1 |
| inflorescence & floral sporophyte, anther developmental series, mature anther, male gametophytic lineage, pollen | DWD | C | 1 |
| inflorescence & floral sporophyte, anther developmental series, mature anther, male gametophytic lineage, pollen | DWD | D | 1 |
| inflorescence & floral sporophyte, anther developmental series, mature anther, male gametophytic lineage, pollen | F-box | FBD | 2 |
| inflorescence & floral sporophyte, anther developmental series, mature anther, male gametophytic lineage, pollen | F-box | FBDUF | 5 |
| inflorescence & floral sporophyte, anther developmental series, mature anther, male gametophytic lineage, pollen | F-box | FBL | 4 |
| inflorescence & floral sporophyte, anther developmental series, mature anther, male gametophytic lineage, pollen | F-box | FBO | 4 |
| inflorescence & floral sporophyte, anther developmental series, mature anther, male gametophytic lineage, pollen | F-box | FBT | 1 |
| inflorescence & floral sporophyte, anther developmental series, mature anther, male gametophytic lineage, pollen | F-box | FBX | 25 |
| inflorescence & floral sporophyte, anther developmental series, mature anther, male gametophytic lineage, pollen | F-box |  | 5 |
| inflorescence & floral sporophyte, anther developmental series, mature anther, male gametophytic lineage, pollen | HECT |  | 1 |
| inflorescence & floral sporophyte, anther developmental series, mature anther, male gametophytic lineage, pollen | RING | H2 | 24 |
| inflorescence & floral sporophyte, anther developmental series, mature anther, male gametophytic lineage, pollen | RING | HC | 16 |
| inflorescence & floral sporophyte, anther developmental series, mature anther, male gametophytic lineage, pollen | RING | Not Determined | 3 |
| inflorescence & floral sporophyte, anther developmental series, mature anther, male gametophytic lineage, pollen | RING | V | 3 |
| inflorescence & floral sporophyte, anther developmental series, mature anther, male gametophytic lineage, pollen | U-box | II | 1 |
| inflorescence & floral sporophyte, anther developmental series, mature anther, male gametophytic lineage, pollen | U-box | VII | 1 |
| inflorescence & floral sporophyte, anther developmental series, mature anther, male gametophytic lineage, sperm | APC |  | 3 |
| inflorescence & floral sporophyte, anther developmental series, mature anther, male gametophytic lineage, sperm | BTB |  | 4 |
| inflorescence & floral sporophyte, anther developmental series, mature anther, male gametophytic lineage, sperm | DWD | A | 3 |
| inflorescence & floral sporophyte, anther developmental series, mature anther, male gametophytic lineage, sperm | DWD | B | 2 |
| inflorescence & floral sporophyte, anther developmental series, mature anther, male gametophytic lineage, sperm | DWD | D | 3 |
| inflorescence & floral sporophyte, anther developmental series, mature anther, male gametophytic lineage, sperm | DWD | K | 1 |
| inflorescence & floral sporophyte, anther developmental series, mature anther, male gametophytic lineage, sperm | F-box | FBA | 1 |
| inflorescence & floral sporophyte, anther developmental series, mature anther, male gametophytic lineage, sperm | F-box | FBD | 1 |
| inflorescence & floral sporophyte, anther developmental series, mature anther, male gametophytic lineage, sperm | F-box | FBDUF | 2 |
| inflorescence & floral sporophyte, anther developmental series, mature anther, male gametophytic lineage, sperm | F-box | FBK | 5 |
| inflorescence & floral sporophyte, anther developmental series, mature anther, male gametophytic lineage, sperm | F-box | FBL | 6 |
| inflorescence & floral sporophyte, anther developmental series, mature anther, male gametophytic lineage, sperm | F-box | FBLD | 1 |
| inflorescence & floral sporophyte, anther developmental series, mature anther, male gametophytic lineage, sperm | F-box | FBO | 4 |
| inflorescence & floral sporophyte, anther developmental series, mature anther, male gametophytic lineage, sperm | F-box | FBT | 2 |
| inflorescence & floral sporophyte, anther developmental series, mature anther, male gametophytic lineage, sperm | F-box | FBW | 1 |
| inflorescence & floral sporophyte, anther developmental series, mature anther, male gametophytic lineage, sperm | F-box | FBX | 45 |
| inflorescence & floral sporophyte, anther developmental series, mature anther, male gametophytic lineage, sperm | F-box |  | 5 |
| inflorescence & floral sporophyte, anther developmental series, mature anther, male gametophytic lineage, sperm | HECT |  | 2 |
| inflorescence & floral sporophyte, anther developmental series, mature anther, male gametophytic lineage, sperm | RING | C2 | 1 |
| inflorescence & floral sporophyte, anther developmental series, mature anther, male gametophytic lineage, sperm | RING | H2 | 12 |
| inflorescence & floral sporophyte, anther developmental series, mature anther, male gametophytic lineage, sperm | RING | HC | 20 |
| inflorescence & floral sporophyte, anther developmental series, mature anther, male gametophytic lineage, sperm | RING | Not Determined | 9 |
| inflorescence & floral sporophyte, anther developmental series, mature anther, male gametophytic lineage, sperm | RING | V | 3 |
| inflorescence & floral sporophyte, anther developmental series, mature anther, male gametophytic lineage, sperm | TAD1 |  | 1 |
| inflorescence & floral sporophyte, anther developmental series, mature anther, male gametophytic lineage, sperm | U-box | II | 3 |
| inflorescence & floral sporophyte, anther developmental series, mature anther, male gametophytic lineage, sperm | U-box | IV | 1 |
| inflorescence & floral sporophyte, gynoecium organs | F-box | FBDUF | 1 |
| inflorescence & floral sporophyte, gynoecium organs | F-box | FBK | 1 |
| inflorescence & floral sporophyte, gynoecium organs | F-box | FBO | 1 |
| inflorescence & floral sporophyte, gynoecium organs | F-box | FBX | 1 |
| inflorescence & floral sporophyte, gynoecium organs | RING | HC | 1 |
| inflorescence & floral sporophyte, panicle | RING | H2 | 1 |
| inflorescence & floral sporophyte, panicle | RING | Not Determined | 3 |
| seed development | DWD | A | 1 |
| seed development | DWD | B | 1 |
| seed development | DWD | N | 1 |
| seed development | F-box | FBX | 6 |
| seed development | RING | H2 | 4 |
| seed development | RING | HC | 1 |
| seed development | RING | Not Determined | 1 |
| seed development, endosperm | BTB |  | 1 |
| seed development, endosperm | F-box | FBX | 3 |
| seed development, endosperm | HECT |  | 1 |
| seed development, endosperm | RING | H2 | 3 |
| seed development, endosperm | RING | HC | 1 |
| seed development, endosperm | RING | Not Determined | 2 |
| seed development, seed | BTB |  | 1 |
| seed development, seed | F-box | FBD | 1 |
| seed development, seed | F-box | FBK | 1 |
| seed development, seed | F-box | FBX | 8 |
| seed development, seed | F-box |  | 1 |
| seed development, seed | RING | H2 | 1 |
| seed development, seed | RING | HC | 2 |
| seed development, seed | RING | Not Determined | 1 |
| vegetative body | DWD | D | 1 |
| vegetative body | F-box | FBX | 1 |
| vegetative body | RING | H2 | 3 |
| vegetative body, callus | BTB |  | 1 |
| vegetative body, callus | Cullin | Cullin 1a-1 | 1 |
| vegetative body, callus | F-box | FBDUF | 1 |
| vegetative body, callus | F-box | FBX | 1 |
| vegetative body, callus | RING | H2 | 2 |
| vegetative body, callus | SKP1 |  | 1 |
| vegetative body, root system | BTB |  | 5 |
| vegetative body, root system | DWD | A | 1 |
| vegetative body, root system | DWD | L | 1 |
| vegetative body, root system | F-box | FBK | 1 |
| vegetative body, root system | F-box | FBL | 1 |
| vegetative body, root system | F-box | FBT | 1 |
| vegetative body, root system | F-box | FBX | 3 |
| vegetative body, root system | RING | H2 | 17 |
| vegetative body, root system | RING | HC | 4 |
| vegetative body, root system | RING | Not Determined | 1 |
| vegetative body, root system | RING | V | 2 |
| vegetative body, root system | SKP1 |  | 1 |
| vegetative body, root system | U-box | I | 1 |
| vegetative body, root system | U-box | II | 2 |
| vegetative body, root system | U-box | III | 4 |
| vegetative body, root system | U-box | IV | 2 |
| vegetative body, root system | U-box | VII | 1 |
| vegetative body, shoot system | RING | HC | 1 |
| vegetative body, shoot system, node | BTB |  | 1 |
| vegetative body, shoot system, node | F-box | FBL | 1 |
| vegetative body, shoot system, node | F-box | FBX | 2 |
| vegetative body, shoot system, node | RING | H2 | 4 |
| vegetative body, shoot system, shoot and leaf tissues | BTB |  | 1 |
| vegetative body, shoot system, shoot and leaf tissues | Cullin | Cullin 1e | 1 |
| vegetative body, shoot system, shoot and leaf tissues | DWD | L | 1 |
| vegetative body, shoot system, shoot and leaf tissues | F-box | FBX | 3 |
| vegetative body, shoot system, shoot and leaf tissues | F-box |  | 1 |
| vegetative body, shoot system, shoot and leaf tissues | RING | H2 | 9 |
| vegetative body, shoot system, shoot and leaf tissues | RING | Not Determined | 2 |
| vegetative body, shoot system, shoot and leaf tissues | SKP1 |  | 1 |
