## Supplementary material for "RE3DB: A multi-omics phylogenomics platform for rice E3 ubiquitin ligases identifies novel regulators of pollen germination": Table S8

| **E3 Family** | **Term** | **Overlap** | **P-value** | **Adjusted P-value** | **Odds Ratio** | **Combined Score** | **Genes** |
| --- | --- | --- | --- | --- | --- | --- | --- |
| U-box | root system | 10/2447 | 0.001942631 | 0.015541047 | 3.400683761 | 21.23289032 | LOC_Os06g13080;LOC_Os06g13090;LOC_Os04g34030;LOC_Os10g40120;LOC_Os12g06410;LOC_Os02g57700;LOC_Os04g49970;LOC_Os04g58920;LOC_Os10g40490;LOC_Os03g31400 |
| F-box | mature anther | 284/12828 | 2.00575E-22 | 4.21208E-21 | 2.165016805 | 108.1660846 | LOC_Os08g34860;LOC_Os03g12940;LOC_Os04g40770;LOC_Os04g35930;LOC_Os06g13850;LOC_Os07g36360;LOC_Os08g35870;LOC_Os04g11890;LOC_Os01g45020;LOC_Os12g27810;LOC_Os01g27680;LOC_Os01g33490;LOC_Os10g06700;LOC_Os04g13010;LOC_Os02g41930;LOC_Os10g03600;LOC_Os01g71440;LOC_Os12g27760;LOC_Os06g43740;LOC_Os02g05700;LOC_Os01g64030;LOC_Os12g41630;LOC_Os03g27250;LOC_Os06g35080;LOC_Os10g10420;LOC_Os10g41829;LOC_Os01g06360;LOC_Os10g41650;LOC_Os01g46500;LOC_Os02g16760;LOC_Os04g19750;LOC_Os02g21110;LOC_Os03g36439;LOC_Os07g09110;LOC_Os10g25680;LOC_Os11g38130;LOC_Os08g16710;LOC_Os10g10390;LOC_Os11g39609;LOC_Os08g24370;LOC_Os01g36940;LOC_Os08g35880;LOC_Os03g12190;LOC_Os10g04370;LOC_Os05g45040;LOC_Os09g21580;LOC_Os03g25240;LOC_Os12g31340;LOC_Os05g27550;LOC_Os03g06100;LOC_Os09g39000;LOC_Os02g56760;LOC_Os02g56820;LOC_Os06g04980;LOC_Os07g48940;LOC_Os09g20650;LOC_Os05g25580;LOC_Os01g17390;LOC_Os04g31540;LOC_Os06g07380;LOC_Os05g49400;LOC_Os08g38520;LOC_Os09g15460;LOC_Os02g04000;LOC_Os10g35920;LOC_Os11g42240;LOC_Os05g30920;LOC_Os09g32870;LOC_Os06g41520;LOC_Os12g30990;LOC_Os10g37570;LOC_Os12g27790;LOC_Os02g03910;LOC_Os07g36520;LOC_Os03g25650;LOC_Os11g33220;LOC_Os04g39080;LOC_Os01g07160;LOC_Os10g05000;LOC_Os08g06710;LOC_Os09g22460;LOC_Os06g05580;LOC_Os02g35530;LOC_Os11g41570;LOC_Os02g36520;LOC_Os11g03680;LOC_Os04g40760;LOC_Os03g46120;LOC_Os05g08350;LOC_Os04g57290;LOC_Os10g17930;LOC_Os03g25190;LOC_Os01g56750;LOC_Os01g28150;LOC_Os04g19800;LOC_Os10g03620;LOC_Os04g40030;LOC_Os11g42160;LOC_Os09g17190;LOC_Os07g31680;LOC_Os08g24830;LOC_Os10g34340;LOC_Os08g24140;LOC_Os04g40960;LOC_Os10g05800;LOC_Os07g08570;LOC_Os11g41560;LOC_Os12g30950;LOC_Os05g02570;LOC_Os08g36960;LOC_Os11g07970;LOC_Os04g26240;LOC_Os02g17210;LOC_Os06g49530;LOC_Os10g24900;LOC_Os04g39070;LOC_Os07g13870;LOC_Os08g09730;LOC_Os04g13040;LOC_Os03g04270;LOC_Os02g54550;LOC_Os07g36530;LOC_Os10g25660;LOC_Os03g25250;LOC_Os02g17180;LOC_Os01g05970;LOC_Os03g47420;LOC_Os08g31690;LOC_Os07g36900;LOC_Os03g07530;LOC_Os12g34210;LOC_Os12g41300;LOC_Os06g07390;LOC_Os03g30920;LOC_Os01g55430;LOC_Os11g36350;LOC_Os07g13930;LOC_Os03g46500;LOC_Os09g34200;LOC_Os12g05609;LOC_Os02g51350;LOC_Os02g03980;LOC_Os07g27030;LOC_Os03g46510;LOC_Os04g40920;LOC_Os07g16800;LOC_Os05g46300;LOC_Os06g07460;LOC_Os12g05709;LOC_Os06g02110;LOC_Os09g17152;LOC_Os05g49530;LOC_Os03g46140;LOC_Os10g05240;LOC_Os06g05620;LOC_Os10g05200;LOC_Os02g33310;LOC_Os10g04590;LOC_Os02g54240;LOC_Os04g52870;LOC_Os11g42310;LOC_Os09g15440;LOC_Os12g34290;LOC_Os05g46320;LOC_Os12g42340;LOC_Os03g51760;LOC_Os09g15570;LOC_Os05g43850;LOC_Os02g15160;LOC_Os11g38180;LOC_Os08g09590;LOC_Os02g55050;LOC_Os04g40780;LOC_Os09g38900;LOC_Os08g38330;LOC_Os11g09640;LOC_Os09g27100;LOC_Os01g60920;LOC_Os03g46530;LOC_Os05g51100;LOC_Os11g06420;LOC_Os06g07430;LOC_Os04g19810;LOC_Os12g40140;LOC_Os07g04790;LOC_Os02g06520;LOC_Os03g51270;LOC_Os07g10710;LOC_Os12g34240;LOC_Os08g40640;LOC_Os12g39520;LOC_Os11g16280;LOC_Os02g19540;LOC_Os01g40160;LOC_Os11g37300;LOC_Os02g33400;LOC_Os07g18600;LOC_Os08g38490;LOC_Os11g38100;LOC_Os08g09460;LOC_Os02g02350;LOC_Os04g57920;LOC_Os07g03090;LOC_Os11g42280;LOC_Os05g49980;LOC_Os03g49250;LOC_Os12g34200;LOC_Os10g10410;LOC_Os10g04750;LOC_Os06g47400;LOC_Os01g72310;LOC_Os06g02400;LOC_Os04g50200;LOC_Os09g39050;LOC_Os01g28520;LOC_Os10g05520;LOC_Os11g42270;LOC_Os04g40330;LOC_Os09g27660;LOC_Os10g24010;LOC_Os09g06620;LOC_Os07g03110;LOC_Os04g40800;LOC_Os03g50050;LOC_Os04g35190;LOC_Os03g40370;LOC_Os04g35940;LOC_Os02g48300;LOC_Os11g10340;LOC_Os08g06780;LOC_Os03g12200;LOC_Os04g42670;LOC_Os06g10290;LOC_Os02g38720;LOC_Os01g71450;LOC_Os10g17940;LOC_Os09g30180;LOC_Os11g33180;LOC_Os09g32410;LOC_Os01g52980;LOC_Os09g28120;LOC_Os02g28600;LOC_Os05g49540;LOC_Os03g56510;LOC_Os11g05590;LOC_Os03g56440;LOC_Os08g35960;LOC_Os05g23810;LOC_Os05g02550;LOC_Os05g01620;LOC_Os05g49680;LOC_Os08g27190;LOC_Os02g52230;LOC_Os04g31580;LOC_Os08g09420;LOC_Os03g43390;LOC_Os08g20492;LOC_Os03g24200;LOC_Os09g27570;LOC_Os04g11450;LOC_Os07g46555;LOC_Os01g05890;LOC_Os07g36870;LOC_Os11g32940;LOC_Os09g21620;LOC_Os02g21230;LOC_Os08g38470;LOC_Os02g33840;LOC_Os08g35930;LOC_Os08g24190;LOC_Os07g02910;LOC_Os03g56450;LOC_Os08g33010;LOC_Os10g04600;LOC_Os03g25220;LOC_Os02g21240;LOC_Os02g38589;LOC_Os07g36280;LOC_Os12g38090;LOC_Os02g38499;LOC_Os10g04700;LOC_Os08g35900;LOC_Os03g43060;LOC_Os07g06730 |
| F-box | sperm | 73/2975 | 2.30365E-07 | 1.20942E-06 | 2.012358624 | 30.75608434 | LOC_Os09g34200;LOC_Os10g05520;LOC_Os10g35920;LOC_Os09g06620;LOC_Os02g51350;LOC_Os07g27030;LOC_Os03g46510;LOC_Os05g46300;LOC_Os04g11890;LOC_Os01g45020;LOC_Os04g35940;LOC_Os11g10340;LOC_Os06g02110;LOC_Os06g10290;LOC_Os04g39080;LOC_Os06g05620;LOC_Os10g05200;LOC_Os02g33310;LOC_Os09g30180;LOC_Os10g03600;LOC_Os06g05580;LOC_Os09g32410;LOC_Os10g04590;LOC_Os02g36520;LOC_Os01g64030;LOC_Os03g27250;LOC_Os03g51760;LOC_Os10g04750;LOC_Os02g15160;LOC_Os10g41650;LOC_Os01g46500;LOC_Os01g28150;LOC_Os02g55050;LOC_Os02g21110;LOC_Os10g03620;LOC_Os09g38900;LOC_Os04g40030;LOC_Os11g09640;LOC_Os10g25680;LOC_Os10g10390;LOC_Os10g05800;LOC_Os01g60920;LOC_Os01g36940;LOC_Os08g20492;LOC_Os11g06420;LOC_Os08g36960;LOC_Os09g27570;LOC_Os11g07970;LOC_Os07g46555;LOC_Os06g49530;LOC_Os10g24900;LOC_Os07g36870;LOC_Os02g06520;LOC_Os03g51270;LOC_Os11g32940;LOC_Os03g04270;LOC_Os11g16280;LOC_Os08g35930;LOC_Os08g24190;LOC_Os07g02910;LOC_Os02g33400;LOC_Os02g56760;LOC_Os02g56820;LOC_Os10g04600;LOC_Os01g05970;LOC_Os02g21240;LOC_Os08g31690;LOC_Os02g02350;LOC_Os01g55430;LOC_Os03g30920;LOC_Os10g10410;LOC_Os06g02400;LOC_Os07g13930 |
| RING | sperm | 45/2975 | 0.000229143 | 0.001527619 | 1.854966356 | 15.54677952 | LOC_Os05g43610;LOC_Os11g05300;LOC_Os09g32690;LOC_Os02g03620;LOC_Os06g17280;LOC_Os09g25260;LOC_Os01g42500;LOC_Os08g06510;LOC_Os03g16480;LOC_Os03g47500;LOC_Os01g49770;LOC_Os07g45350;LOC_Os03g58390;LOC_Os12g01190;LOC_Os01g72480;LOC_Os10g20600;LOC_Os02g13810;LOC_Os03g44810;LOC_Os02g03760;LOC_Os01g03100;LOC_Os03g59540;LOC_Os07g49030;LOC_Os09g33740;LOC_Os01g57110;LOC_Os12g05370;LOC_Os07g37140;LOC_Os01g68060;LOC_Os03g22830;LOC_Os08g29590;LOC_Os07g31850;LOC_Os09g25220;LOC_Os05g19970;LOC_Os08g05560;LOC_Os11g36970;LOC_Os04g43220;LOC_Os04g53720;LOC_Os04g46450;LOC_Os05g07070;LOC_Os08g31930;LOC_Os02g47870;LOC_Os07g32730;LOC_Os03g15730;LOC_Os08g08220;LOC_Os09g15430;LOC_Os05g48970 |
| F-box | anther developmental series | 304/15916 | 1.30307E-14 | 1.36823E-13 | 1.808747172 | 57.82829773 | LOC_Os08g34860;LOC_Os03g12940;LOC_Os04g40770;LOC_Os04g35930;LOC_Os12g37130;LOC_Os06g13850;LOC_Os07g36360;LOC_Os08g35870;LOC_Os04g11890;LOC_Os01g45020;LOC_Os12g27810;LOC_Os01g27680;LOC_Os01g33490;LOC_Os10g06700;LOC_Os04g13010;LOC_Os02g41930;LOC_Os10g03600;LOC_Os01g71440;LOC_Os12g27760;LOC_Os06g43740;LOC_Os02g05700;LOC_Os01g64030;LOC_Os12g41630;LOC_Os03g27250;LOC_Os06g35080;LOC_Os10g10420;LOC_Os10g41829;LOC_Os01g06360;LOC_Os10g41650;LOC_Os01g46500;LOC_Os02g16760;LOC_Os04g19750;LOC_Os02g21110;LOC_Os03g36439;LOC_Os07g09110;LOC_Os10g25680;LOC_Os11g38130;LOC_Os08g16710;LOC_Os10g10390;LOC_Os11g39609;LOC_Os08g24370;LOC_Os01g36940;LOC_Os08g35880;LOC_Os03g12190;LOC_Os10g04370;LOC_Os05g45040;LOC_Os09g21580;LOC_Os03g25240;LOC_Os12g31340;LOC_Os05g27550;LOC_Os03g06100;LOC_Os09g39000;LOC_Os02g56760;LOC_Os02g56820;LOC_Os06g04980;LOC_Os07g48940;LOC_Os05g46050;LOC_Os09g20650;LOC_Os05g25580;LOC_Os01g17390;LOC_Os04g31540;LOC_Os06g07380;LOC_Os05g46060;LOC_Os05g49400;LOC_Os08g38520;LOC_Os09g15460;LOC_Os02g04000;LOC_Os10g35920;LOC_Os11g42240;LOC_Os04g13170;LOC_Os05g30920;LOC_Os09g32870;LOC_Os06g41520;LOC_Os12g30990;LOC_Os10g37570;LOC_Os12g27790;LOC_Os02g03910;LOC_Os07g36370;LOC_Os07g36520;LOC_Os03g25650;LOC_Os11g33220;LOC_Os04g39080;LOC_Os01g07160;LOC_Os10g05000;LOC_Os08g06710;LOC_Os09g22460;LOC_Os06g05580;LOC_Os02g35530;LOC_Os11g41570;LOC_Os01g71430;LOC_Os11g03680;LOC_Os04g40760;LOC_Os02g36520;LOC_Os01g28520;LOC_Os03g46120;LOC_Os05g08350;LOC_Os04g57290;LOC_Os10g17930;LOC_Os03g25190;LOC_Os01g56750;LOC_Os01g28150;LOC_Os04g19800;LOC_Os10g03620;LOC_Os04g40030;LOC_Os11g42160;LOC_Os09g17190;LOC_Os07g31680;LOC_Os08g24830;LOC_Os10g34340;LOC_Os08g24140;LOC_Os04g40960;LOC_Os10g05800;LOC_Os07g08570;LOC_Os11g41560;LOC_Os12g30950;LOC_Os05g02570;LOC_Os08g36960;LOC_Os07g36330;LOC_Os11g07970;LOC_Os04g26240;LOC_Os02g17210;LOC_Os06g49530;LOC_Os10g24900;LOC_Os04g39070;LOC_Os07g13870;LOC_Os08g09730;LOC_Os04g13040;LOC_Os03g04270;LOC_Os02g54550;LOC_Os07g36530;LOC_Os10g25660;LOC_Os03g25250;LOC_Os02g17180;LOC_Os01g05970;LOC_Os03g47420;LOC_Os08g31690;LOC_Os12g40370;LOC_Os07g36900;LOC_Os03g07530;LOC_Os12g34210;LOC_Os12g41300;LOC_Os06g07390;LOC_Os03g30920;LOC_Os01g55430;LOC_Os11g36350;LOC_Os07g13930;LOC_Os12g40360;LOC_Os03g46500;LOC_Os09g34200;LOC_Os12g05609;LOC_Os02g51350;LOC_Os02g03980;LOC_Os07g27030;LOC_Os03g46510;LOC_Os04g40920;LOC_Os07g16800;LOC_Os05g46300;LOC_Os06g07460;LOC_Os12g05709;LOC_Os06g02110;LOC_Os09g17152;LOC_Os05g49530;LOC_Os03g46140;LOC_Os10g05240;LOC_Os06g05620;LOC_Os10g05200;LOC_Os02g33310;LOC_Os10g04590;LOC_Os02g54240;LOC_Os04g52870;LOC_Os11g42310;LOC_Os09g15440;LOC_Os12g34290;LOC_Os05g46320;LOC_Os05g40500;LOC_Os12g42340;LOC_Os06g44920;LOC_Os03g51760;LOC_Os09g15570;LOC_Os05g43850;LOC_Os02g15160;LOC_Os11g38180;LOC_Os08g09590;LOC_Os02g55050;LOC_Os04g40780;LOC_Os09g38900;LOC_Os08g38330;LOC_Os11g09640;LOC_Os09g27100;LOC_Os01g60920;LOC_Os03g46530;LOC_Os05g51100;LOC_Os11g06420;LOC_Os06g07430;LOC_Os04g19810;LOC_Os12g40140;LOC_Os06g13990;LOC_Os07g04790;LOC_Os02g06520;LOC_Os03g51270;LOC_Os07g10710;LOC_Os03g22990;LOC_Os12g34240;LOC_Os08g40640;LOC_Os12g39520;LOC_Os11g16280;LOC_Os02g19540;LOC_Os01g40160;LOC_Os12g40320;LOC_Os11g37300;LOC_Os02g33400;LOC_Os07g18600;LOC_Os08g38490;LOC_Os11g38100;LOC_Os08g09460;LOC_Os02g02350;LOC_Os04g57920;LOC_Os07g03090;LOC_Os11g42280;LOC_Os05g49980;LOC_Os03g49250;LOC_Os01g55210;LOC_Os10g10410;LOC_Os10g04750;LOC_Os06g47400;LOC_Os12g34200;LOC_Os01g72310;LOC_Os04g50200;LOC_Os09g39050;LOC_Os06g02400;LOC_Os10g05520;LOC_Os11g42270;LOC_Os01g71460;LOC_Os06g45460;LOC_Os04g40330;LOC_Os09g27660;LOC_Os10g24010;LOC_Os09g06620;LOC_Os07g03110;LOC_Os04g40800;LOC_Os03g50050;LOC_Os04g35190;LOC_Os03g40370;LOC_Os04g35940;LOC_Os02g48300;LOC_Os11g10340;LOC_Os08g06780;LOC_Os03g12200;LOC_Os04g42670;LOC_Os06g10290;LOC_Os02g38720;LOC_Os01g71450;LOC_Os10g17940;LOC_Os09g30180;LOC_Os11g33180;LOC_Os09g32410;LOC_Os01g52980;LOC_Os09g28120;LOC_Os02g28600;LOC_Os05g49540;LOC_Os03g56510;LOC_Os11g05590;LOC_Os03g56440;LOC_Os08g35960;LOC_Os05g23810;LOC_Os05g02550;LOC_Os05g01620;LOC_Os05g49680;LOC_Os08g27190;LOC_Os02g52230;LOC_Os04g31580;LOC_Os08g09420;LOC_Os03g43390;LOC_Os08g20492;LOC_Os03g24200;LOC_Os05g40520;LOC_Os09g27570;LOC_Os04g11450;LOC_Os07g46555;LOC_Os01g05890;LOC_Os07g36870;LOC_Os12g40310;LOC_Os11g32940;LOC_Os09g21620;LOC_Os02g21230;LOC_Os08g38470;LOC_Os02g33840;LOC_Os08g35930;LOC_Os08g24190;LOC_Os07g02910;LOC_Os03g56450;LOC_Os08g33010;LOC_Os10g04600;LOC_Os03g25220;LOC_Os02g21240;LOC_Os02g38589;LOC_Os07g36280;LOC_Os12g38090;LOC_Os05g46160;LOC_Os02g38499;LOC_Os10g04700;LOC_Os08g35900;LOC_Os03g43060;LOC_Os07g06730 |
| RING | male gametophytic lineage | 100/7157 | 8.03323E-07 | 8.03323E-06 | 1.792052063 | 25.15057165 | LOC_Os09g32690;LOC_Os09g24650;LOC_Os02g03620;LOC_Os09g25260;LOC_Os08g06510;LOC_Os03g47500;LOC_Os03g08000;LOC_Os05g39380;LOC_Os12g01190;LOC_Os04g43300;LOC_Os06g05200;LOC_Os07g27950;LOC_Os01g03100;LOC_Os03g59540;LOC_Os07g43740;LOC_Os12g43930;LOC_Os01g68060;LOC_Os07g31850;LOC_Os03g16570;LOC_Os04g46450;LOC_Os04g41470;LOC_Os06g21390;LOC_Os07g32730;LOC_Os08g08220;LOC_Os03g10890;LOC_Os08g06090;LOC_Os05g48970;LOC_Os03g01790;LOC_Os06g17280;LOC_Os01g42500;LOC_Os05g11720;LOC_Os03g16480;LOC_Os01g72480;LOC_Os01g13370;LOC_Os10g20600;LOC_Os02g13810;LOC_Os09g33740;LOC_Os07g49030;LOC_Os07g31650;LOC_Os03g13100;LOC_Os07g37140;LOC_Os08g29590;LOC_Os01g08340;LOC_Os01g53130;LOC_Os11g36970;LOC_Os04g43220;LOC_Os04g53720;LOC_Os05g25180;LOC_Os04g44820;LOC_Os07g26490;LOC_Os01g61420;LOC_Os08g31930;LOC_Os05g43610;LOC_Os12g35320;LOC_Os06g12560;LOC_Os01g49770;LOC_Os02g09820;LOC_Os04g10680;LOC_Os03g58390;LOC_Os12g04660;LOC_Os02g49550;LOC_Os02g08200;LOC_Os09g33670;LOC_Os02g03760;LOC_Os05g06270;LOC_Os12g05370;LOC_Os03g22830;LOC_Os05g19970;LOC_Os08g05560;LOC_Os09g26400;LOC_Os05g07140;LOC_Os03g15730;LOC_Os03g57500;LOC_Os01g72310;LOC_Os12g24530;LOC_Os11g05300;LOC_Os05g14860;LOC_Os06g34360;LOC_Os07g45350;LOC_Os01g58780;LOC_Os08g38060;LOC_Os10g26610;LOC_Os01g16950;LOC_Os10g31850;LOC_Os03g44810;LOC_Os03g53080;LOC_Os02g36300;LOC_Os02g40810;LOC_Os02g36740;LOC_Os01g57110;LOC_Os01g68900;LOC_Os10g34030;LOC_Os09g25220;LOC_Os12g16690;LOC_Os05g07070;LOC_Os02g47870;LOC_Os10g35190;LOC_Os06g42700;LOC_Os09g15430;LOC_Os01g01420 |
| RING | mature anther | 165/12828 | 1.39148E-08 | 2.78296E-07 | 1.750289324 | 31.66328321 | LOC_Os09g32690;LOC_Os09g24650;LOC_Os02g03620;LOC_Os09g25260;LOC_Os08g06510;LOC_Os03g47500;LOC_Os03g08000;LOC_Os05g39380;LOC_Os12g01190;LOC_Os12g05280;LOC_Os05g39260;LOC_Os04g43300;LOC_Os06g05200;LOC_Os02g45780;LOC_Os01g38710;LOC_Os07g27950;LOC_Os01g03100;LOC_Os03g59540;LOC_Os07g43740;LOC_Os12g43930;LOC_Os01g68060;LOC_Os07g31850;LOC_Os03g26370;LOC_Os03g16570;LOC_Os06g34640;LOC_Os04g46450;LOC_Os06g14640;LOC_Os04g41470;LOC_Os06g21390;LOC_Os07g43380;LOC_Os07g32730;LOC_Os06g48040;LOC_Os01g11490;LOC_Os11g05200;LOC_Os03g10890;LOC_Os08g08220;LOC_Os08g06090;LOC_Os05g48970;LOC_Os08g42640;LOC_Os11g36560;LOC_Os03g01790;LOC_Os09g38640;LOC_Os06g17280;LOC_Os01g42500;LOC_Os05g11720;LOC_Os03g16480;LOC_Os03g04890;LOC_Os01g69040;LOC_Os01g72480;LOC_Os01g13370;LOC_Os05g39940;LOC_Os10g20600;LOC_Os02g13810;LOC_Os03g44636;LOC_Os06g16940;LOC_Os09g33740;LOC_Os07g49030;LOC_Os07g31650;LOC_Os01g38700;LOC_Os03g13100;LOC_Os01g35100;LOC_Os07g37140;LOC_Os08g29590;LOC_Os01g50750;LOC_Os01g08340;LOC_Os08g37760;LOC_Os12g02220;LOC_Os10g32740;LOC_Os01g38760;LOC_Os01g53130;LOC_Os11g36970;LOC_Os04g43220;LOC_Os04g53720;LOC_Os02g46740;LOC_Os04g44820;LOC_Os02g31150;LOC_Os07g26490;LOC_Os05g25180;LOC_Os01g61420;LOC_Os04g49550;LOC_Os08g31930;LOC_Os03g22110;LOC_Os08g33860;LOC_Os06g34400;LOC_Os11g04280;LOC_Os05g43610;LOC_Os12g35320;LOC_Os06g12560;LOC_Os06g09310;LOC_Os01g49770;LOC_Os02g09820;LOC_Os04g10680;LOC_Os03g58390;LOC_Os12g04660;LOC_Os02g49550;LOC_Os02g08200;LOC_Os08g44950;LOC_Os11g19500;LOC_Os09g33670;LOC_Os06g47270;LOC_Os11g04690;LOC_Os01g48310;LOC_Os02g03760;LOC_Os05g06270;LOC_Os02g44700;LOC_Os05g11860;LOC_Os12g05370;LOC_Os03g22830;LOC_Os06g34470;LOC_Os05g19970;LOC_Os02g54830;LOC_Os09g38630;LOC_Os08g05560;LOC_Os11g04680;LOC_Os03g15000;LOC_Os02g46100;LOC_Os01g35120;LOC_Os09g26400;LOC_Os02g52210;LOC_Os05g07140;LOC_Os03g15730;LOC_Os03g57500;LOC_Os01g72310;LOC_Os12g24530;LOC_Os04g41070;LOC_Os11g02260;LOC_Os11g05300;LOC_Os01g06590;LOC_Os05g14860;LOC_Os01g38720;LOC_Os06g14650;LOC_Os07g45350;LOC_Os01g58780;LOC_Os06g34360;LOC_Os08g38060;LOC_Os01g47740;LOC_Os10g26610;LOC_Os04g31390;LOC_Os01g16950;LOC_Os10g31850;LOC_Os03g44810;LOC_Os03g53080;LOC_Os10g42390;LOC_Os02g36300;LOC_Os02g40810;LOC_Os03g19020;LOC_Os02g36740;LOC_Os01g57110;LOC_Os01g68900;LOC_Os07g43830;LOC_Os10g32750;LOC_Os10g34030;LOC_Os09g25220;LOC_Os11g04281;LOC_Os12g16690;LOC_Os01g38690;LOC_Os05g07070;LOC_Os02g47870;LOC_Os05g28730;LOC_Os10g35190;LOC_Os03g22080;LOC_Os06g42700;LOC_Os09g15430;LOC_Os01g01420;LOC_Os08g31720 |
| F-box | female gametophytic lineage | 47/2205 | 0.000821784 | 0.002876245 | 1.706135807 | 12.12044444 | LOC_Os09g32600;LOC_Os12g43770;LOC_Os02g09580;LOC_Os11g09670;LOC_Os12g30940;LOC_Os04g11660;LOC_Os11g09580;LOC_Os01g59910;LOC_Os07g36910;LOC_Os06g06600;LOC_Os07g36840;LOC_Os11g09630;LOC_Os08g09480;LOC_Os11g10200;LOC_Os10g03740;LOC_Os04g36020;LOC_Os07g02770;LOC_Os08g09650;LOC_Os07g04750;LOC_Os09g27090;LOC_Os03g02210;LOC_Os08g09220;LOC_Os08g09720;LOC_Os10g37540;LOC_Os01g05880;LOC_Os09g39190;LOC_Os10g03750;LOC_Os04g36000;LOC_Os10g03730;LOC_Os11g37340;LOC_Os06g02100;LOC_Os07g13900;LOC_Os07g02290;LOC_Os08g09380;LOC_Os11g16330;LOC_Os07g36830;LOC_Os03g25640;LOC_Os12g03440;LOC_Os01g08830;LOC_Os07g36300;LOC_Os08g03300;LOC_Os04g35990;LOC_Os11g09600;LOC_Os01g56740;LOC_Os11g12480;LOC_Os10g04850;LOC_Os09g37570 |
| F-box | inflorescence & floral sporophyte | 318/17563 | 2.9317E-12 | 2.05219E-11 | 1.697741038 | 45.08425925 | LOC_Os08g34860;LOC_Os03g12940;LOC_Os04g40770;LOC_Os12g37130;LOC_Os04g11890;LOC_Os01g06360;LOC_Os07g09110;LOC_Os10g25680;LOC_Os03g25240;LOC_Os05g27550;LOC_Os09g39000;LOC_Os02g57890;LOC_Os09g20650;LOC_Os01g17390;LOC_Os05g49400;LOC_Os08g38520;LOC_Os11g36610;LOC_Os09g32870;LOC_Os10g37570;LOC_Os12g27790;LOC_Os08g06710;LOC_Os09g22460;LOC_Os03g46120;LOC_Os03g25190;LOC_Os01g56750;LOC_Os10g03620;LOC_Os11g42160;LOC_Os09g17190;LOC_Os07g08570;LOC_Os11g41560;LOC_Os07g36330;LOC_Os02g17210;LOC_Os10g25660;LOC_Os12g34210;LOC_Os12g41300;LOC_Os06g07390;LOC_Os03g30920;LOC_Os09g34200;LOC_Os07g27030;LOC_Os04g40920;LOC_Os12g05709;LOC_Os06g02110;LOC_Os06g05620;LOC_Os09g15440;LOC_Os12g34290;LOC_Os05g46320;LOC_Os05g40500;LOC_Os09g15570;LOC_Os02g15160;LOC_Os08g09590;LOC_Os09g38900;LOC_Os08g38330;LOC_Os09g27100;LOC_Os06g07430;LOC_Os12g40140;LOC_Os07g04790;LOC_Os07g10710;LOC_Os08g34820;LOC_Os12g34240;LOC_Os08g40640;LOC_Os02g19540;LOC_Os11g37300;LOC_Os11g38100;LOC_Os04g57920;LOC_Os10g10410;LOC_Os10g04750;LOC_Os06g47400;LOC_Os04g50200;LOC_Os09g39050;LOC_Os10g05520;LOC_Os01g71460;LOC_Os06g45460;LOC_Os02g03660;LOC_Os09g06620;LOC_Os04g40800;LOC_Os11g33210;LOC_Os09g30180;LOC_Os01g52980;LOC_Os05g23810;LOC_Os05g01620;LOC_Os05g49680;LOC_Os03g43390;LOC_Os08g20492;LOC_Os04g11450;LOC_Os01g05890;LOC_Os07g36870;LOC_Os08g38470;LOC_Os08g35930;LOC_Os07g02910;LOC_Os10g04600;LOC_Os07g36280;LOC_Os07g06730;LOC_Os06g13850;LOC_Os01g45020;LOC_Os02g41930;LOC_Os06g43740;LOC_Os08g24370;LOC_Os01g36940;LOC_Os08g35880;LOC_Os09g21580;LOC_Os12g31340;LOC_Os03g06100;LOC_Os02g56760;LOC_Os06g04980;LOC_Os09g15460;LOC_Os10g35920;LOC_Os05g30920;LOC_Os07g36520;LOC_Os04g39080;LOC_Os02g35530;LOC_Os05g08350;LOC_Os04g57290;LOC_Os10g17930;LOC_Os01g28150;LOC_Os07g31680;LOC_Os08g24830;LOC_Os04g40960;LOC_Os12g30950;LOC_Os11g07970;LOC_Os07g13870;LOC_Os04g13040;LOC_Os03g25250;LOC_Os03g47420;LOC_Os08g31690;LOC_Os12g40370;LOC_Os07g36900;LOC_Os12g04130;LOC_Os12g40360;LOC_Os02g03980;LOC_Os09g17152;LOC_Os05g49530;LOC_Os10g05200;LOC_Os02g33310;LOC_Os12g42340;LOC_Os11g38180;LOC_Os05g51100;LOC_Os11g06420;LOC_Os02g06520;LOC_Os12g39520;LOC_Os02g33400;LOC_Os07g03090;LOC_Os03g49250;LOC_Os11g42270;LOC_Os07g03110;LOC_Os04g35190;LOC_Os04g35940;LOC_Os08g06780;LOC_Os03g12200;LOC_Os04g42670;LOC_Os09g32410;LOC_Os02g28600;LOC_Os03g56440;LOC_Os05g02550;LOC_Os04g30810;LOC_Os08g33010;LOC_Os12g38090;LOC_Os05g46160;LOC_Os12g27810;LOC_Os10g06700;LOC_Os04g13010;LOC_Os10g03600;LOC_Os01g71440;LOC_Os12g27760;LOC_Os02g05700;LOC_Os01g64030;LOC_Os12g41630;LOC_Os03g27250;LOC_Os10g41829;LOC_Os10g41650;LOC_Os02g21110;LOC_Os03g36439;LOC_Os11g38130;LOC_Os05g45040;LOC_Os02g56820;LOC_Os05g25580;LOC_Os04g31540;LOC_Os03g30160;LOC_Os05g46060;LOC_Os06g41520;LOC_Os02g03910;LOC_Os07g36370;LOC_Os01g07160;LOC_Os10g05000;LOC_Os11g03680;LOC_Os08g38480;LOC_Os04g40030;LOC_Os10g34340;LOC_Os05g02570;LOC_Os08g36960;LOC_Os04g26240;LOC_Os06g49530;LOC_Os10g24900;LOC_Os04g39070;LOC_Os08g09730;LOC_Os03g04270;LOC_Os02g54550;LOC_Os02g17180;LOC_Os01g55430;LOC_Os07g13930;LOC_Os12g05609;LOC_Os02g51350;LOC_Os03g46510;LOC_Os07g16800;LOC_Os05g46300;LOC_Os02g54240;LOC_Os04g52870;LOC_Os11g42310;LOC_Os06g44920;LOC_Os03g46530;LOC_Os04g19810;LOC_Os03g51270;LOC_Os08g09460;LOC_Os02g02350;LOC_Os11g42280;LOC_Os01g55210;LOC_Os12g34200;LOC_Os01g72310;LOC_Os06g02400;LOC_Os04g40330;LOC_Os10g24010;LOC_Os06g10290;LOC_Os01g71450;LOC_Os11g33180;LOC_Os09g28120;LOC_Os05g49540;LOC_Os03g56510;LOC_Os02g52230;LOC_Os03g24200;LOC_Os05g40520;LOC_Os09g27570;LOC_Os12g40310;LOC_Os11g32940;LOC_Os06g02054;LOC_Os09g21620;LOC_Os02g33840;LOC_Os08g24190;LOC_Os03g56450;LOC_Os02g38720;LOC_Os03g44980;LOC_Os04g35930;LOC_Os07g36360;LOC_Os08g35870;LOC_Os01g27680;LOC_Os01g33490;LOC_Os06g35080;LOC_Os10g10420;LOC_Os01g46500;LOC_Os04g19750;LOC_Os02g16760;LOC_Os10g10390;LOC_Os08g16710;LOC_Os11g39609;LOC_Os03g12190;LOC_Os10g04370;LOC_Os07g48940;LOC_Os07g17570;LOC_Os05g46050;LOC_Os06g07380;LOC_Os02g04000;LOC_Os11g42240;LOC_Os04g13170;LOC_Os12g30990;LOC_Os03g25650;LOC_Os11g33220;LOC_Os06g05580;LOC_Os11g41570;LOC_Os01g71430;LOC_Os04g40760;LOC_Os02g36520;LOC_Os04g19800;LOC_Os08g24140;LOC_Os10g05800;LOC_Os07g36530;LOC_Os01g05970;LOC_Os03g07530;LOC_Os11g36350;LOC_Os03g46500;LOC_Os02g03810;LOC_Os06g07460;LOC_Os03g46140;LOC_Os10g05240;LOC_Os10g04590;LOC_Os03g51760;LOC_Os05g43850;LOC_Os02g55050;LOC_Os04g40780;LOC_Os11g09640;LOC_Os01g60920;LOC_Os06g13990;LOC_Os03g22990;LOC_Os11g16280;LOC_Os01g40160;LOC_Os12g40320;LOC_Os07g18600;LOC_Os08g38490;LOC_Os05g49980;LOC_Os01g28520;LOC_Os09g27660;LOC_Os03g50050;LOC_Os03g40370;LOC_Os02g48300;LOC_Os11g10340;LOC_Os10g17940;LOC_Os03g44920;LOC_Os11g05590;LOC_Os08g35960;LOC_Os08g27190;LOC_Os04g31580;LOC_Os08g09420;LOC_Os07g46555;LOC_Os02g21230;LOC_Os03g25220;LOC_Os02g21240;LOC_Os02g38589;LOC_Os02g38499;LOC_Os10g04700;LOC_Os08g35900;LOC_Os03g43060 |
| RING | pollen | 47/3487 | 0.001898035 | 0.009490176 | 1.640873911 | 10.28325186 | LOC_Os12g24530;LOC_Os12g35320;LOC_Os05g14860;LOC_Os06g12560;LOC_Os05g11720;LOC_Os02g09820;LOC_Os04g10680;LOC_Os06g34360;LOC_Os03g08000;LOC_Os05g39380;LOC_Os08g38060;LOC_Os10g26610;LOC_Os02g49550;LOC_Os01g13370;LOC_Os01g16950;LOC_Os10g31850;LOC_Os03g53080;LOC_Os02g36300;LOC_Os02g40810;LOC_Os04g43300;LOC_Os06g05200;LOC_Os02g36740;LOC_Os07g27950;LOC_Os05g06270;LOC_Os07g43740;LOC_Os07g31650;LOC_Os12g43930;LOC_Os03g13100;LOC_Os01g68900;LOC_Os01g72310;LOC_Os10g34030;LOC_Os01g08340;LOC_Os03g16570;LOC_Os06g42700;LOC_Os01g53130;LOC_Os12g16690;LOC_Os04g44820;LOC_Os07g26490;LOC_Os01g61420;LOC_Os04g41470;LOC_Os06g21390;LOC_Os09g26400;LOC_Os05g07140;LOC_Os10g35190;LOC_Os03g10890;LOC_Os03g57500;LOC_Os08g06090 |
| F-box | male gametophytic lineage | 139/7157 | 1.02685E-06 | 4.31279E-06 | 1.617499162 | 22.30371202 | LOC_Os03g12940;LOC_Os08g35870;LOC_Os04g11890;LOC_Os01g45020;LOC_Os01g33490;LOC_Os10g03600;LOC_Os02g05700;LOC_Os01g64030;LOC_Os03g27250;LOC_Os01g46500;LOC_Os10g41650;LOC_Os02g21110;LOC_Os03g36439;LOC_Os10g25680;LOC_Os11g38130;LOC_Os08g16710;LOC_Os10g10390;LOC_Os01g36940;LOC_Os08g35880;LOC_Os05g45040;LOC_Os12g31340;LOC_Os03g06100;LOC_Os02g56760;LOC_Os02g56820;LOC_Os06g04980;LOC_Os09g20650;LOC_Os05g25580;LOC_Os01g17390;LOC_Os10g35920;LOC_Os09g32870;LOC_Os12g27790;LOC_Os04g39080;LOC_Os10g05000;LOC_Os06g05580;LOC_Os11g41570;LOC_Os02g36520;LOC_Os11g03680;LOC_Os05g08350;LOC_Os04g57290;LOC_Os10g17930;LOC_Os01g28150;LOC_Os10g03620;LOC_Os04g40030;LOC_Os11g42160;LOC_Os08g24140;LOC_Os08g24830;LOC_Os10g05800;LOC_Os11g41560;LOC_Os08g36960;LOC_Os11g07970;LOC_Os02g17210;LOC_Os06g49530;LOC_Os10g24900;LOC_Os03g04270;LOC_Os10g25660;LOC_Os01g05970;LOC_Os08g31690;LOC_Os12g34210;LOC_Os06g07390;LOC_Os01g55430;LOC_Os03g30920;LOC_Os07g13930;LOC_Os09g34200;LOC_Os12g05609;LOC_Os02g51350;LOC_Os07g27030;LOC_Os03g46510;LOC_Os04g40920;LOC_Os05g46300;LOC_Os06g02110;LOC_Os06g05620;LOC_Os10g05200;LOC_Os02g33310;LOC_Os10g04590;LOC_Os05g46320;LOC_Os03g51760;LOC_Os09g15570;LOC_Os05g43850;LOC_Os02g15160;LOC_Os08g09590;LOC_Os02g55050;LOC_Os09g38900;LOC_Os11g09640;LOC_Os01g60920;LOC_Os11g06420;LOC_Os12g40140;LOC_Os02g06520;LOC_Os03g51270;LOC_Os12g39520;LOC_Os11g16280;LOC_Os02g19540;LOC_Os11g37300;LOC_Os02g33400;LOC_Os07g18600;LOC_Os08g09460;LOC_Os02g02350;LOC_Os05g49980;LOC_Os04g57920;LOC_Os03g49250;LOC_Os01g72310;LOC_Os10g10410;LOC_Os10g04750;LOC_Os06g02400;LOC_Os10g05520;LOC_Os04g40330;LOC_Os09g27660;LOC_Os09g06620;LOC_Os07g03110;LOC_Os03g50050;LOC_Os04g35190;LOC_Os04g35940;LOC_Os11g10340;LOC_Os08g06780;LOC_Os04g42670;LOC_Os06g10290;LOC_Os01g71450;LOC_Os09g30180;LOC_Os09g32410;LOC_Os01g52980;LOC_Os09g28120;LOC_Os02g28600;LOC_Os08g35960;LOC_Os08g09420;LOC_Os03g43390;LOC_Os08g20492;LOC_Os09g27570;LOC_Os04g11450;LOC_Os07g46555;LOC_Os01g05890;LOC_Os07g36870;LOC_Os11g32940;LOC_Os08g35930;LOC_Os08g24190;LOC_Os07g02910;LOC_Os10g04600;LOC_Os03g25220;LOC_Os02g21240;LOC_Os10g04700;LOC_Os07g06730 |
| RING | anther developmental series | 166/15916 | 0.002858751 | 0.011435004 | 1.317424544 | 7.716643568 | LOC_Os09g32690;LOC_Os09g24650;LOC_Os02g03620;LOC_Os09g25260;LOC_Os08g06510;LOC_Os03g47500;LOC_Os03g08000;LOC_Os05g39380;LOC_Os12g01190;LOC_Os12g05280;LOC_Os05g39260;LOC_Os04g43300;LOC_Os06g05200;LOC_Os02g45780;LOC_Os01g38710;LOC_Os07g27950;LOC_Os01g03100;LOC_Os03g59540;LOC_Os07g43740;LOC_Os12g43930;LOC_Os01g68060;LOC_Os07g31850;LOC_Os03g26370;LOC_Os03g16570;LOC_Os06g34640;LOC_Os04g46450;LOC_Os06g14640;LOC_Os04g41470;LOC_Os06g21390;LOC_Os07g43380;LOC_Os07g32730;LOC_Os06g48040;LOC_Os01g11490;LOC_Os11g05200;LOC_Os03g10890;LOC_Os08g08220;LOC_Os08g06090;LOC_Os05g48970;LOC_Os08g42640;LOC_Os11g36560;LOC_Os03g01790;LOC_Os09g38640;LOC_Os06g17280;LOC_Os01g42500;LOC_Os05g11720;LOC_Os03g16480;LOC_Os03g04890;LOC_Os01g69040;LOC_Os01g72480;LOC_Os01g13370;LOC_Os05g39940;LOC_Os10g20600;LOC_Os02g13810;LOC_Os03g44636;LOC_Os06g16940;LOC_Os09g33740;LOC_Os07g49030;LOC_Os07g31650;LOC_Os01g38700;LOC_Os03g13100;LOC_Os01g35100;LOC_Os07g37140;LOC_Os08g29590;LOC_Os01g50750;LOC_Os01g08340;LOC_Os08g37760;LOC_Os12g02220;LOC_Os10g32740;LOC_Os01g38760;LOC_Os01g53130;LOC_Os11g36970;LOC_Os04g43220;LOC_Os04g53720;LOC_Os02g46740;LOC_Os04g44820;LOC_Os02g31150;LOC_Os07g26490;LOC_Os05g25180;LOC_Os01g61420;LOC_Os04g49550;LOC_Os08g31930;LOC_Os03g22110;LOC_Os08g33860;LOC_Os06g34400;LOC_Os11g04280;LOC_Os05g43610;LOC_Os12g35320;LOC_Os06g12560;LOC_Os06g09310;LOC_Os01g49770;LOC_Os02g09820;LOC_Os04g10680;LOC_Os03g58390;LOC_Os12g04660;LOC_Os04g41080;LOC_Os02g49550;LOC_Os02g08200;LOC_Os08g44950;LOC_Os11g19500;LOC_Os09g33670;LOC_Os06g47270;LOC_Os11g04690;LOC_Os01g48310;LOC_Os02g03760;LOC_Os05g06270;LOC_Os02g44700;LOC_Os05g11860;LOC_Os12g05370;LOC_Os03g22830;LOC_Os06g34470;LOC_Os05g19970;LOC_Os02g54830;LOC_Os09g38630;LOC_Os08g05560;LOC_Os11g04680;LOC_Os03g15000;LOC_Os02g46100;LOC_Os01g35120;LOC_Os09g26400;LOC_Os02g52210;LOC_Os05g07140;LOC_Os03g15730;LOC_Os03g57500;LOC_Os01g72310;LOC_Os12g24530;LOC_Os04g41070;LOC_Os11g02260;LOC_Os11g05300;LOC_Os01g06590;LOC_Os05g14860;LOC_Os01g38720;LOC_Os06g14650;LOC_Os07g45350;LOC_Os01g58780;LOC_Os06g34360;LOC_Os08g38060;LOC_Os01g47740;LOC_Os10g26610;LOC_Os04g31390;LOC_Os01g16950;LOC_Os10g31850;LOC_Os03g44810;LOC_Os03g53080;LOC_Os10g42390;LOC_Os02g36300;LOC_Os02g40810;LOC_Os03g19020;LOC_Os02g36740;LOC_Os01g57110;LOC_Os01g68900;LOC_Os07g43830;LOC_Os10g32750;LOC_Os10g34030;LOC_Os09g25220;LOC_Os11g04281;LOC_Os12g16690;LOC_Os01g38690;LOC_Os05g07070;LOC_Os02g47870;LOC_Os05g28730;LOC_Os10g35190;LOC_Os03g22080;LOC_Os06g42700;LOC_Os09g15430;LOC_Os01g01420;LOC_Os08g31720 |
| RING | inflorescence & floral sporophyte | 178/17563 | 0.006588407 | 0.021961357 | 1.274860708 | 6.402916126 | LOC_Os09g32690;LOC_Os02g32570;LOC_Os09g24650;LOC_Os02g03620;LOC_Os09g25260;LOC_Os08g06510;LOC_Os03g47500;LOC_Os03g08000;LOC_Os05g39380;LOC_Os12g01190;LOC_Os12g05280;LOC_Os05g39260;LOC_Os04g43300;LOC_Os06g05200;LOC_Os02g45780;LOC_Os01g38710;LOC_Os07g27950;LOC_Os01g03100;LOC_Os03g59540;LOC_Os07g43740;LOC_Os12g43930;LOC_Os01g68060;LOC_Os07g31850;LOC_Os03g26370;LOC_Os03g16570;LOC_Os06g34640;LOC_Os04g46450;LOC_Os06g14640;LOC_Os04g41470;LOC_Os06g21390;LOC_Os07g43380;LOC_Os07g32730;LOC_Os06g48040;LOC_Os01g11490;LOC_Os11g05200;LOC_Os03g10890;LOC_Os08g08220;LOC_Os08g06090;LOC_Os05g48970;LOC_Os08g42640;LOC_Os10g41580;LOC_Os11g36560;LOC_Os03g44642;LOC_Os03g01790;LOC_Os09g38640;LOC_Os06g17280;LOC_Os01g42500;LOC_Os05g11720;LOC_Os03g16480;LOC_Os03g04890;LOC_Os01g69040;LOC_Os01g72480;LOC_Os01g13370;LOC_Os05g39940;LOC_Os10g20600;LOC_Os02g13810;LOC_Os03g44636;LOC_Os06g16940;LOC_Os09g33740;LOC_Os07g49030;LOC_Os07g31650;LOC_Os01g38700;LOC_Os03g13100;LOC_Os01g35100;LOC_Os07g37140;LOC_Os08g29590;LOC_Os01g50750;LOC_Os01g08340;LOC_Os08g37760;LOC_Os12g02220;LOC_Os10g32740;LOC_Os01g38760;LOC_Os01g53130;LOC_Os11g36970;LOC_Os04g43220;LOC_Os04g53720;LOC_Os02g46740;LOC_Os04g44820;LOC_Os02g31150;LOC_Os07g26490;LOC_Os05g25180;LOC_Os01g61420;LOC_Os04g49550;LOC_Os08g31930;LOC_Os03g22110;LOC_Os08g33860;LOC_Os06g34400;LOC_Os12g04130;LOC_Os01g44240;LOC_Os11g04280;LOC_Os05g43610;LOC_Os12g35320;LOC_Os06g12560;LOC_Os06g09310;LOC_Os01g49770;LOC_Os02g09820;LOC_Os04g10680;LOC_Os03g58390;LOC_Os12g04660;LOC_Os02g35329;LOC_Os04g41080;LOC_Os02g49550;LOC_Os10g30310;LOC_Os02g08200;LOC_Os08g44950;LOC_Os11g19500;LOC_Os09g33670;LOC_Os06g47270;LOC_Os11g04690;LOC_Os01g48310;LOC_Os02g03760;LOC_Os12g10250;LOC_Os05g06270;LOC_Os12g01340;LOC_Os02g44700;LOC_Os05g11860;LOC_Os12g05370;LOC_Os03g22830;LOC_Os06g34470;LOC_Os05g19970;LOC_Os02g54830;LOC_Os09g38630;LOC_Os08g05560;LOC_Os11g04680;LOC_Os03g15000;LOC_Os02g46100;LOC_Os01g35120;LOC_Os09g26400;LOC_Os02g52210;LOC_Os05g07140;LOC_Os03g15730;LOC_Os06g44275;LOC_Os03g57500;LOC_Os01g72310;LOC_Os12g24530;LOC_Os04g41070;LOC_Os10g32980;LOC_Os11g02260;LOC_Os11g05300;LOC_Os01g06590;LOC_Os05g14860;LOC_Os01g38720;LOC_Os06g14650;LOC_Os07g45350;LOC_Os01g58780;LOC_Os06g34360;LOC_Os08g38060;LOC_Os01g47740;LOC_Os10g26610;LOC_Os04g31390;LOC_Os01g16950;LOC_Os10g31850;LOC_Os03g44810;LOC_Os03g53080;LOC_Os10g42390;LOC_Os02g36300;LOC_Os02g40810;LOC_Os03g19020;LOC_Os02g36740;LOC_Os01g57110;LOC_Os01g68900;LOC_Os07g43830;LOC_Os10g32750;LOC_Os10g34030;LOC_Os09g25220;LOC_Os11g04281;LOC_Os02g35347;LOC_Os12g16690;LOC_Os01g38690;LOC_Os05g07070;LOC_Os02g47870;LOC_Os05g28730;LOC_Os10g35190;LOC_Os03g22080;LOC_Os06g42700;LOC_Os09g15430;LOC_Os01g01420;LOC_Os08g31720 |
