## Supplementary material for "RE3DB: A multi-omics phylogenomics platform for rice E3 ubiquitin ligases identifies novel regulators of pollen germination": Table S9

| **E3 Family** | **Enriched Term** | **Symbol** | **RAPdb** | **MSU** |
| --- | --- | --- | --- | --- |
| U-box | root system | OsPUB67 | Os10g0552400 | LOC_Os10g40490 |
| U-box | root system | OsPUB1 | Os03g0427900 | LOC_Os03g31400 |
| U-box | root system | OsPUB19 | Os06g0238200 | LOC_Os06g13090 |
| U-box | root system | OsPUB20 | Os06g0238000 | LOC_Os06g13080 |
| U-box | root system | OsPUB32 | Os04g0686000 | LOC_Os04g58920 |
| U-box | root system | OsPUB34 | None | LOC_Os04g34030 |
| U-box | root system | OsPUB35 | Os04g0589700 | LOC_Os04g49970 |
| U-box | root system | OsPUB37 | Os12g0161100 | LOC_Os12g06410 |
| U-box | root system | OsPUB49 | Os10g0548850 | LOC_Os10g40120 |
| U-box | root system | OsPUB58 | Os10g0548850 | LOC_Os10g40120 |
| U-box | root system | OsPUB62 | Os02g0822900 | LOC_Os02g57700 |
| F-box | mature anther | GIC | Os04g0675800 | LOC_Os04g57920 |
| F-box | mature anther | MOF | Os04g0464966 | LOC_Os04g39080 |
| F-box | mature anther | OsAFB4 | Os02g0759700 | LOC_Os02g52230 |
| F-box | mature anther | OsDRF1 | Os04g0431200 | LOC_Os04g35190 |
| F-box | mature anther | OsFBK12 | Os03g0171600 | LOC_Os03g07530 |
| F-box | mature anther | OsLKP2 | Os02g0150800 | LOC_Os02g05700 |
| F-box | mature anther | CHR708 | Os01g0952200 | LOC_Os01g72310 |
| F-box | mature anther | OsJMJ-C8 | Os03g0389900 | LOC_Os03g27250 |
| F-box | mature anther | OsSTA12 | Os01g0281000 | LOC_Os01g17390 |
| F-box | mature anther | OsSTA237 | Os10g0183800 | LOC_Os10g10420 |
| F-box | mature anther | OsSTA265 | Os11g0593600 | LOC_Os11g38130 |
| F-box | mature anther | OsSTA64 | Os02g0543200 | LOC_Os02g33840 |
| F-box | mature anther | OsTLP10\|OsTLP10a\|OsTLP10b | Os05g0514300 | LOC_Os05g43850 |
| F-box | mature anther | OsTLP12 | Os01g0759100 | LOC_Os01g55430 |
| F-box | mature anther | OsTLP6 | Os11g0163600 | LOC_Os11g06420 |
| F-box | mature anther | OsWD40-110 | Os04g0619700 | LOC_Os04g52870 |
| F-box | mature anther | OsWD40-184 | Os11g0182400 | LOC_Os11g07970 |
| F-box | sperm | MOF | Os04g0464966 | LOC_Os04g39080 |
| F-box | sperm | OsJMJ-C8 | Os03g0389900 | LOC_Os03g27250 |
| F-box | sperm | OsTLP12 | Os01g0759100 | LOC_Os01g55430 |
| F-box | sperm | OsTLP6 | Os11g0163600 | LOC_Os11g06420 |
| F-box | sperm | OsWD40-184 | Os11g0182400 | LOC_Os11g07970 |
| RING | sperm | ALT1 | Os01g0779400 | LOC_Os01g57110 |
| RING | sperm | BRHIS1 | Os08g0180300 | LOC_Os08g08220 |
| RING | sperm | EBR1 | Os05g0279400 | LOC_Os05g19970 |
| RING | sperm | HEI10 | Os02g0232100 | LOC_Os02g13810 |
| RING | sperm | OsHTAS | Os09g0323100 | LOC_Os09g15430 |
| RING | sperm | OsHUB1\|MIP1 | Os04g0550400 | LOC_Os04g46450 |
| RING | sperm | OsNLA2 | Os03g0650900 | LOC_Os03g44810 |
| RING | sperm | OsPIE1 | Os01g0954400 | LOC_Os01g72480 |
| RING | sperm | OsRINGC2-2 | Os12g0102400 | LOC_Os12g01190 |
| RING | sperm | OsSIRP2 | Os03g0798200 | LOC_Os03g58390 |
| RING | sperm | CHR715 | Os04g0629300 | LOC_Os04g53720 |
| RING | sperm | CHR731 | Os07g0511500 | LOC_Os07g32730 |
| RING | sperm | OsRFPH2-23 | Os01g0692700 | LOC_Os01g49770 |
| RING | sperm | OsRFPHC-9 | Os02g0128800 | LOC_Os02g03620 |
| RING | sperm | OsRFPV-4 | Os01g0610700 | LOC_Os01g42500 |
| RING | sperm | OsZFP96 | Os11g0578200\|Os11g0578225\|Os11g0578250 | LOC_Os11g36970 |
| F-box | anther developmental series | APO1\|OsAPO1\|SCM2 | Os06g0665400 | LOC_Os06g45460 |
| F-box | anther developmental series | GIC | Os04g0675800 | LOC_Os04g57920 |
| F-box | anther developmental series | MOF | Os04g0464966 | LOC_Os04g39080 |
| F-box | anther developmental series | OsAFB4 | Os02g0759700 | LOC_Os02g52230 |
| F-box | anther developmental series | OsDRF1 | Os04g0431200 | LOC_Os04g35190 |
| F-box | anther developmental series | OsFBK12 | Os03g0171600 | LOC_Os03g07530 |
| F-box | anther developmental series | OsLKP2 | Os02g0150800 | LOC_Os02g05700 |
| F-box | anther developmental series | CHR708 | Os01g0952200 | LOC_Os01g72310 |
| F-box | anther developmental series | OsJMJ-C8 | Os03g0389900 | LOC_Os03g27250 |
| F-box | anther developmental series | OsSTA12 | Os01g0281000 | LOC_Os01g17390 |
| F-box | anther developmental series | OsSTA237 | Os10g0183800 | LOC_Os10g10420 |
| F-box | anther developmental series | OsSTA265 | Os11g0593600 | LOC_Os11g38130 |
| F-box | anther developmental series | OsSTA64 | Os02g0543200 | LOC_Os02g33840 |
| F-box | anther developmental series | OsTLP10\|OsTLP10a\|OsTLP10b | Os05g0514300 | LOC_Os05g43850 |
| F-box | anther developmental series | OsTLP12 | Os01g0759100 | LOC_Os01g55430 |
| F-box | anther developmental series | OsTLP6 | Os11g0163600 | LOC_Os11g06420 |
| F-box | anther developmental series | OsWD40-110 | Os04g0619700 | LOC_Os04g52870 |
| F-box | anther developmental series | OsWD40-184 | Os11g0182400 | LOC_Os11g07970 |
| RING | male gametophytic lineage | ALT1 | Os01g0779400 | LOC_Os01g57110 |
| RING | male gametophytic lineage | APIP6 | Os05g0154600 | LOC_Os05g06270 |
| RING | male gametophytic lineage | BRHIS1 | Os08g0180300 | LOC_Os08g08220 |
| RING | male gametophytic lineage | DCA1 | Os10g0456800 | LOC_Os10g31850 |
| RING | male gametophytic lineage | EBR1 | Os05g0279400 | LOC_Os05g19970 |
| RING | male gametophytic lineage | HEI10 | Os02g0232100 | LOC_Os02g13810 |
| RING | male gametophytic lineage | MKRN | Os06g0318700 | LOC_Os06g21390 |
| RING | male gametophytic lineage | OsCLR1\|DSNP1 | Os06g0633500 | LOC_Os06g42700 |
| RING | male gametophytic lineage | OsCOIN | Os01g0104100 | LOC_Os01g01420 |
| RING | male gametophytic lineage | OsDSG1 | Os09g0434200 | LOC_Os09g26400 |
| RING | male gametophytic lineage | OsHTAS | Os09g0323100 | LOC_Os09g15430 |
| RING | male gametophytic lineage | OsHUB1\|MIP1 | Os04g0550400 | LOC_Os04g46450 |
| RING | male gametophytic lineage | OsNLA2 | Os03g0650900 | LOC_Os03g44810 |
| RING | male gametophytic lineage | OsPIE1 | Os01g0954400 | LOC_Os01g72480 |
| RING | male gametophytic lineage | OsRDCP1\|OsREIW1 | Os04g0530500 | LOC_Os04g44820 |
| RING | male gametophytic lineage | OsRINGC2-2 | Os12g0102400 | LOC_Os12g01190 |
| RING | male gametophytic lineage | OsSDIR1 | Os03g0272300 | LOC_Os03g16570 |
| RING | male gametophytic lineage | OsSIRP2 | Os03g0798200 | LOC_Os03g58390 |
| RING | male gametophytic lineage | SDEL1\|OsMSRFP | Os12g0538500 | LOC_Os12g35320 |
| RING | male gametophytic lineage | CHR708 | Os01g0952200 | LOC_Os01g72310 |
| RING | male gametophytic lineage | CHR715 | Os04g0629300 | LOC_Os04g53720 |
| RING | male gametophytic lineage | CHR731 | Os07g0511500 | LOC_Os07g32730 |
| RING | male gametophytic lineage | OsRFPH2-11 | Os10g0406200 | LOC_Os10g26610 |
| RING | male gametophytic lineage | OsRFPH2-12 | Os01g0802000 | LOC_Os01g58780 |
| RING | male gametophytic lineage | OsRFPH2-19 | Os03g0788800 | LOC_Os03g57500 |
| RING | male gametophytic lineage | OsRFPH2-23 | Os01g0692700 | LOC_Os01g49770 |
| RING | male gametophytic lineage | OsRFPH2-7 | Os06g0231600 | LOC_Os06g12560 |
| RING | male gametophytic lineage | OsRFPHC-12 | Os04g0512400 | LOC_Os04g43300 |
| RING | male gametophytic lineage | OsRFPHC-14 | Os01g0234900 | LOC_Os01g13370 |
| RING | male gametophytic lineage | OsRFPHC-9 | Os02g0128800 | LOC_Os02g03620 |
| RING | male gametophytic lineage | OsRFPV-4 | Os01g0610700 | LOC_Os01g42500 |
| RING | male gametophytic lineage | OsSTA11 | Os01g0276600 | LOC_Os01g16950 |
| RING | male gametophytic lineage | OsSTA5 | Os01g0178700 | LOC_Os01g08340 |
| RING | male gametophytic lineage | OsSTA9 | Os01g0234900 | LOC_Os01g13370 |
| RING | male gametophytic lineage | OsZFP96 | Os11g0578200\|Os11g0578225\|Os11g0578250 | LOC_Os11g36970 |
| RING | male gametophytic lineage | XBOS34 | Os07g0446100 | LOC_Os07g26490 |
| RING | male gametophytic lineage | ZOS10-06 | Os10g0494500 | LOC_Os10g35190 |
| RING | mature anther | ALT1 | Os01g0779400 | LOC_Os01g57110 |
| RING | mature anther | APIP6 | Os05g0154600 | LOC_Os05g06270 |
| RING | mature anther | BRHIS1 | Os08g0180300 | LOC_Os08g08220 |
| RING | mature anther | DCA1 | Os10g0456800 | LOC_Os10g31850 |
| RING | mature anther | DHS | Os02g0682300 | LOC_Os02g45780 |
| RING | mature anther | EBR1 | Os05g0279400 | LOC_Os05g19970 |
| RING | mature anther | HEI10 | Os02g0232100 | LOC_Os02g13810 |
| RING | mature anther | MKRN | Os06g0318700 | LOC_Os06g21390 |
| RING | mature anther | OsCLR1\|DSNP1 | Os06g0633500 | LOC_Os06g42700 |
| RING | mature anther | OsCOIN | Os01g0104100 | LOC_Os01g01420 |
| RING | mature anther | OsDIRP1 | Os06g0687200 | LOC_Os06g47270 |
| RING | mature anther | OsDSG1 | Os09g0434200 | LOC_Os09g26400 |
| RING | mature anther | OsHIRP1 | Os03g0302200 | LOC_Os03g19020 |
| RING | mature anther | OsHTAS | Os09g0323100 | LOC_Os09g15430 |
| RING | mature anther | OsHUB1\|MIP1 | Os04g0550400 | LOC_Os04g46450 |
| RING | mature anther | OsMAR1 | Os06g0695600 | LOC_Os06g48040 |
| RING | mature anther | OsNLA2 | Os03g0650900 | LOC_Os03g44810 |
| RING | mature anther | OsPIE1 | Os01g0954400 | LOC_Os01g72480 |
| RING | mature anther | OsRDCP1\|OsREIW1 | Os04g0530500 | LOC_Os04g44820 |
| RING | mature anther | OsRFPH2-10 | Os08g0539300 | LOC_Os08g42640 |
| RING | mature anther | OsRING-1\|OsATL38 | Os02g0759400 | LOC_Os02g52210 |
| RING | mature anther | OsRINGC2-2 | Os12g0102400 | LOC_Os12g01190 |
| RING | mature anther | OsSDIR1 | Os03g0272300 | LOC_Os03g16570 |
| RING | mature anther | OsSIRP2 | Os03g0798200 | LOC_Os03g58390 |
| RING | mature anther | SDEL1\|OsMSRFP | Os12g0538500 | LOC_Os12g35320 |
| RING | mature anther | UCIP2 | Os01g0667700 | LOC_Os01g47740 |
| RING | mature anther | AM29 | Os06g0535900 | LOC_Os06g34470 |
| RING | mature anther | CHR708 | Os01g0952200 | LOC_Os01g72310 |
| RING | mature anther | CHR715 | Os04g0629300 | LOC_Os04g53720 |
| RING | mature anther | CHR731 | Os07g0511500 | LOC_Os07g32730 |
| RING | mature anther | OsRFPH2-11 | Os10g0406200 | LOC_Os10g26610 |
| RING | mature anther | OsRFPH2-12 | Os01g0802000 | LOC_Os01g58780 |
| RING | mature anther | OsRFPH2-19 | Os03g0788800 | LOC_Os03g57500 |
| RING | mature anther | OsRFPH2-23 | Os01g0692700 | LOC_Os01g49770 |
| RING | mature anther | OsRFPH2-7 | Os06g0231600 | LOC_Os06g12560 |
| RING | mature anther | OsRFPH2-9 | Os03g0142500 | LOC_Os03g04890 |
| RING | mature anther | OsRFPHC-12 | Os04g0512400 | LOC_Os04g43300 |
| RING | mature anther | OsRFPHC-14 | Os01g0234900 | LOC_Os01g13370 |
| RING | mature anther | OsRFPHC-6 | Os07g0626900 | LOC_Os07g43380 |
| RING | mature anther | OsRFPHC-9 | Os02g0128800 | LOC_Os02g03620 |
| RING | mature anther | OsRFPV-2 | Os05g0355300 | LOC_Os05g28730 |
| RING | mature anther | OsRFPV-4 | Os01g0610700 | LOC_Os01g42500 |
| RING | mature anther | OsSTA11 | Os01g0276600 | LOC_Os01g16950 |
| RING | mature anther | OsSTA5 | Os01g0178700 | LOC_Os01g08340 |
| RING | mature anther | OsSTA9 | Os01g0234900 | LOC_Os01g13370 |
| RING | mature anther | OsZFP96 | Os11g0578200\|Os11g0578225\|Os11g0578250 | LOC_Os11g36970 |
| RING | mature anther | XBOS34 | Os07g0446100 | LOC_Os07g26490 |
| RING | mature anther | ZOS10-06 | Os10g0494500 | LOC_Os10g35190 |
| F-box | female gametophytic lineage | OsFBDUF48 | Os10g0519800 | LOC_Os10g37540 |
| F-box | inflorescence & floral sporophyte | APO1\|OsAPO1\|SCM2 | Os06g0665400 | LOC_Os06g45460 |
| F-box | inflorescence & floral sporophyte | GIC | Os04g0675800 | LOC_Os04g57920 |
| F-box | inflorescence & floral sporophyte | MOF | Os04g0464966 | LOC_Os04g39080 |
| F-box | inflorescence & floral sporophyte | OsAFB4 | Os02g0759700 | LOC_Os02g52230 |
| F-box | inflorescence & floral sporophyte | OsDRF1 | Os04g0431200 | LOC_Os04g35190 |
| F-box | inflorescence & floral sporophyte | OsFBK12 | Os03g0171600 | LOC_Os03g07530 |
| F-box | inflorescence & floral sporophyte | OsLKP2 | Os02g0150800 | LOC_Os02g05700 |
| F-box | inflorescence & floral sporophyte | CHR708 | Os01g0952200 | LOC_Os01g72310 |
| F-box | inflorescence & floral sporophyte | OsJMJ-C8 | Os03g0389900 | LOC_Os03g27250 |
| F-box | inflorescence & floral sporophyte | OsSTA12 | Os01g0281000 | LOC_Os01g17390 |
| F-box | inflorescence & floral sporophyte | OsSTA237 | Os10g0183800 | LOC_Os10g10420 |
| F-box | inflorescence & floral sporophyte | OsSTA265 | Os11g0593600 | LOC_Os11g38130 |
| F-box | inflorescence & floral sporophyte | OsSTA64 | Os02g0543200 | LOC_Os02g33840 |
| F-box | inflorescence & floral sporophyte | OsTLP10\|OsTLP10a\|OsTLP10b | Os05g0514300 | LOC_Os05g43850 |
| F-box | inflorescence & floral sporophyte | OsTLP12 | Os01g0759100 | LOC_Os01g55430 |
| F-box | inflorescence & floral sporophyte | OsTLP6 | Os11g0163600 | LOC_Os11g06420 |
| F-box | inflorescence & floral sporophyte | OsWD40-110 | Os04g0619700 | LOC_Os04g52870 |
| F-box | inflorescence & floral sporophyte | OsWD40-184 | Os11g0182400 | LOC_Os11g07970 |
| RING | pollen | APIP6 | Os05g0154600 | LOC_Os05g06270 |
| RING | pollen | DCA1 | Os10g0456800 | LOC_Os10g31850 |
| RING | pollen | MKRN | Os06g0318700 | LOC_Os06g21390 |
| RING | pollen | OsCLR1\|DSNP1 | Os06g0633500 | LOC_Os06g42700 |
| RING | pollen | OsDSG1 | Os09g0434200 | LOC_Os09g26400 |
| RING | pollen | OsRDCP1\|OsREIW1 | Os04g0530500 | LOC_Os04g44820 |
| RING | pollen | OsSDIR1 | Os03g0272300 | LOC_Os03g16570 |
| RING | pollen | SDEL1\|OsMSRFP | Os12g0538500 | LOC_Os12g35320 |
| RING | pollen | CHR708 | Os01g0952200 | LOC_Os01g72310 |
| RING | pollen | OsRFPH2-11 | Os10g0406200 | LOC_Os10g26610 |
| RING | pollen | OsRFPH2-19 | Os03g0788800 | LOC_Os03g57500 |
| RING | pollen | OsRFPH2-7 | Os06g0231600 | LOC_Os06g12560 |
| RING | pollen | OsRFPHC-12 | Os04g0512400 | LOC_Os04g43300 |
| RING | pollen | OsRFPHC-14 | Os01g0234900 | LOC_Os01g13370 |
| RING | pollen | OsSTA11 | Os01g0276600 | LOC_Os01g16950 |
| RING | pollen | OsSTA5 | Os01g0178700 | LOC_Os01g08340 |
| RING | pollen | OsSTA9 | Os01g0234900 | LOC_Os01g13370 |
| RING | pollen | XBOS34 | Os07g0446100 | LOC_Os07g26490 |
| RING | pollen | ZOS10-06 | Os10g0494500 | LOC_Os10g35190 |
| F-box | male gametophytic lineage | GIC | Os04g0675800 | LOC_Os04g57920 |
| F-box | male gametophytic lineage | MOF | Os04g0464966 | LOC_Os04g39080 |
| F-box | male gametophytic lineage | OsDRF1 | Os04g0431200 | LOC_Os04g35190 |
| F-box | male gametophytic lineage | OsLKP2 | Os02g0150800 | LOC_Os02g05700 |
| F-box | male gametophytic lineage | CHR708 | Os01g0952200 | LOC_Os01g72310 |
| F-box | male gametophytic lineage | OsJMJ-C8 | Os03g0389900 | LOC_Os03g27250 |
| F-box | male gametophytic lineage | OsSTA12 | Os01g0281000 | LOC_Os01g17390 |
| F-box | male gametophytic lineage | OsSTA265 | Os11g0593600 | LOC_Os11g38130 |
| F-box | male gametophytic lineage | OsTLP10\|OsTLP10a\|OsTLP10b | Os05g0514300 | LOC_Os05g43850 |
| F-box | male gametophytic lineage | OsTLP12 | Os01g0759100 | LOC_Os01g55430 |
| F-box | male gametophytic lineage | OsTLP6 | Os11g0163600 | LOC_Os11g06420 |
| F-box | male gametophytic lineage | OsWD40-184 | Os11g0182400 | LOC_Os11g07970 |
| RING | anther developmental series | ALT1 | Os01g0779400 | LOC_Os01g57110 |
| RING | anther developmental series | APIP6 | Os05g0154600 | LOC_Os05g06270 |
| RING | anther developmental series | BRHIS1 | Os08g0180300 | LOC_Os08g08220 |
| RING | anther developmental series | DCA1 | Os10g0456800 | LOC_Os10g31850 |
| RING | anther developmental series | DHS | Os02g0682300 | LOC_Os02g45780 |
| RING | anther developmental series | EBR1 | Os05g0279400 | LOC_Os05g19970 |
| RING | anther developmental series | HEI10 | Os02g0232100 | LOC_Os02g13810 |
| RING | anther developmental series | MKRN | Os06g0318700 | LOC_Os06g21390 |
| RING | anther developmental series | OsCLR1\|DSNP1 | Os06g0633500 | LOC_Os06g42700 |
| RING | anther developmental series | OsCOIN | Os01g0104100 | LOC_Os01g01420 |
| RING | anther developmental series | OsDIRP1 | Os06g0687200 | LOC_Os06g47270 |
| RING | anther developmental series | OsDSG1 | Os09g0434200 | LOC_Os09g26400 |
| RING | anther developmental series | OsHIRP1 | Os03g0302200 | LOC_Os03g19020 |
| RING | anther developmental series | OsHTAS | Os09g0323100 | LOC_Os09g15430 |
| RING | anther developmental series | OsHUB1\|MIP1 | Os04g0550400 | LOC_Os04g46450 |
| RING | anther developmental series | OsMAR1 | Os06g0695600 | LOC_Os06g48040 |
| RING | anther developmental series | OsNLA2 | Os03g0650900 | LOC_Os03g44810 |
| RING | anther developmental series | OsPIE1 | Os01g0954400 | LOC_Os01g72480 |
| RING | anther developmental series | OsRDCP1\|OsREIW1 | Os04g0530500 | LOC_Os04g44820 |
| RING | anther developmental series | OsRFPH2-10 | Os08g0539300 | LOC_Os08g42640 |
| RING | anther developmental series | OsRING-1\|OsATL38 | Os02g0759400 | LOC_Os02g52210 |
| RING | anther developmental series | OsRINGC2-2 | Os12g0102400 | LOC_Os12g01190 |
| RING | anther developmental series | OsSDIR1 | Os03g0272300 | LOC_Os03g16570 |
| RING | anther developmental series | OsSIRP2 | Os03g0798200 | LOC_Os03g58390 |
| RING | anther developmental series | SDEL1\|OsMSRFP | Os12g0538500 | LOC_Os12g35320 |
| RING | anther developmental series | UCIP2 | Os01g0667700 | LOC_Os01g47740 |
| RING | anther developmental series | AM29 | Os06g0535900 | LOC_Os06g34470 |
| RING | anther developmental series | CHR708 | Os01g0952200 | LOC_Os01g72310 |
| RING | anther developmental series | CHR715 | Os04g0629300 | LOC_Os04g53720 |
| RING | anther developmental series | CHR731 | Os07g0511500 | LOC_Os07g32730 |
| RING | anther developmental series | OsRFPH2-11 | Os10g0406200 | LOC_Os10g26610 |
| RING | anther developmental series | OsRFPH2-12 | Os01g0802000 | LOC_Os01g58780 |
| RING | anther developmental series | OsRFPH2-19 | Os03g0788800 | LOC_Os03g57500 |
| RING | anther developmental series | OsRFPH2-23 | Os01g0692700 | LOC_Os01g49770 |
| RING | anther developmental series | OsRFPH2-7 | Os06g0231600 | LOC_Os06g12560 |
| RING | anther developmental series | OsRFPH2-9 | Os03g0142500 | LOC_Os03g04890 |
| RING | anther developmental series | OsRFPHC-12 | Os04g0512400 | LOC_Os04g43300 |
| RING | anther developmental series | OsRFPHC-14 | Os01g0234900 | LOC_Os01g13370 |
| RING | anther developmental series | OsRFPHC-6 | Os07g0626900 | LOC_Os07g43380 |
| RING | anther developmental series | OsRFPHC-9 | Os02g0128800 | LOC_Os02g03620 |
| RING | anther developmental series | OsRFPV-2 | Os05g0355300 | LOC_Os05g28730 |
| RING | anther developmental series | OsRFPV-4 | Os01g0610700 | LOC_Os01g42500 |
| RING | anther developmental series | OsSTA11 | Os01g0276600 | LOC_Os01g16950 |
| RING | anther developmental series | OsSTA5 | Os01g0178700 | LOC_Os01g08340 |
| RING | anther developmental series | OsSTA9 | Os01g0234900 | LOC_Os01g13370 |
| RING | anther developmental series | OsZFP96 | Os11g0578200\|Os11g0578225\|Os11g0578250 | LOC_Os11g36970 |
| RING | anther developmental series | XBOS34 | Os07g0446100 | LOC_Os07g26490 |
| RING | anther developmental series | ZOS10-06 | Os10g0494500 | LOC_Os10g35190 |
| RING | inflorescence & floral sporophyte | ALT1 | Os01g0779400 | LOC_Os01g57110 |
| RING | inflorescence & floral sporophyte | APIP6 | Os05g0154600 | LOC_Os05g06270 |
| RING | inflorescence & floral sporophyte | BRHIS1 | Os08g0180300 | LOC_Os08g08220 |
| RING | inflorescence & floral sporophyte | DCA1 | Os10g0456800 | LOC_Os10g31850 |
| RING | inflorescence & floral sporophyte | DHS | Os02g0682300 | LOC_Os02g45780 |
| RING | inflorescence & floral sporophyte | EBR1 | Os05g0279400 | LOC_Os05g19970 |
| RING | inflorescence & floral sporophyte | EL5 | Os02g0559800 | LOC_Os02g35329 |
| RING | inflorescence & floral sporophyte | EL5.6 | Os02g0561800 | LOC_Os02g35347 |
| RING | inflorescence & floral sporophyte | HEI10 | Os02g0232100 | LOC_Os02g13810 |
| RING | inflorescence & floral sporophyte | MKRN | Os06g0318700 | LOC_Os06g21390 |
| RING | inflorescence & floral sporophyte | OsCesA7 | Os10g0467800 | LOC_Os10g32980 |
| RING | inflorescence & floral sporophyte | OsCLR1\|DSNP1 | Os06g0633500 | LOC_Os06g42700 |
| RING | inflorescence & floral sporophyte | OsCOIN | Os01g0104100 | LOC_Os01g01420 |
| RING | inflorescence & floral sporophyte | OsDIRP1 | Os06g0687200 | LOC_Os06g47270 |
| RING | inflorescence & floral sporophyte | OsDSG1 | Os09g0434200 | LOC_Os09g26400 |
| RING | inflorescence & floral sporophyte | OsHIRP1 | Os03g0302200 | LOC_Os03g19020 |
| RING | inflorescence & floral sporophyte | OsHTAS | Os09g0323100 | LOC_Os09g15430 |
| RING | inflorescence & floral sporophyte | OsHUB1\|MIP1 | Os04g0550400 | LOC_Os04g46450 |
| RING | inflorescence & floral sporophyte | OsMAR1 | Os06g0695600 | LOC_Os06g48040 |
| RING | inflorescence & floral sporophyte | OsNLA2 | Os03g0650900 | LOC_Os03g44810 |
| RING | inflorescence & floral sporophyte | OsPIE1 | Os01g0954400 | LOC_Os01g72480 |
| RING | inflorescence & floral sporophyte | OsRDCP1\|OsREIW1 | Os04g0530500 | LOC_Os04g44820 |
| RING | inflorescence & floral sporophyte | OsRFPH2-10 | Os08g0539300 | LOC_Os08g42640 |
| RING | inflorescence & floral sporophyte | OsRING-1\|OsATL38 | Os02g0759400 | LOC_Os02g52210 |
| RING | inflorescence & floral sporophyte | OsRINGC2-2 | Os12g0102400 | LOC_Os12g01190 |
| RING | inflorescence & floral sporophyte | OsSDIR1 | Os03g0272300 | LOC_Os03g16570 |
| RING | inflorescence & floral sporophyte | OsSIRP2 | Os03g0798200 | LOC_Os03g58390 |
| RING | inflorescence & floral sporophyte | SDEL1\|OsMSRFP | Os12g0538500 | LOC_Os12g35320 |
| RING | inflorescence & floral sporophyte | UCIP2 | Os01g0667700 | LOC_Os01g47740 |
| RING | inflorescence & floral sporophyte | AM29 | Os06g0535900 | LOC_Os06g34470 |
| RING | inflorescence & floral sporophyte | CHR708 | Os01g0952200 | LOC_Os01g72310 |
| RING | inflorescence & floral sporophyte | CHR710 | Os02g0527100 | LOC_Os02g32570 |
| RING | inflorescence & floral sporophyte | CHR715 | Os04g0629300 | LOC_Os04g53720 |
| RING | inflorescence & floral sporophyte | CHR731 | Os07g0511500 | LOC_Os07g32730 |
| RING | inflorescence & floral sporophyte | OsRFPH2-11 | Os10g0406200 | LOC_Os10g26610 |
| RING | inflorescence & floral sporophyte | OsRFPH2-12 | Os01g0802000 | LOC_Os01g58780 |
| RING | inflorescence & floral sporophyte | OsRFPH2-19 | Os03g0788800 | LOC_Os03g57500 |
| RING | inflorescence & floral sporophyte | OsRFPH2-23 | Os01g0692700 | LOC_Os01g49770 |
| RING | inflorescence & floral sporophyte | OsRFPH2-7 | Os06g0231600 | LOC_Os06g12560 |
| RING | inflorescence & floral sporophyte | OsRFPH2-9 | Os03g0142500 | LOC_Os03g04890 |
| RING | inflorescence & floral sporophyte | OsRFPHC-12 | Os04g0512400 | LOC_Os04g43300 |
| RING | inflorescence & floral sporophyte | OsRFPHC-14 | Os01g0234900 | LOC_Os01g13370 |
| RING | inflorescence & floral sporophyte | OsRFPHC-6 | Os07g0626900 | LOC_Os07g43380 |
| RING | inflorescence & floral sporophyte | OsRFPHC-9 | Os02g0128800 | LOC_Os02g03620 |
| RING | inflorescence & floral sporophyte | OsRFPV-2 | Os05g0355300 | LOC_Os05g28730 |
| RING | inflorescence & floral sporophyte | OsRFPV-4 | Os01g0610700 | LOC_Os01g42500 |
| RING | inflorescence & floral sporophyte | OsSTA11 | Os01g0276600 | LOC_Os01g16950 |
| RING | inflorescence & floral sporophyte | OsSTA5 | Os01g0178700 | LOC_Os01g08340 |
| RING | inflorescence & floral sporophyte | OsSTA9 | Os01g0234900 | LOC_Os01g13370 |
| RING | inflorescence & floral sporophyte | OsZFP96 | Os11g0578200\|Os11g0578225\|Os11g0578250 | LOC_Os11g36970 |
| RING | inflorescence & floral sporophyte | XBOS34 | Os07g0446100 | LOC_Os07g26490 |
| RING | inflorescence & floral sporophyte | ZOS10-06 | Os10g0494500 | LOC_Os10g35190 |
