## Supplementary material for "RE3DB: A multi-omics phylogenomics platform for rice E3 ubiquitin ligases identifies novel regulators of pollen germination": Table S10

| Locus id | rupo | osmtd2 | gori | madstri | tape | osrac6 | ralf17/19 | abcg16/28 | gtrd5 | Group |
| --- | --- | --- | --- | --- | --- | --- | --- | --- | --- | --- |
| LOC_Os04g40770 |  |  |  |  |  |  |  | O |  | AD |
| LOC_Os02g41930 |  |  |  | O |  |  |  |  |  | MD |
| LOC_Os01g71440 |  |  |  |  |  |  |  | O |  | AD |
| LOC_Os06g35080 |  |  |  |  |  |  | O |  |  | RAD |
| LOC_Os10g10420 | O |  | O | O |  |  |  |  |  | RD/GD/MD |
| LOC_Os01g06360 | O |  |  | O | O |  |  |  |  | RD/MD/TD |
| LOC_Os07g09110 |  |  |  | O |  |  |  |  |  | MD |
| LOC_Os03g25240 | O |  |  | O |  | O |  |  |  | RD/MD/R6D |
| LOC_Os01g17390 |  |  |  | O |  |  |  |  |  | MD |
| LOC_Os04g31540 |  |  |  |  |  |  |  |  | O | G5D |
| LOC_Os09g15460 |  |  |  |  |  |  |  | O |  | AD |
| LOC_Os12g27790 |  |  |  |  |  |  | O |  |  | RAD |
| LOC_Os07g36520 |  |  |  |  |  |  |  |  | O | G5D |
| LOC_Os03g25650 |  |  |  | O |  |  |  |  |  | MD |
| LOC_Os03g25190 | O |  |  |  |  |  |  |  |  | RD |
| LOC_Os01g56750 |  |  |  | O |  |  |  |  |  | MD |
| LOC_Os09g17190 | O |  |  |  | O |  |  |  |  | RD/TD |
| LOC_Os10g34340 |  |  | O |  |  |  |  |  |  | GD |
| LOC_Os04g40960 | O |  |  | O |  |  |  | O | O | RD/MD/AD/G5D |
| LOC_Os08g36960 |  |  |  |  |  |  |  | O | O | AD/G5D |
| LOC_Os04g26240 |  |  |  |  |  |  |  |  | O | G5D |
| LOC_Os02g17210 |  |  |  | O |  |  |  |  |  | MD |
| LOC_Os03g47420 |  |  |  |  |  |  |  |  | O | G5D |
| LOC_Os11g36350 |  | O |  |  | O |  |  |  |  | M2D/TD |
| LOC_Os03g46500 |  |  |  |  |  |  |  |  | O | G5D |
| LOC_Os07g16800 |  |  |  |  |  |  |  |  | O | G5D |
| LOC_Os09g17152 |  |  | O |  |  |  |  |  |  | GD |
| LOC_Os03g46140 | O |  |  |  |  |  |  |  |  | RD |
| LOC_Os02g54240 |  |  |  |  |  |  |  |  | O | G5D |
| LOC_Os11g42310 | O |  |  |  |  |  |  |  |  | RD |
| LOC_Os12g34290 |  |  |  |  |  | O |  |  | O | R6D/G5D |
| LOC_Os11g38180 |  |  |  |  |  | O | O |  |  | R6D/RAD |
| LOC_Os05g51100 | O |  |  |  |  |  | O |  |  | RD/RAD |
| LOC_Os12g40140 |  |  |  | O |  |  |  |  |  | MD |
| LOC_Os01g40160 | O |  |  |  |  | O |  |  |  | RD/R6D |
| LOC_Os11g38100 | O |  |  |  |  |  |  |  |  | RD |
| LOC_Os07g03090 |  |  |  |  |  |  |  |  | O | G5D |
| LOC_Os11g42280 | O |  |  |  |  |  |  |  |  | RD |
| LOC_Os11g42270 |  |  |  |  |  |  |  | O |  | AD |
| LOC_Os04g40800 |  |  | O | O |  | O |  |  |  | GD/MD/R6D |
| LOC_Os04g35190 |  |  |  | O |  |  |  |  |  | MD |
| LOC_Os05g49540 | O |  | O | O |  | O |  |  |  | RD/GD/MD/R6D |
| LOC_Os03g56440 |  |  |  |  |  |  |  |  | O | G5D |
| LOC_Os05g02550 |  |  |  |  |  |  |  |  | O | G5D |
| LOC_Os05g01620 |  |  |  | O |  |  |  |  |  | MD |
| LOC_Os02g33840 | O |  |  | O |  |  |  |  |  | RD/MD |
| LOC_Os03g56450 |  |  | O |  |  |  |  |  |  | GD |
| LOC_Os03g25220 | O |  |  |  |  |  |  |  |  | RD |
| LOC_Os07g36280 |  |  | O |  | O |  |  |  |  | GD/TD |
