## Supplementary material for "RE3DB: A multi-omics phylogenomics platform for rice E3 ubiquitin ligases identifies novel regulators of pollen germination": Table S11

| **Upstream genes** | **Downstream genes** | **Regulatory relationship** | **Evidence** | **Experimental validation** | **Sources** |
| --- | --- | --- | --- | --- | --- |
| OsMADS62/63/68 | RUPO | Downregulation | Downregulation shown in the RNA-Seq of OsMADS triple knockout mutant | No | 10.3390/ijms23010239 10.1111/jipb.13900 |
| OsMADS62/63/68 | TAPE | Downregulation | Downregulation shown in the RNA-Seq of OsMADS triple knockout mutant | No | 10.1111/jipb.13900 |
| OsMADS62/63/68 | OsMTD2 | Downregulation | Downregulation shown in the RNA-Seq of OsMADS triple knockout mutant | No | 10.1111/jipb.13900 |
| OsMADS62/63/68 | GORI | Downregulation | Downregulation shown in the RNA-Seq of OsMADS triple knockout mutant | No | 10.1111/jipb.13900 |
| OsMADS62/63/68 | OsRALF17/19 | Downregulation | Downregulation shown in the RNA-Seq of OsMADS triple knockout mutant | No | 10.1111/jipb.13900 |
| OsMADS62/63/68 | OsABCG16/28 | Downregulation | Downregulation shown in the RNA-Seq of OsMADS triple knockout mutant | No | 10.1111/jipb.13900 |
| OsMADS62/63/68 | GTrD5 | Downregulation | Downregulation shown in the RNA-Seq of OsMADS triple knockout mutant | No | 10.1111/jipb.13900 |
| RUPO | GORI | Downregulation | Downregulation shown in the RNA-Seq of RUPO knockout mutant | No | 10.1111/jipb.13900 |
| OsMADS62/68 | RUPO | Negative feedback | Reduced OsMADS62/68 expression in knockout mutant of RUPO | No | 10.1111/jipb.13900 |
| RUPO | TAPE | Downregulation | Downregulation shown in the RNA-Seq of RUPO knockout mutant | No | 10.1111/jipb.13900 |
| RUPO | OsRac6 | Downregulation | Downregulation shown in the RNA-Seq of RUPO knockout mutant | No | 10.1111/jipb.13900 |
| OsRALF19 | RUPO | Negative feedback | Reduced OsRALF19 expression in knockout mutant of RUPO | No | 10.1111/jipb.13900 |
| RUPO | OsABCG16/28 | Downregulation | Downregulation shown in the RNA-Seq of RUPO knockout mutant | No | 10.1111/jipb.13900 |
| RUPO | GTrD5 | Downregulation | Downregulation shown in the RNA-Seq of RUPO knockout mutant | No | 10.1111/jipb.13900 |
| OsMADS63 | OsMTD2 | Negative feedback | Reduced OsMADS63 expression in knockout mutant of OsMTD2 | No | 10.1111/jipb.13900 |
| OsMADS63 | TAPE | Negative feedback | Reduced OsMADS63 expression in knockout mutant of TAPE | No | 10.1111/jipb.13900 |
| RUPO | OsRac6 | Negative feedback | Reduced OsMADS63 expression in knockout mutant of OsRac6 | No | 10.1007/s12374-023-09403-7 10.1111/jipb.13900 |
| OsRALF17/19 | OsMTD2 | Protein interaction | Co-immunoprecipitation, bimolecular fluorescence complementation, and yeast two-hybrid assays revealed that mature peptides OsRALF17 and OsRALF19 directly bind the extracellular domains of both OsMTD2 and RUPO forming a heteromeric receptor complex | Yes | 10.1016/j.jplph.2025.154421 10.1111/jipb.13508 |
| OsRALF17/19 | RUPO | Protein interaction | Co-immunoprecipitation, bimolecular fluorescence complementation, and yeast two-hybrid assays revealed that mature peptides OsRALF17 and OsRALF19 directly bind the extracellular domains of both OsMTD2 and RUPO forming a heteromeric receptor complex | Yes | 10.1016/j.jplph.2025.154421 10.1111/jipb.13508 |
| GORI | OsRac6 | Downregulation | Co-immunoprecipitation reveals direct protein interaction, and over-expression crossing lines shows that extra GORI dampens Rac6-driven hyper-elongation | Yes | 10.1080/15592324.2022.2082678 |
